# Efficient cell-free evolution of RNA polymerases by droplet microfluidics

**DOI:** 10.64898/2026.02.23.707346

**Authors:** Naohiro Terasaka, Taro Furubayashi, Kenya Tajima, Hiroyuki Noji

## Abstract

Directed evolution enables the rapid creation of biomolecules with new functions, yet linking genotype and phenotype often requires laborious cell-based workflows. Here we integrate a boosted *in vitro* transcription–translation system capable of protein expression from single-copy DNA with ultrahigh-throughput fluorescence-activated droplet sorting, developing a simple one-day workflow for efficient protein evolution. Using this system, we evolved SP6 RNA polymerase into a highly robust variant that functions in multiple environments, including cell-free reactions and mammalian cells. We further engineered the evolved enzyme into a proximity-dependent split RNA polymerase that converts molecular interactions into transcriptional activity with minimal background. The resulting biosensors detect diverse molecular targets *in vitro*, including proteins, peptides, RNAs, and molecular glues. This cell-free platform provides versatile routes for evolving diverse functional biomolecules, accelerating the development of advanced biotechnologies such as diagnostics and therapeutics.

## Introduction

Directed evolution, which emulates natural selection in laboratory settings, is a powerful technology for creating functional biomolecules with novel activities, such as enzymes, artificial antibodies, and biosensors^1^. The success of directed evolution depends on the size of accessible sequence space and efficiency of screening variants with desired functions. In recent years, droplet microfluidics has enabled ultrahigh-throughput screening, allowing for the compartmentalization and analysis of millions of individual genes and their protein products^2^. This approach, often coupled with fluorescence-activated droplet sorting (FADS), has been successfully used to evolve a wide array of molecules such as enzymes^3^, antibodies^4^, and RNA polymerases^5^.

Despite these remarkable successes, conventional FADS platforms that rely on living cells suffer from several inherent limitations. The typical workflow involves multiple steps, including plasmid library preparation, transformation of cells, culturing cells to express proteins, encapsulation of these cells with substrates and lysis reagents, incubation, sorting, recovery of genes from sorted droplets, and finally amplification of DNA for the next round^2^. This cell-based approach hampers the use of cytotoxic proteins and risks loss of library diversity due to transformation inefficiencies, thereby requiring time-consuming cell culturing steps.

To circumvent these constraints in FADS, *in vitro* transcription-translation (IVTT) systems have emerged as a powerful alternative for expressing proteins in droplets^6,7^. In principle, combining IVTT with FADS provides a direct link between genotype and phenotype in a droplet, enabling rapid and simplified screening. For accurate single-variant assessment, the encapsulation process should follow Poisson statistics, ensuring that most droplets contain one or zero DNA molecules. This requirement typically results in an effective DNA concentration in the several hundred femtomolar (fM) range, which is approximately 10,000 times lower than the nanomolar (nM) concentrations typically used in IVTT systems^8^.

Several strategies have been developed to overcome this low-concentration barrier, such as coupling IVTT with isothermal amplification methods like rolling circle amplification (RCA)^9^, single-template amplification on magnetic beads (BEAMing)^10^ or multi-step reagent exchange using functional agarose beads^11^. However, these techniques often require multiple experimental steps, specialized instrumentation such as pico-injection^12^, or impose limits on the length of the available DNA^13^. In our recent work, we addressed this fundamental hurdle by engineering the IVTT system itself. We optimized the concentration of RNA polymerase and ribosome in a bacterial reconstituted IVTT system (PURE system) for sub-picomolar linear DNA templates, enabling robust protein expression from a single DNA molecule in a picoliter droplet without any prior DNA amplification, making it directly applicable to FADS platforms^8^. Building on our cell-free FADS platform, we next aimed to broaden the repertoire of orthogonal transcription systems and proximity-dependent split RNA polymerase (RNAP) biosensors for cell-free synthetic biology. While split T7 RNAP-based sensors enable modular detection schemes, their performance *in vitro* can be limited by background arising from spontaneous fragment reassembly, often necessitating protein engineering and assay tuning to maximize signal-to-noise^14–16^. We chose SP6 RNA polymerase (SP6 RNAP) as a well-characterized single-subunit phage polymerase that recognizes a distinct promoter, offering inherent orthogonality, yet whose broader use has been limited by pronounced salt sensitivity^17^. In this study, we first evolved SP6 RNAP into a highly robust variant that functions across cell-free systems and mammalian cells. We then demonstrate the utility of our platform by engineering the evolved enzyme into an orthogonal, proximity-dependent split SP6 RNAP to provide an orthogonal transcription module that complements the T7 RNAP-based toolbox for cell-free genetic circuits. Cell-free evolution markedly improved the signal-to-noise ratio of this biosensor, enabling the detection of diverse molecular interactions, including those involving peptides, antibodies, RNA, and small molecules.

## Results

### A single-molecule IVTT platform enables robust SP6 RNAP activity detection in picoliter droplets

To enable the evolution of SP6 RNAP, we first established a FADS-compatible assay capable of detecting its activity from a single DNA template. Building on our previously developed boosted cell-free gene expression system optimized for sub-picomolar DNA concentrations (hereafter b-PURE)^8^, we designed two DNA constructs: a gene encoding SP6 RNAP under the control of a T7 promoter, and a reporter gene encoding the green fluorescent protein mNeonGreen (mNG) under an SP6 promoter^18^ (Fig. 1a). The activity of expressed SP6 RNAP was evaluated by monitoring mNG fluorescence when these two linear double-stranded DNA (dsDNA) templates were introduced into the b-PURE system. Initially, b-PURE reaction components were optimized to improve the expression of SP6 RNAP at a DNA concentration of 200 fM in a test tube, mimicking the conditions of a single DNA copy in a droplet with an approximate diameter of 25 µm. We found that the addition of chaperone proteins (1.25 μM of DnaK, 0.25 μM of DnaJ, and 0.25 μM of GrpE) and 0.25 μM of elongation factor-P (EF-P) enhanced SP6 RNAP expression (Extended Data Fig. 1a). To further improve the transcription by SP6 RNAP, we hypothesized that increasing the affinity of the RNAP for the DNA template should be beneficial in preventing the RNAP from dissociating during transcription. Inspired by previous work showing that DNA polymerase processivity can be improved by fusion to a non-specific dsDNA-binding protein, we fused the Sso7d protein to the N-terminus of SP6 RNAP via linkers of varying lengths^19^. IVTT experiments showed that a 41-amino-acid linker was optimal for enhancing SP6 RNAP activity (Extended Data Fig. 1b).

**Fig. 1:**
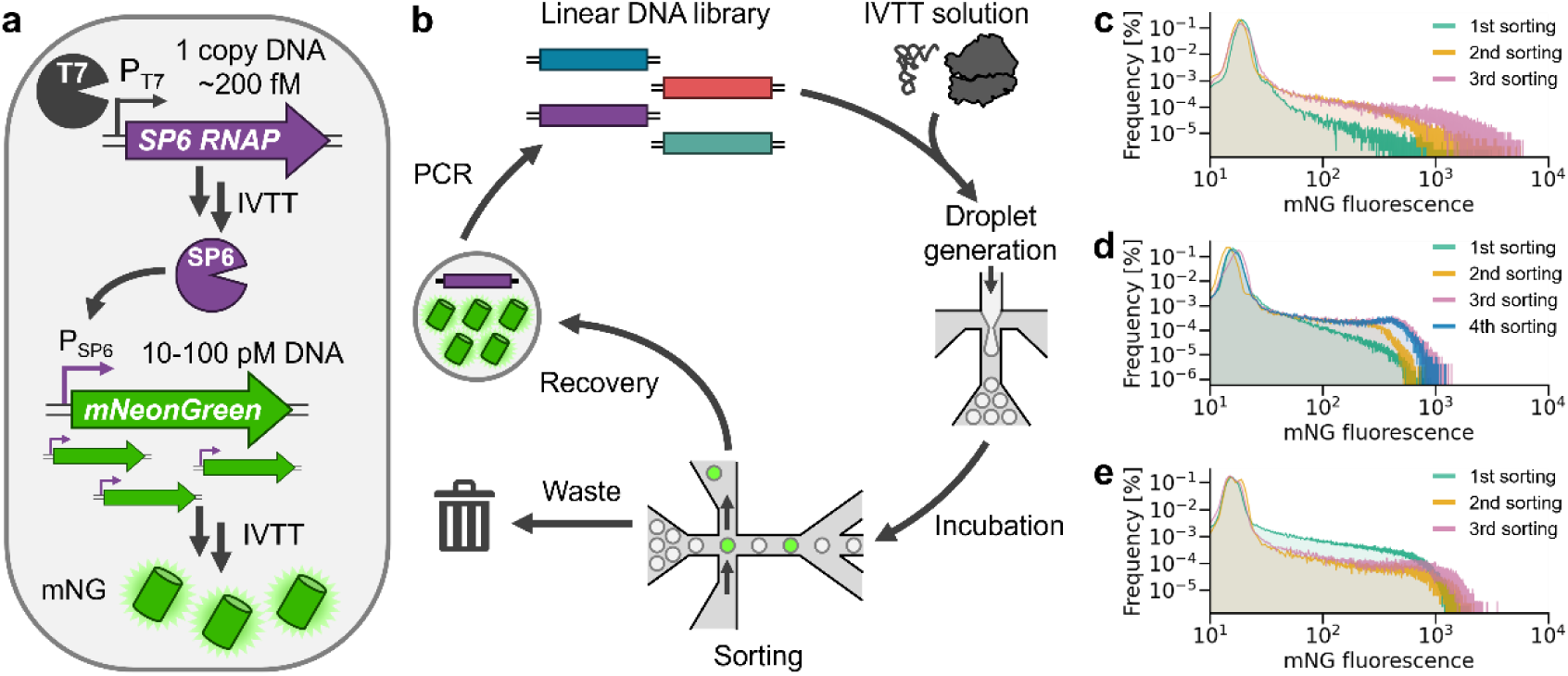
Screening of SP6 RNAP expressed from single molecule DNA in water-in-oil emulsion. **(a)** Schematic illustration of the detection scheme for SP6 RNA polymerase activity expressed from single-copy DNA in water-in-oil emulsion. **(b)** Schematic illustration of cell-free fluorescence-activated droplet sorting (FADS) platform. **(c-e)** Histograms of the fluorescence signal distribution of droplets screened for SP6 RNA polymerase activity in FADS. In each round of directed evolution, sorting was repeated 3-4 times. The histograms corresponding to each sorting are colored in green (1st), orange (2nd), pink (3rd), blue (4th). The directed evolution using Library 1 is shown in (**c**), Library 2 is in (**d**) and Library 3 is in (**e**). The individual histograms are shown in Extended Data Figs 2 and 3.

### Directed evolution of SP6 RNAP using cell-free FADS

We then proceeded with the directed evolution of SP6 RNAP to enhance its activity in the PURE solution (Fig. 1b). Although the detailed structure of SP6 RNAP has not been experimentally determined, ColabFold-predicted models^20^ indicate that SP6 RNAP adopts an overall structure similar to the elongation state of T7 RNAP, with both RNAPs belonging to the bacteriophage single-subunit RNA polymerase family^21^ (Extended Data Fig. 1c-f). Transcription by T7 RNAP involves three distinct stages: initiation, elongation, and termination. The transition from the initiation to the elongation state requires a rearrangement of the N-terminal domain (NTD). Mutations in the NTD that facilitate this transition have been shown to improve T7 RNAP activity^22^. Inspired by this work, we constructed a dsDNA library by randomly introducing mutations into the NTD (Gln2–Ala240) of the Sso7d-fused SP6 RNAP using error-prone PCR (epPCR) (Library 1). The Sso7d-SP6 RNAP DNA library (20 fM) and the mNG reporter DNA (80 pM) were mixed with the b-PURE system to generate water-in-oil droplets containing, on average, a single DNA molecule encoding an SP6 RNAP mutant per ∼10 droplets (λ = 0.1) and ∼400 copies of DNA encoding mNG per droplet. The diameter of droplets was approximately 25 µm and the concentration of single-copy DNA in this droplet was ∼200 fM. After off-chip incubation at 37 °C for 18 hours, droplets exhibiting the top 0.4–0.5% of green fluorescence intensity were sorted, and mRNA was extracted from the sorted droplets (Fig. 1c and Extended Data Fig. 2a). The recovered mRNA was reverse-transcribed and amplified by PCR, and the resulting DNA library was used for the next round of screening. After three such cycles, the final amplified DNA library (Library 1.3) was ligated into a plasmid vector for cloning. Randomly selected clones were sequenced, and the activity of these variants was evaluated by monitoring mNG fluorescence levels in the b-PURE system in test tubes (Extended Data Fig. 2b and c). Variant #15, which had eight amino acid mutations (I31N, S37T, N42S, S110R, K123E, E177G, E199V, and T208S), showed the highest activity, and we named it Sso7d-SP6 RNAP-v2. To further improve its activity, random mutations were introduced into the NTD of Sso7d-SP6 RNAP-v2 to create a second-generation library (Library 2), and the concentration of mNG reporter DNA was reduced from 80 pM to 8 pM (∼400 to ∼40 copies per droplet) during the second round of evolution. After four cycles, we obtained Library 2.4 (Fig. 1d and Extended Data Fig. 3a). Library 3 was then prepared by DNA shuffling of Libraries 1.3 and 2.4 to merge beneficial mutations, followed by three additional rounds of screening to produce Library 3.3 (Fig. 1e and Extended Data Fig. 3b). We randomly selected clones from Library 3.3 and analyzed their sequences (Extended Data Fig. 3c). At this point, we also checked the activities of the evolved variants with or without the Sso7d domain and found that the Sso7d fusion was beneficial for the initial template, but it was no longer required for the activity of the evolved variants (Extended Data Fig. 4a and b). Consequently, variant #20 from Library 3.3, lacking the Sso7d domain, was named SP6 RNAP-v3 and chosen for further study due to the ease of expression and purification. SP6 RNAP-v3 contained five additional mutations (A25T, K68R, S142F, V153A, and G210R) and two reversions to wild-type residues (G177E, S208T) (Fig. 2a and Extended Data Fig. 4c).

**Fig. 2:**
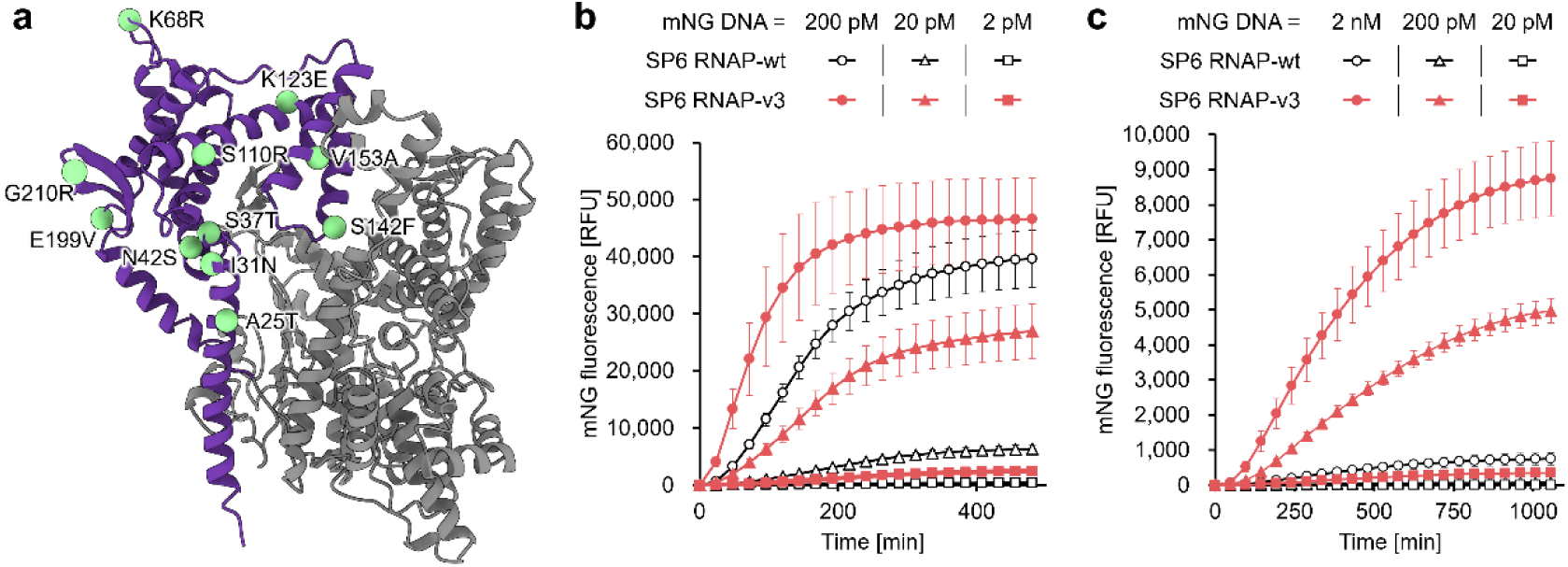
Directed evolution of SP6 RNA polymerase by cell-free FADS platform. **(a)** Mapping the mutations of SP6 RNAP-v3 onto the predicted structure of SP6 RNAP-wt (shown in light green spheres). The N-terminal domain where random mutations were introduced in the library preparation is colored purple. **(b, c)** The real-time fluorescence values of mNeonGreen (mNG) in the PURE system at 37 °C (**b**) or Wheat Germ system at 25 °C (**c**). DNA encoding mNG under SP6 promoter (2 nM, 200 pM, 20 pM or 2 pM) and purified SP6 RNAPs were mixed with PURE system and fluorescence values were monitored in a real-time PCR system. The data are presented as the mean ± SD from three independent experiments.

### Characterization of evolved SP6 RNA polymerase-v3

We next characterized the SP6 RNAP-v3. Wild-type SP6 RNAP and SP6 RNAP-v3 were expressed in *E. coli* and purified using immobilized metal affinity chromatography followed by anion-exchange chromatography. The purified polymerases were mixed with various concentrations of mNG reporter DNA in the PURE system, and their activities were evaluated by monitoring mNG fluorescence intensity in test tubes (Fig. 2b). Time-course measurements revealed that the purified SP6 RNAP-v3 exhibited higher activity than SP6 RNAP-wt, especially at low concentrations of mNG DNA (1.3-fold at 200 pM mNG DNA and 4.0-fold at 20 pM mNG DNA based on the amount of expressed mNG at the endpoint).

SP6 RNA polymerase is known to be sensitive to high salt concentrations^17^, a condition that is suboptimal for IVTT reactions, which typically contain high concentrations of anions such as glutamate^23^. We evaluated the transcriptional activity of the SP6 RNAP variants under different buffer conditions (Extended Data Fig. 5). Transcriptional activities were evaluated relative to the activity of SP6 RNAP-wt measured in a standard transcription buffer containing 6 mM magnesium chloride^24^. Under high-salt conditions (300 mM potassium glutamate and 20 mM magnesium acetate), which mimic the PURE system buffer^7,23^, SP6 RNAP-wt retained only 17.8% activity. In contrast, SP6 RNAP-v3 exhibited enhanced activity under the high-salt conditions, reaching 125% of the wild-type activity measured in the standard transcription buffer, but showed only 14.9% activity in the standard transcription buffer. These results indicate that SP6 RNAP-v3 was successfully optimized for the PURE system buffer conditions during the directed evolution campaign. To investigate the applicability of evolved SP6 RNAP-v3 in different IVTT system, we tested the activity of the SP6 RNAP variants in a wheat germ extract-based cell-free system, which typically contains ∼100 mM of potassium acetate and 2.7 mM magnesium acetate^25^ (Fig. 2c). In this system, the activity difference between SP6 RNAP-wt and -v3 was even more pronounced. At a 2 nM reporter DNA concentration, SP6 RNAP-v3 showed a 9.1-fold higher activity than SP6 RNAP-wt based on the amount of expressed mNG at the endpoint. These results demonstrate the robust and versatile activity of the evolved SP6 RNAP-v3 across various IVTT systems.

### Directed evolution of proximity-dependent split SP6 RNA polymerase

Split RNAPs function as biosensors by converting biomolecular interactions into transcriptional activity^14,26^. Their N- and C-terminal fragments assemble into an active polymerase only when fused interaction partners bring them into proximity. The laboratory-evolved split T7 RNA polymerase (N-29-1) has been utilized in a modular cell-free biosensing platform called TLISA^14,15^. To expand the cell-free toolbox, we sought to develop a proximity-dependent split SP6 RNAP (spSP6 RNAP) that is orthogonal to T7 RNAP. For such a biosensor to be effective, the fused proteins must not sterically hinder assembly, yet the fragments should not self-assemble in the absence of the target interaction to ensure a good signal-to-noise ratio. Inspired by the reported spT7 RNAP (N-29-1), we first tested eight potential split sites (S47, Q79, A129, H149, F168, N197, I227, R279) within SP6 RNAP-v3, selected because these residues are predicted to be solvent-exposed (Extended Data Fig. 6a-b). The rapamycin-induced dimerization domains (FRB and FKBP) were fused to the corresponding N-terminal (N-SP6 RNAP-v3-FRB) and C-terminal (FKBP-C-SP6 RNAP-v3) fragments. The rapamycin-dependent activities of these spSP6 RNAPs were evaluated in the b-PURE system. The results identified H149 and N197 as promising split sites with relatively low background signals (Extended Data Fig. 6c). We selected the N197 site (Rapa-spSP6-v3) for further study to ensure the resulting spSP6 RNAP would be orthogonal to existing spT7 RNAPs, which have been split at the site corresponding to residue H149 in SP6 RNAP^14^.

As our initial Rapa-spSP6-v3 design exhibited a poor signal-to-noise ratio (8.97 times) as a practical biosensor in cell-free translation systems, we proceeded with cell-free microfluidics-based directed evolution. A bicistronic DNA construct was designed to express both FKBP-C-SP6-v3 and N-SP6-v3-FRB under a single T7 promoter (Fig. 3a). Random mutations were introduced by epPCR into the N-terminal fragment (Library 4) or into residues 198-400 of the C-terminal fragment, which are close to the N-terminal fragment in the predicted structure (Library 5). These two libraries were screened in parallel using cell-free FADS platform, selecting highly fluorescent droplets in the presence of rapamycin (Fig. 3b-c, Extended Data Fig. 6d-e). After three cycles of screening, clones were arbitrarily chosen from each campaign for sequencing characterization. The evolved variants showed a 20- to 70-fold improvement in rapamycin-dependent activity recovery compared to the parent Rapa-spSP6-v3 (Extended Data Fig. 6f-i).

**Fig. 3:**
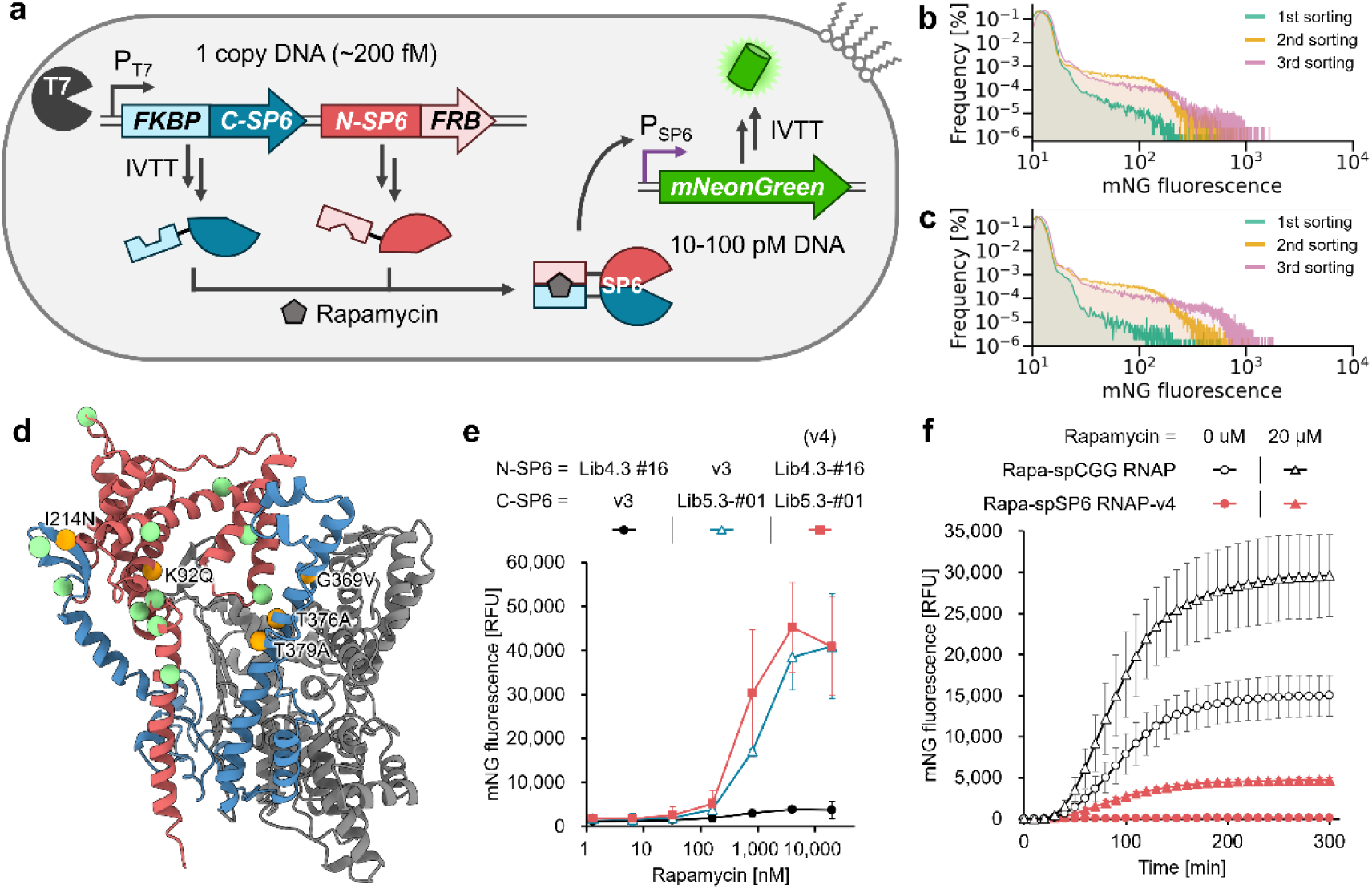
Directed evolution of proximity-dependent spSP6 RNAPs by cell-free FADS platform. **(a)** Schematic illustration of measuring the rapamycin-dependent activity of spSP6 RNA polymerases expressed by cell-free translation system from single-copy DNA in water-in-oil emulsion. **(b-c)** Histograms of the fluorescence signal distribution of droplets screened for Rapa-spSP6 RNA polymerase activity in fluorescence-activated droplet sorting (FADS). In each round of directed evolution, sorting was repeated three times. The histograms corresponding to each sorting are colored in green (1st), orange (2nd), pink (3rd). The directed evolution using Library 4 is shown in (**b**) and Library 5 is in (**c**). The individual histograms are shown in Extended Data Fig. 6. **(d)** Mapping the mutations of spSP6 RNAP-v4 onto the predicted structure of SP6 RNAP-wt. The N-terminal fragment of spSP6 RNAP is colored red and the region where random mutations were introduced in the C-terminal fragments is colored blue. The mutations from SP6 RNAP-v3 to spSP6 RNAP-v4 are shown in orange spheres. Light green spheres indicate the mutations in SP6 RNAP-v3 from SP6 RNAP-wt. **(e)** Endpoint measurement of rapamycin-induced mNG expression by spSP6 RNAPs in the PURE system. **(f)** Time course measurement of mNG fluorescence values in the PURE system by split RNAPs. The data are presented as the mean ± SD from three independent experiments.

We then evaluated the rapamycin dose-response of the promising variant from each library (Lib4.3-#16 and Lib5.3-#01) (Fig. 3e). These results identified the combination of N-SP6-Lib4.3-#16 (bearing the K92Q mutation) and C-SP6-Lib5.3-#01 (bearing I214N, G369V, T376A, and T379A mutations) as the optimal biosensor, which we named spSP6 RNAP-v4.

To be broadly applicable, proximity-dependent RNAPs require a high signal-to-noise ratio and orthogonality to other RNAP systems. However, a previously reported evolved split T7 RNA polymerase (spT7 RNAP) system suffers from substantial background activity caused by spontaneous fragment reassembly under IVTT conditions^15^. To compare the performance of our evolved spSP6 RNAP-v4 to the previously reported split RNAP in IVTT solution containing T7 RNAP, we employed a split CGG RNA polymerase (spCGG RNAP), which is orthogonal to the T7 RNAP used in the PURE system as a reference. The N-terminal fragment of spCGG RNAP is identical to that of evolved split T7 RNA polymerase (N-29-1), whereas its C-terminal fragment derives from engineered T7 RNA polymerase recognizing the “CGG” promoter^14,27^ (Extended Data Fig. 7a). For both systems, the split RNAP fragments were fused to FKBP and FRB and expressed under the control of a T7 promoter in the PURE system. While the spCGG RNAP exhibited substantial background activity and showed only 1.97-fold induction after 300 minutes, Rapa-spSP6 RNAP-v4 showed negligible activity in the absence of rapamycin and robust activation in time-course measurement (Fig. 3f). Although the absolute output of spSP6 RNAP-v4 was lower than that of spCGG RNAP, its signal-to-noise ratio reached 39.5-fold at 300 minutes. Endpoint analysis further demonstrated that spSP6 RNAP-v4 was highly orthogonal to CGG promoters (Extended Data Fig. 7b). Moreover, non-cognate pairs of spCGG RNAP fragments and spSP6 RNAP-v4 fragments showed no detectable activity, highlighting the potential of this system to construct multilayer genetic circuits^16^ and screen protein-protein interaction (PPI) networks^28^.

### Detection of various molecular interactions by evolved spSP6 RNAP-v4 *in vitro*

To showcase the breadth of applications enabled by the evolved proximity-dependent RNAPs, we next applied spSP6 RNAP-v4 as a general molecular interaction sensor capable of detecting diverse protein, peptide, RNA, and small-molecule interactions *in vitro*^29^. We first fused the oncoprotein human Aurora A kinase (hAurA) and its peptide ligand TPX2 (reported *K*_D_ = 5.4 μM) to the N- and C-terminal fragments of spSP6 RNAP-v4^30^ (Fig. 4a). In the PURE system, the cognate interaction between hAurA and TPX2 reconstituted polymerase activity, resulting in SP6 RNAP-dependent mNG expression (Fig. 4a). Consistent with a previous study, the hAurA(Y199K) mutant, which does not bind TPX2, showed a 6.5-fold lower signal, confirming the specificity of the sensor^30^. In a similar manner, anti-hAurA monobodies (Mb1 and Mb6) were fused to spSP6 RNAP-v4 and their specific binding to hAurA-wt were detected (Extended Data Fig. 8a and b).

**Fig. 4:**
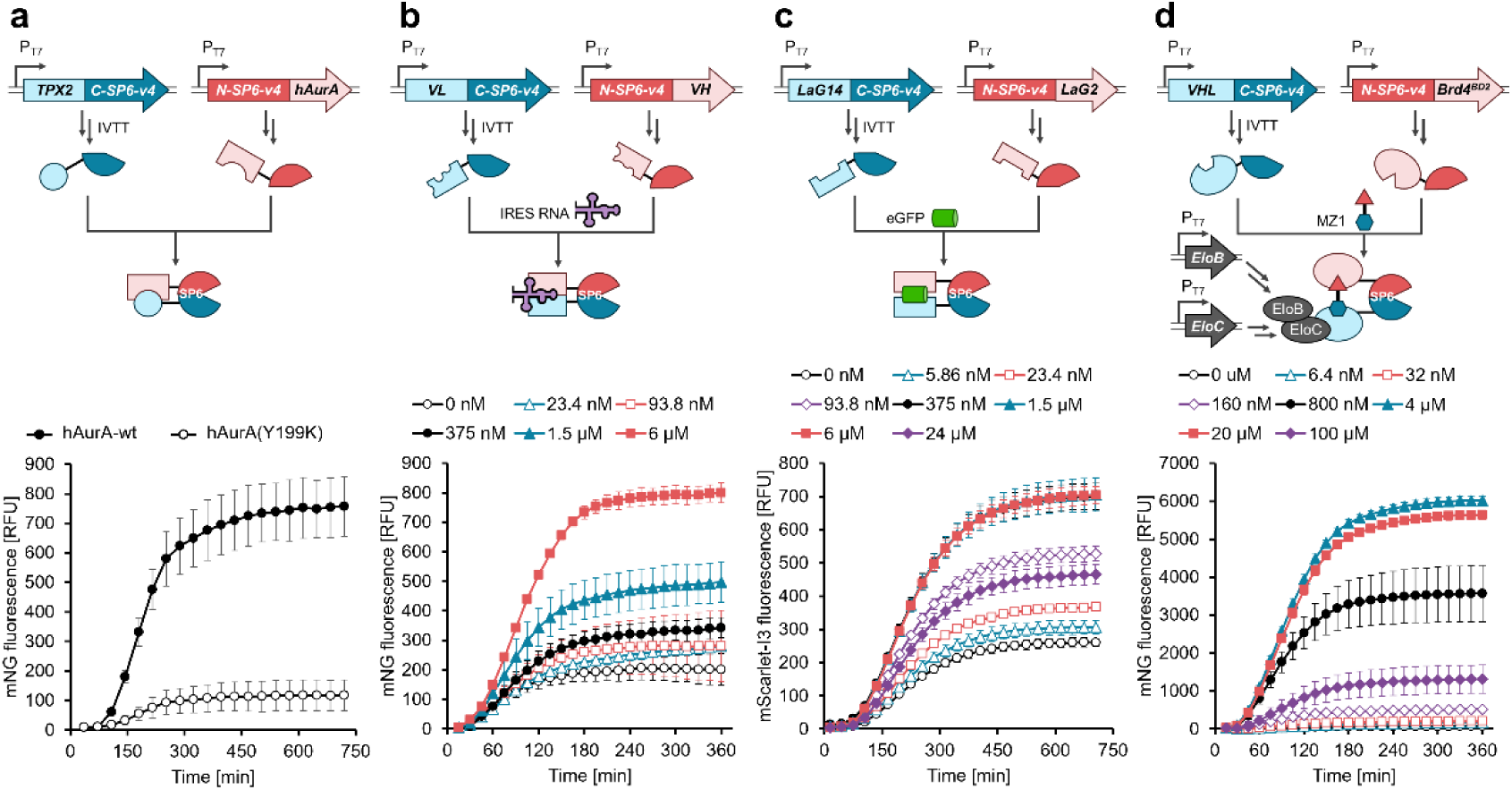
Cell-free detection of molecular interaction by evolved spSP6 RNAP-v4. Each protein is encoded on linear DNA under the T7 promoter. The interactions between molecules fused to the fragments of spSP6 RNAP-v4 activate RNAP. The expression level of reporter fluorescent protein under the SP6 promoter was monitored in a time-dependent manner. **(a)** Human Aurora A kinase (hAurA) or its negative mutant (Y199K) was fused to the N-terminal fragment of spSP6 RNAP-v4 as a target, and a peptide ligand (TPX2) was fused to the C-terminal fragment of spSP6 RNAP-v4^30^. **(b)** VH and VL domains of anti-HCV IRES RNA antibody were fused to the N- and C-terminal fragments of spSP6 RNAP-v4^16^. *In vitro* transcribed HCV IRES RNA at different concentrations was added as a target molecule. **(c)** Anti-eGFP nanobodies (LaG2 and LaG14) were fused to the N- and C-terminal fragments of spSP6 RNAP-v4^15^. Purified eGFP at different concentrations was added as a target molecule. **(d)** Human Brd4 bromodomain 2 (Brd4^BD2^) and VHL were fused to the N- and C-terminal fragments of spSP6 RNAP-v4^35^. Elongin B and Elongin C proteins were co-expressed in the PURE system to stabilize VHL. PROTAC molecule MZ1 was added as a molecular glue at different conditions. **(a-d)** The data are presented as the mean ± SD from three independent experiments.

We next applied spSP6 RNAP-v4 to detect RNA-binding interactions. Separately expressed VH and VL antibody domains are known to assemble into a heterotrimeric complex upon binding their targets^31^. This concept has been applied to the split T7 RNA polymerase platform in mammalian cells to detect antigen molecules such as proteins, RNA and small molecules^16^. When the VH and VL domains of an anti-HCV IRES antibody were fused to the spSP6 RNAP-v4 fragments, the transcriptional activity increased in a dose-dependent manner upon addition of *in vitro* transcribed IRES RNA (Fig. 4b).

We then evaluated whether spSP6 RNAP-v4 could support sandwich-type detection of protein antigens^15^. Our evolved spSP6 RNAP-v4 was also applied to this system. Anti-eGFP nanobodies LaG2 and LaG14 were fused to the RNAP fragments, yielding robust eGFP-dependent expression of mScarlet-I3 (Fig. 4c)^32^. At high eGFP concentrations (24 μM), the signal decreased, consistent with a hook effect in which each nanobody binds separate antigen molecules, preventing productive reassembly of the RNAP^33^.

Finally, we assessed whether spSP6 RNAP-v4 could detect molecular glues such as proteolysis targeting chimeras (PROTACs)^34^. Brd4 bromodomain 2 (Brd4^BD2^) and human von Hippel-Lindau (VHL) Cullin RING E3 ligase, which are targets of the PROTAC drug MZ1, were fused to the N- and C-terminal fragments of spSP6 RNAP-v4 and Elongin C and Elongin B proteins were co-expressed to stabilize VHL^35^. The PROTAC MZ1, which links the BD2 ligand JQ1 and the VHL ligand VH032, induced dose-dependent activation of spSP6 RNAP-v4 with a hook effect observed above 20 μM (Fig. 4d)^36^. Competition with free JQ1 or VH032 inhibited activation, indicating that MZ1 specifically recruits both Brd4BD2 and VHL to induce RNAP reconstitution (Extended Data Fig. 8c, d).

### Application of evolved SP6 RNAPs in living cells

Finally, we evaluated the activity of the evolved SP6 RNA polymerases in living cells using the rapamycin-inducible split RNAP assay as a representative system. To assess their activity in *E. coli*, the polymerases were placed under an arabinose-inducible promoter on a pBAD vector, while a mScarlet-I3^32^ reporter gene was placed under an SP6 or T7 promoter on a pACYC vector (Extended Data Fig. 9a). *E. coli* cells were co-transformed with these two plasmids. Upon arabinose induction, the RNAPs were expressed and could subsequently transcribe the mScarlet-I3 mRNA in the presence of rapamycin. After overnight culture, the fluorescence of mScarlet-I3 relative to cell density (OD700)^37^ indicated rapamycin-inducible protein expression. Although Rapa-spSP6-v4 showed no rapamycin-dependent activity, Rapa-spSP6-v3 exhibited a better signal-to-noise ratio than Rapa-spT7 RNAP in *E. coli* cells (13.4-fold vs. 3.78-fold) (Extended Data Fig. 9b).

We next examined the activity of SP6 RNAPs in mammalian cells, preparing vectors encoding both RNAPs and reporter genes under internal ribosome entry sites (IRES) (Fig. 5a)^14^. HEK293 cells transfected with vectors encoding eGFP as a reporter gene upon treatment with 20 nM rapamycin showed enhanced intracellular fluorescence for Rapa-spSP6-v3 and Rapa-spSP6-v4, similar to that observed for Rapa-spT7 RNAP (N-29-1) in fluorescence microscopy images (Fig. 5b). To quantify RNA polymerase activities, we next transfected HEK293 cells with plasmids encoding NanoLuc as a reporter gene^38^. Both Rapa-spSP6-v3 and Rapa-spSP6-v4 displayed rapamycin-dependent activation comparable to Rapa-spT7 RNAP (Fig. 5c–e), and SP6 RNAP-v3 exhibited 5.22-fold higher activity than SP6 RNAP-wt (Fig. 5f). These results differ from those observed in *E. coli*. These contrasting behaviors among the PURE system, bacterial and mammalian cells likely reflect differences in intracellular environments, including ionic composition, molecular crowding, chaperone availability, and DNA-binding proteins.

**Fig. 5:**
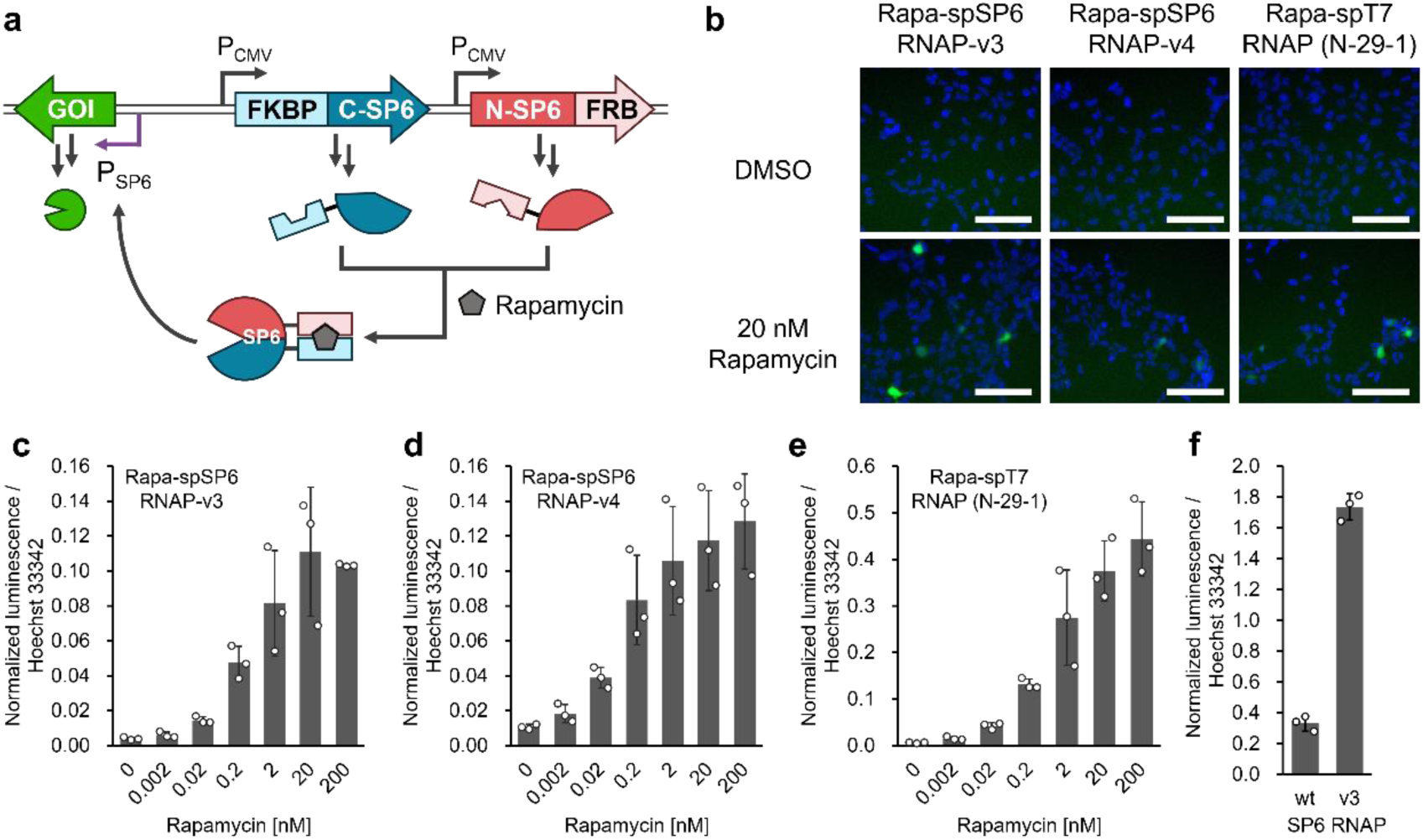
Evolved RNA polymerases are active in mammalian cells. **(a)** Schematic illustration of rapamycin-induced gene of interest (GOI) expression in mammalian cells. **(b)** Fluorescence microscopy images of HEK293 cells expressing eGFP (green) upon rapamycin treatment. Nuclei were stained with Hoechst 33342 (blue). Scale bars are 100 μm. **(c-e)** Rapamycin-induced NanoLuc expression in HEK293 cells. **(f)** NanoLuc expression in HEK293 cells transfected with pCMV-SP6 RNAP-wt-NanoLuc or pCMV-SP6 RNAP-v3-NanoLuc. Luminescence signal was normalized by the fluorescence intensity of cells stained with Hoechst 33342. The data are presented as the mean ± SD from three independent experiments.

In summary, SP6 RNAP-v3 displayed comparable or superior activity compared with wild-type SP6 RNAP in IVTT, *E. coli*, and mammalian cells. The proximity-dependent spSP6 RNAP-v4 functioned in both IVTT and mammalian cells, whereas spSP6 RNAP-v3 outperformed v4 in *E. coli*. These results clearly demonstrate the broad applicability of our evolved SP6 RNAPs across diverse biological systems.

## Discussion

In this study, we developed the cell-free FADS platform for the directed evolution of complex molecular machinery. This platform provides the genotype–phenotype coupling and selection entirely in a cell-free, amplification-free droplet workflow, enabling rapid evolution. By utilizing this technology, we evolved SP6 RNA polymerase, which is sensitive to its environment, into a robust variant optimized for cell-free systems (Extended Data Fig. 10). We then demonstrated the utility of the platform by evolving this variant into a proximity-dependent biosensor, spSP6 RNAP-v4, which exhibits superior signal-to-noise ratio and orthogonality to existing tools particularly in cell-free platforms.

The power of our cell-free FADS platform lies in its simple workflow, which enables one-day cycle including DNA preparation, protein expression in droplets and high-throughput screening. A key advantage is that it avoids the need for any DNA amplification within the droplets, utilizing linear dsDNA templates generated by PCR. This eliminates the need for time-consuming plasmid cloning and cell culturing. While not performed in this study, the output of our screening is directly compatible with long-read next-generation sequencing, enabling a fully cell-free directed evolution pipeline^39^. This cell-free nature expands the scope of evolvable targets including cytotoxic proteins. In addition, combined with genetic code reprogramming and posttranslational modification methods, it further opens the door to evolving proteins containing diverse non-canonical building blocks^40–42^.

This platform enabled us to tackle the evolution of SP6 RNAP-v3 and gain valuable insights. Before the directed evolution, DNA-binding domain Sso7d was fused to the N-terminus of SP6 RNAP-wt to enhance its activity in the PURE system. It has recently been reported that Sso7d tethering to T7 RNAP improved salt tolerance and decreased dsRNA impurity^43^. These findings indicate that fusion of DNA-binding domain to RNAP has potential to be a general strategy to engineer RNAPs. The N-terminal domain (NTD) of SP6 RNAP was a primary target for mutagenesis, a strategy inspired by previous work on the related T7 RNAP, where the NTD is known to be critical for transcriptional activity^22^. While the precise mechanism of each mutation requires further study, our success showed the importance of this domain in polymerase function especially in high-salt buffer conditions. We also found that the Sso7d domain was no longer required for SP6 RNAP-v3 activity in the PURE system. This suggests that the evolution campaign improved the intrinsic properties of the enzyme itself such as its expression level, solubility, folding and processivity. Consistent with this, purified SP6 RNAP-v3 showed enhanced transcriptional activity under PURE-like high-salt conditions. To our knowledge, wild-type SP6 RNAP has not been widely used in cell-free applications, but SP6 RNAP-v3, with its high activity in robust conditions, is now a valuable new component for constructing complex and robust synthetic gene circuits^44^.

Split RNA polymerases are already recognized as powerful tools for various applications, from detecting molecular interactions to building cellular logic gates both *in vitro* and in cells^14–16,28,29^. The spSP6 RNAP-v4 we developed represents an advancement in this class of biosensors for *in vitro* usage. Its most critical feature is its lower background levels than other split RNAP systems and comparable signal-to-noise ratio to the other enzyme-based molecular detection systems such as split luciferase and biotin ligase^15,45^. Our study demonstrates the broad applicability of spSP6 RNAP-v4 to convert the various types of molecular interactions, such as peptide, protein, nucleic acids and molecular glues, to reporter gene expression. Furthermore, the demonstrated orthogonality to the spT7 RNAP system expands the synthetic biology toolbox, allowing for the construction of multi-layered genetic circuits that can process multiple inputs independently. Importantly, in applications involving large numbers of samples, multi-step signal cascades, or low-abundance targets, suppressing background leakage becomes more critical. The near-zero background activity of spSP6 RNAP-v4 therefore provides a decisive advantage for robust diagnostics, therapeutics, and the engineering of artificial cells.

While our system enabled direct detection of gene expression signals from a single DNA molecule, some proteins would still remain as difficult targets for direct activity screening due to insufficient gene expression level. One potential solution is to combine one-pot DNA amplification with IVTT. However, previous methods, while achieving protein expression in liposomes, suffer from high variability in amplification that leads to inconsistent protein yields, a major issue for quantitative directed evolution^46^. An alternative approach could be to reduce the volume of the compartments, thereby increasing the effective concentration of the expressed protein^47^. Another drawback is that substrates which inhibit the IVTT system cannot be used in this system. To address this issue, a method has recently been reported in which DNA is amplified inside agarose beads, followed by expression of enzymes via IVTT, after which buffer exchange is performed to introduce the substrate^11^. This condition control strategy may help reduce over-optimization of target proteins to the PURE system-specific environment. In this study, we observed that evolution of spSP6 RNAP-v4 under PURE system conditions did not confer improved activity *in vivo*, highlighting the context dependence of evolved function. Screening under conditions that more closely mimic the intended application may facilitate the identification of variants that retain activity in downstream demonstrations. While these approaches increase experimental complexity, careful selection of the screening environment remains an important consideration for effective directed evolution.

In conclusion, our work establishes a powerful and versatile cell-free directed evolution platform enabling us to engineer both a highly robust SP6 RNA polymerase and proximity-dependent spSP6 RNAP-based biosensor. These engineered molecular tools and the simplified cell-free platform accelerate synthetic biology research, diagnostics, and drug discovery by enabling the design and optimization of molecular systems with unprecedented precision and speed.

## Acknowledgements

We thank the technical staff Ms. Sakura Shimizu for molecular cloning and IVTT experiments. This work was supported by Japan Society for the Promotion of Science Grant-in-Aid for Transformative Research Areas (B) 21H05119 (to N.T. and T.F.), Grant in Aid for Scientific Research (B) 25K01913 (to N.T. and T.F.), Grant-in-Aid for Early-Career Scientists 24K17798 (to T.F.), The Naito Foundation (to N.T.), and Chugai Foundation for Innovative Drug Discovery Science C-FINDs (to N.T.), and Japan Society for the Promotion of Science Research Fellowships for Young Scientists 24KJ1042 (to K.T.), and Japan Science and Technology Green Technologies of Excellence (GteX) Program JPMJGX23B1 (to H.N.). Molecular graphics were made with UCSF ChimeraX, developed by the Resource for Biocomputing, Visualization, and Informatics at the University of California, San Francisco, with support from National Institutes of Health R01-GM129325 and the Office of Cyber Infrastructure and Computational Biology, National Institute of Allergy and Infectious Diseases.

## Author contributions

T.F. performed optimization of b-PURE reaction conditions. N.T. performed directed evolution and characterization of SP6 RNA polymerases. K.T. performed protein expression and purification. N.T. supervised this project and wrote the manuscript with input from all the other authors.

## Materials and Methods

Polymerase Chain Reaction (PCR) was carried out using KOD One PCR Master Mix (KMM-201, TOYOBO). PCR products were purified by FastGene Gel/PCR Extraction Kit (FG-91302, NIPPON Genetics). Restriction enzymes were purchased from New England Biolabs. DNA ligation reaction was carried out using Ligation high Ver.2 (LGK-201, TOYOBO). The plasmids were purified using FastGene PlasmidMini Kit (FG-90502, NIPPON Genetics) or PureLink™ HiPure Plasmid Midiprep Kit (K210004, Thermo Fisher). Plasmid sequences were confirmed by Sanger sequencing (Eurofins Genomics). DNA oligos were purchased from Integrated DNA Technologies or Eurofins Genomics. The gene fragments were purchased from Integrated DNA Technologies or GenScript. The sequences of DNA oligos, proteins, DNA templates for IVTT and DNA templates for in vitro transcription are listed in Supplementary Tables 1-4.

### Molecular cloning

pRSETb-His-SKIK-eGFP: The gene fragment encoding eGFP was digested by NdeI and EcoRI, and ligated into a pRSETb vector, which had been digested with the same restriction enzymes, to give plasmid pRSETb-eGFP.

pRSETb-His-mNG: The gene fragment encoding mNG was digested by NdeI and EcoRI, and ligated into a pRSETb vector, which had been digested with the same restriction enzymes, to give plasmid pRSETb-His-mNG.

pRSETb-SP6pro-mNG: The gene fragment encoding SP6 promoter was introduced by NEBuilder HiFi DNA Assembly (E2621, New England Biolabs) into the vector prepared by inverse PCR with primers Oligo01, Oligo02 and pRSETb-His-mNG to give plasmid pRSETb-SP6pro-mNG.

pRSETb-SP6 RNAP-wt: The gene fragment encoding HAT-tagged SP6 RNAP was digested by NdeI and EcoRI, and ligated into a pRSETb vector, which had been digested with the same restriction enzymes, to give plasmid pRSETb-HAT-SP6 RNAP-wt.

pRSETb-HAT-Sso7d-Linker(60)-SP6 RNAP-wt: The gene fragment encoding a 60-amino-acid linker was introduced by NEBuilder HiFi DNA Assembly into the vector prepared by inverse PCR with primers Oligo03, Oligo04 and pRSETb-HAT-SP6 RNAP-wt to give plasmid pRSETb-HAT-Linker(60)-SP6 RNAP-wt. The gene fragment encoding Sso7d was digested by SacI and SpeI, and ligated into a pRSETb-HAT-Linker(60)-SP6 RNAP-wt, which had been digested by the same restriction enzymes, to give plasmid pRSETb-HAT-Sso7d-Linker(60)-SP6 RNAP-wt.

pRSETb-HAT-Sso7d-Linker(41)-SP6 RNAP-wt: The gene fragment encoding Sso7d was digested by SacI and SpeI, and ligated into a pRSETb-HAT-Linker(60)-SP6 RNAP-wt, which had been digested by SacI and NheI, to give plasmid pRSETb-HAT-Sso7d-Linker(41)-SP6 RNAP-wt.

pRSETb-HAT-Sso7d-Linker(21)-SP6 RNAP-wt: The gene fragment encoding Sso7d was digested by SacI and KpnI, and ligated into a pRSETb-HAT-Linker(60)-SP6 RNAP-wt, which had been digested by the same restriction enzymes, to give plasmid pRSETb-HAT-Sso7d-Linker(21)-SP6 RNAP-wt.

pRSETb-SP6 RNAP: Sso7d-linker regions of pRSETb-HAT-Sso7d-linker(41)-SP6 RNAPs were deleted by site-directed mutagenesis using Oligo05, Oligo06 and Oligo07.

pET22b-SP6 RNAP: The gene fragments encoding RNAPs were amplified by PCR using Oligo08 and Oligo09 with pRSETb-RNAP as templates. Resulting gene fragments were digested by XbaI and BlpI, and ligated into a pET22b vector, which had been digested by the same restriction enzymes.

pF3A-SP6-E01-WG (BYDV)-SKIK-mNG: The gene fragment encoding SP6 promoter, E01 enhancer (CellFree Science) and SKIK-mNG was introduced by NEBuilder HiFi DNA Assembly into the vector prepared by inverse PCR with primers Oligo10, Oligo11 and PF3A-WG(BYDV) Flexi Vector (L576A, Promega) to give plasmid pF3A-SP6-E01-WG (BYDV)-SKIK-mNG.

pET22b-FKBP-C-SP6 RNAP-v3: The gene fragment encoding FKBP was introduced by NEBuilder HiFi DNA Assembly into the vector prepared by inverse PCR with reverse primer (Oligo12), forward primers (Oligo13, Oligo14, Oligo15, Oligo16, Oligo17, Oligo18, Oligo19 or Oligo20) and pET22b-SP6 RNAP-v3.

pET22b-N-SP6 RNAP-v3-FRB: The gene fragment encoding FRB was introduced by NEBuilder HiFi DNA Assembly into the vector prepared by inverse PCR with forward primer (Oligo21), reverse primers (Oligo22, Oligo23, Oligo24, Oligo25, Oligo26, Oligo27, Oligo28 or Oligo29) and pET22b-SP6 RNAP-v3.

pET22b-Rapa-spSP6 RNAP-v3: The gene fragment encoding ribosomal binding site and N-SP6 RNAP-v3 (−197)-FRB was digested by EcoRI and HindIII, and ligated into a pET22b-FKBP-C-SP6 RNAP-v3 (198+), which had been digested by the same restriction enzymes.

pET22b-Rapa-spSP6 RNAP-v4: The gene fragment encoding N-SP6 RNAP Lib4-3-#16 (−197)-FRB was amplified by PCR using Oligo30 and Oligo31 with pET22b-FKBP-C-SP6 RNAP-v3 (198+)_N-SP6 RNAP Lib4-3-#16 (−197)-FRB as templates. Resulting gene fragment was introduced by NEBuilder HiFi DNA Assembly into the vector prepared by inverse PCR with Oligo32, Oligo33 and pET22b-FKBP-C-SP6 RNAP Lib5-3-#01 (198+) (v4)_N-SP6 RNAP-v3 (−197).

pET22b-Rapa-spT7 RNAP: The gene fragment encoding N-29-1 split N-terminal T7 RNAP-FRB was amplified by PCR using Oligo34 and Oligo35 with p5-79 N-29-1 split N-terminal T7 RNAP variant-GGSGSGSS-FRB expression plasmid (#118087, Addgene)^14^ as templates. Resulting gene fragment was digested by NcoI and HindIII, and ligated into a pET22b-Rapa-spSP6 RNAP-v4, which had been digested by the same restriction enzymes, to give plasmid pET22b-FKBP-C-SP6 RNAP-v4_N-29-1 split N-terminal T7 RNAP-FRB. The gene fragment encoding FKBP-split C-terminal T7 RNAP was amplified by PCR using Oligo36, Oligo37, Oligo38, Oligo39 with p5-39 FKBP-TSGGSG-Split C-terminal T7 RNAP expression plasmid (#118088, Addgene)^14^ as templates. Resulting gene fragment was digested by XbaI and EcoRI, and ligated into a pET22b-Rapa-spSP6 RNAP-v4 or pET22b-FKBP-C-SP6 RNAP-v4_N-29-1 split N-terminal T7 RNAP-FRB, which had been digested by the same restriction enzymes, to give plasmid pET22b-FKBP-split C-terminal T7 RNAP_N-SP6 RNAP-v4-FRB or pET22b-Rapa-spT7 RNAP.

pET22b-Rapa-spCGG RNAP: The gene fragment encoding CGG promoter recognition sequence was introduced by NEBuilder HiFi DNA Assembly into the vector prepared by inverse PCR with primers Oligo40, Oligo41 and pET22b-FKBP-split C-terminal T7 RNAP_N-SP6 RNAP-v4-FRB or pET22b-Rapa-spT7 RNAP to give plasmid pET22b-FKBP-split C-terminal CGG RNAP_N-SP6 RNAP-v4-FRB or pET22b-Rapa-spCGG RNAP.

pET22b-FKBP-C-SP6 RNAP-v4: The gene fragment encoding C-SP6 RNAP-v4 was digested by SacI and EcoRI, and ligated into a pET22b-FKBP-C-SP6 RNAP-v3, which had been digested by the same restriction enzymes.

pET22b-TPX2-C-SP6 RNAP-v4, pET22b-Mb1-C-SP6 RNAP-v4, pET22b-Mb6-C-SP6 RNAP-v4, pET22b-VL-C-SP6 RNAP-v4, pET22b-LaG14-C-SP6 RNAP-v4, pET22b-VHL-C-SP6 RNAP-v4: The gene fragment encoding TPX2, Mb1, Mb6, VL, LaG14 or VHL were digested by SacI and BamHI, and ligated into a pET22b-FKBP-C-SP6 RNAP-v4, which had been digested by the same restriction enzymes.

pET22b-N-SP6 RNAP-v4-FRB: The gene fragment encoding RBS and N-SP6 RNAP-v4 was digested by XbaI and SacI, and ligated into a pET22b-N-SP6 RNAP-v3-FRB, which had been digested by the same restriction enzymes.

pET22b-N-SP6 RNAP-v4-hAurA, pET22b-N-SP6 RNAP-v4-hAurA(Y199K), pET22b-N-SP6 RNAP-v4-VH, pET22b-N-SP6 RNAP-v4-LaG2, pET22b-N-SP6 RNAP-v4-BD2: The gene fragment encoding hAurA, hAurA(Y199K), VH, LaG2 or BD2 were digested by SacI and EcoRI, and ligated into a pET22b-N-SP6 RNAP-v4-FRB, which had been digested by the same restriction enzymes.

pET22b-T7 RNAP-wt, pET22b-Elongin B, pET22b-Elongin C: The gene fragment encoding T7 RNAP, Elongin B or Elongin C were digested by XbaI and XhoI, and ligated into a pET22b vector, which had been digested with the same restriction enzymes.

pET22b-CGG RNAP: The gene fragment encoding CGG promoter recognition sequence was introduced by NEBuilder HiFi DNA Assembly into the vector prepared by inverse PCR with Oligo42, Oligo43 and pET22b-T7 RNAP.

pBAD-RNAPs: The gene fragments encoding SP6 RNAPs were amplified by PCR using Oligo45 and Oligo44 with pRSETb-RNAPs as templates. Resulting gene fragments were digested by NcoI and EcoRI, and ligated into a pBAD vector, which had been digested with the same restriction enzymes, to give plasmid pBAD-SP6 RNAPs. Recognition site by XbaI was introduced under Arabinose promoter in pBAD-vector by site-directed mutagenesis using Oligo46 and Oligo47 to produce pBAD-XbaI vector. The gene fragment encoding T7 RNAP was amplified by PCR using Oligo48 and Oligo49 with pET22b-T7 RNAP-wt as templates. Resulting gene fragment was digested by XbaI and HindIII, and ligated into a pBAD-XbaI vector, which had been digested by the same restriction enzymes, to give plasmid pBAD-T7 RNAP-wt pBAD-Rapa-spRNAPs: The gene fragment encoding Rapa-spSP6 RNAP was amplified by PCR using Oligo50 and Oligo51 with pET22b-spSP6 RNAP-v3 or v4 as templates. Resulting gene fragments were digested by XbaI and HindIII, and ligated into a pBAD-XbaI vector, which had been digested by the same restriction enzymes, to give plasmid pBAD-Rapa-spSP6 RNAP-v3 and v4. The gene fragment encoding Rapa-split T7 RNAP was amplified by PCR using Oligo52 and Oligo53 with pET22b-spT7 RNAP as template. Resulting gene fragments were digested by XbaI and HindIII, and ligated into a pBAD-XbaI vector, which had been digested by the same restriction enzymes, to give plasmid pBAD-Rapa-spT7 RNAP.

pACYC-mScarlet-I3: The gene fragments encoding T7pro-mScarlet-I3 or SP6pro-mScarlet-I3 were introduced by NEBuilder HiFi DNA Assembly into the vector prepared by inverse PCR with primers Oligo54, Oligo55 and pACYC-pTet vector^48^ to give plasmid pACYC-T7pro-mScarlet-I3 and pACYC-SP6pro-mScarlet-I3.

pCMV-RNAPs-eGFP: The gene fragments encoding T7 or SP6 RNA promoter (reverse) and RNA polymerases under CMVd1 promoter (positive) were digested by NsiI and BlpI, and ligated into a p6-8 rapa-T7-mRNA(GFP) plasmid (#118089, Addgene)^14^, which had been digested by the same restriction enzymes, to give plasmid pCMV-SP6 RNAP-wt-eGFP, pCMV-SP6 RNAP-v3-eGFP, pCMV-Rapa-spSP6 RNAP-v3-eGFP and pCMV-Rapa-spSP6 RNAP-v4-eGFP.

pCMV-RNAPs-NanoLuc: The gene fragments encoding NanoLuc were introduced by NEBuilder HiFi DNA Assembly into the vector prepared by inverse PCR with primers Oligo68, Oligo69 and pCMV-SP6 RNAP-wt-eGFP, pCMV-SP6 RNAP-v3-eGFP, pCMV-Rapa-spSP6 RNAP-v3-eGFP, pCMV-Rapa-spSP6 RNAP-v4-eGFP or p6-8 rapa-T7-mRNA(GFP) to give plasmid pCMV-SP6 RNAP-wt-NanoLuc, pCMV-SP6 RNAP-v3-NanoLuc, pCMV-Rapa-spSP6 RNAP-v3-NanoLuc, pCMV-Rapa-spSP6 RNAP-v4-NanoLuc and pCMV-Rapa-spT7 RNAP-NanoLuc.

pRSETb-HCV-IRES-RNA: The gene fragment encoding HCV-IRES-RNA was digested by XbaI and EcoRI, and ligated into a pRSETb vector, which had been digested with the same restriction enzymes.

pRSETb-SP6pro-Broccoli: The gene fragment encoding Broccoli RNA aptamer^49^ was digested by XbaI and BlpI, and ligated into a pRSETb-SP6pro vector, which had been digested with the same restriction enzymes.

### Preparation of linear DNA templates for IVTT and *in vitro* transcription

Linear DNA templates for IVTT and *in vitro* transcription were prepared by PCR with respective plasmids as templates and primers. PCR products were purified by FastGene Gel/PCR Extraction Kit (FG-91302, NIPPON Genetics). Purified PCR products were quantified by Qubit dsDNA Quantification Assay Kits (Q32854, Thermo Fisher Scientific). The amino acids sequences of proteins and nucleotide sequences of DNA templates are listed in Supplementary Tables 2-4.

### Library construction

Error-prone PCR (epPCR) was carried out using the JBS Error-Prone Kit (PP-102, Jena Bioscience) according to the manufacturer’s protocol. Two primers (Oligo56 and Oligo57) were used with pRSETb-HAT-SSo7d-Linker(41)-SP6 RNAP-WT as a template for epPCR. N-terminal and C-terminal gene fragments were amplified by PCR using primers (Oligo58 and Oligo59 for N-terminal, Oligo60 and Oligo61 for C-terminal respectively). The amplified DNAs from epPCR and PCR reactions were purified by agarose gel electrophoresis, and assembled by PCR into full-length DNA templates encoding HAT-Sso7d-Linker(41)-SP6 RNA polymerases Library 1. pRSETb-HAT-Sso7d-Linker(41)-SP6 RNAP Lib1-3-#15 was used as a template to produce HAT-Sso7d-Linker(41)-SP6 RNAP Library 2 following the above-mentioned procedures.

HAT-Sso7d-Linker(41)-SP6 RNA polymerases Library 3 was prepared by staggered extension process (StEP)^50^. PCR containing 0.3 µM Primers (Oligo56 and Oligo57), 0.75 nM of DNA fragment libraries Lib1-3 and Lib2-4, rTaq polymerase (TAP-211, TOYOBO), 0.2 mM dNTP and 1×PCR buffer was carried out for 80 cycles of 94 °C for 30 s and 55 °C for 5s. PCR products were purified by agarose gel electrophoresis. Purified DNA fragments were assembled into DNA templates encoding full-length HAT-Sso7d-Linker(41)-SP6 RNA polymerases Library 3.

Two primers (Oligo30 and Oligo31) were used with pET22b-Rapa-spSP6 RNAP-v3 as a template for epPCR to construct Library 4. N-terminal and C-terminal gene fragments were amplified by PCR using primers (Oligo62 and Oligo32 for N-terminal, Oligo33 and Oligo63 for C-terminal respectively). The amplified DNAs from epPCR or PCR reactions were purified by agarose gel electrophoresis. Purified DNA fragments were assembled by PCR into DNA templates encoding full-length Rapa-spSP6 RNAP Library 4.

Two primers (Oligo64 and Oligo65) were used with pET22b-Rapa-spSP6 RNAP-v3 as a template for epPCR to construct Library 5. N-terminal and C-terminal gene fragments were amplified by PCR using primers (Oligo62 and Oligo66 for N-terminal, Oligo67 and Oligo63 for C-terminal respectively). The amplified DNAs from epPCR or PCR reactions were purified by agarose gel electrophoresis. Purified DNA fragments were assembled by PCR into DNA templates encoding full-length Rapa-spSP6 RNAP Library 5.

### Cell-free IVTT using PURE system

Reconstituted bacterial IVTT was carried out using PURE*frex* 1.0 (PF001, GeneFrontier). The standard reaction solutions consisted of 50% Solution I, 5% Solution II, 5% Solution III, 0.625% EF-P (PFS052, GeneFrontier) and 1.25% DnaK mix (PF003, GeneFrontier). Additional components in each experiment are as follows; 400 nM of SP6 RNAPs (Fig. 2b), 12.8 nM-20 μM of Rapamycin (R0097, Tokyo Chemical Industry) (Fig. 3e), 0.2% dimethyl sulfoxide or 20 μM Rapamycin (Fig. 3f, Extended Data Fig. 6h, i and 7b), 93.8 nM-6 μM HCV IRES RNA (Fig. 4b), 5.86 nM-24 μM eGFP (Fig. 4c), 0 nM-100 μM MZ-1 (BD00797551, BLD pharm) (Fig. 4d), 4 μM MZ-1 and 8 μM-200 μM JQ1 (S7110, Selleck) or 8 μM-200 μM VH032 (BD00758812, BLD pharm) (Fig. 4d, Extended Data Fig. 8c-d). Linear DNA templates in each experiment are as follows; 2-200 pM SP6pro-mNG DNA (Fig. 2b), 800 pM bicistronic T7pro-Rapa-split RNAP DNA and 800 pM SP6pro-mNG DNA or CGGpro-mNG DNA (Fig. 3e-f, Extended Data Fig. 6h, i and 7b), 400 pM of T7pro-split N-terminal SP6 RNAP-v4-VH DNA, 400 pM of T7pro-VL-split C-terminal SP6 RNAP-v4 DNA and 800 pM SP6pro-mNG DNA (Fig. 4b), 400 pM of T7pro-split N-terminal SP6 RNAP-v4-LaG2 DNA, 400 pM of T7pro-LaG4-split C-terminal SP6 RNAP-v4 DNA and 800 pM SP6pro-mScarlet-I3 DNA (Fig. 4c), 24 pM of T7pro-Elongin B DNA, 24 pM of T7pro-Elongin C DNA, 96 pM T7pro-Split N-terminal SP6 RNAP-v4-BD2 DNA and 96 pM T7pro-VHL-split-C-terminal SP6 RNAP-v4 DNA (Fig. 4d).

The b-PURE reaction mixture consisted of 50% Solution I, 5% Solution II, 2.5% Solution III, 10% RNasin Plus Ribonuclease inhibitor (N2615, Promega), 0.625% EF-P (PFS052, GeneFrontier), 1.25% DnaK mix (PF003, GeneFrontier) and 10% T7 RNA polymerase ver.2.0 (2541A, TaKaRa). PURE system reactions were conducted at 37 °C. Additional components in each experiment are as follows; 0.2% dimethyl sulfoxide or 20 μM Rapamycin (Extended Data Fig. 6c). DNAs in each experiment are as follows; 2 pM T7pro-N-SP6 RNAP-v4-hAurA DNA, 2 pM T7pro-ligand-C-SP6 RNAP DNA and 400 pM SP6pro-mNG DNA (Fig. 4a, Extended Data Fig. 8a-b), 200 fM T7pro-SP6 RNAP DNA and 80 pM SP6pro-mNG DNA (Extended Data Fig. 1a-b, 2c), 200 fM T7pro-SP6 RNAP DNA and 8 pM SP6pro-mNG DNA (Extended Data Fig. 4a-b), 200 fM T7pro-N-SP6 RNAP-v3-FRB DNA, 200 fM T7pro-FKBP-C-SP6 RNAP-v3 DNA and 80 pM SP6pro-mNG DNA (Extended Data Fig. 6c).

### Cell-free IVTT using Wheat Germ extract

Wheat germ IVTT was carried out using WEPRO7240H core kit (CFS-C7H, CellFree Sciences). The standard reaction conditions consisted of 10.9 OD/mL of WEPRO7240H, 3.64 μg/mL of creatine kinase, 0.91×SUB-AMIX SGC S1, 0.91×SUB-AMIX SGC S2, 0.91× SUB-AMIX SGC S3, 0.91×SUB-AMIX SGC S4, 150 μM of NTP mix (N0466, New England Biolabs), 1.6 ng/μL of Salmon sperm DNA (15632011, Thermo Fisher Scientific) and 25 nM of SP6 RNAPs. The reactions were conducted in a single layer format at 25 °C. Linear DNA template in Fig. 2c is 20 pM to 2 nM SP6pro-E01-SKIK-mNG DNA.

### Fluorescence Detection

In the case of endpoint measurement, reaction mixture after 18 hours of incubation was transferred to a black 384-well plate (4511, CORNING) and the amount of expressed fluorescent protein was quantified by EnSpire plate reader (PerkinElmer) or Synergy H1 (BioTek). Excitation and emission wavelengths for mNG were 490 nm and 530 nm respectively. In the case of real-time measurement in Fig. 2b-c, Fig. 3f, Fig. 4a and Extended Data Fig. 8a-b, reaction mixture was transferred to 0.2 ml 8-Strip PCR Tube (3247-00, SSIbio) and the fluorescence was measured in Mx3005P Real-Time QPCR System (Agilent Technologies) using the FAM filter set (Ex, 492 nm; Em, 516 nm) to measure the fluorescence of mNG. In the case of Fig. 4b-d and Extended Data Fig. 8c-d, reaction mixture was transferred to white 0.2 ml 8-Tube PCR Strips (TLS0851, Bio-Rad) and the fluorescence was measured in CFX Opus 96 Real-Time PCR Instrument (Bio-Rad) using the FAM filter set (Ex, 450-490 nm; Em, 510-530 nm) for mNG or Texas Red filter set (Ex, 560-590 nm; Em, 610-650 nm) for mScarlet-I3.

### Droplet generation

The water-in-oil emulsion droplets were generated by On-chip Droplet Generator (On-Chip Biotechnologies) in 2D chip-800DG chips (1003002, On-Chip Biotechnologies) at 10 °C. The oil phase was prepared by mixing 008-FluoroSurfactant-5wtH (2003001, On-Chip Biotechnologies) and 008-FluoroSurfactant-0.1wtH (2003002, On-Chip Biotechnologies) at 2:3 ratio. During the evolution of SP6 RNAP, the b-PURE system containing 100 pM sulforhodamine B (152479, Fujifilm WAKO), 20 fM T7pro-SP6 RNAP DNA library and SP6pro-mNG DNA (80 pM for Library 1, 8 pM for Library 2 and 3) was used as the aqueous phase. During the evolution of spSP6 RNAP, the b-PURE system containing 100 pM sulforhodamine B, 10 μM Rapamycin, 1% dimethyl sulfoxide (08904-14, Nacalai Tesque), 20 fM T7pro-SP6 RNAP DNA library and 80 pM SP6pro-mNG DNA was used as the aqueous phase. Air pressures for the aqueous sample and the oil sample were set to 45 kPa and 50 kPa respectively. The droplets were approximately 25 µm in diameter.

### Droplet-based microfluidics screening of SP6 RNA polymerases

The droplets containing cell-free translation mixture were collected in a microtube and incubated at 37 °C for 18 hours to express proteins. The droplets were sorted by fluorescence-activated droplet sorting using an On-chip Sort machine (On-Chip Biotechnologies) in 2D Chip-Z1001 chips (1002004, On-Chip Biotechnologies). The sheath fluid was 008-FluoroSurfactant-0.1wtH. Mineral oil (M5904, Merck) and 1 µL of droplets containing PBS were added to the collection reservoir to improve the recovery of sorted droplets. The droplets bearing ∼25 μm diameter were first gated based on the value of forward scatter and sulforhodamine B fluorescence (FL-4, Ex. 561 nm, Em. 676 ± 37 nm). Then the droplets showing higher mNG fluorescence (FL-2, Ex. 488 nm, Em. 543 ± 22 nm) were sorted. The histogram of mNG fluorescence (FL-2) in each screening experiment was shown in Extended Data Fig. 2a, 3a-b, 6d-e.

The sorted droplets were recovered from the collection reservoir and transferred to a microtube. Droplets were lysed by adding 100 µL of droplet lysis buffer (10 mM Tris-HCl at pH 8.0, 1 mM EDTA, 1% Triton X-100) and lysed droplets were transferred to a new microtube. An equal volume of Phenol/chloroform/isoamyl alcohol (25970-56, Nacalai Tesque) was added to the tube and vigorously mixed by vortex mixer. The tube was centrifuged at 15,000 g for 5 minutes at room temperature and the aqueous phase was recovered. Sodium acetate solution was added to the recovered solution at final concentration 300 mM and two volume of ethanol was added to the mixture. After mixing the solution by vortex, nucleic acids were precipitated by centrifugation at 15,000 g for 15 minutes at room temperature.

DNA fragments were amplified by reverse transcription PCR reaction. The pellet after the ethanol precipitation was dissolved in 5 µL of water and reverse transcription reaction was carried out by SuperScript IV reverse transcriptase (18090010, Thermo Fisher Scientific) following the manufacturer’s protocol. The primer for reverse transcription was Oligo57 (Library 1, 2 and 3), Oligo31 (Library 4) or Oligo65 (Library 5). Reverse transcribed DNA was amplified by PCR and purified by agarose gel electrophoresis. The primers for PCR reaction were Oligo56 and Oligo57 (Library 1, 2 and 3), Oligo30 and Oligo31 (Library 4) or Oligo64 and Oligo65 (Library 5). DNA library encoding full-length SP6 RNAP for next screening was prepared by assembly PCR as described in library preparation.

After several repeats of droplet-based microfluidics screening, sorted DNA library was introduced by NEBuilder HiFi DNA Assembly (E2621X, NEB) into pRSETb vector (Library 1, 2 and 3) or pET-22b vector (Library 4 and 5). XL1-Blue Competent Cells (200249, Agilent Technologies) were transformed with plasmid libraries and plasmids were isolated to identify the sequence of evolved RNA polymerases.

### Expression and Purification of RNA polymerases

All RNA polymerases were produced in *E. coli* BL21-AI cells (C607003, Thermo Fisher Scientific) bearing pET22b-RNA polymerases. Two-liter Erlenmeyer flasks containing 1 L LB medium with 100 μg/mL ampicillin were inoculated with 40 mL overnight cultures and incubated at 37 °C and 200 rpm until the OD600 reached 0.4–0.6. Protein production was induced by adding IPTG and arabinose to a final concentration of 1 mM and 1 mg/mL respectively. Cells were cultured at 37 °C for 4 hours and then harvested by centrifugation at 5,000 g at 4 °C for 20 min. The cell pellet from one 1 L culture was resuspended in 20 mL LB medium, transferred and split into two 50 mL Falcon tubes. The medium used for transfer was removed by centrifugation at 12,000 g and 4 °C for 10 min, decanted, and aliquots of the cell pellet were frozen in liquid nitrogen and stored at −80 °C until purification.

For purification, a cell pellet corresponding to 2 L of culture volume was resuspended in 20 mL lysis buffer (20 mM Tris-HCl buffer at pH 8.0, 20 mM imidazole, 300 mM KCl, and 1 mM DTT). After cell lysis by sonication (60 cycles of 2 sec on, 2 sec off, with amplitude = 50, Sonic Tower UDS-200 TOMY SEIKO) and clearance by centrifugation at 15,000 g at 4 °C for 10 min, the supernatant was loaded onto 1 mL slurry of Nuvia IMAC Metal Affinity Resin (7800802, Bio-Rad) in a gravity flow column at 4 °C. After incubation for 10 min and washing with lysis buffer, proteins were eluted with elution buffer (20 mM Tris-HCl buffer at pH 8.0, 250 mM imidazole, 20 mM KCl, 10 mM MgCl_2_ and 1 mM DTT). The eluted fractions were diluted 10 times in anion exchange buffer (20 mM Tris-HCl buffer at pH 8.0, 20 mM KCl, 10 mM MgCl_2_ and 1 mM DTT) and purified by anion exchange chromatography using a Foresight Nuvia Q Column (7324741, Bio-Rad). The mobile phase consisted of anion exchange buffer containing 20–1000 mM NaCl. Fractions containing RNA polymerases were concentrated using Microsep Advance centrifugal devices with Omega membrane 3K (MCP003C46, Cytiva) and buffer was exchanged to storage buffer (HEPES-KOH at pH 8.0, 100 mM KOAc, 10 mM Mg(OAc)_2_ 1 mM DTT). Protein concentrations were quantified by Qubit protein Assay Kits (Q33212, Thermo Fisher Scientific).

### Expression and purification of eGFP

*E. coli* BL21-Gold(DE3) cells (Agilent Technologies) were transformed with pRSETb-His-SKIK-eGFP. Two-liter Erlenmeyer flasks containing 800 mL LB medium with 100 μg/mL ampicillin were inoculated with 20 mL overnight cultures and incubated at 37 °C and at 150 rpm until OD600 reached 0.6. Protein expression was induced by adding IPTG (M&S TechnoSystems) to a final concentration of 0.4 mM. Cells were cultured at 20 °C overnight and then harvested by centrifugation at 5,000 *g* and 4 °C for 10 min. The cell pellet from one 1000-mL culture was resuspended in 40 mL LB medium, transferred to a 50 mL tube. The medium used for transfer was removed by centrifugation at 5,000 *g* and 4 °C for 10 min, decanted, and aliquots of the cell pellets were frozen in liquid nitrogen and stored at −80 °C until purification. A cell pellet corresponding to 1000 mL of culture volume was resuspended in 10 mL of lysis buffer (50 mM sodium phosphate, 1 M NaCl, 20 mM imidazole at pH 7.4) and lysed by sonication (150 cycles of 1 sec on and 1 sec off with amplitude = 40, Q500 Sonicator, QSONICA). The soluble fraction in the lysate was recovered after centrifugation at 12,000 *g* and 4 °C for 10 min and loaded onto 1 mL slurry of Ni(II)-NTA resin (Nuvia IMAC resin, BIO-RAD) in a gravity flow column. After incubation for 30 min, the resin was washed with 10 mL of binding buffer (50 mM Tris-HCl, 300 mM NaCl, 20 mM imidazole at pH = 8.0) and eluted with 4 aliquots of 1 mL of elution buffer (50 mM Tris-HCl, 300 mM NaCl, 100, 200, 300, or 400 mM imidazole at pH 8.0). The eluted fractions were combined and dialyzed in 1 L of lysis buffer without imidazole at 4 °C twice. The protein quantity was determined by using Pierce 660-nm Protein Assay Reagent (Thermo Fisher Scientific) with NanoDrop OneC (Thermo Fisher Scientific). Final solution was frozen by liquid nitrogen and stored at −80 °C.

### *In vitro* transcription

To measure the transcriptional activity of RNAPs in various buffer conditions, 2 nM of DNA template encoding RNA aptamer under SP6 RNA promoter (SP6pro-Broccoli) was mixed with 400 nM of purified SP6 RNAP with various buffer conditions (see Extended Data Fig. 5). After the incubation at 37 °C for 3 hours, 20% volume of 500 mM EDTA solution was added to the reaction mixture to stop the transcription. Equal amount of 2× RNA loading buffer (8 M Urea, 2 mM EDTA, 2 mM Tris-HCl, pH 7.5, 0.5 g/L Bromophenol blue) was added to the stopped reaction mixture and the solution was incubated at 95 °C for 5 min. Denatured RNA samples were loaded onto denaturing PAGE gel (8% acrylamide, 8 M urea, 1× TBE). The gels were stained with WSE-7135 EzPreStain DNA&RNA (2332397, ATTO) according to the manufacturer’s protocol. Stained gels were visualized by Typhoon FLA 9500 (GE Healthcare). HCV-IRES RNA was prepared by *in vitro* transcription. *In vitro* transcription reactions were performed using in-house expressed T7 RNA polymerase in reaction mixture (40 mM Tris-HCl pH = 8.0, 1 mM spermidine, 0.01% (v/v) Triton X-100, 10 mM DTT, 30 mM MgCl_2_, 5 mM each ATP, UTP, CTP (Combi-Blocks) and GTP (FUJIFILM), 30 mM KOH, 0.12 μM T7 RNA polymerase) with DNA template (T7pro-HCV IRES RNA) at 37 °C for 18 hours. Template DNA was digested by adding RQ1 DNase (Promega) with DNase reaction buffer (40 mM Tris-HCl at pH = 8.0, 10 mM NaCl, 6 mM MgCl_2_, 1 mM CaCl_2_ as final concentration) at 37 °C for 1 hour. RNA was precipitated with isopropanol and dissolved in water. The RNA concentration was quantified by Qubit RNA Quantification Broad Range Assay Kits (Q10210, Thermo Fisher Scientific).

### Reporter assays in bacterial cells

TOP10 chemically competent cells (C404003, Thermo Fisher Scientific) were transformed with two plasmids: (i) arabinose inducible pBAD-RNAP, and (ii) pACYC vector encoding mScarlet-I3 under T7 or SP6 promoter. The transformed cells were then plated onto agar plates with 50 μg/mL ampicillin and 20 μg/mL chloramphenicol. Single colonies were inoculated in 2 mL of LB medium containing 50 μg/mL ampicillin, 20 μg/mL chloramphenicol, and cultured overnight at 37 °C. Each well of a 96-well deep well plate (4-4967-04, VIOLAMO) containing 0.6 mL of LB with antibiotics, 8 μg/mL arabinose (A0515, Tokyo Chemical Industry), and rapamycin (0 nM or 100 μM) was inoculated with 30 μL of the overnight culture. After growth with shaking at 1,700 rpm in a deep well maximizer (MBR-022R, TAITECH) at 37 °C for 4 h, 200 μL of each culture was transferred to a 96-well black wall, clear bottom plate (3603, Corning), and mScarlet-I3 fluorescence (Ex 560 nm, Em. 600 nm) and OD700 were measured on a Synergy H1 (BioTek). After the additional shaking at 1,700 rpm in a deep well maximizer at 25 °C for 18 h, mScarlet-I3 fluorescence and OD700 were measured in the same manner. The data were analyzed by dividing the background-corrected mScarlet-I3 fluorescence values by the background-corrected OD700 value^37^.

### Mammalian cell culture

HEK293 cells (JCRB9068, JCRB Cell Bank) were cultured in DMEM with 1.0 g/L Glucose, L-Gln and sodium pyruvate (08456-65, Nacalai Tesque) supplemented with 10% fetal bovine serum (FBS, F7524, Sigma, non-USA origin) and 1% penicillin/streptomycin (P/S, 26253-84, Nacalai Tesque) at 37 °C and 5% CO_2_ atmosphere. Tissue-culture treated dishes (4-5776-03, VIOLAMO) were used for passages.

### Reporter assays in mammalian cells

For fluorescence microscopy imaging, HEK293 cells were plated on a 96-well Black/Clear Bottom Plate, TC Surface (3603, Corning) at 10,000 cells per well with 200 µL of DMEM containing 10% FBS and 1% P/S. After 24 hours, the medium was exchanged with DMEM containing 10% FBS, 1% P/S and rapamycin (or 0.2% DMSO) and 100 ng of the plasmids were transfected into cells using 0.6 μL of ViaFect Transfection Reagent (E4981, Promega) according to the manufacturer’s protocol. After an additional 48 h incubation, the cell nuclei were stained using Cell Count Normalization Kit (342-09393, Fujifilm) according to the manufacturer’s culture medium exchange method. The cells were imaged on an ECHO Revolve microscope using FITC filter set (Ex, 470 nm; Em, 525 nm) and DAPI filter set (Ex, 380 nm; Em, 450 nm). To quantify the luciferase expression level, HEK293 cells were cultured on a 96-well White/Clear Bottom Plate, TC Surface (3610, Corning). Plasmids encoding NanoLuc as reporter gene were transfected into cells as described above. After an additional 48 hours, the cell nuclei were stained using Cell Count Normalization Kit (342-09393, Fujifilm) according to the manufacturer’s culture medium exchange method. The number of cells was quantified by measuring fluorescence (Ex 350 nm, Em. 461 nm) on a Synergy H1 (BioTek). Then, the activity of NanoLuc expressed in cells was measured using Nano-Glo Luciferase Assay System (N1120, Promega) according to the manufacturer’s lytic method. Chemiluminescence values were normalized by fluorescence values (Ex 350 nm, Em. 461 nm).

## Supplementary information

### Supplementary Tables

**Supplementary Table 1:**
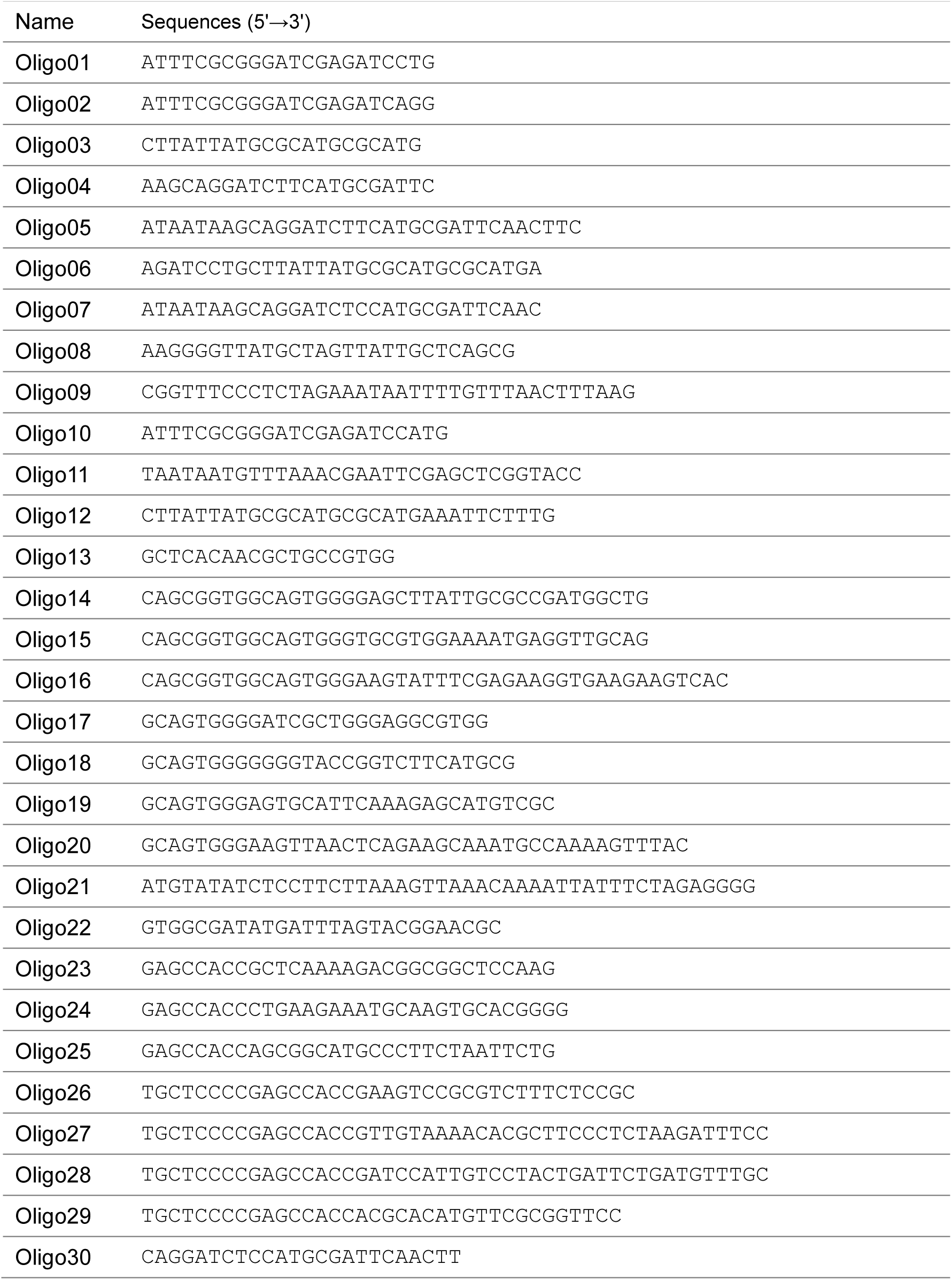
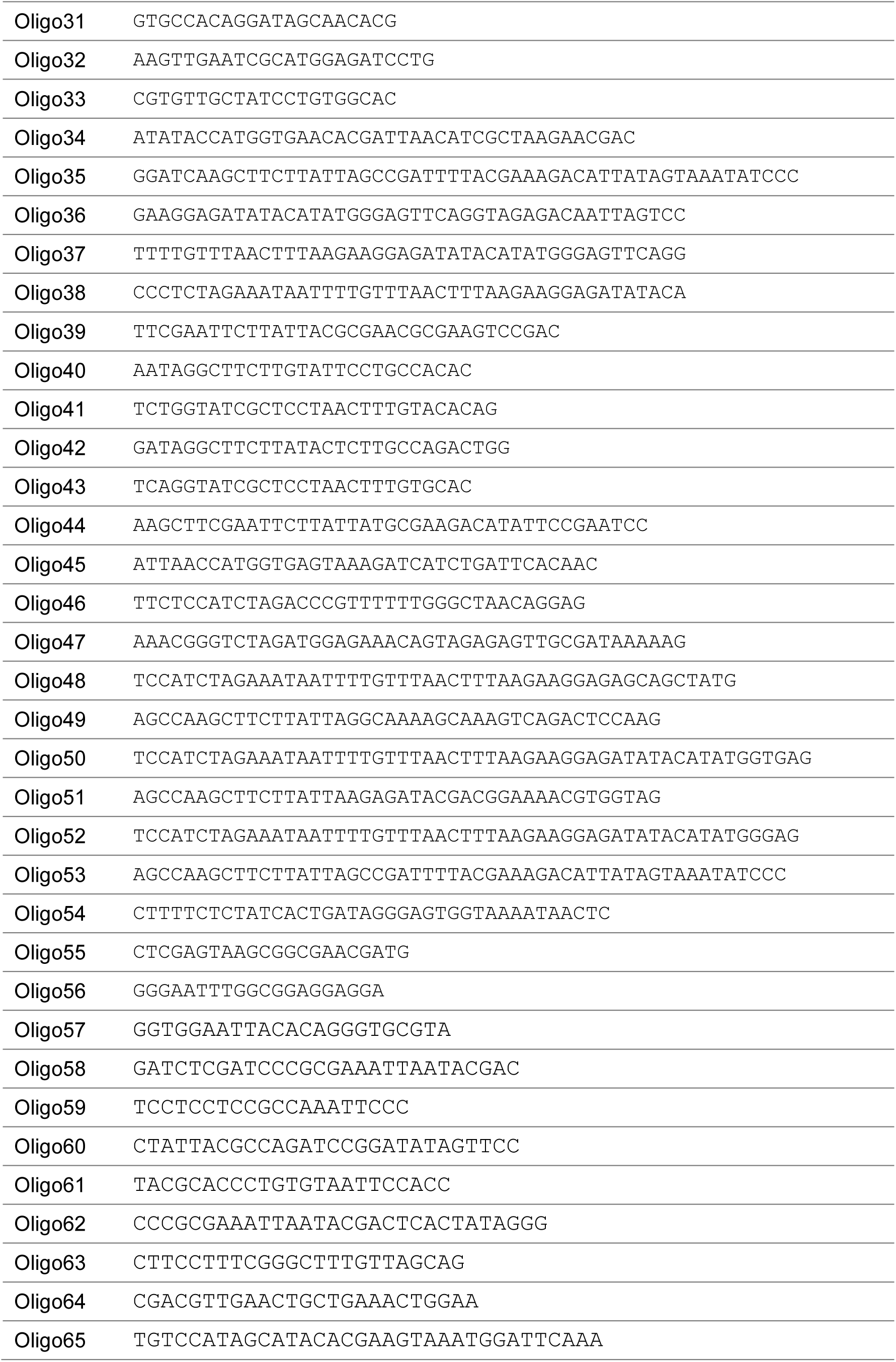
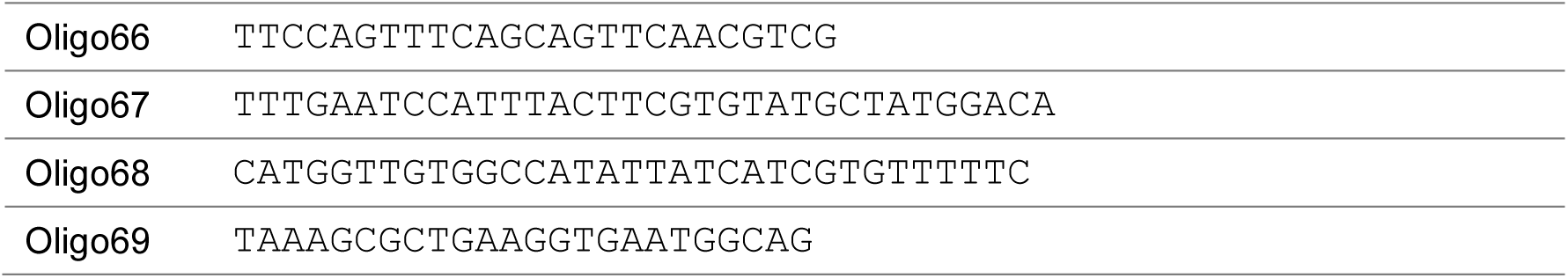
Primers.

**Supplementary Table 2:**
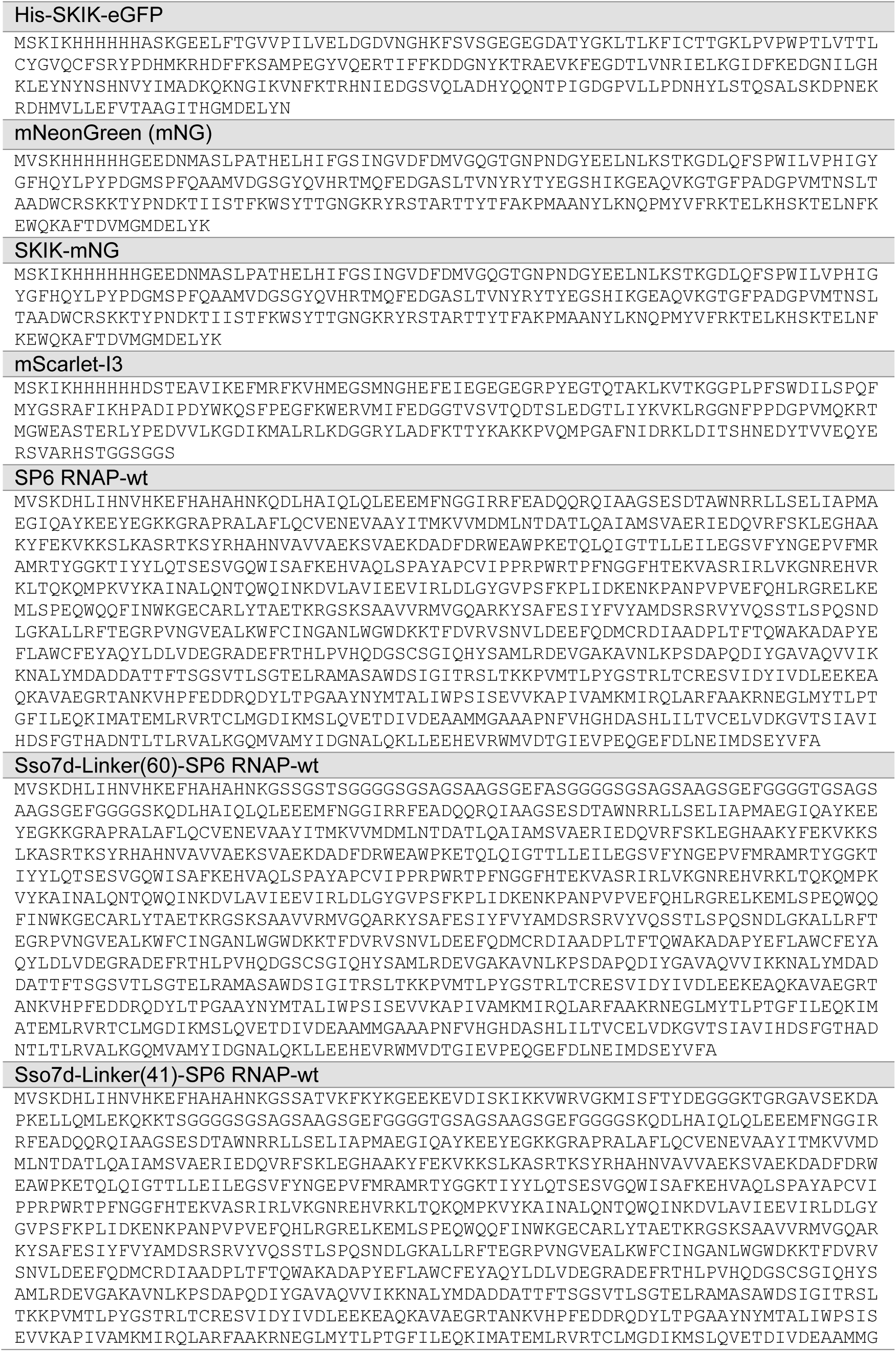
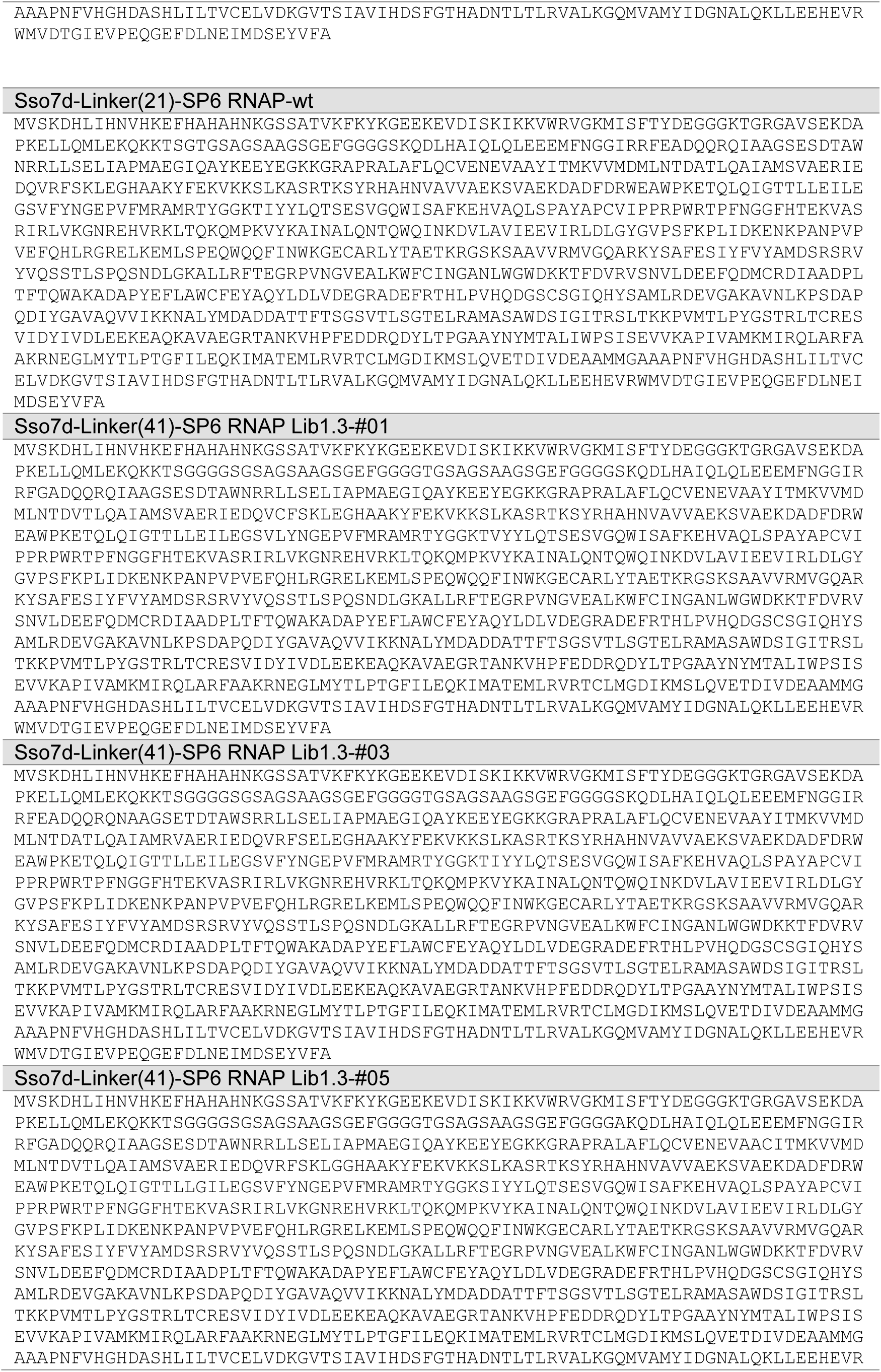
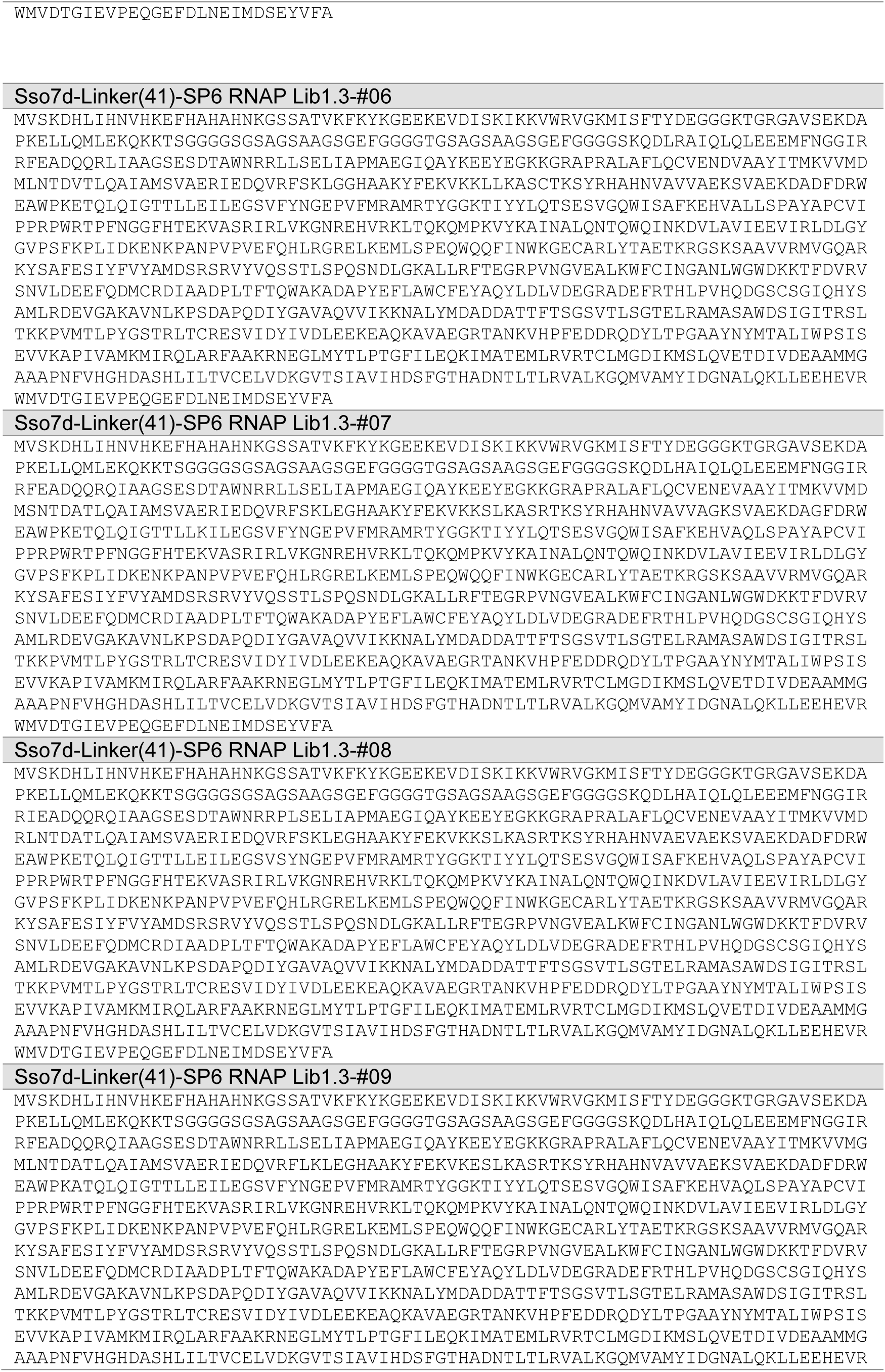
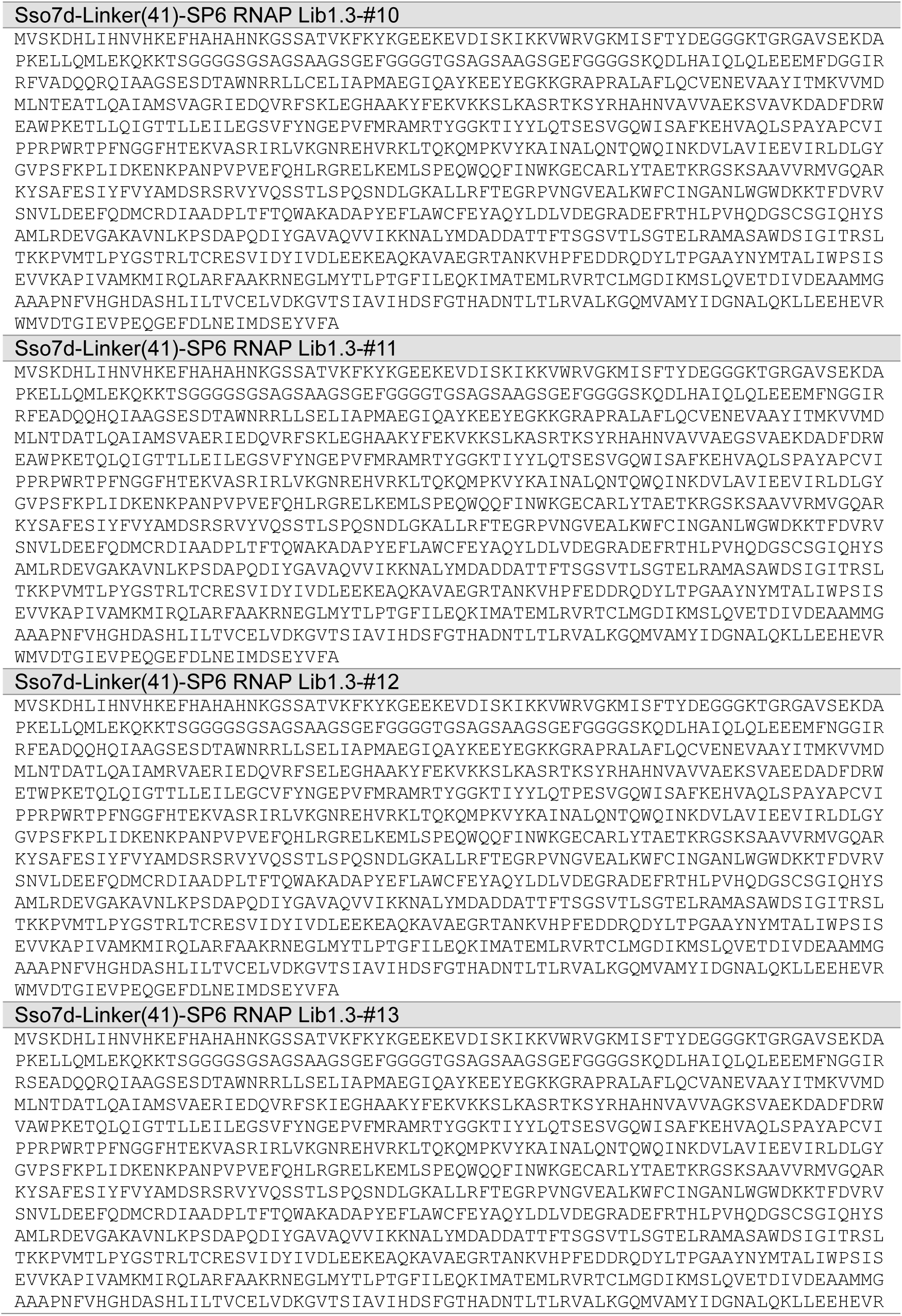
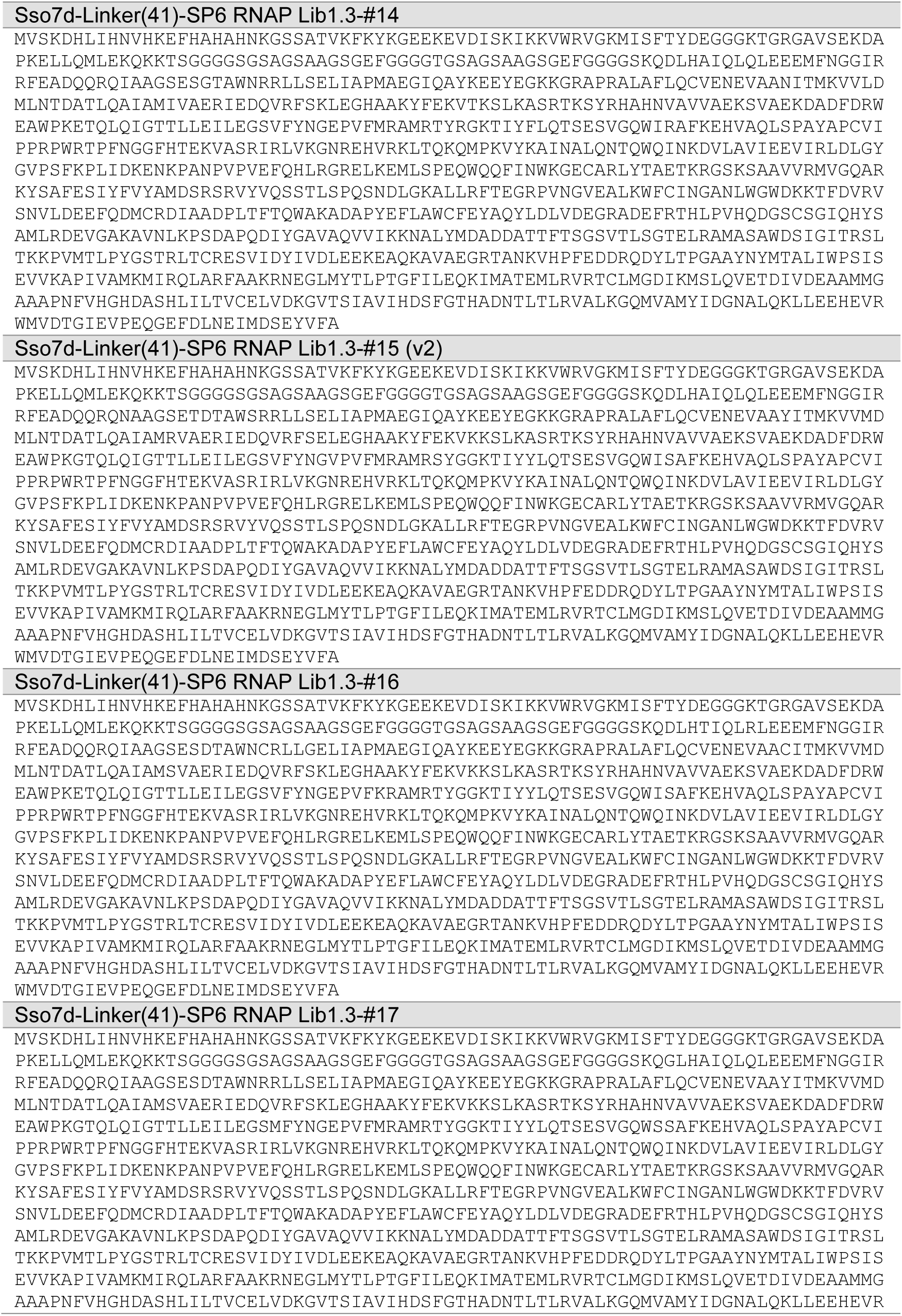
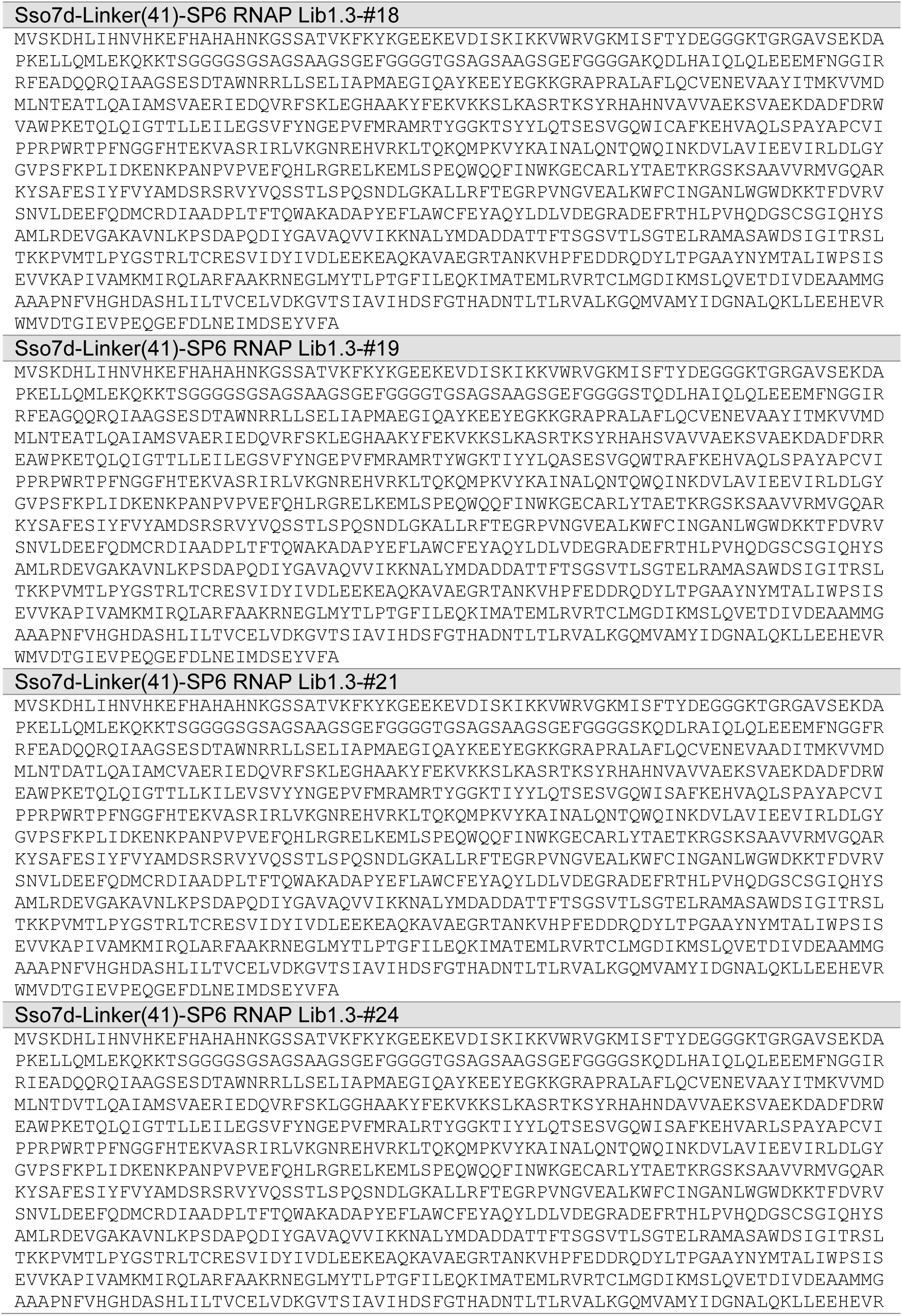
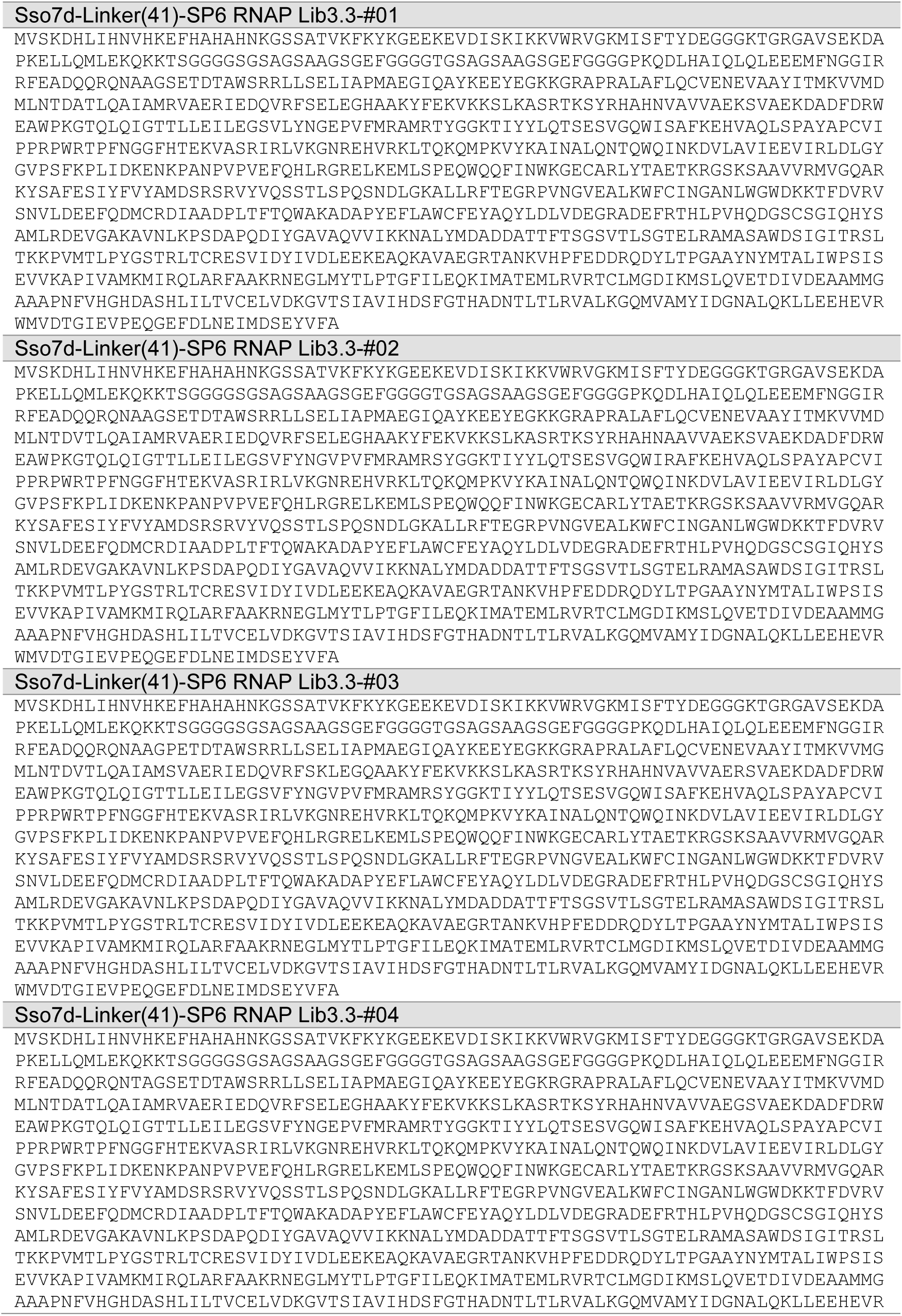
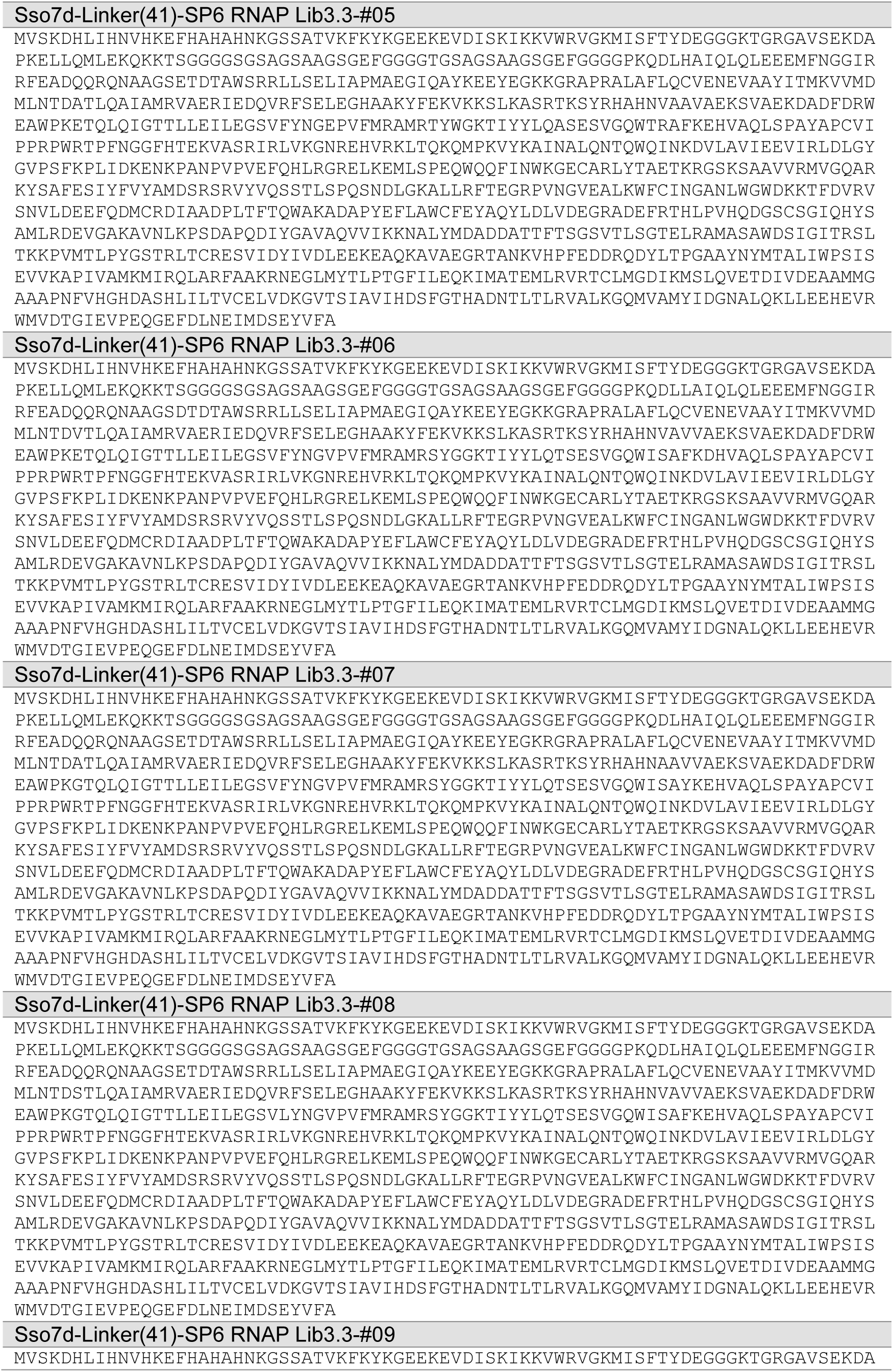
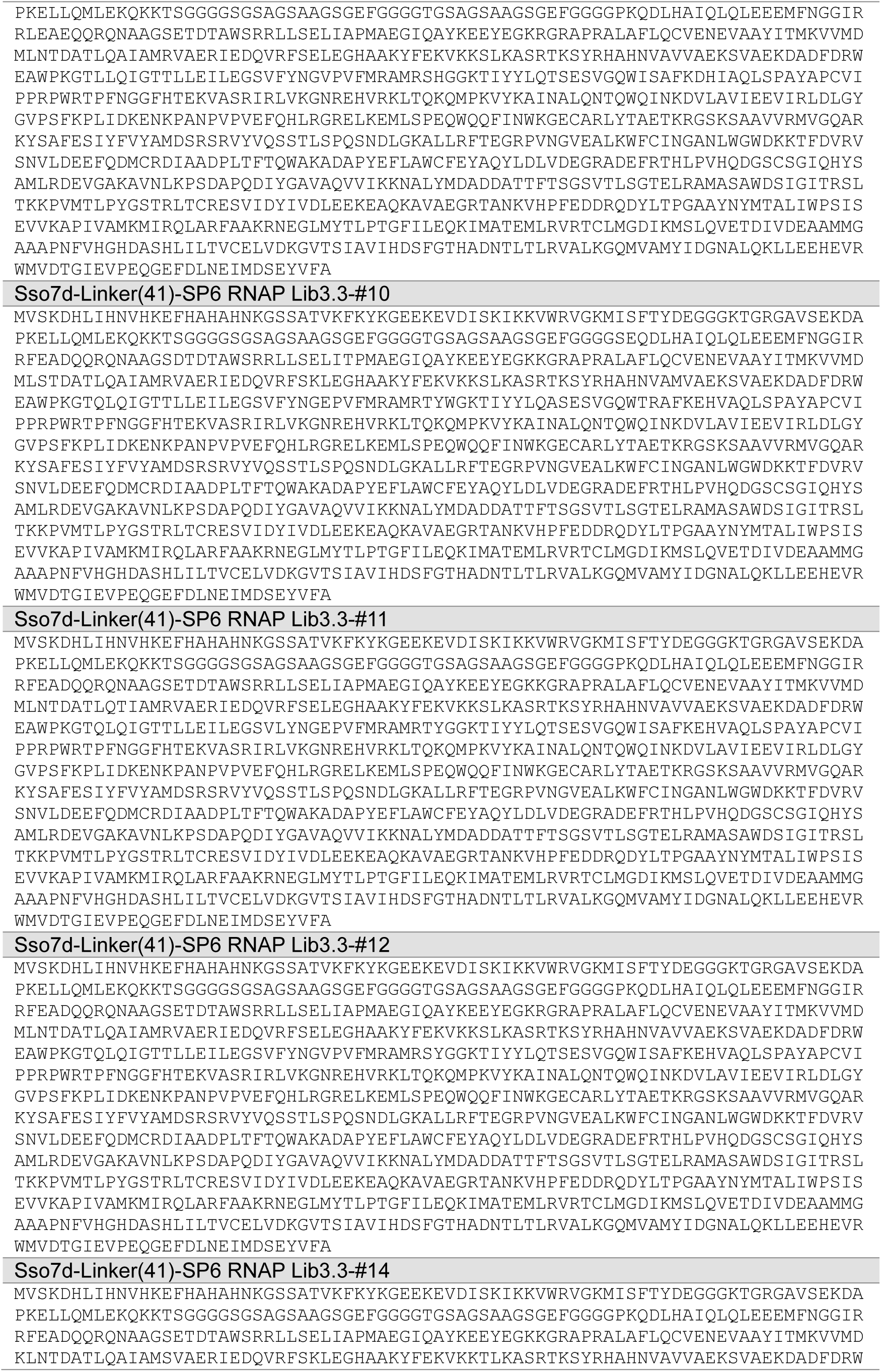
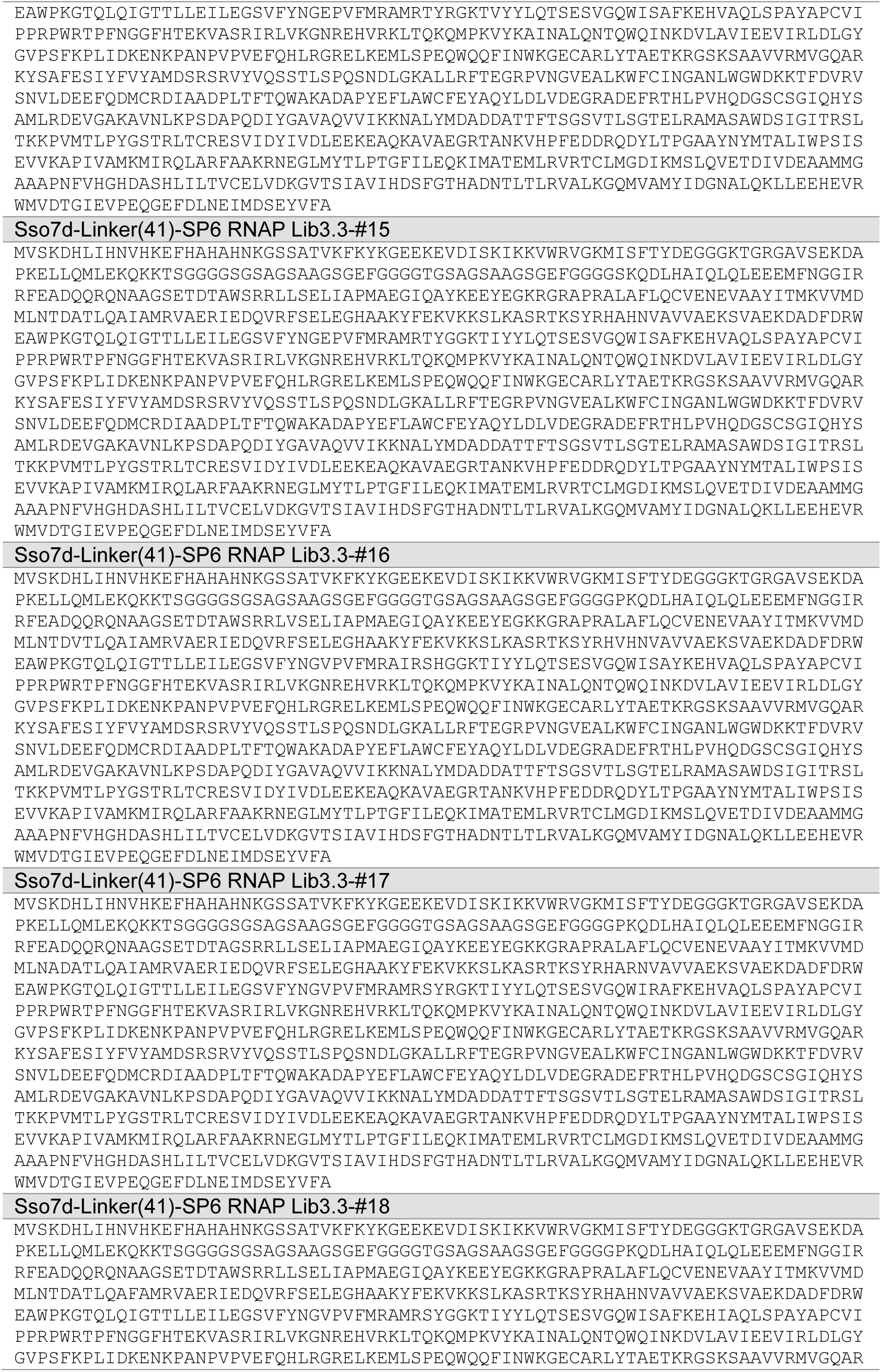
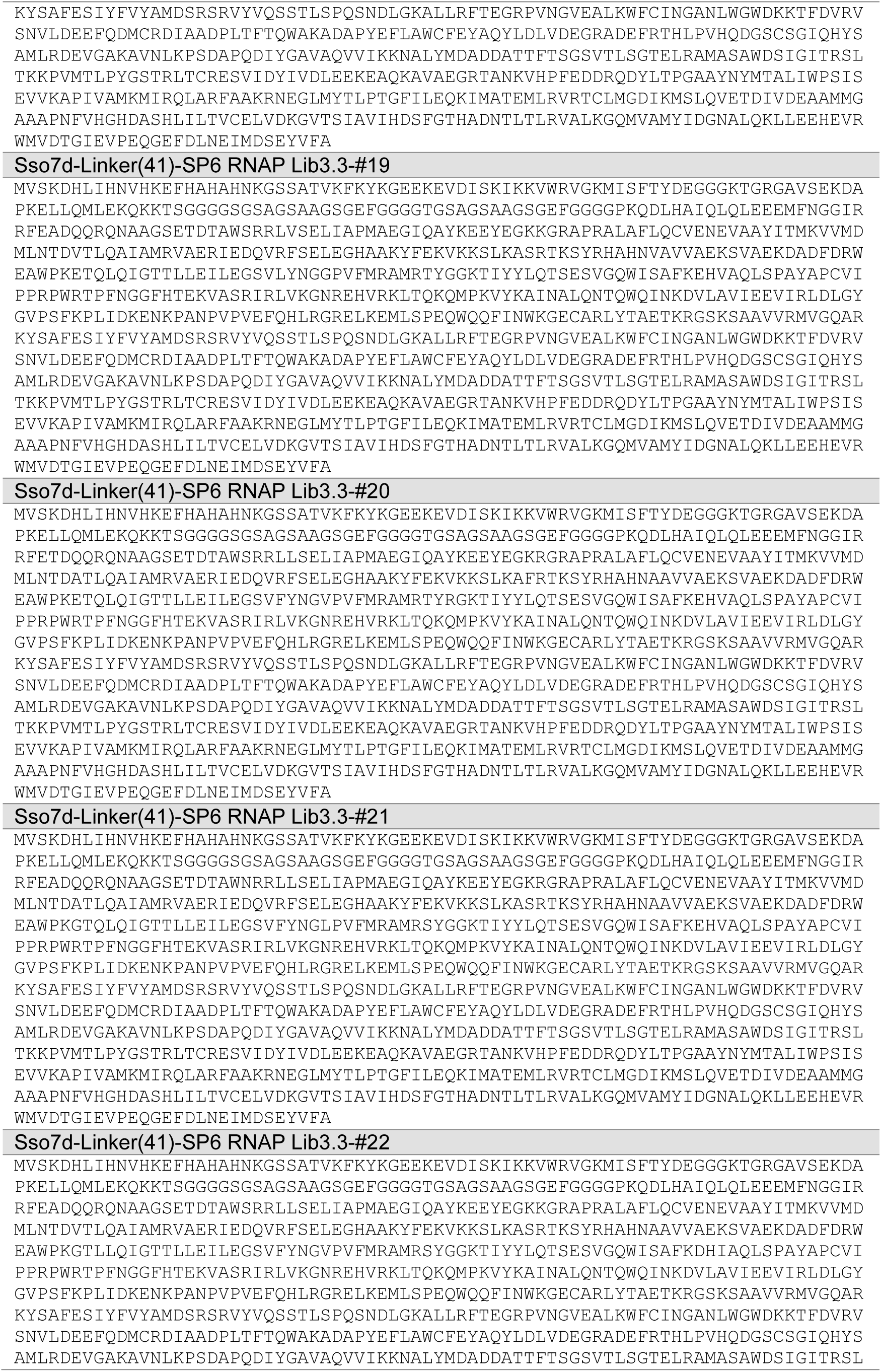
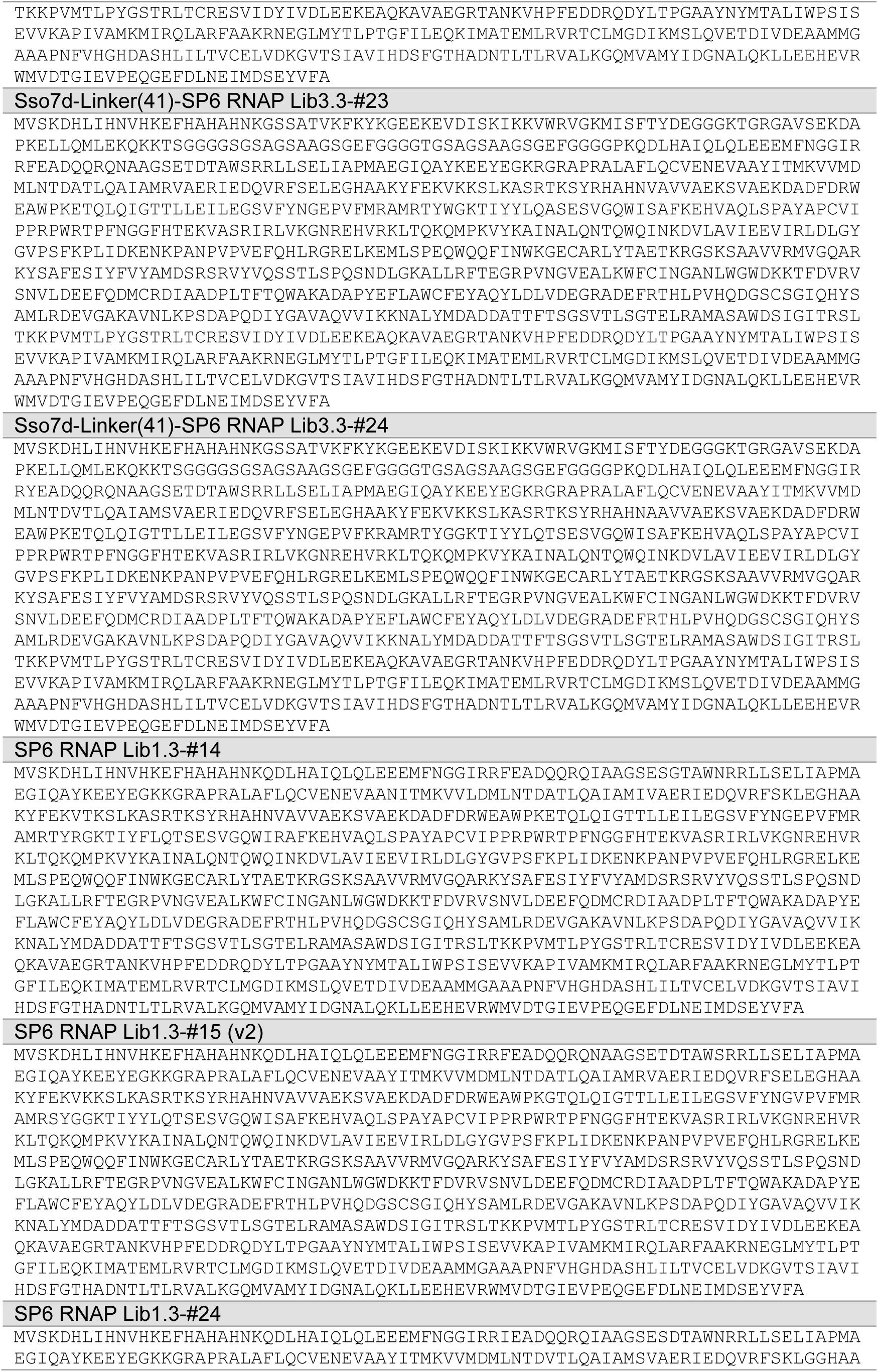
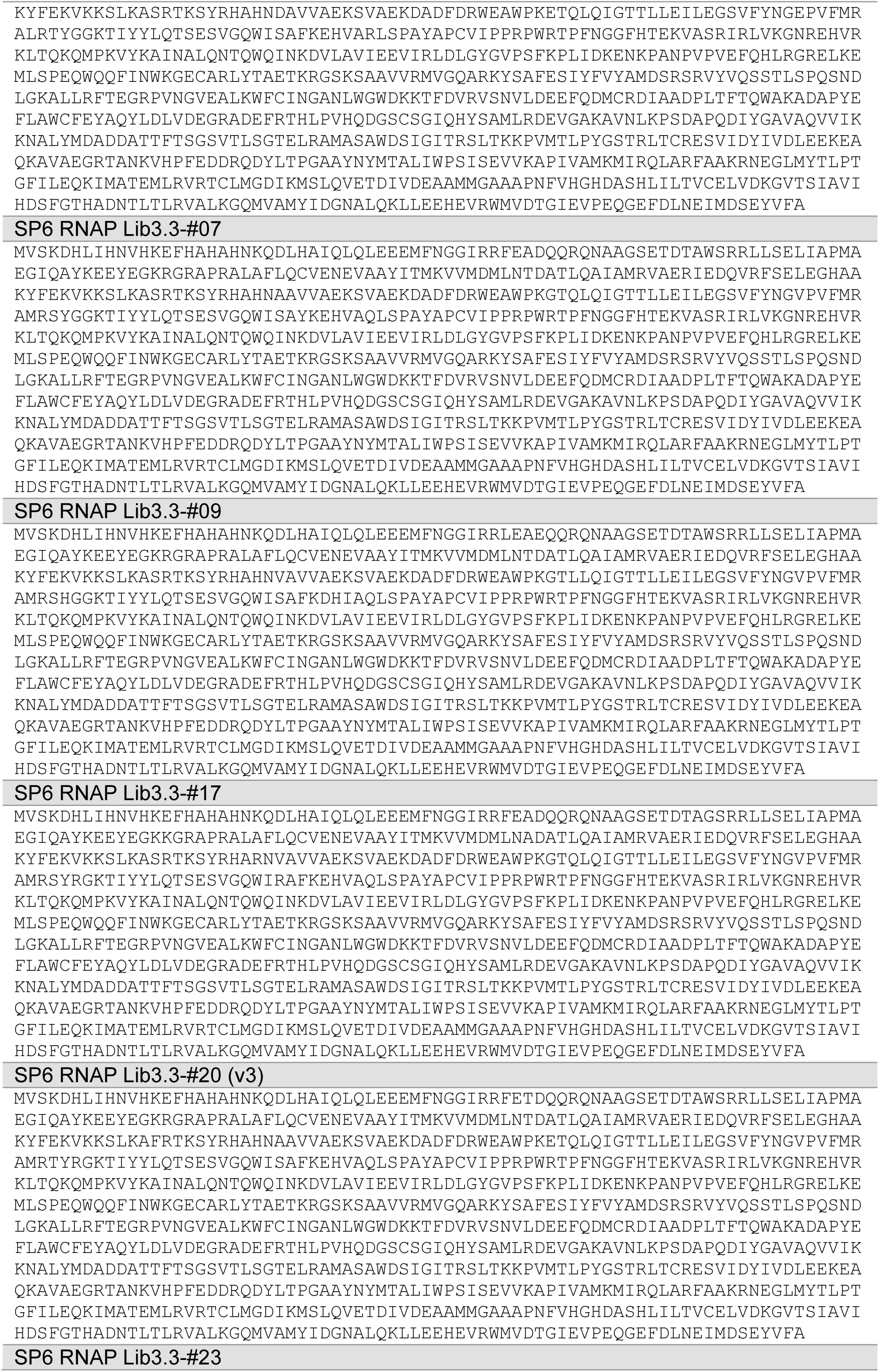
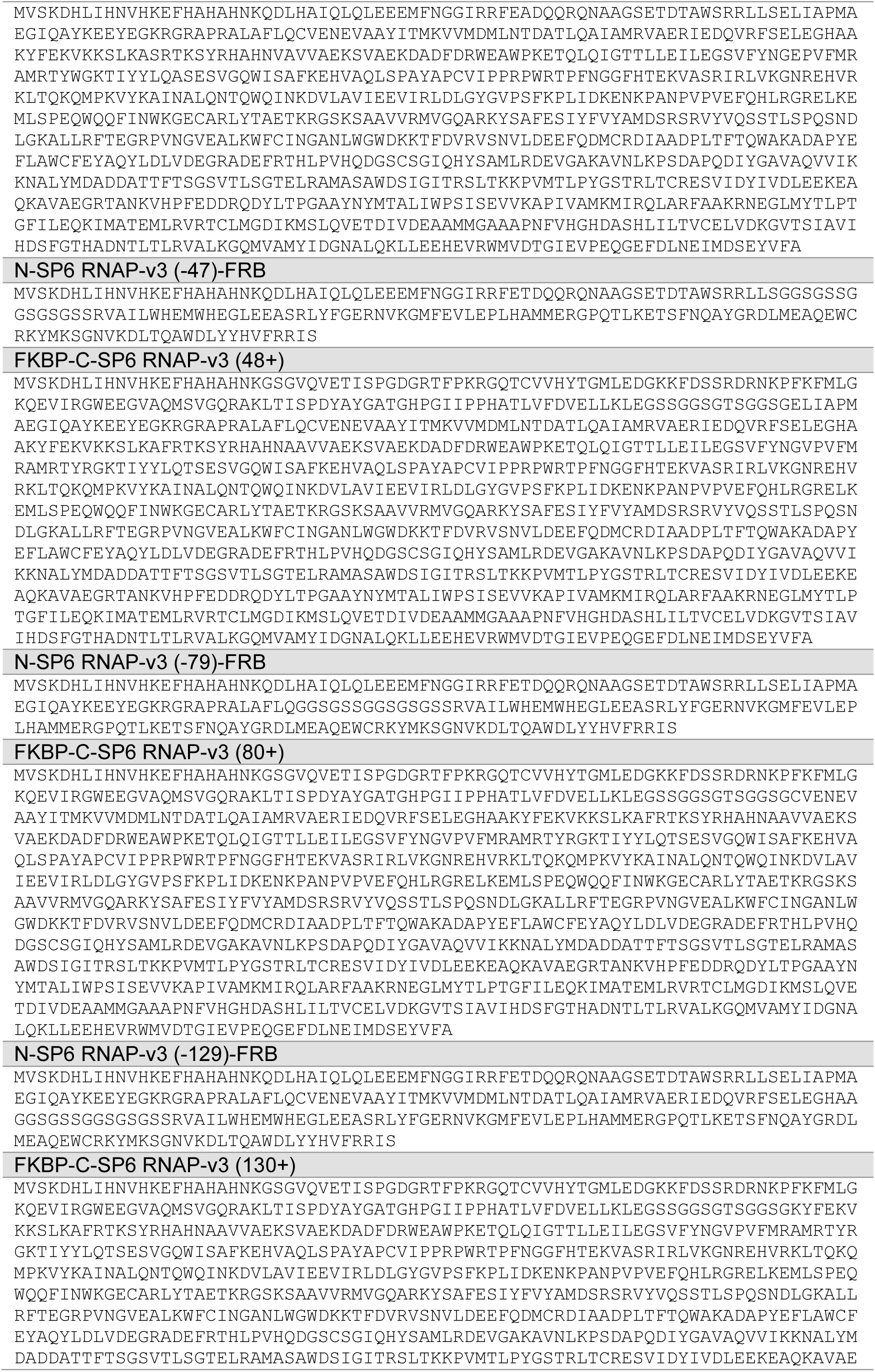
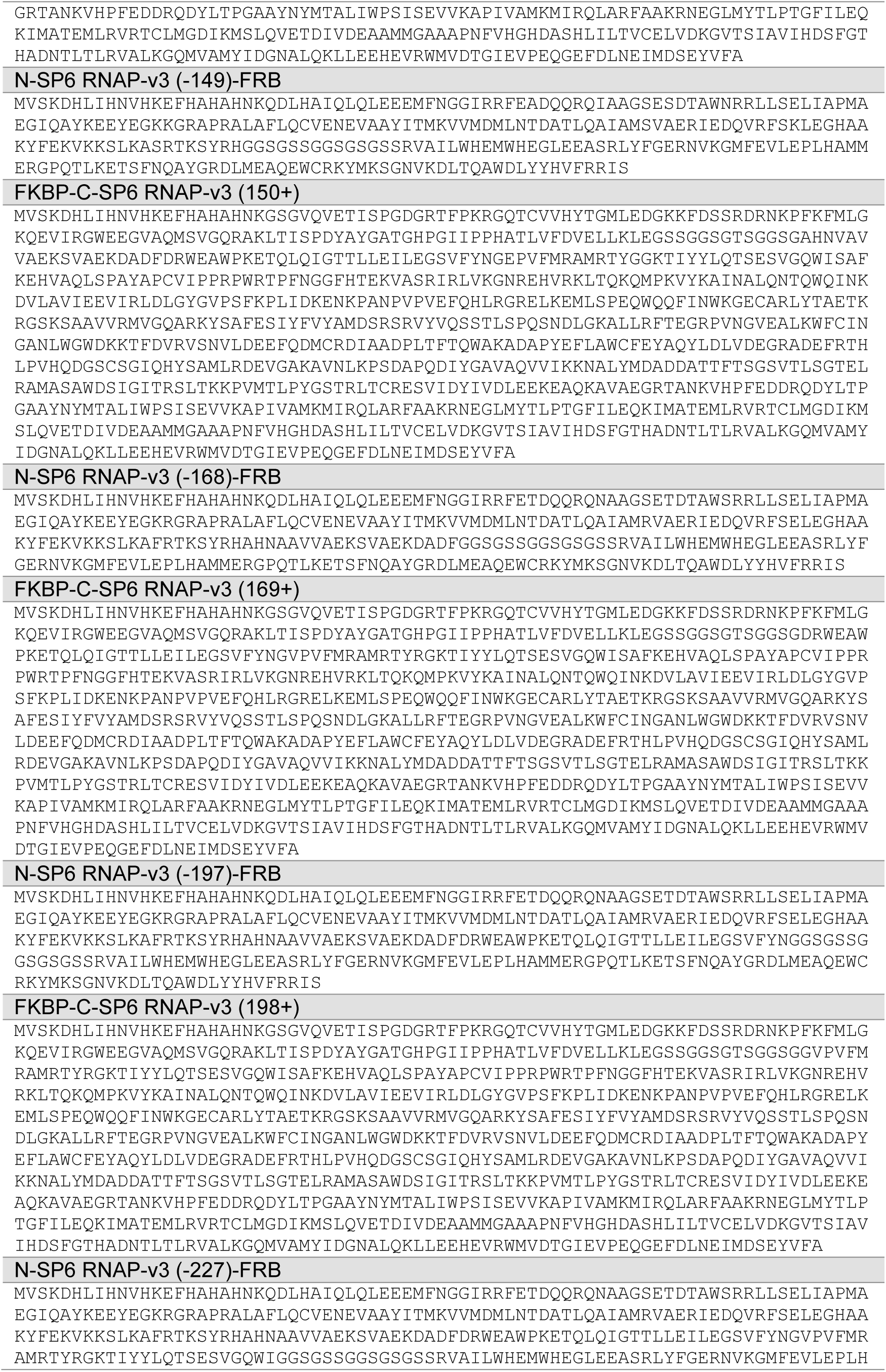
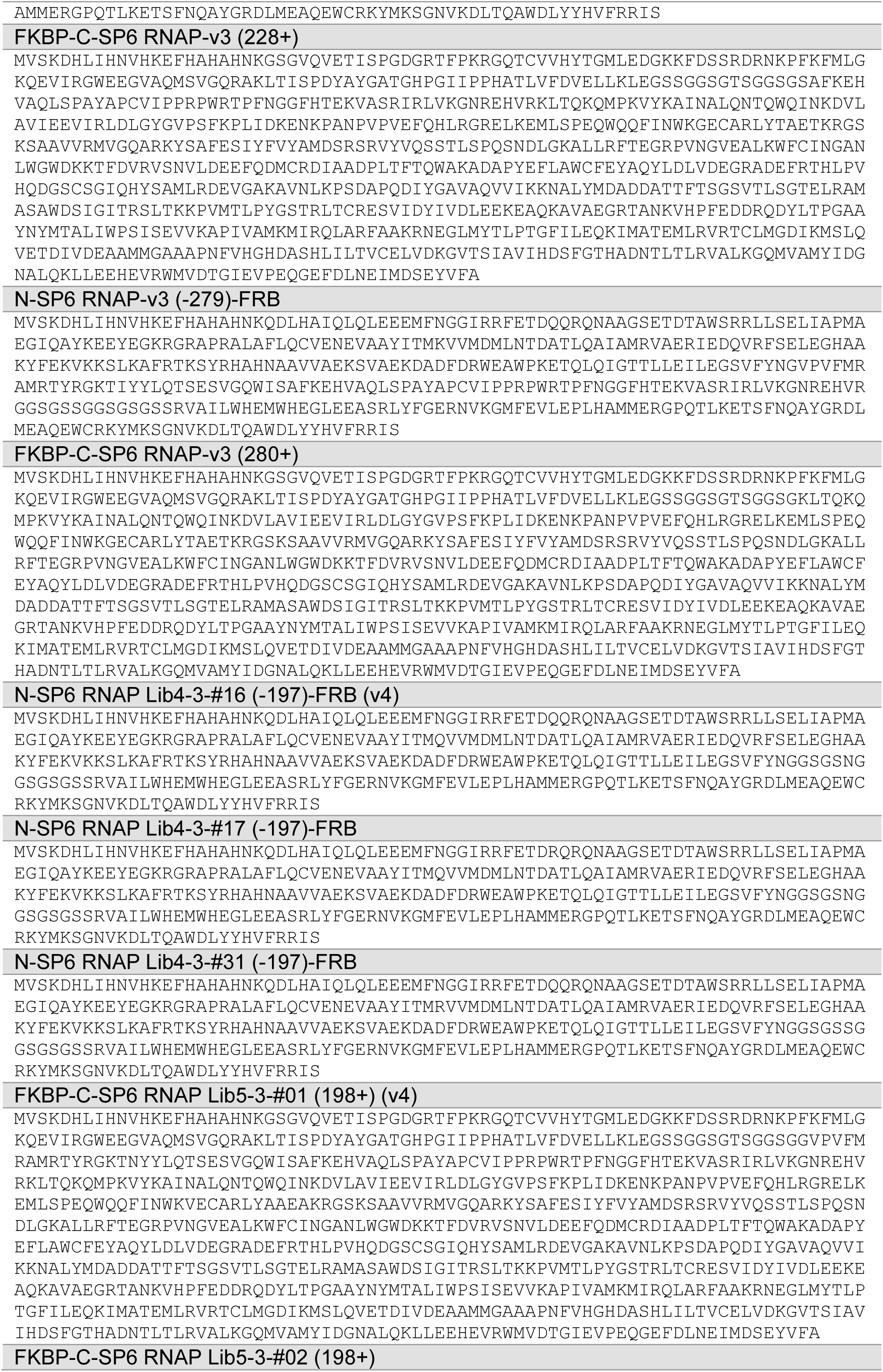
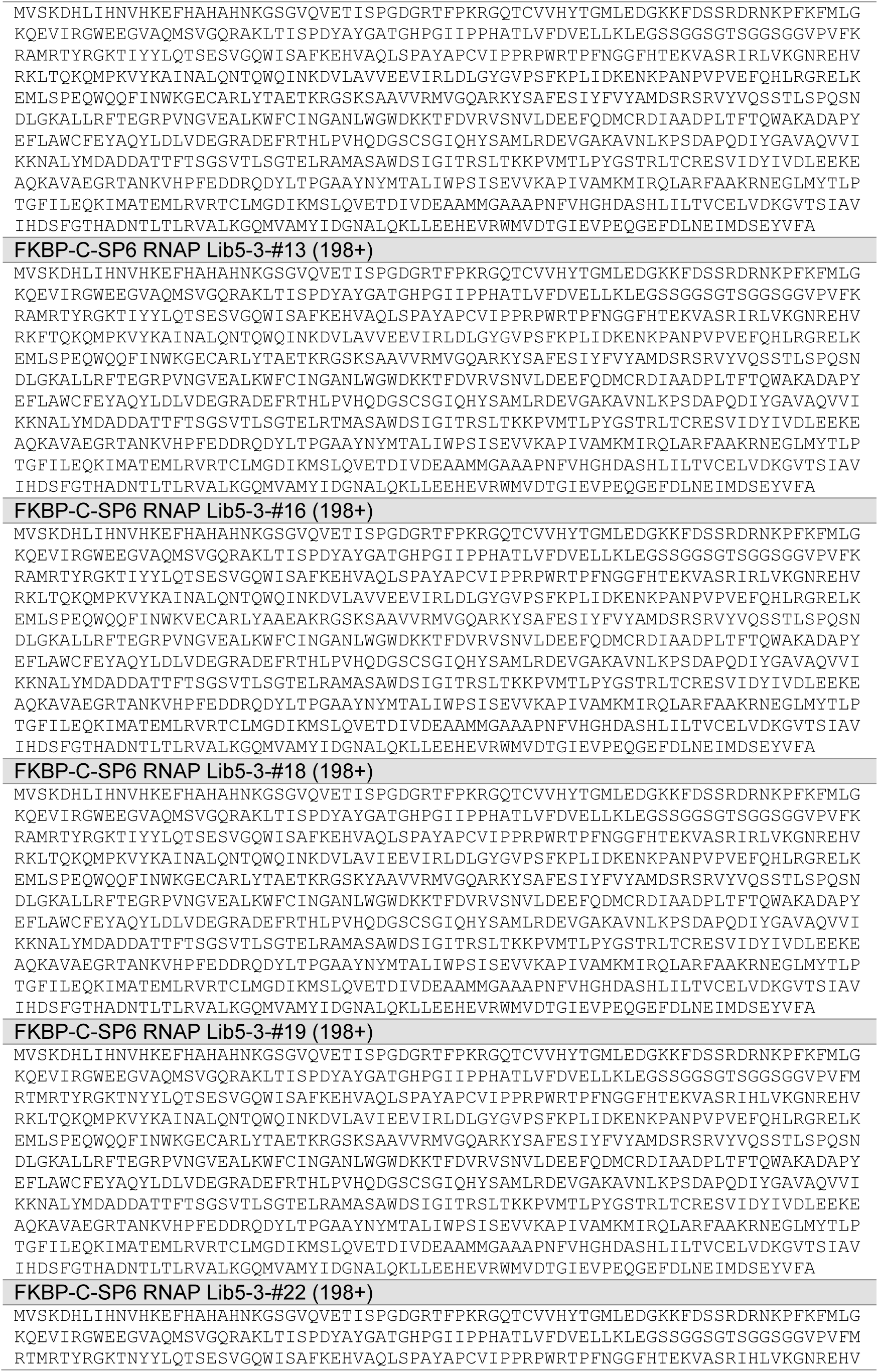
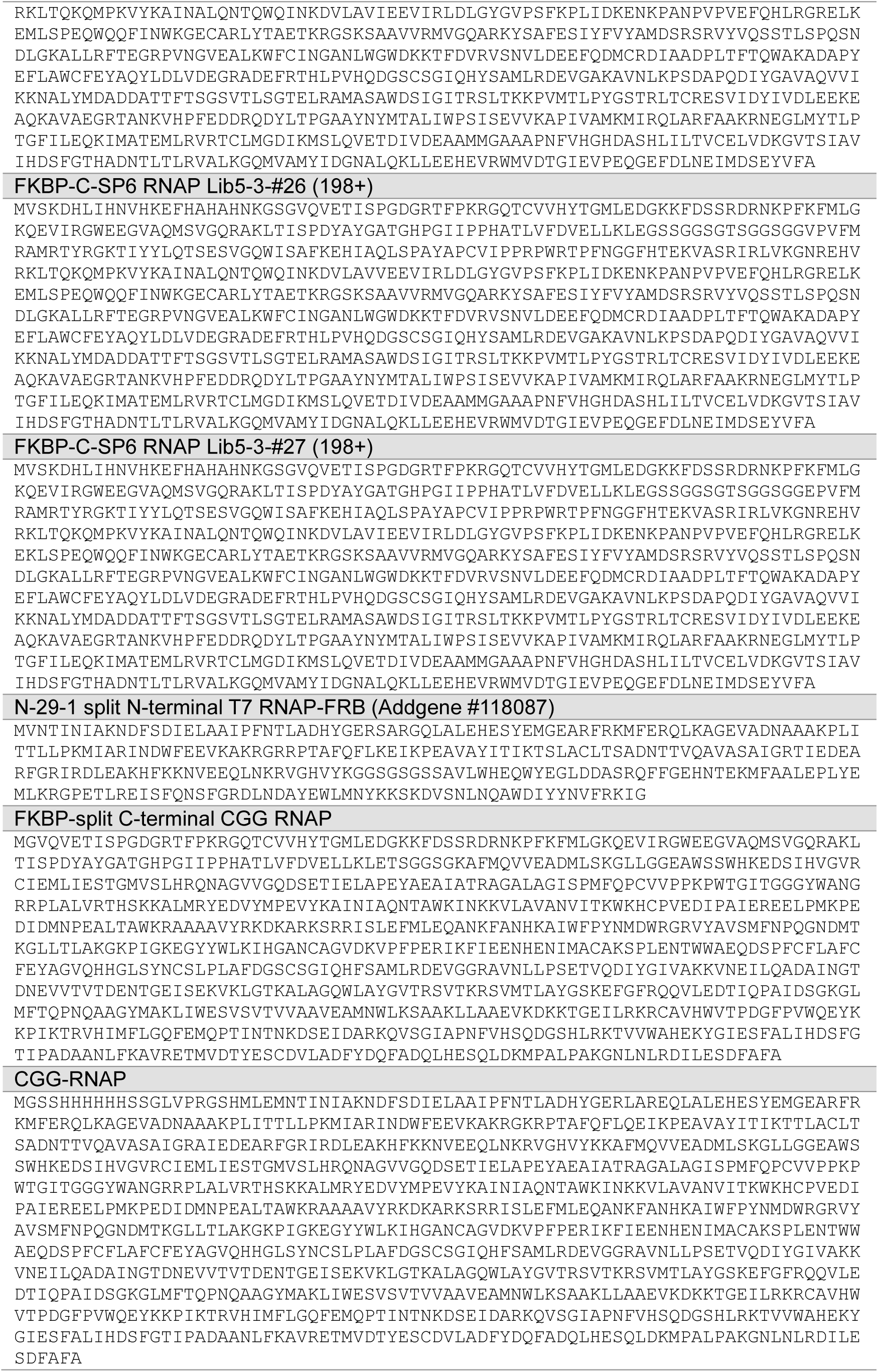
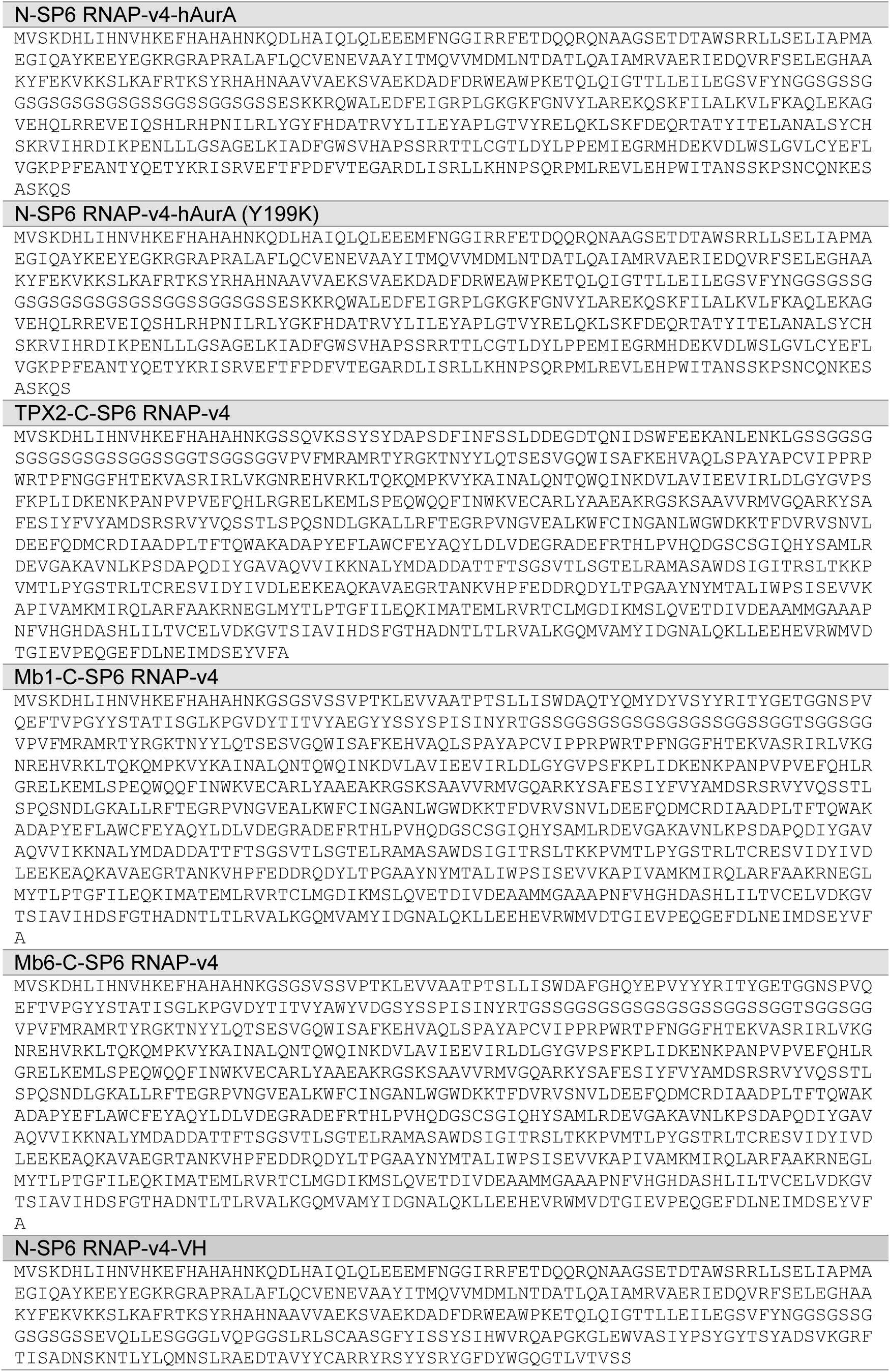
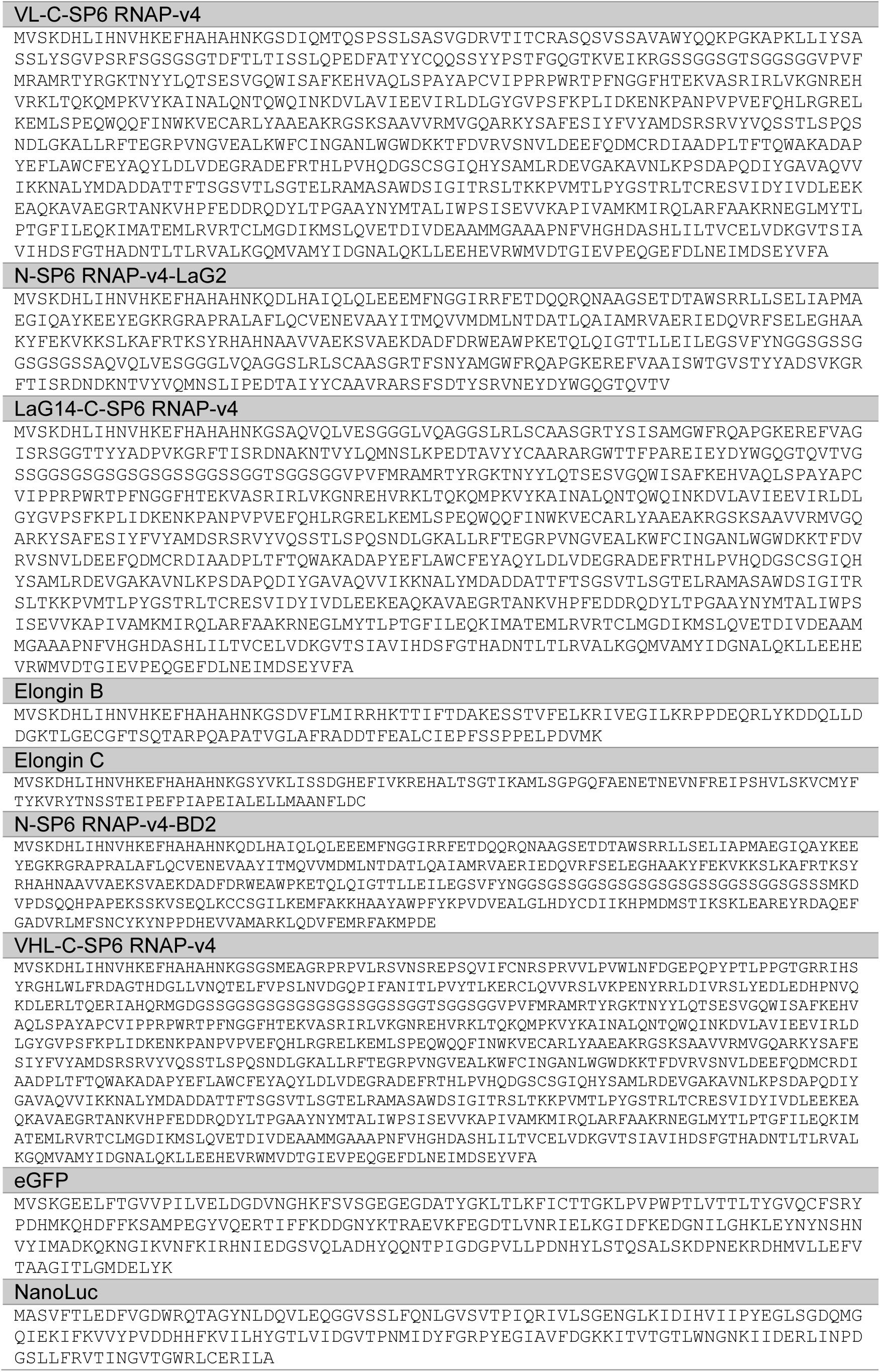
Amino acid sequences of individual proteins used in this study.

**Supplementary Table 3:**
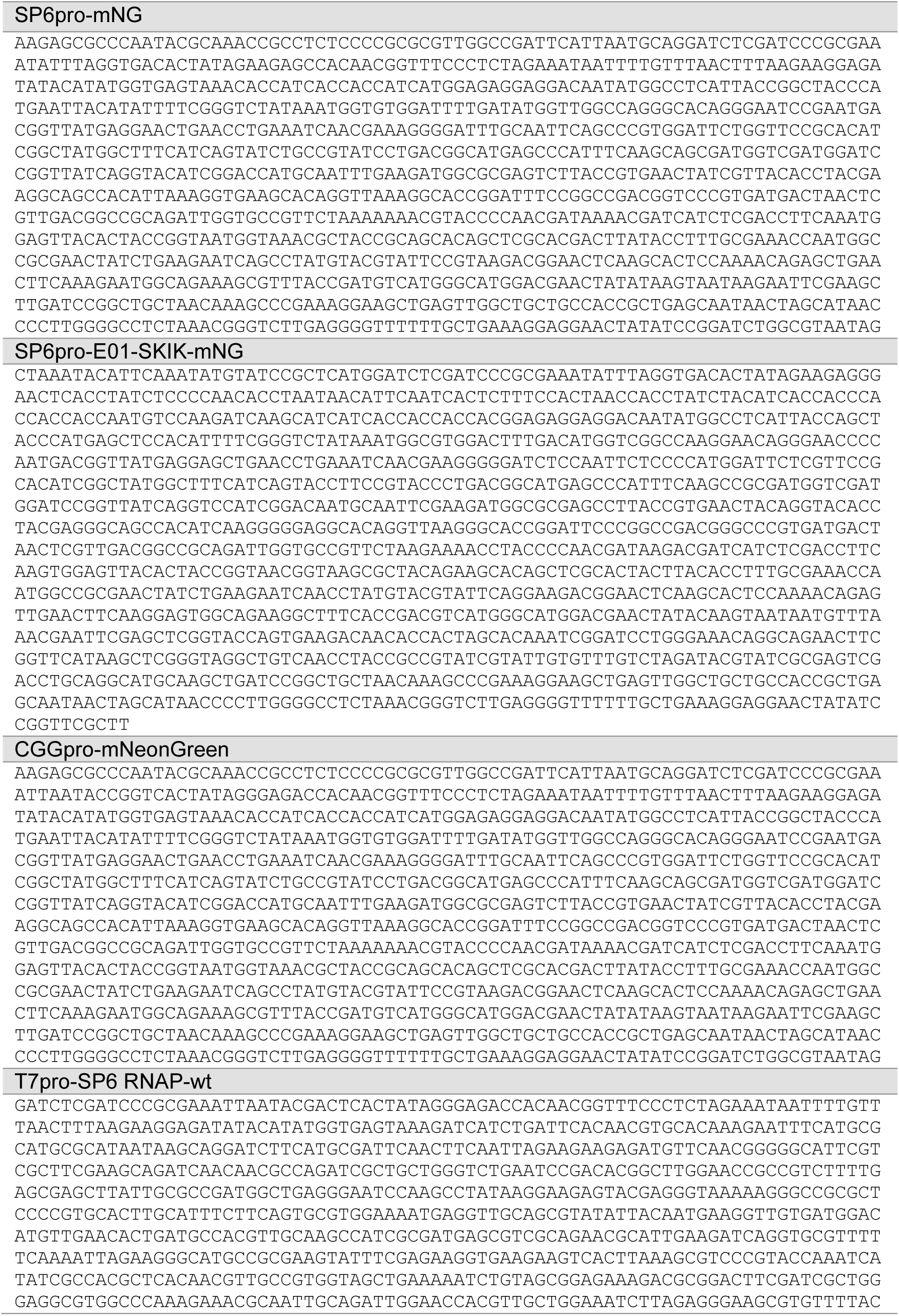
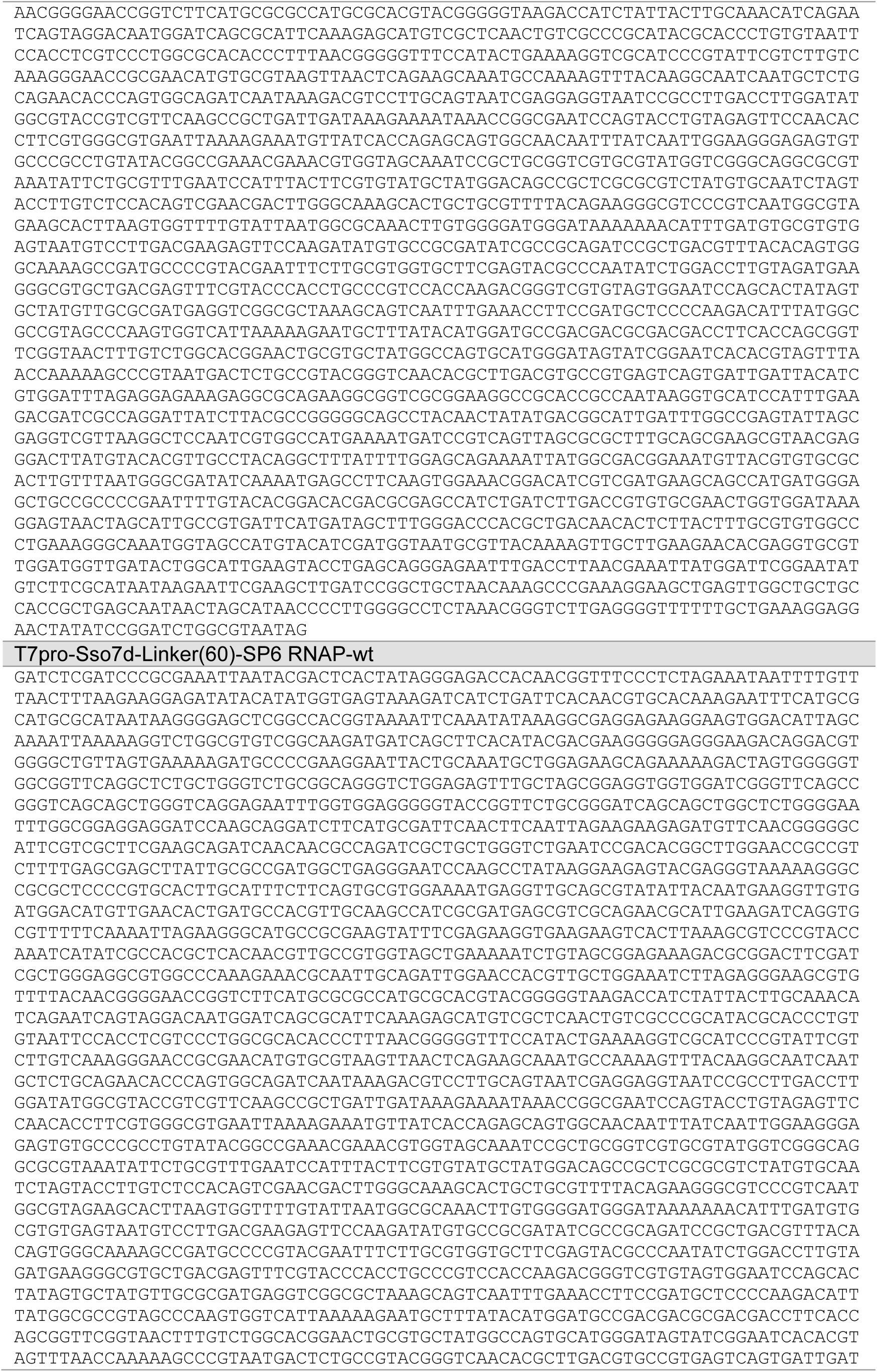
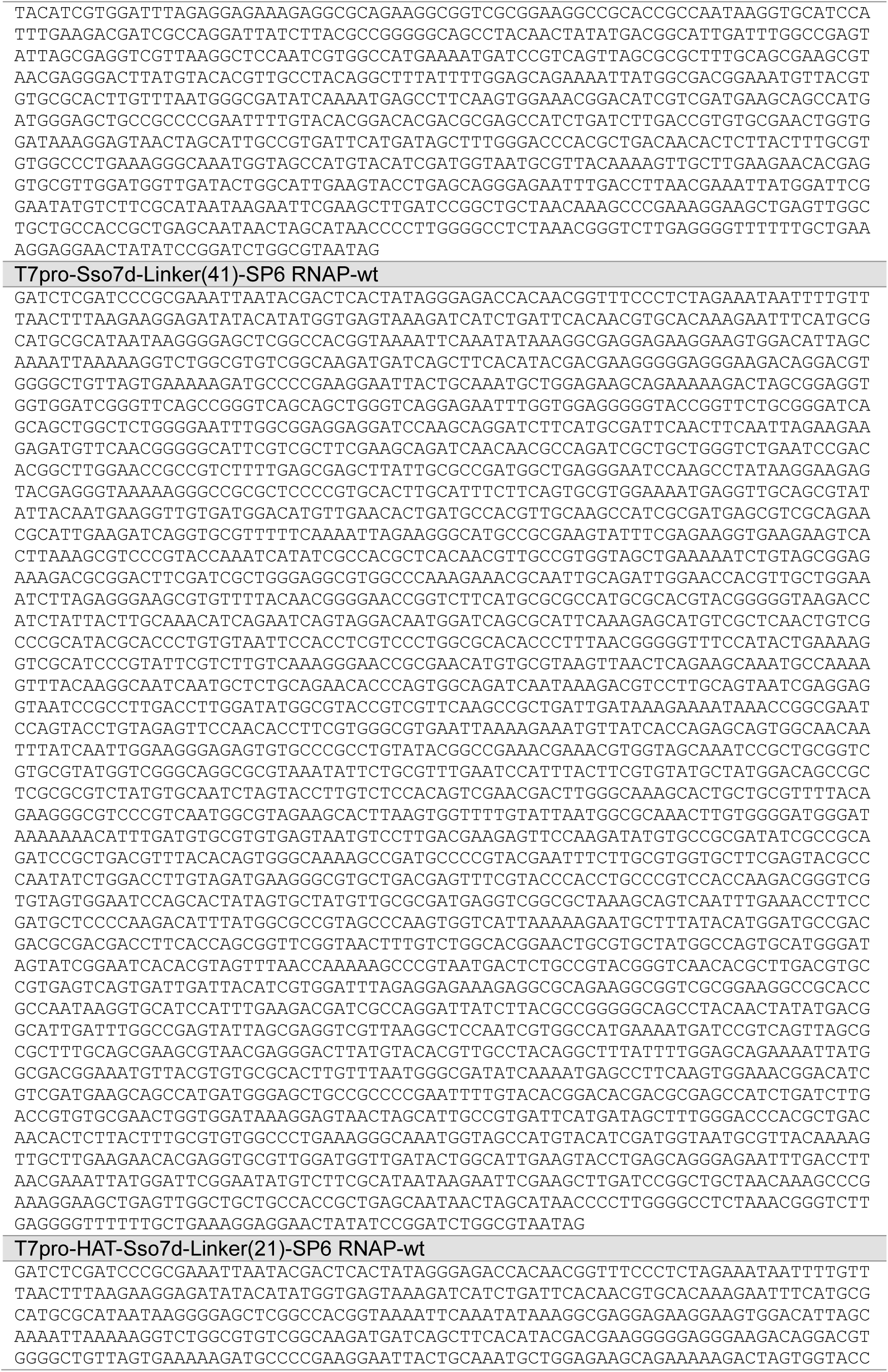
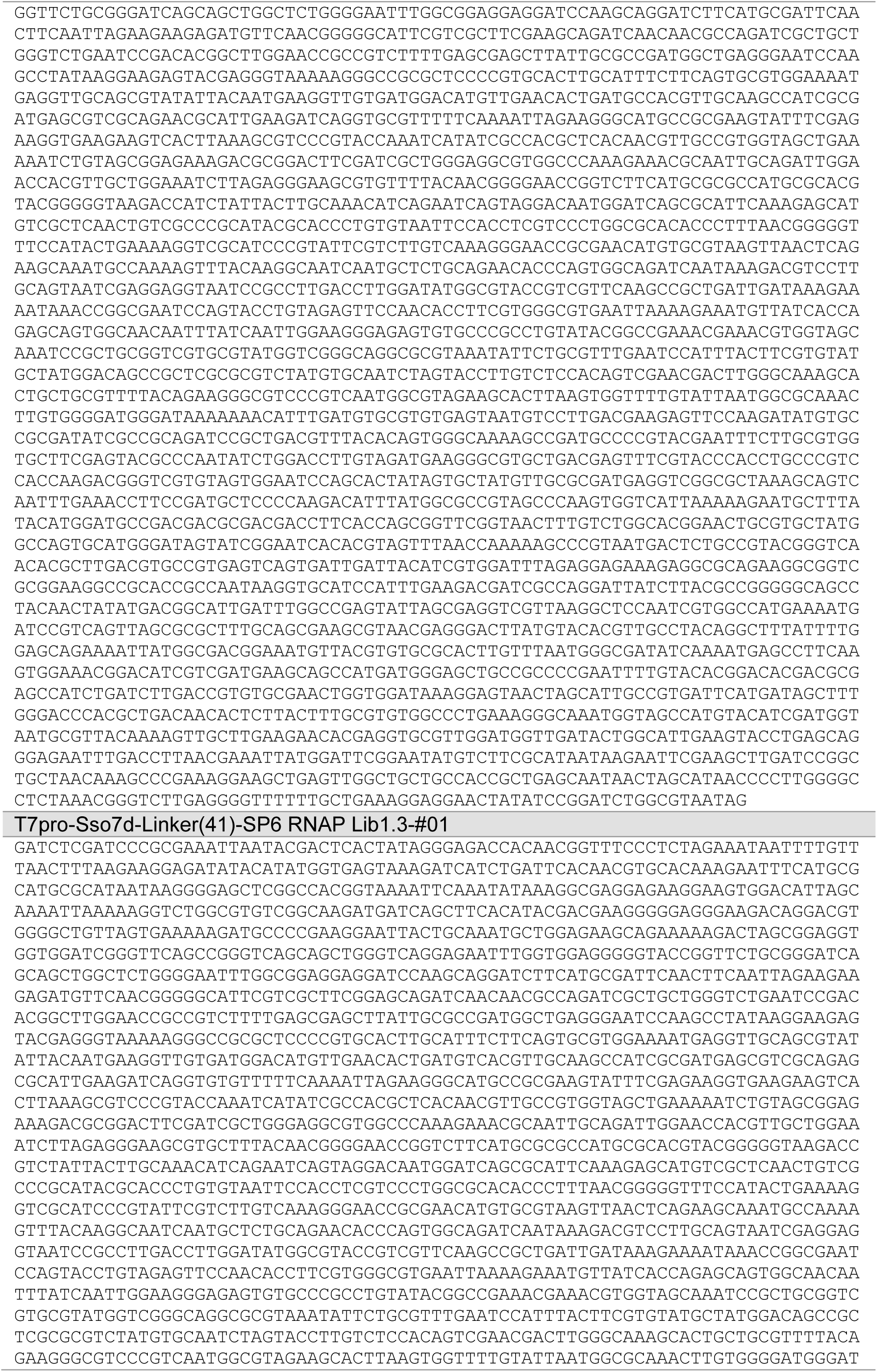
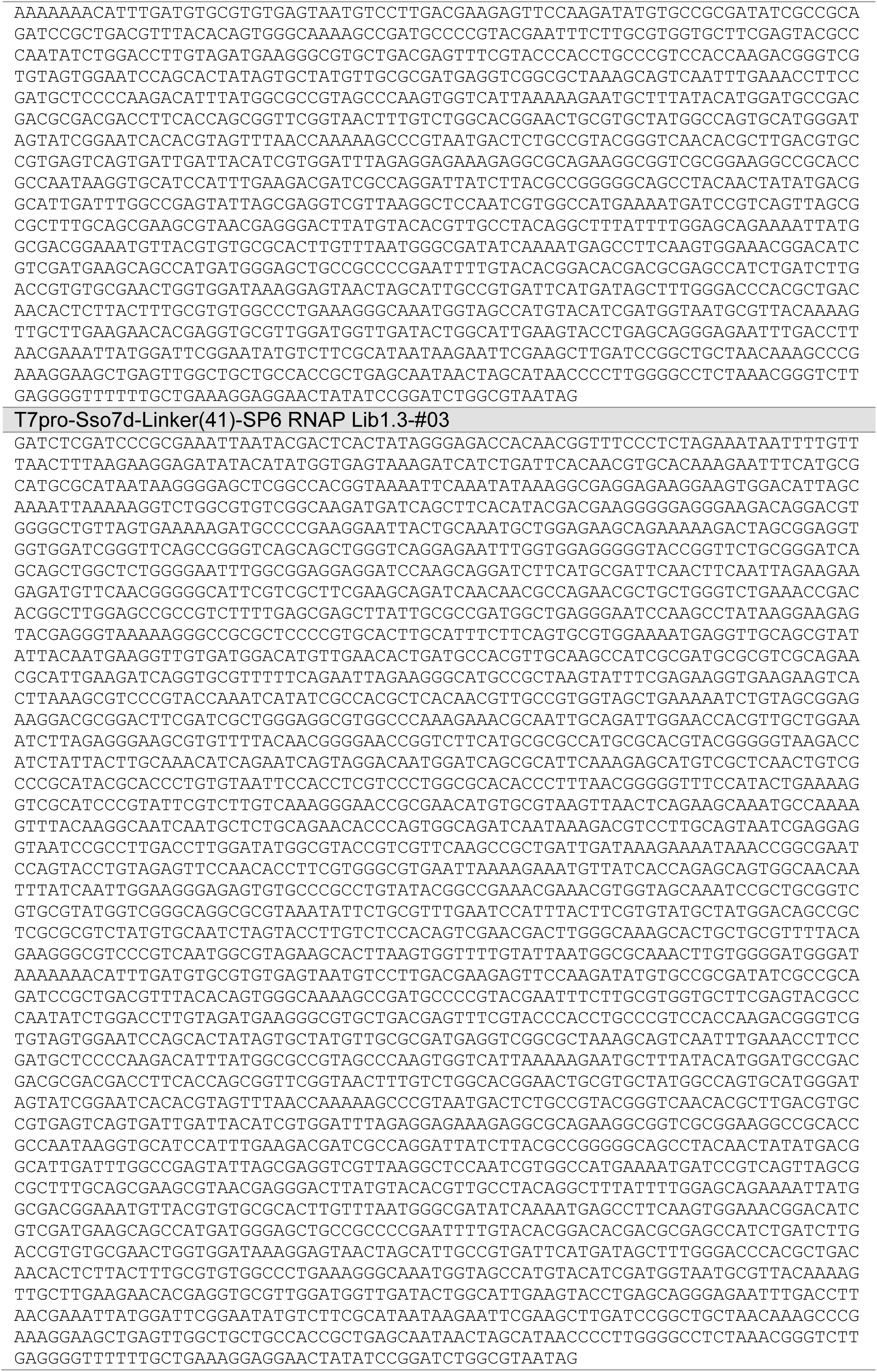
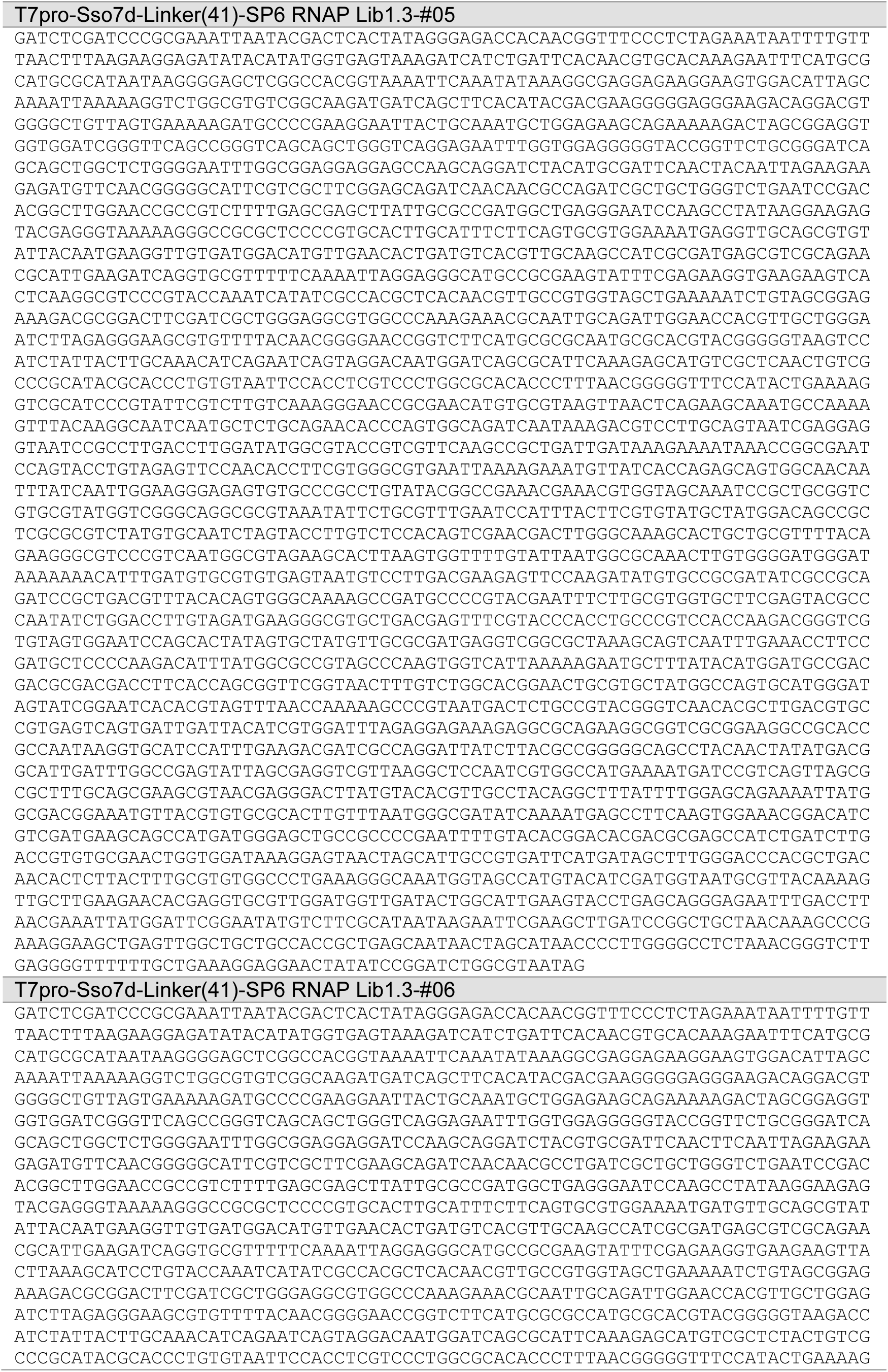
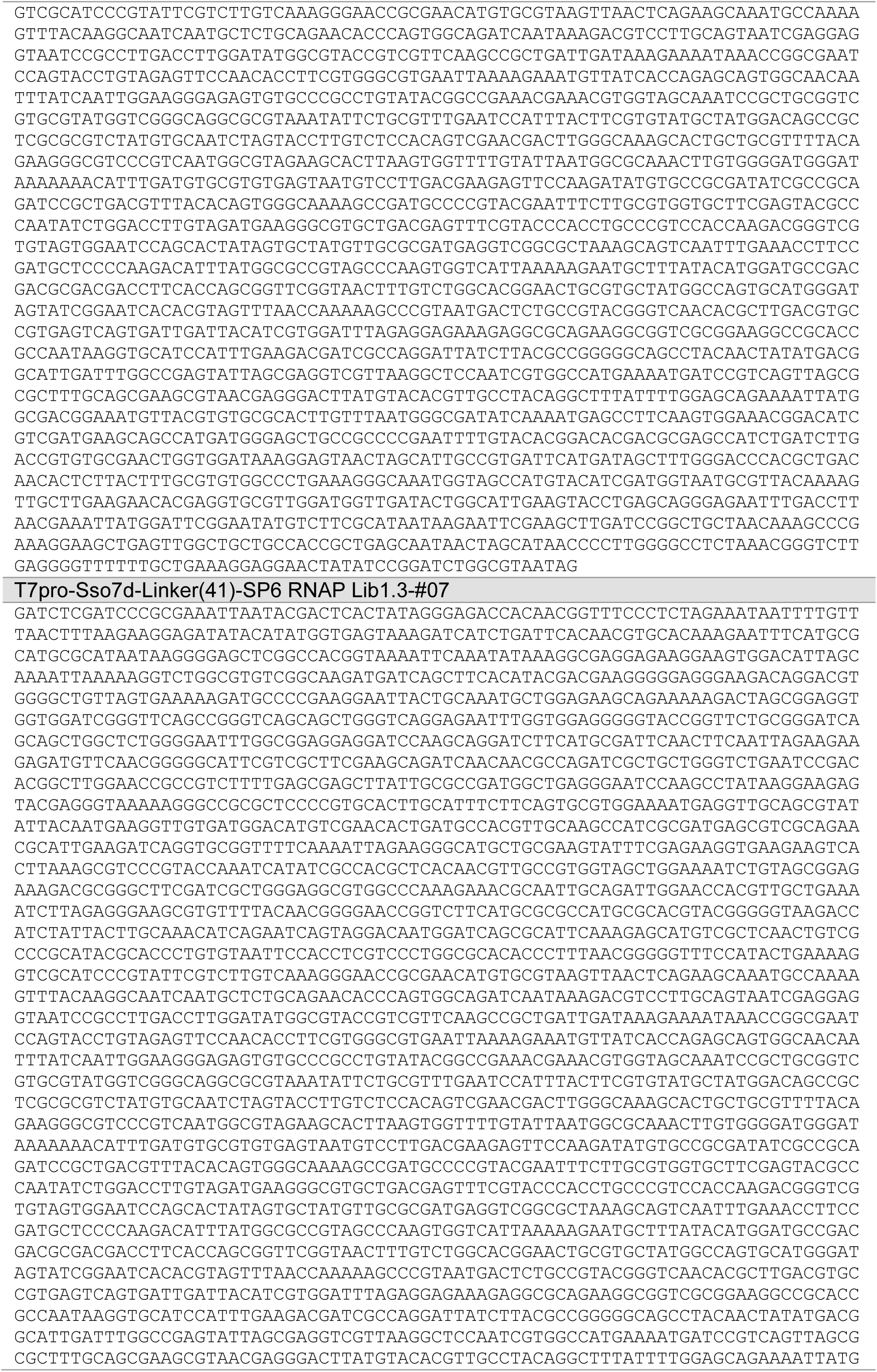
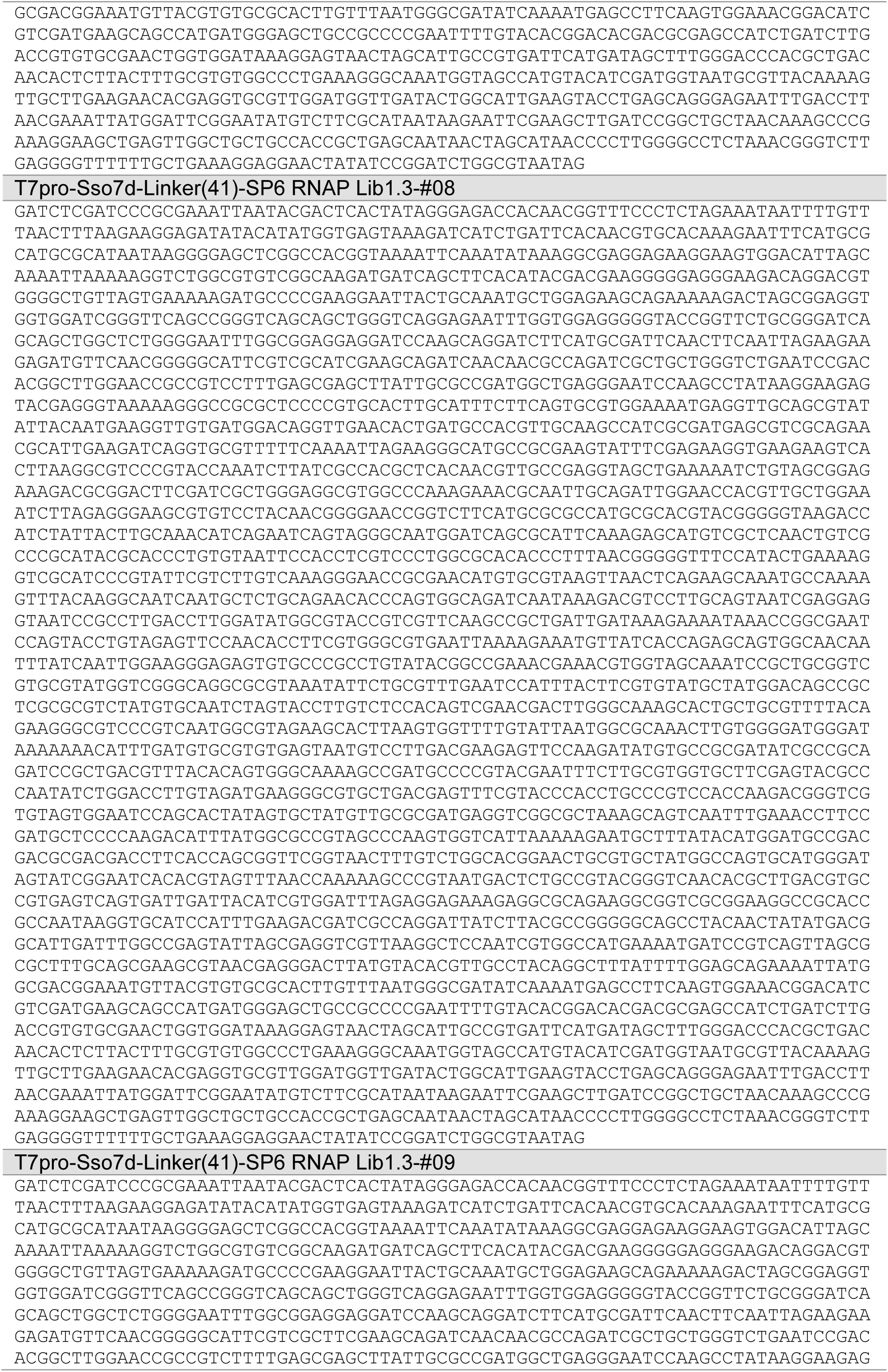
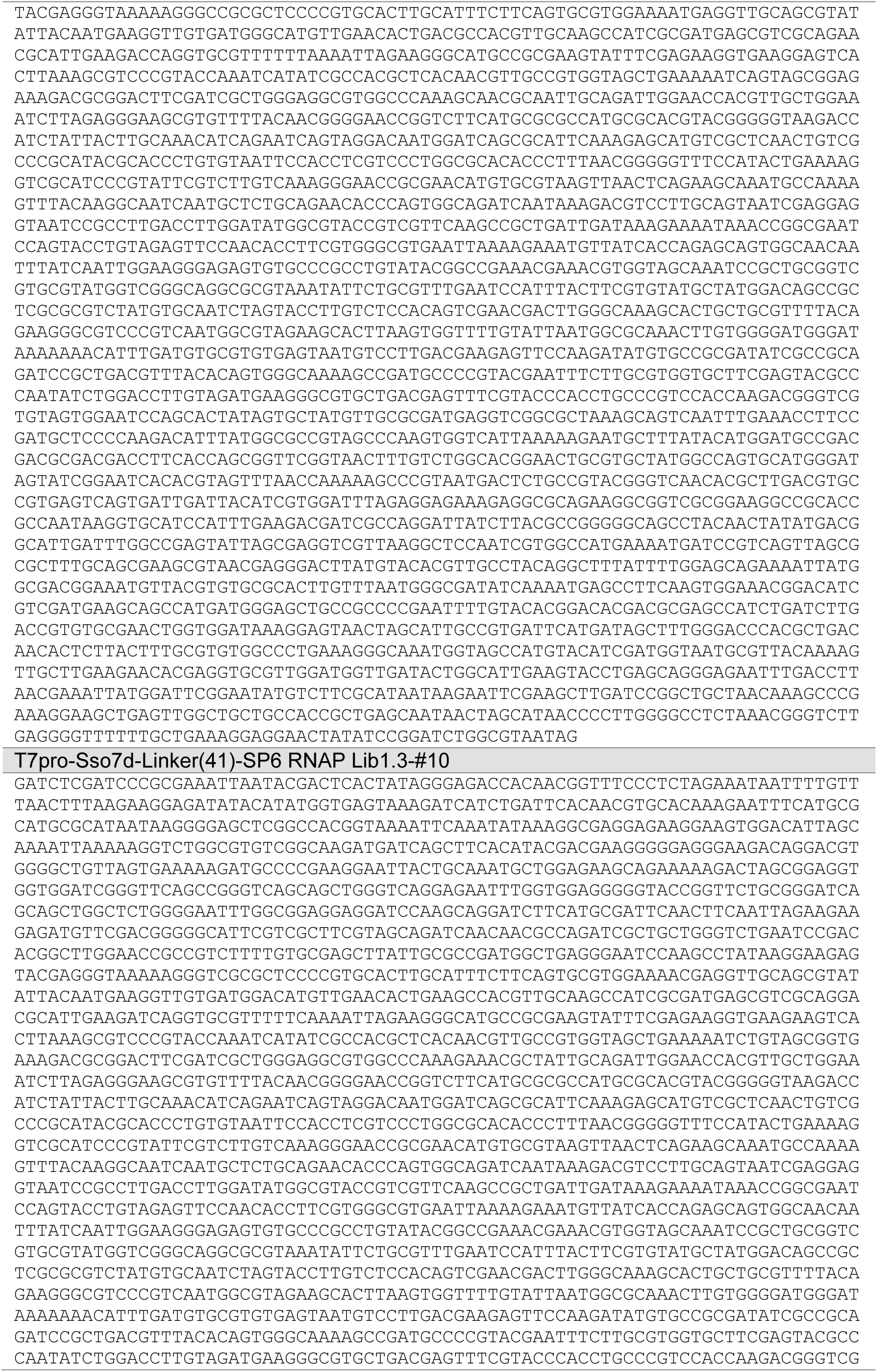
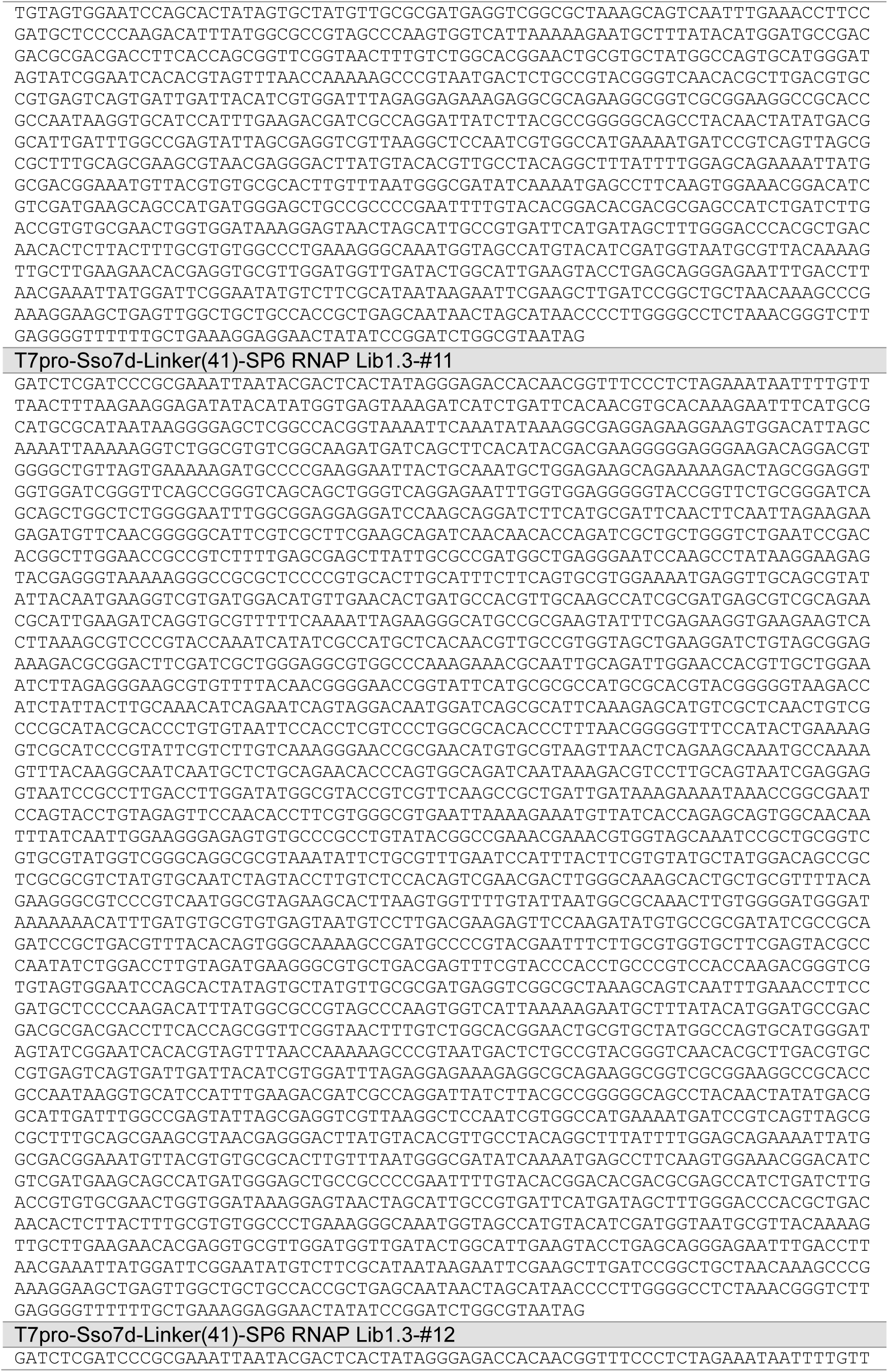
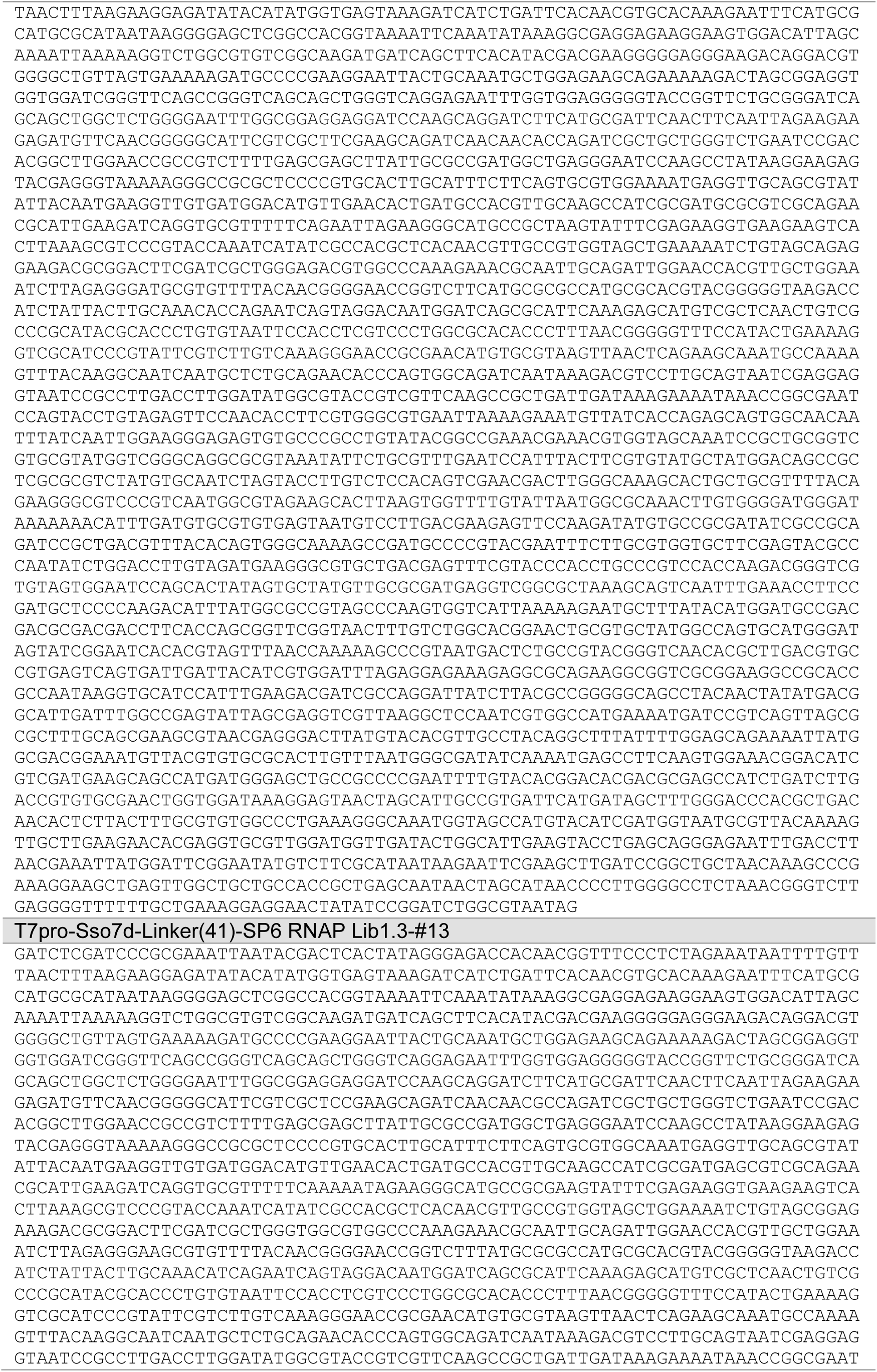
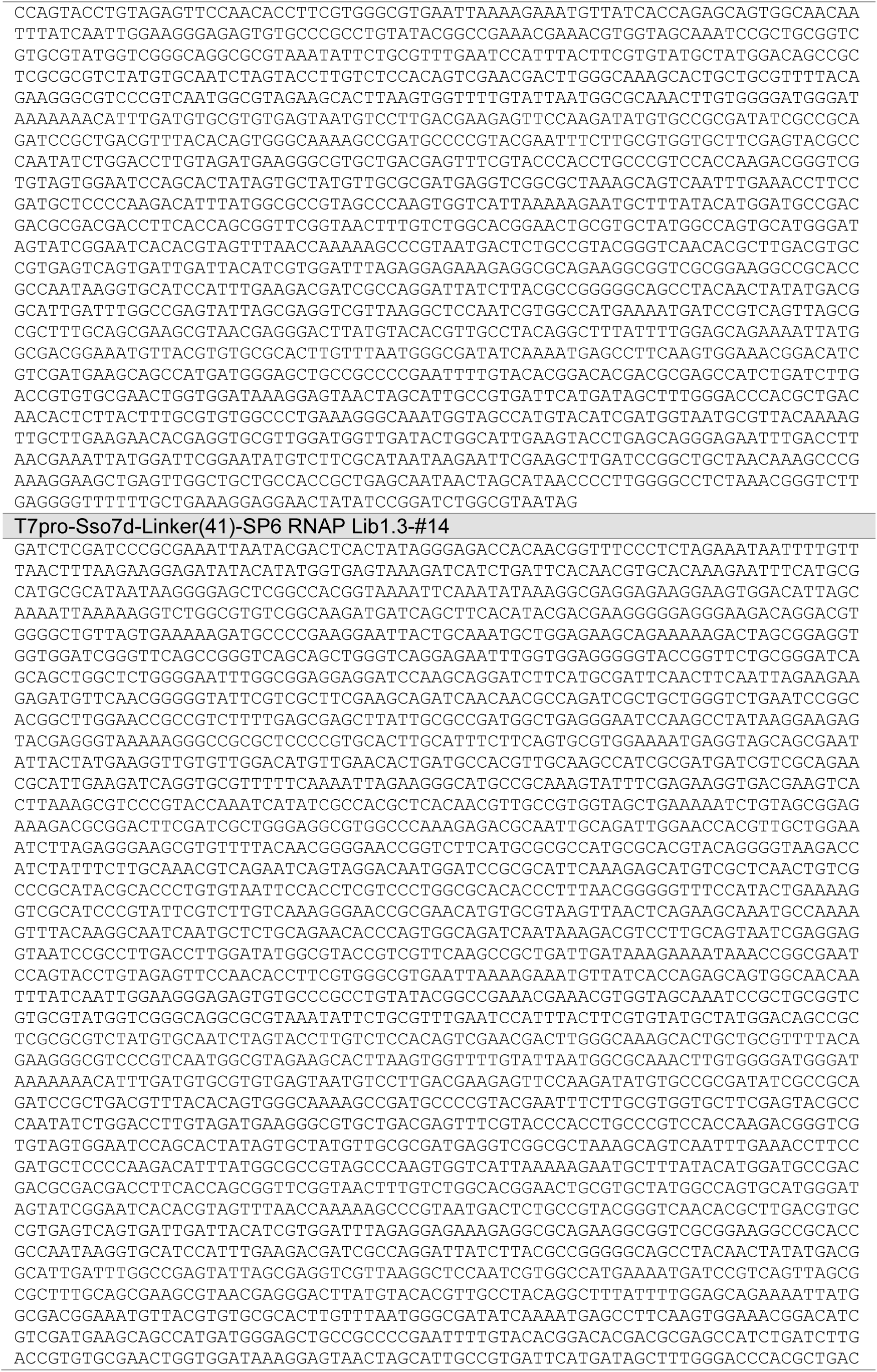
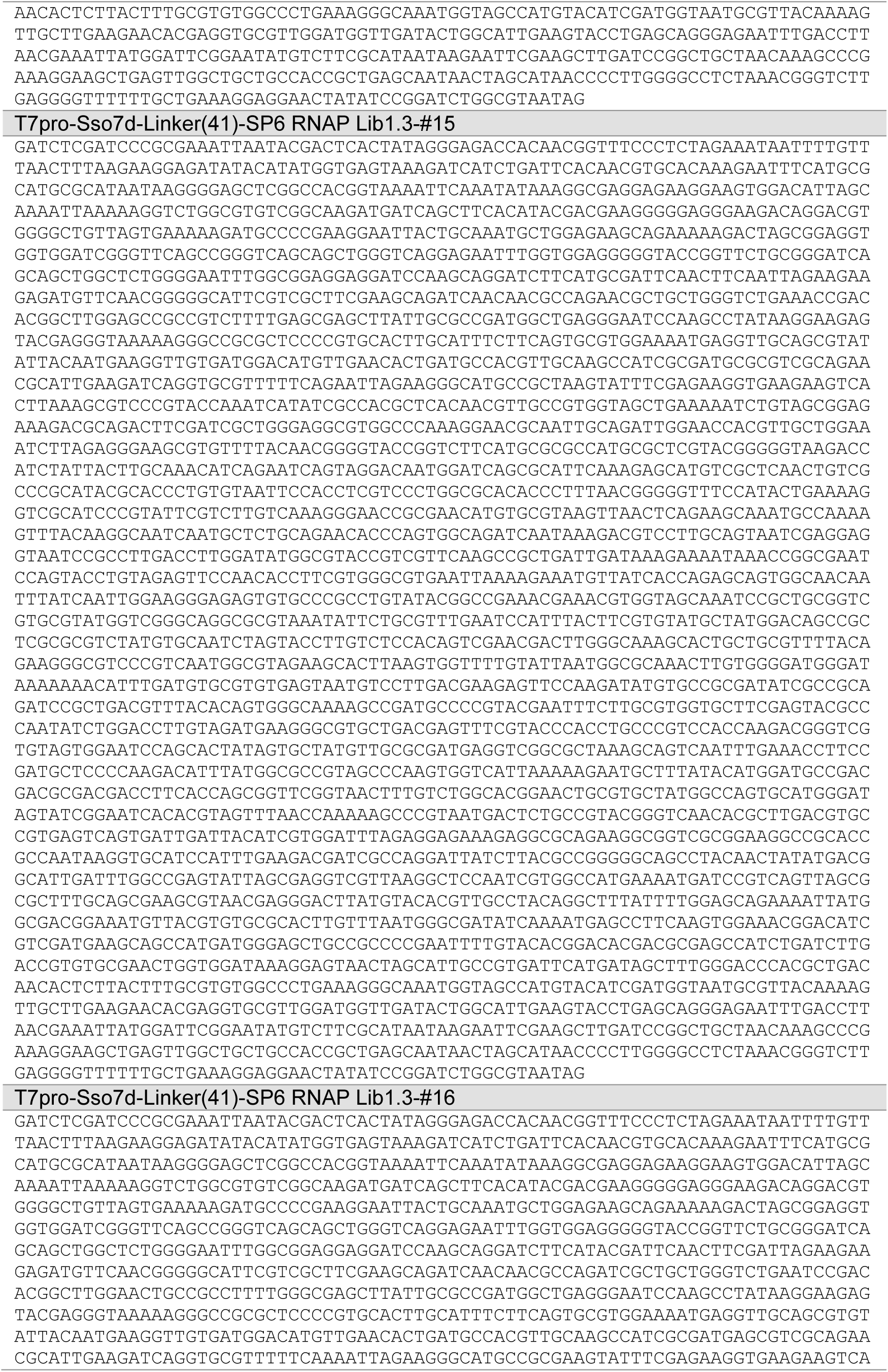
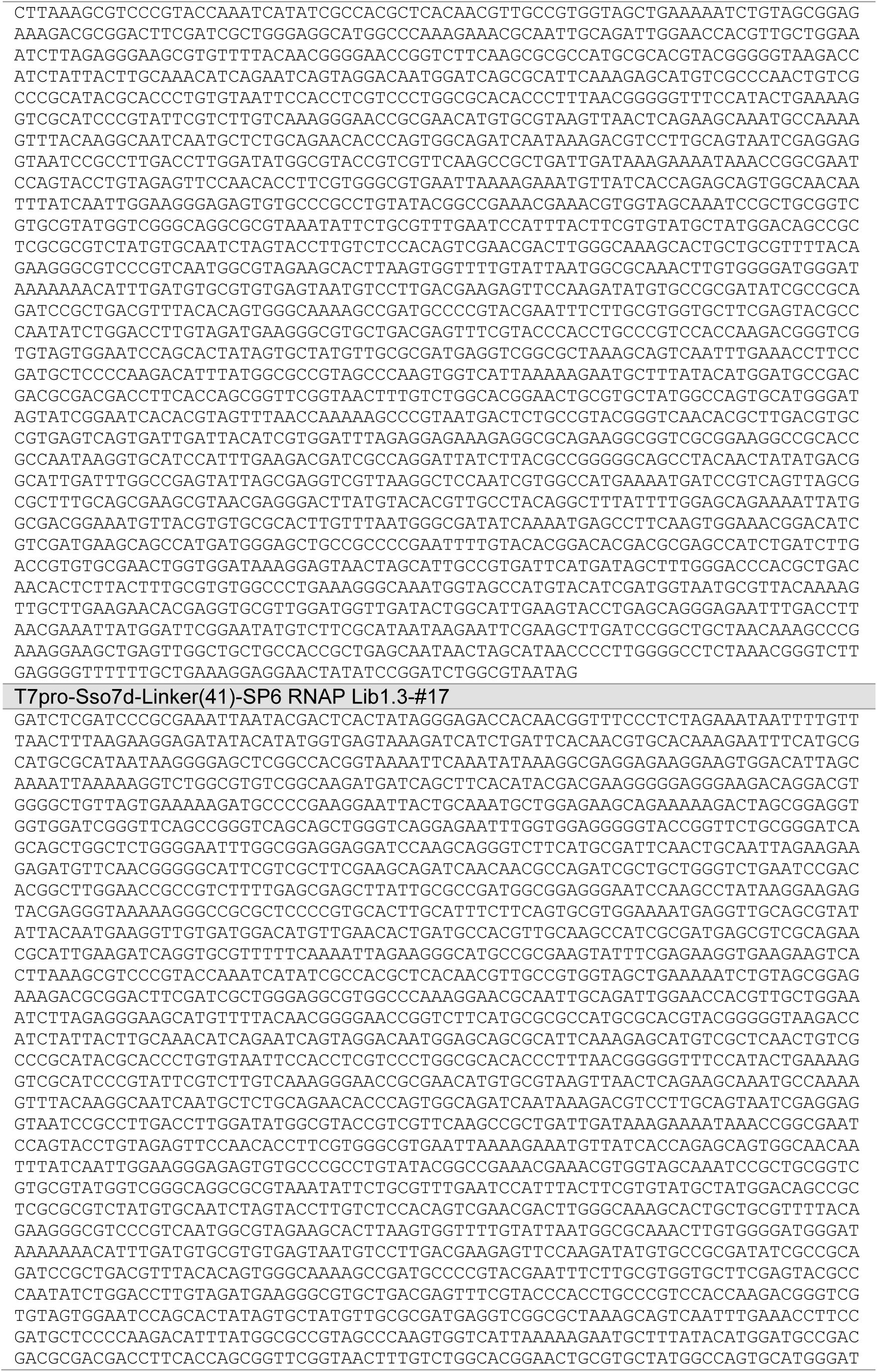
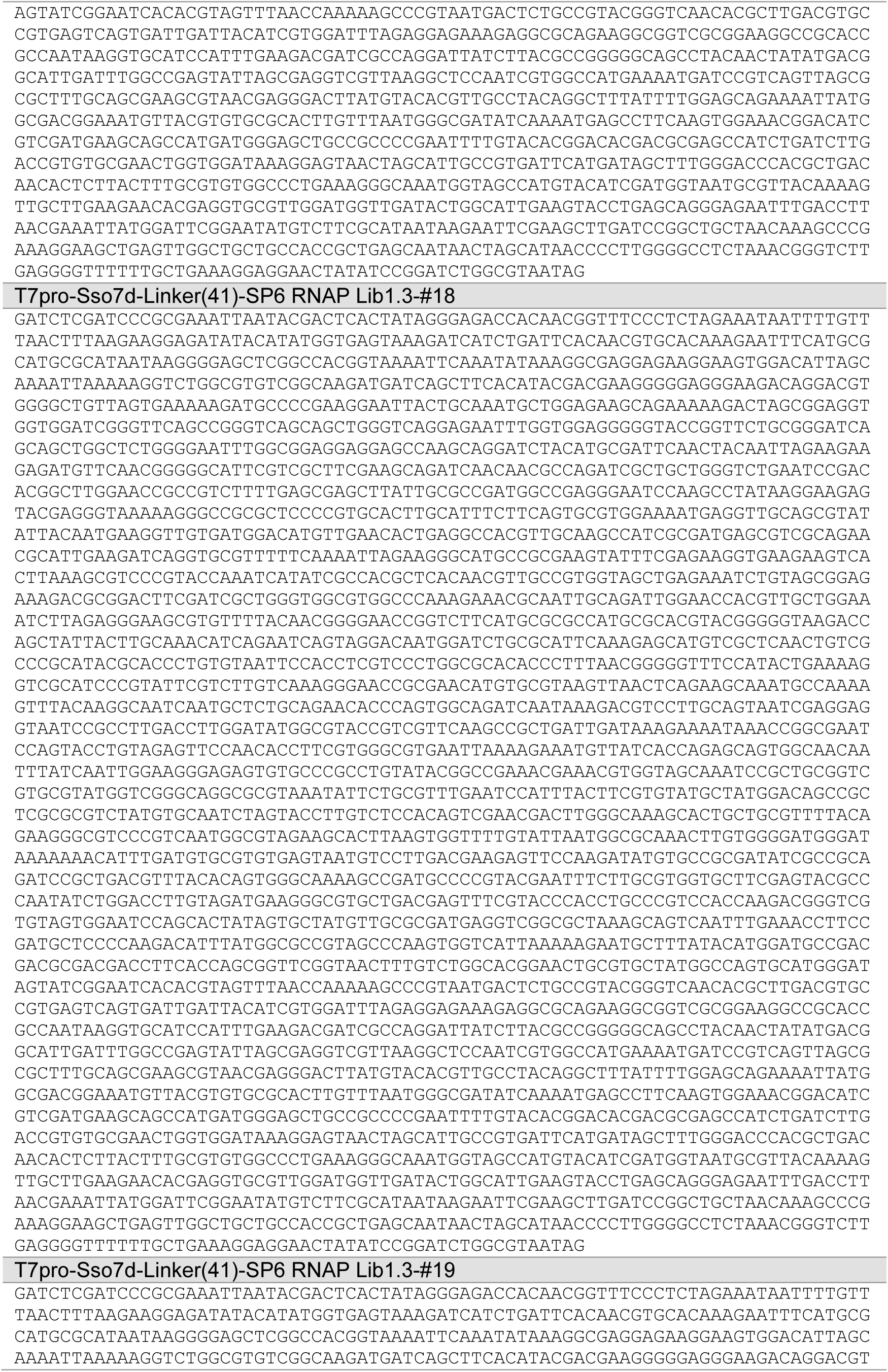
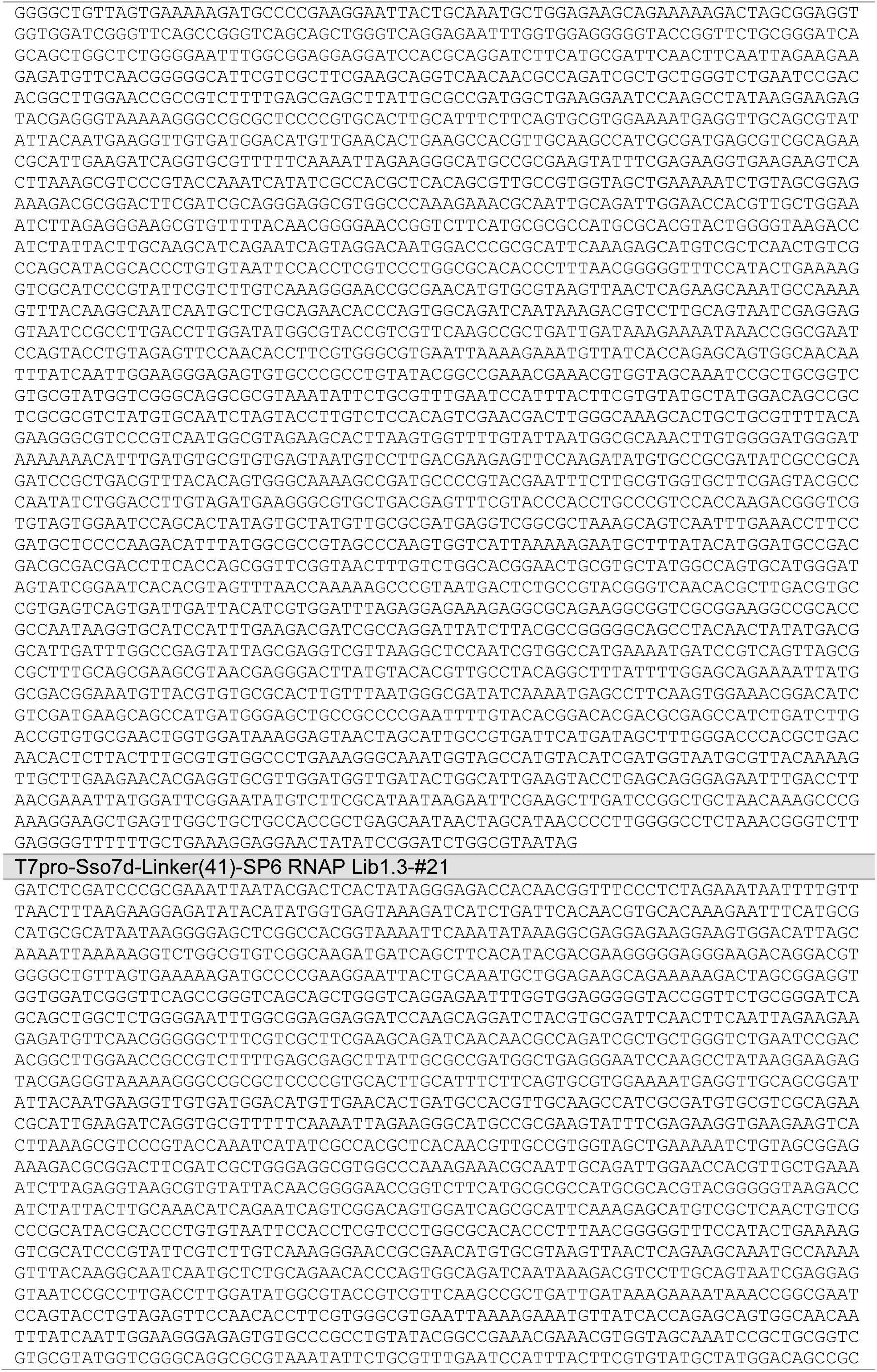
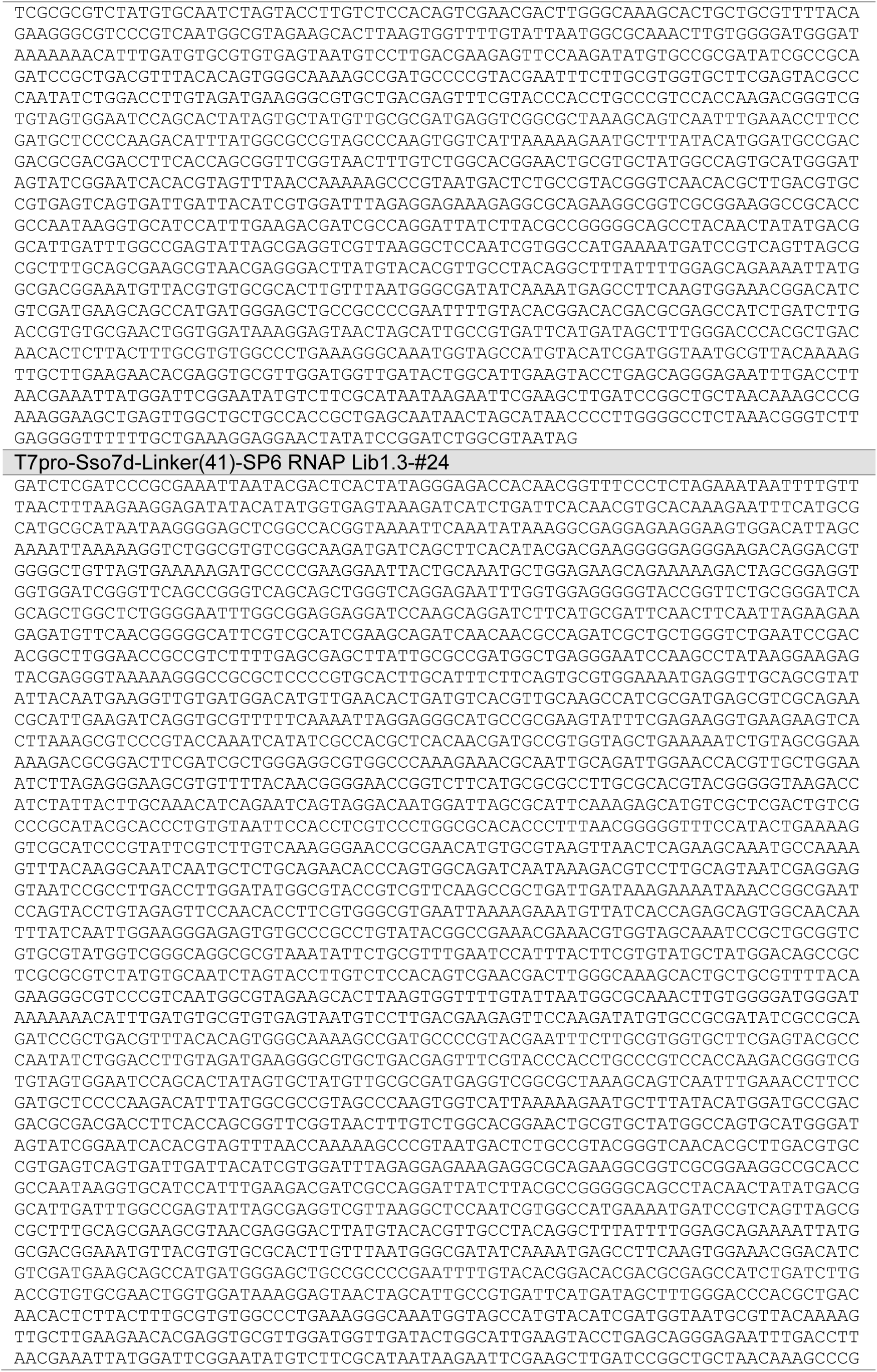
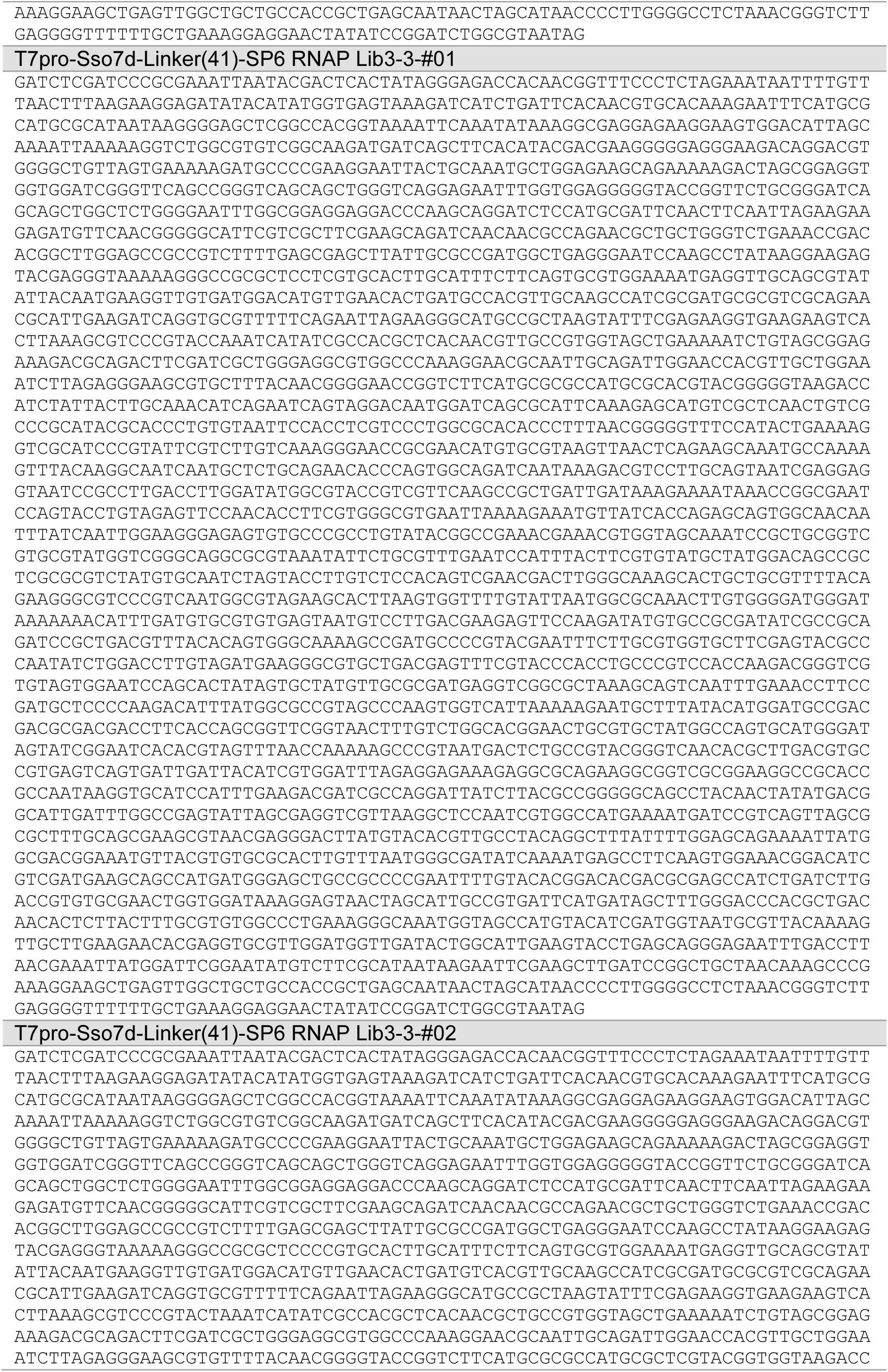
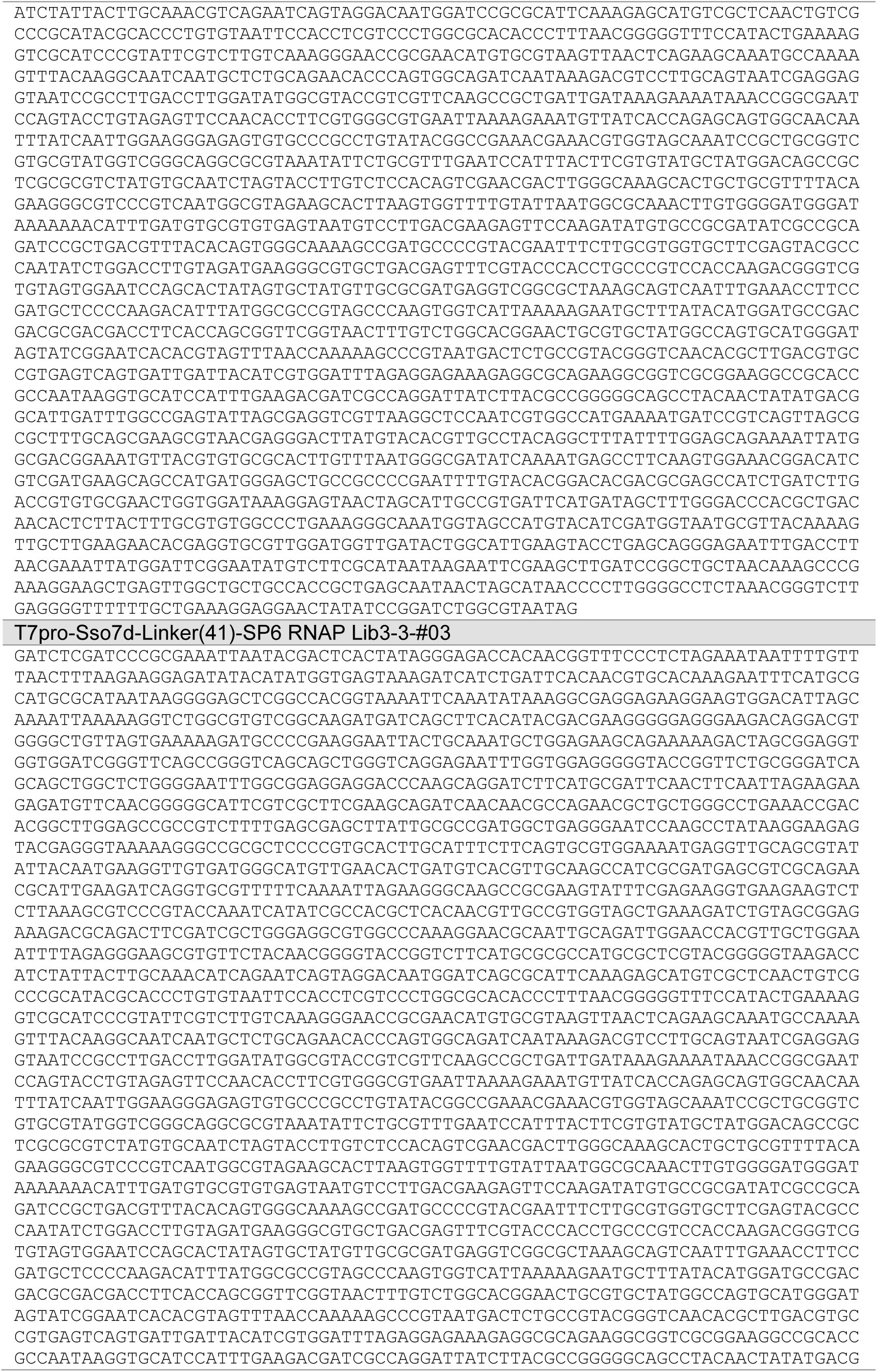
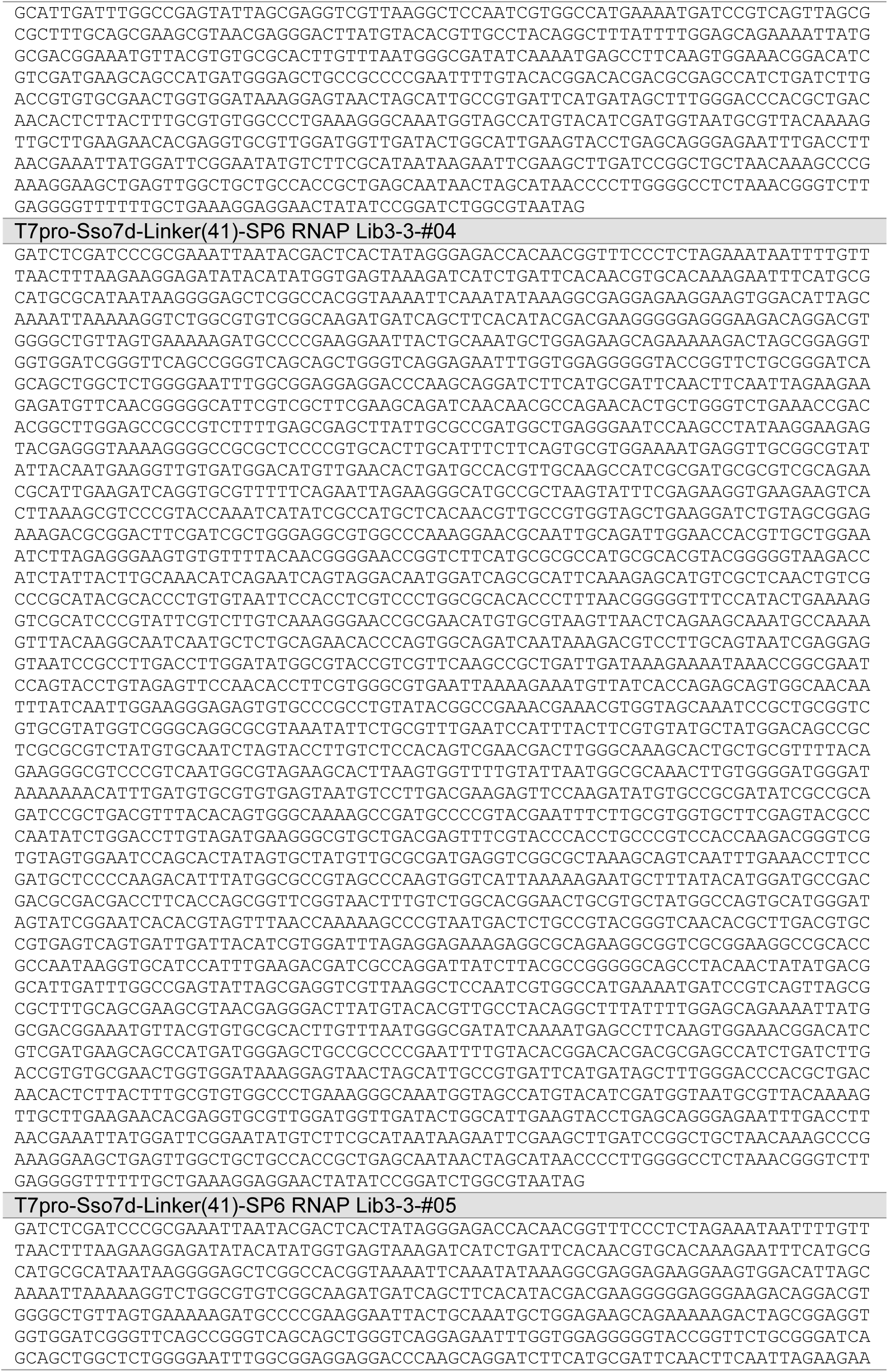
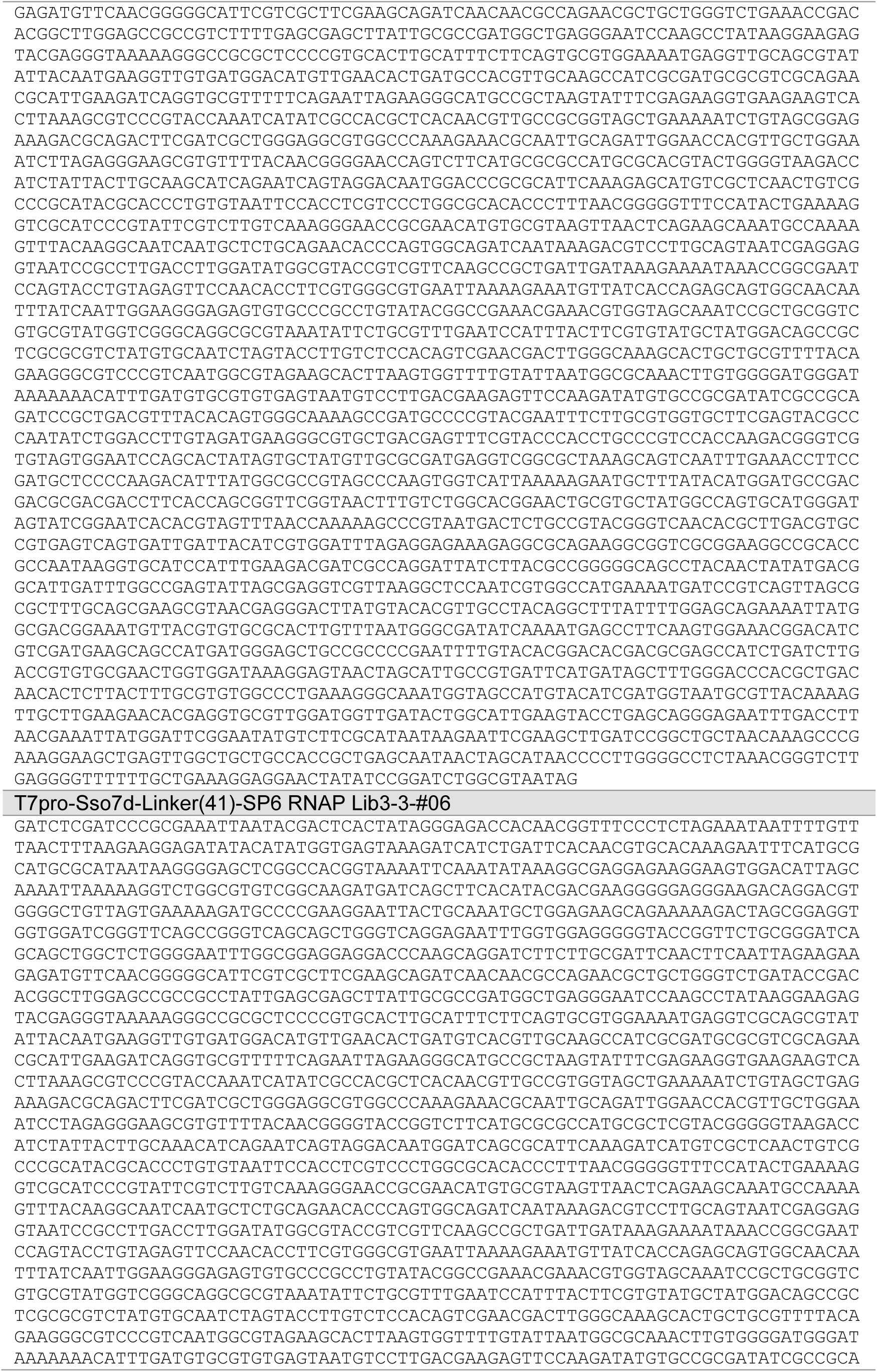
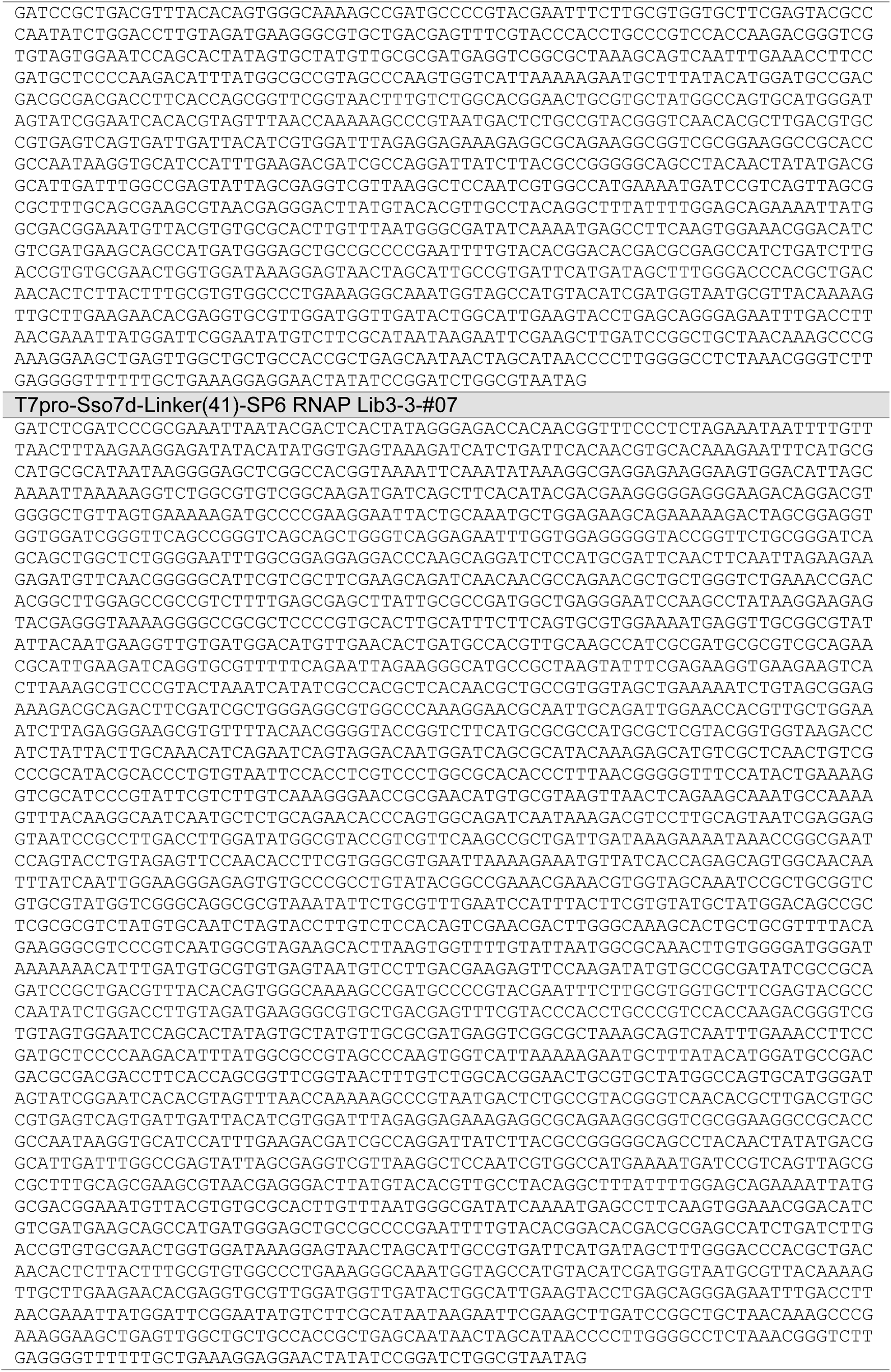
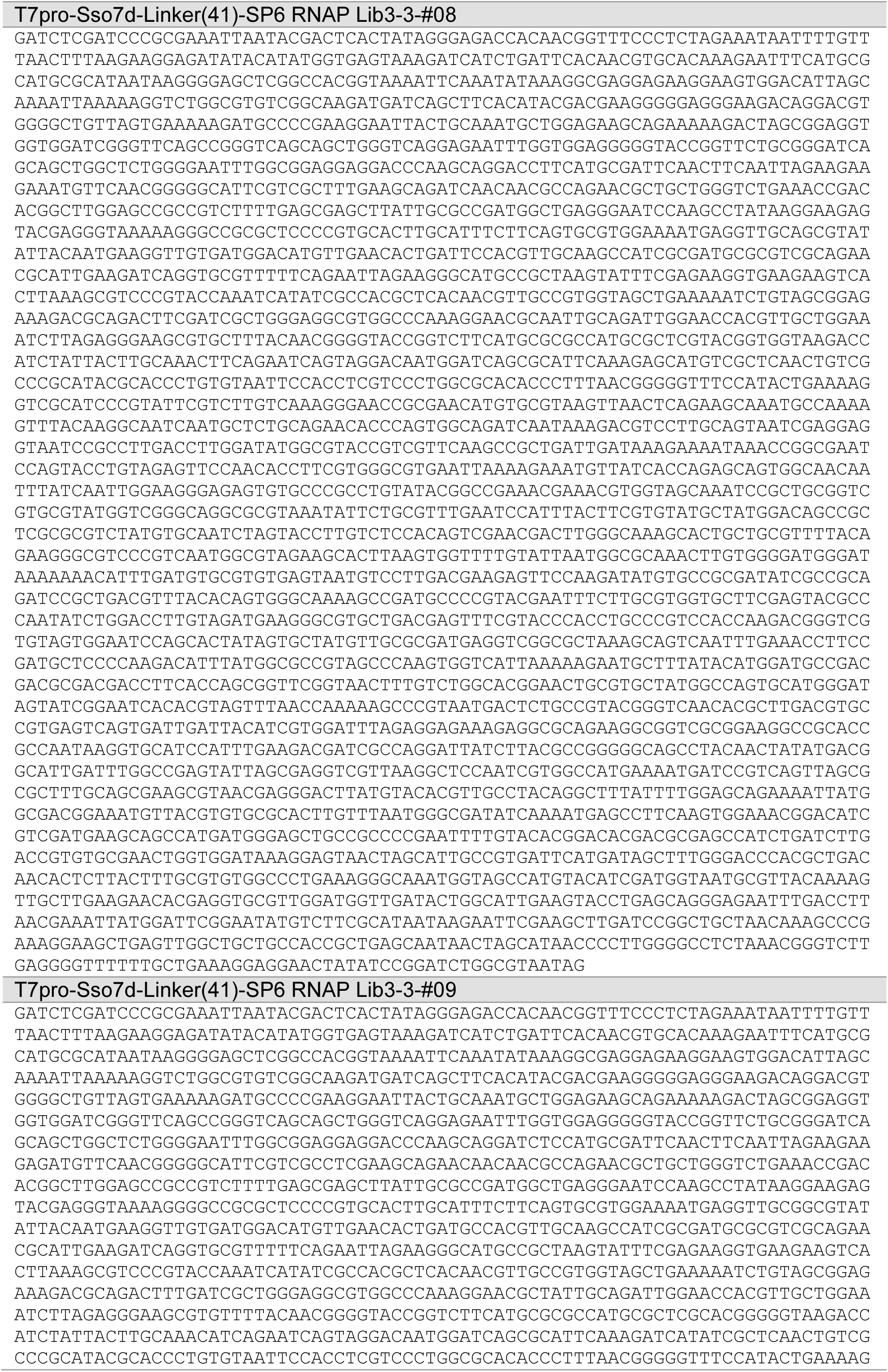
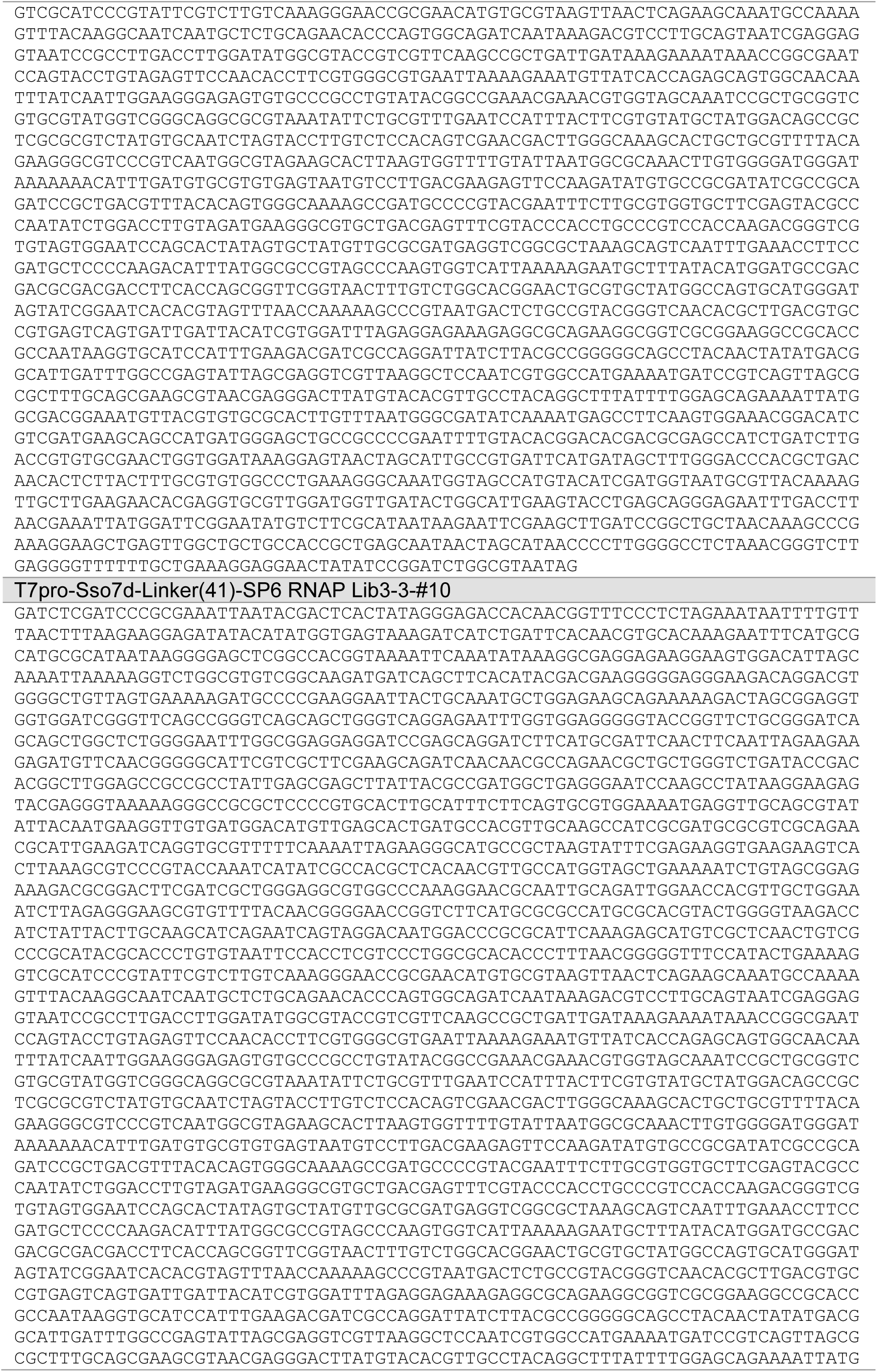
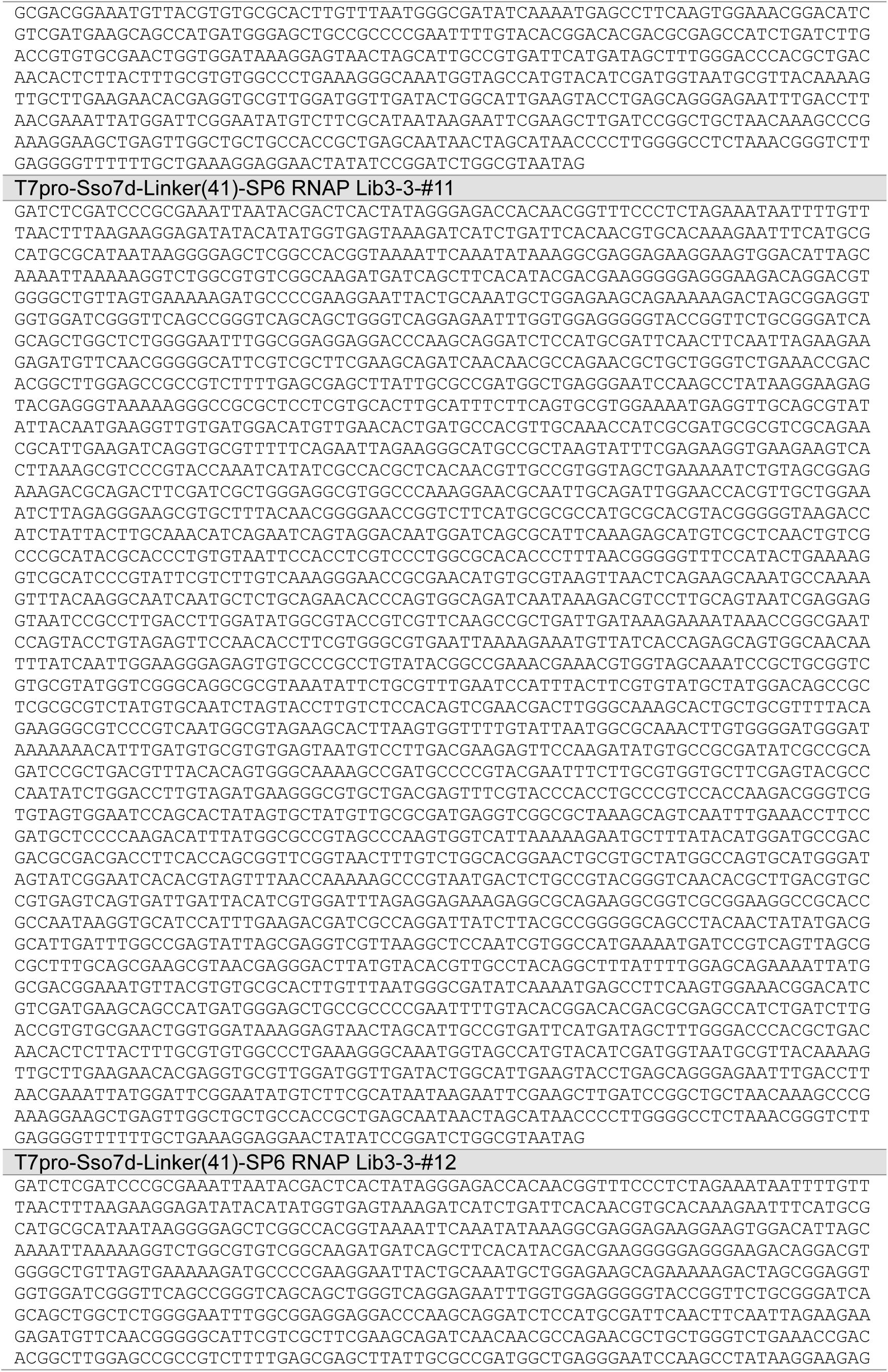
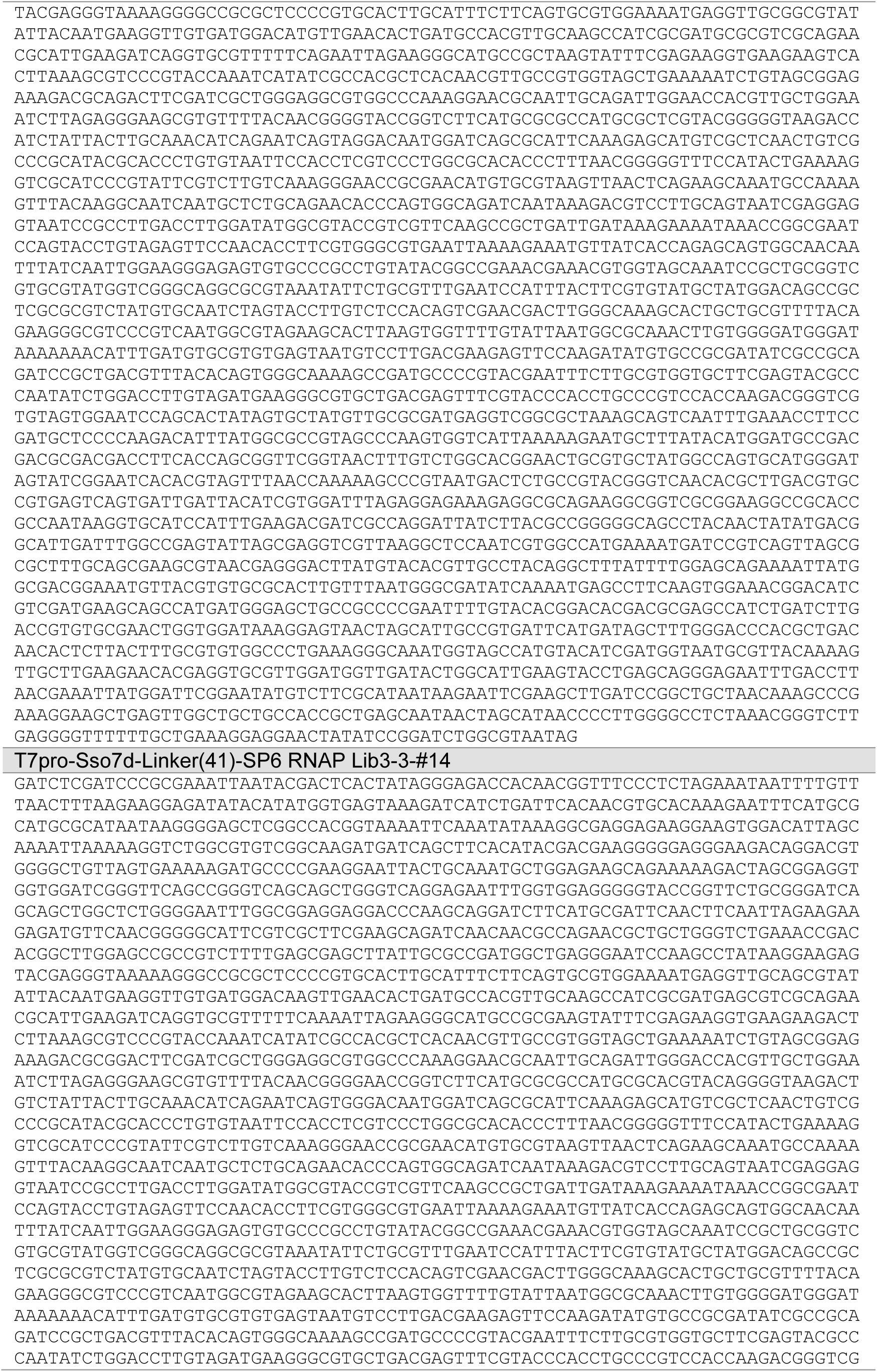
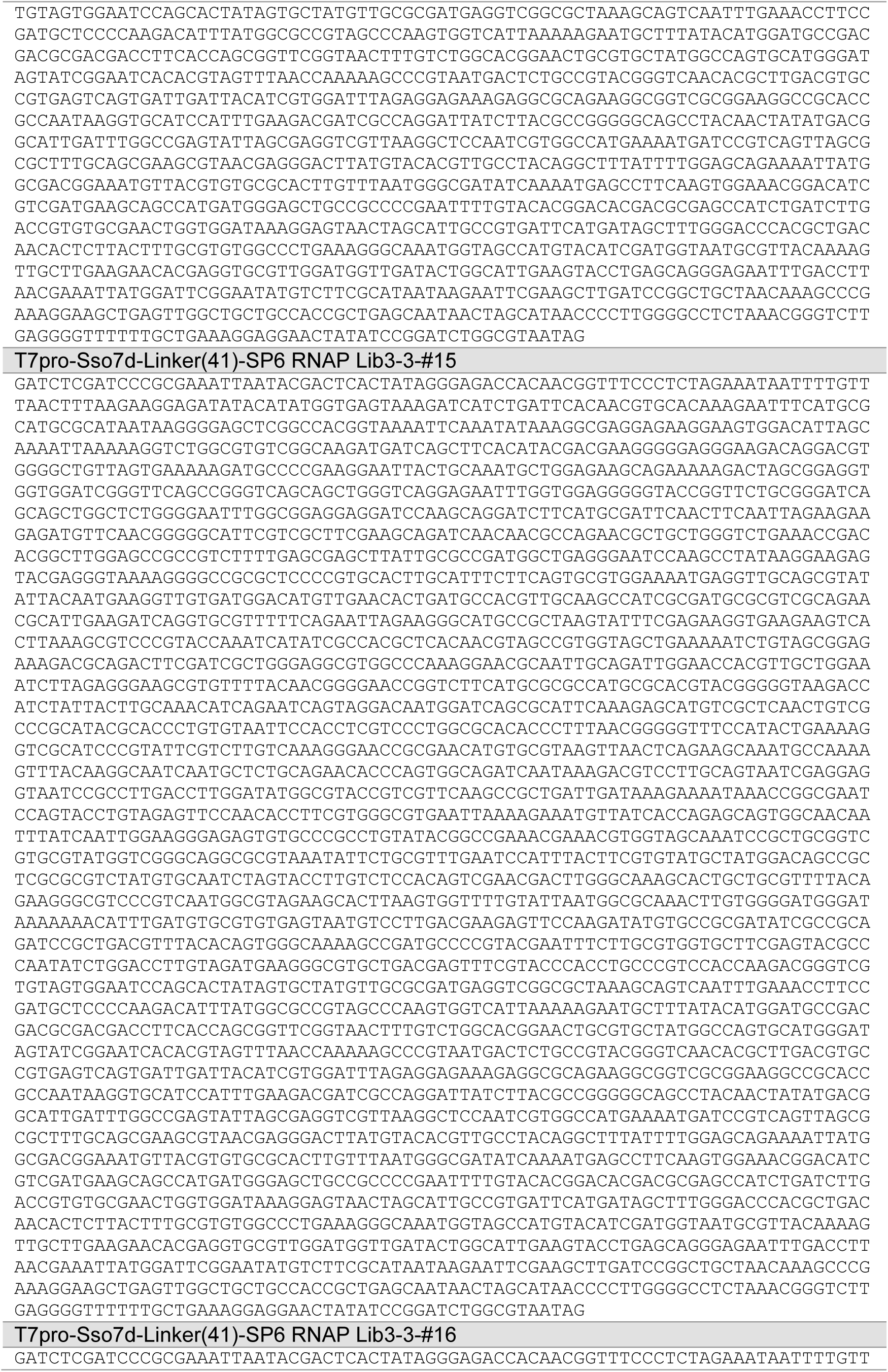
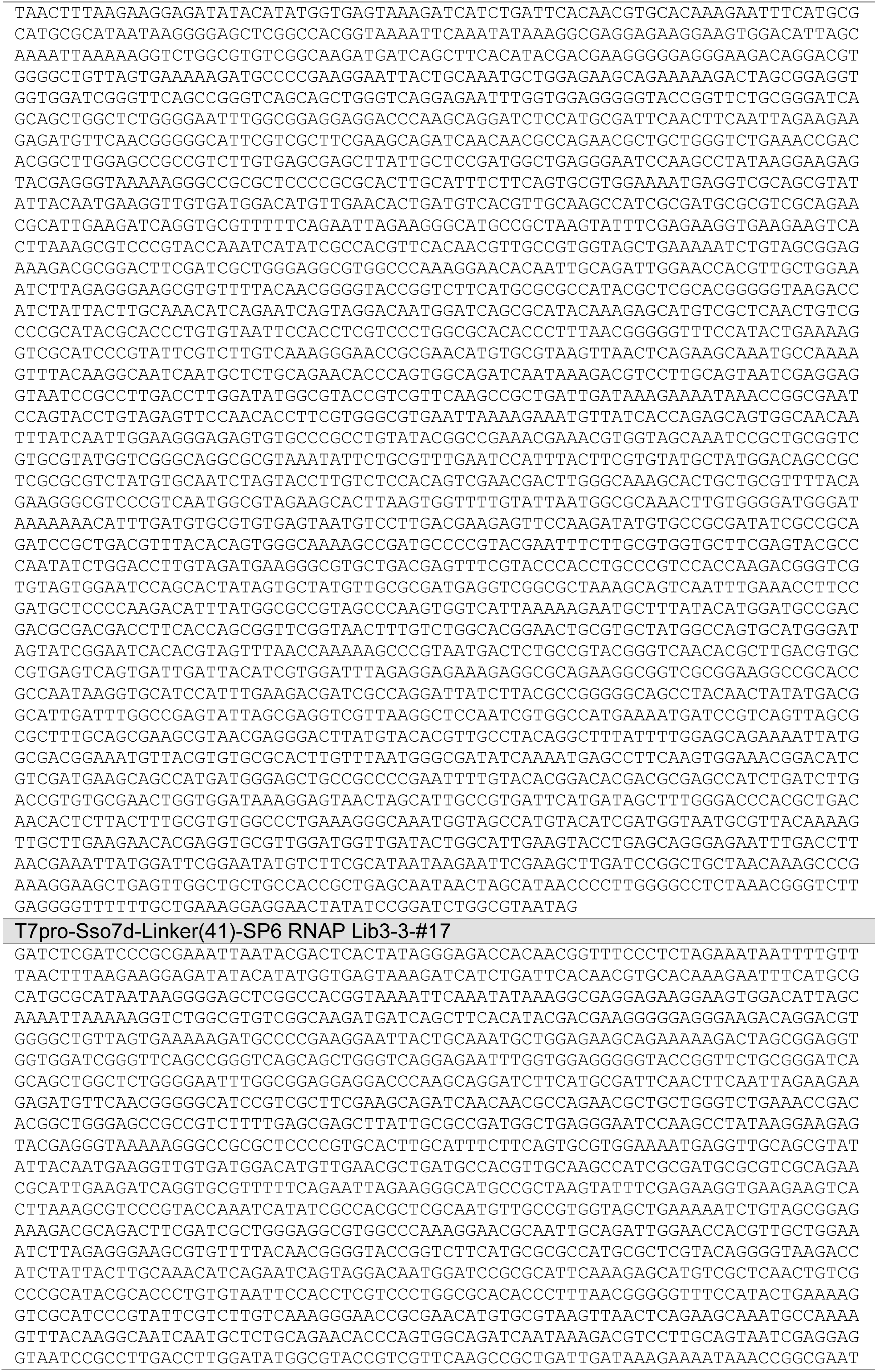
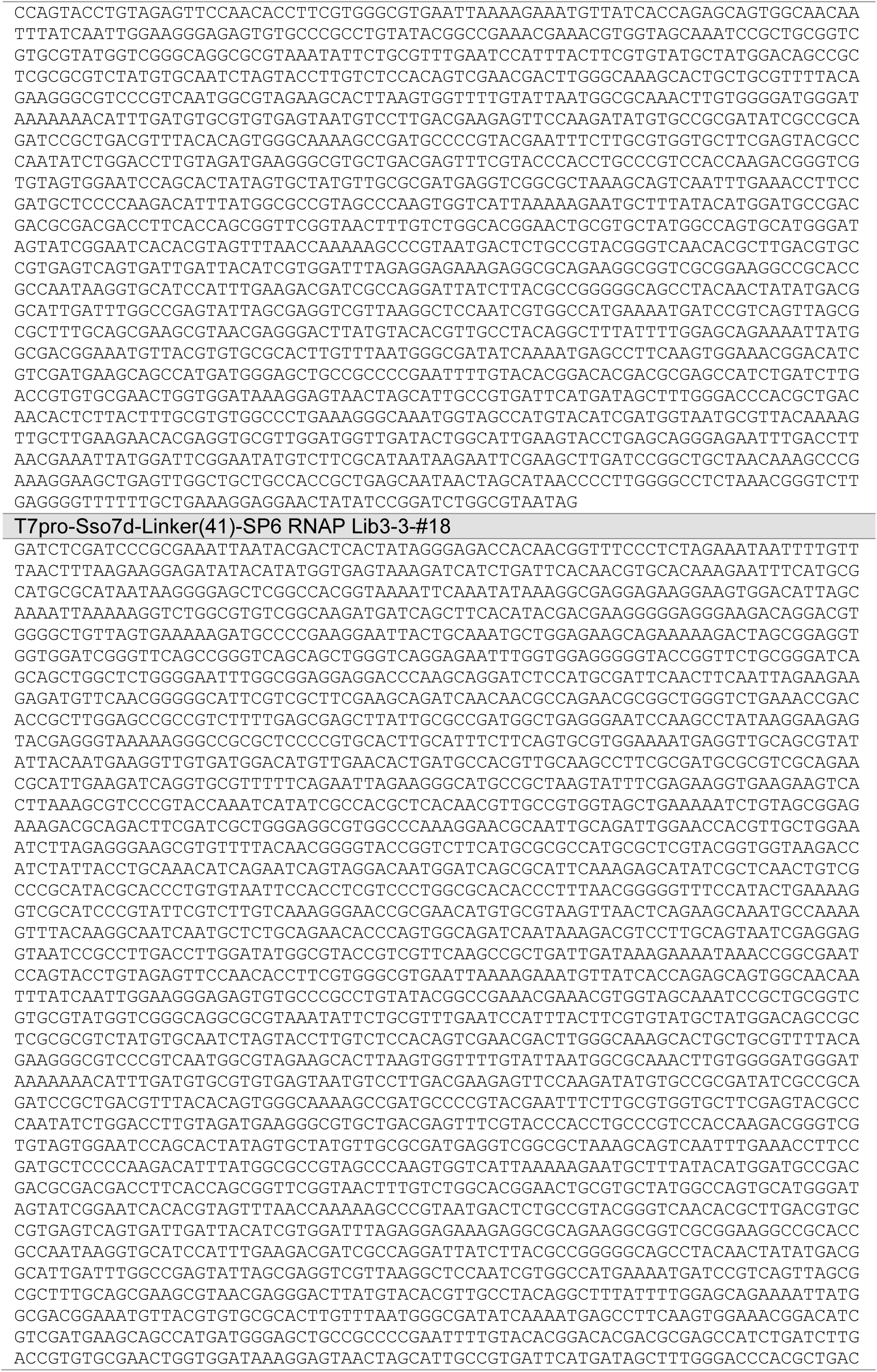
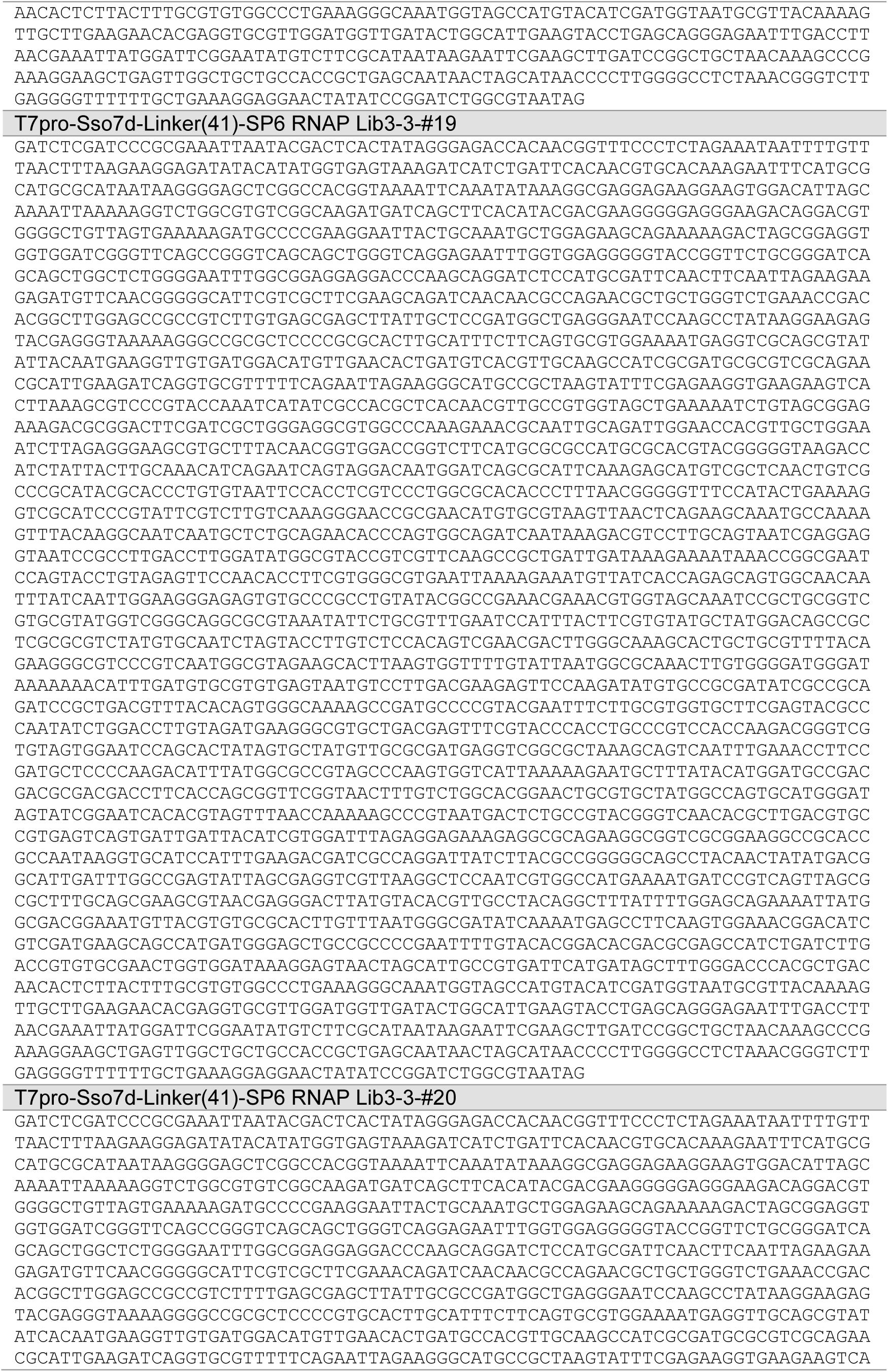
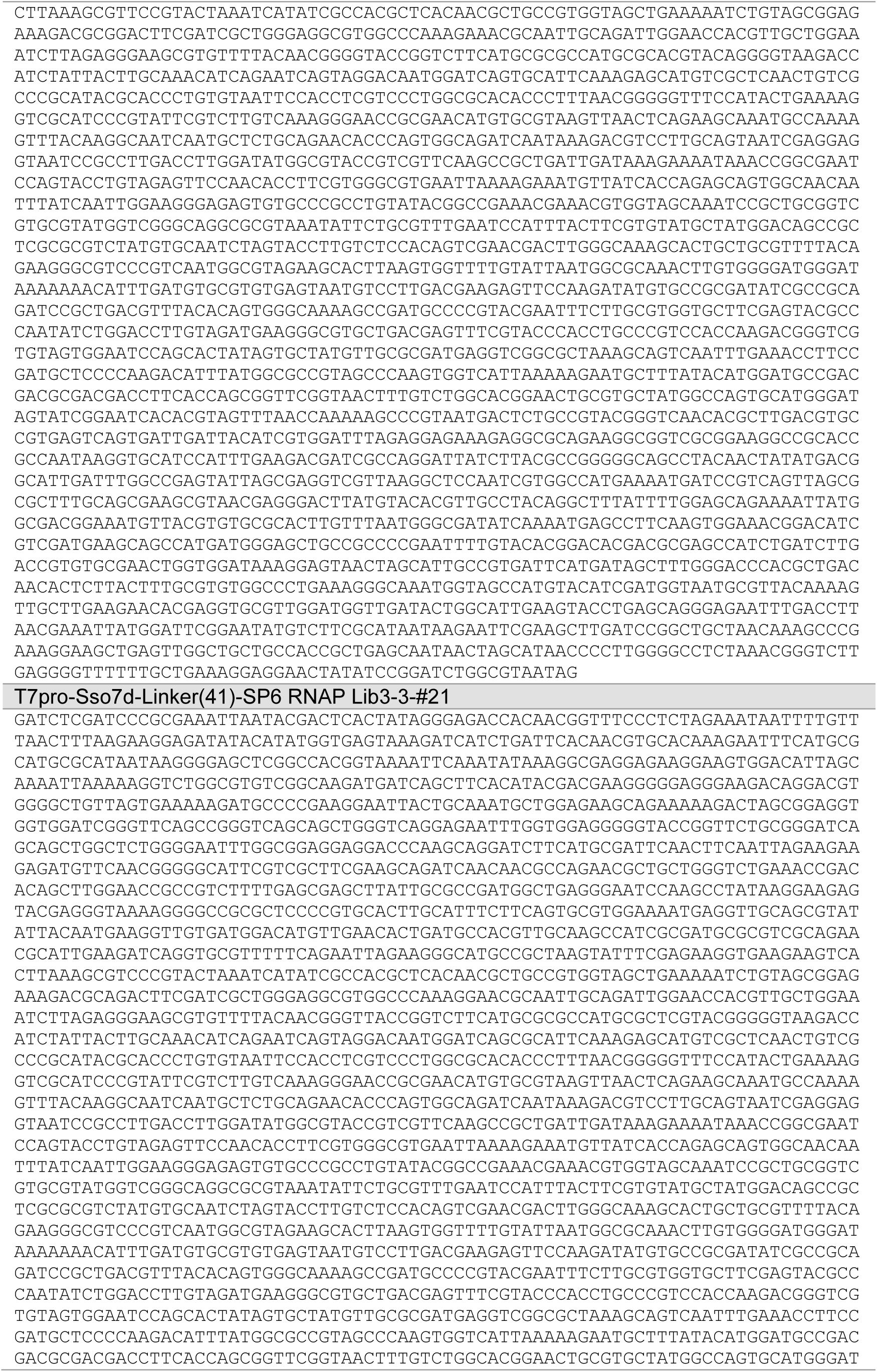
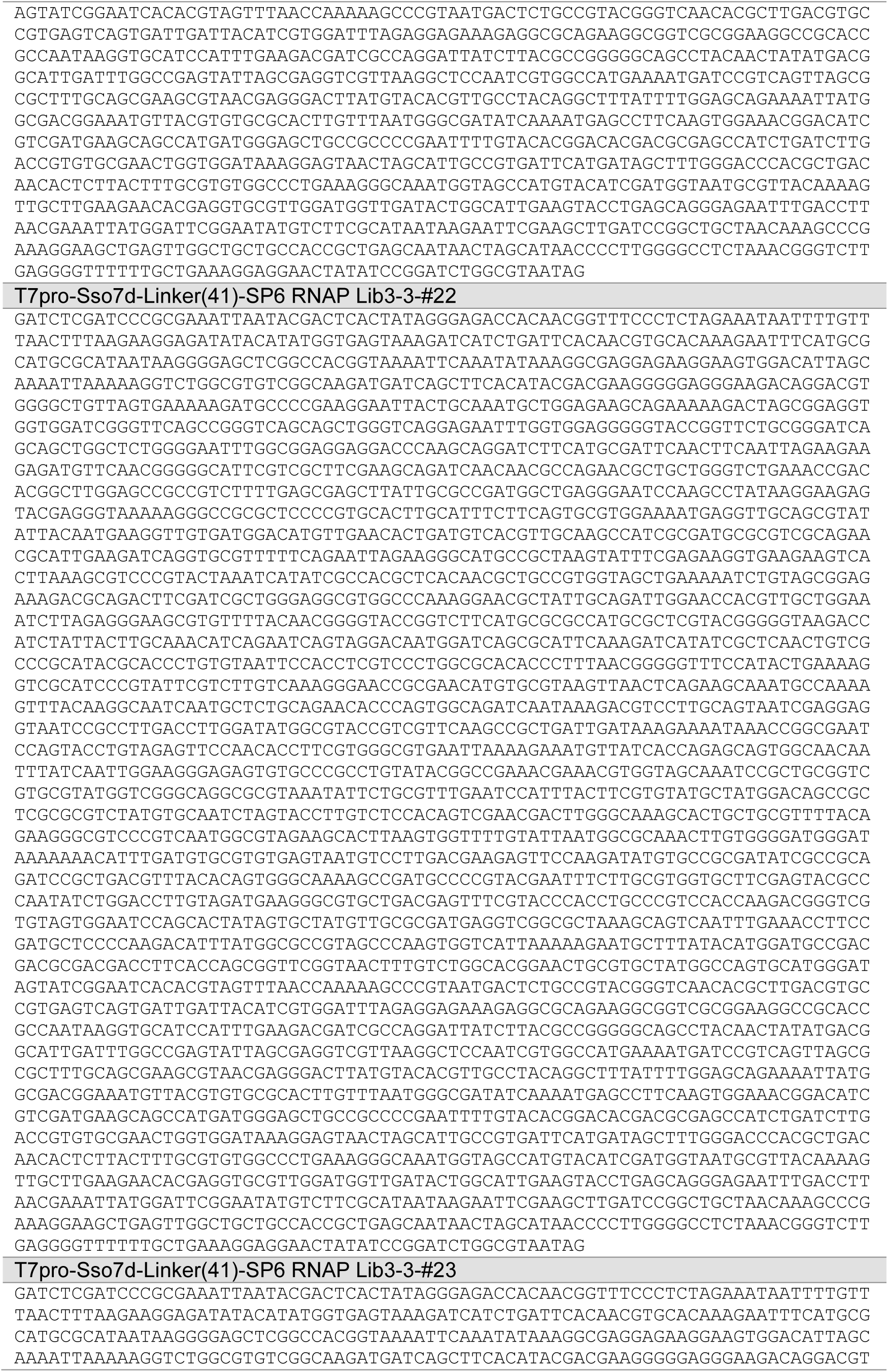
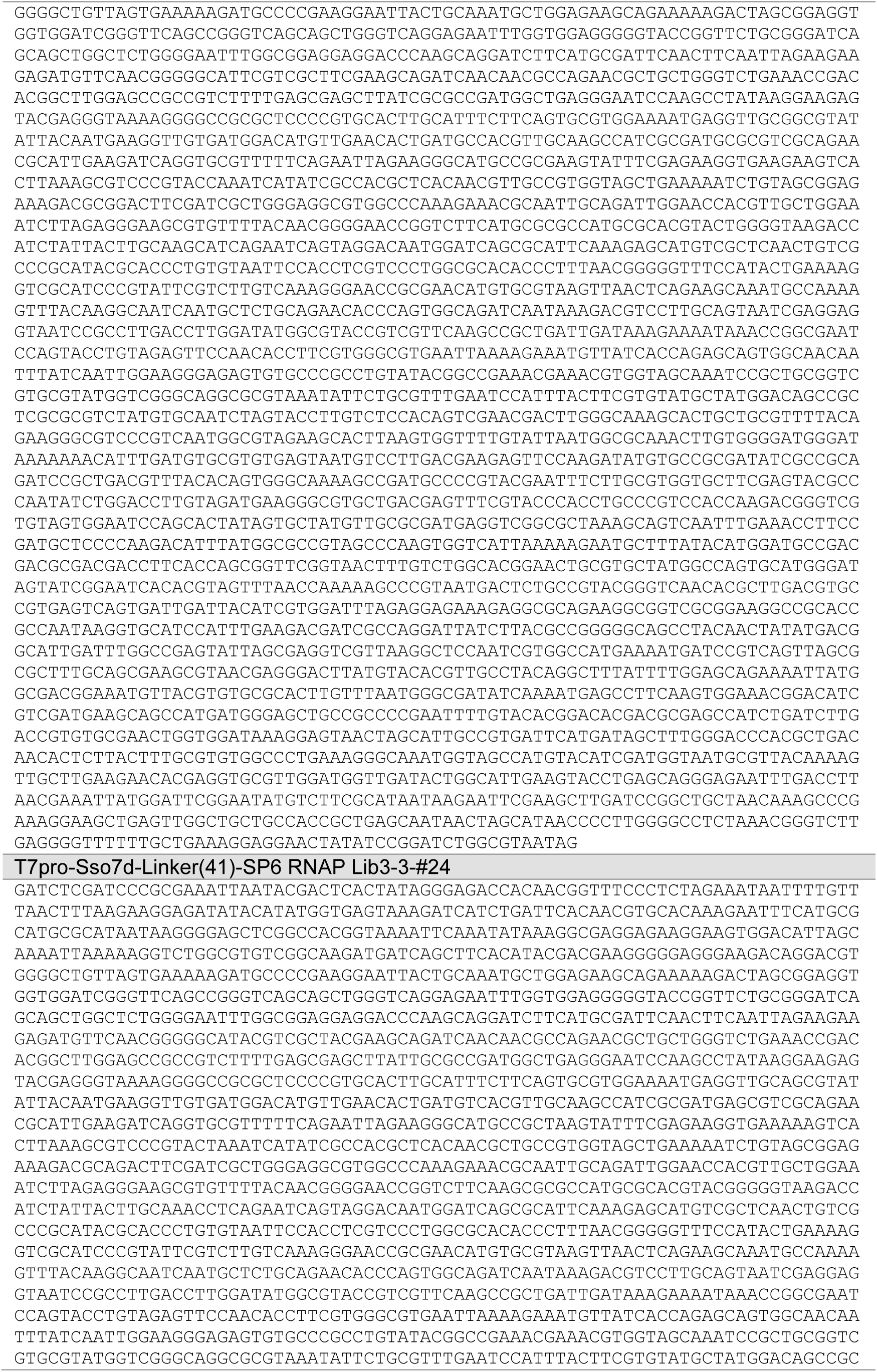
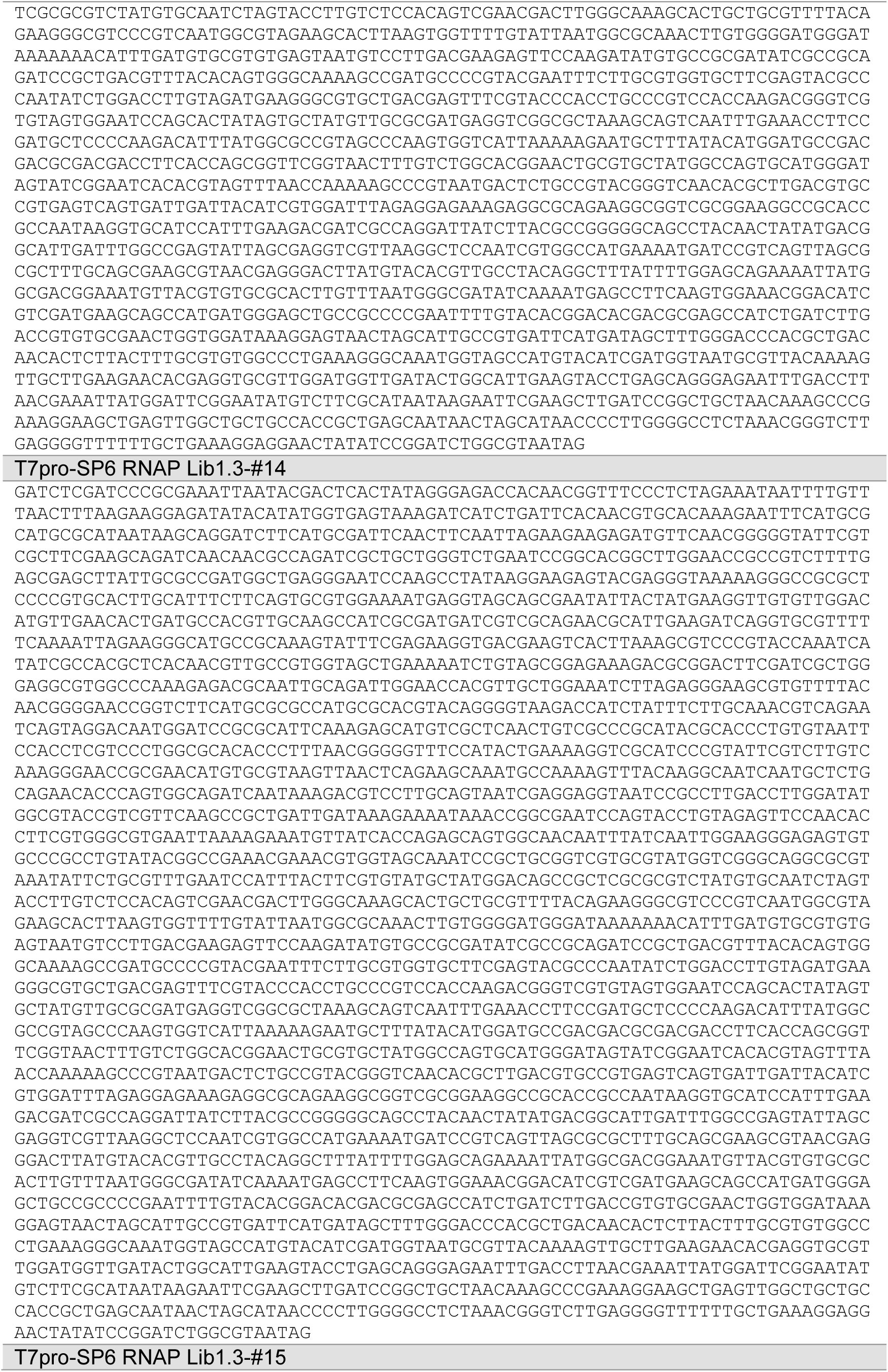
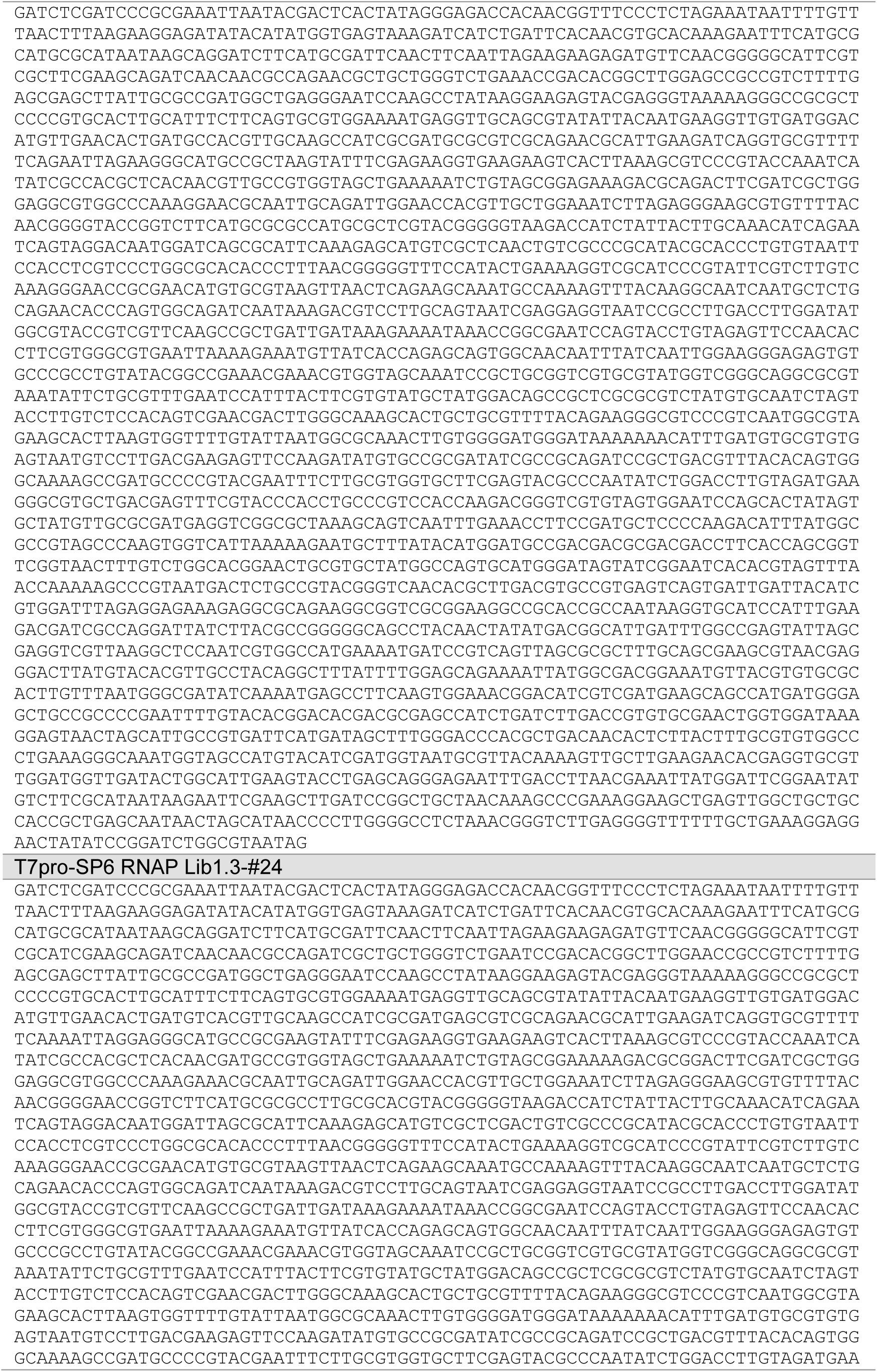
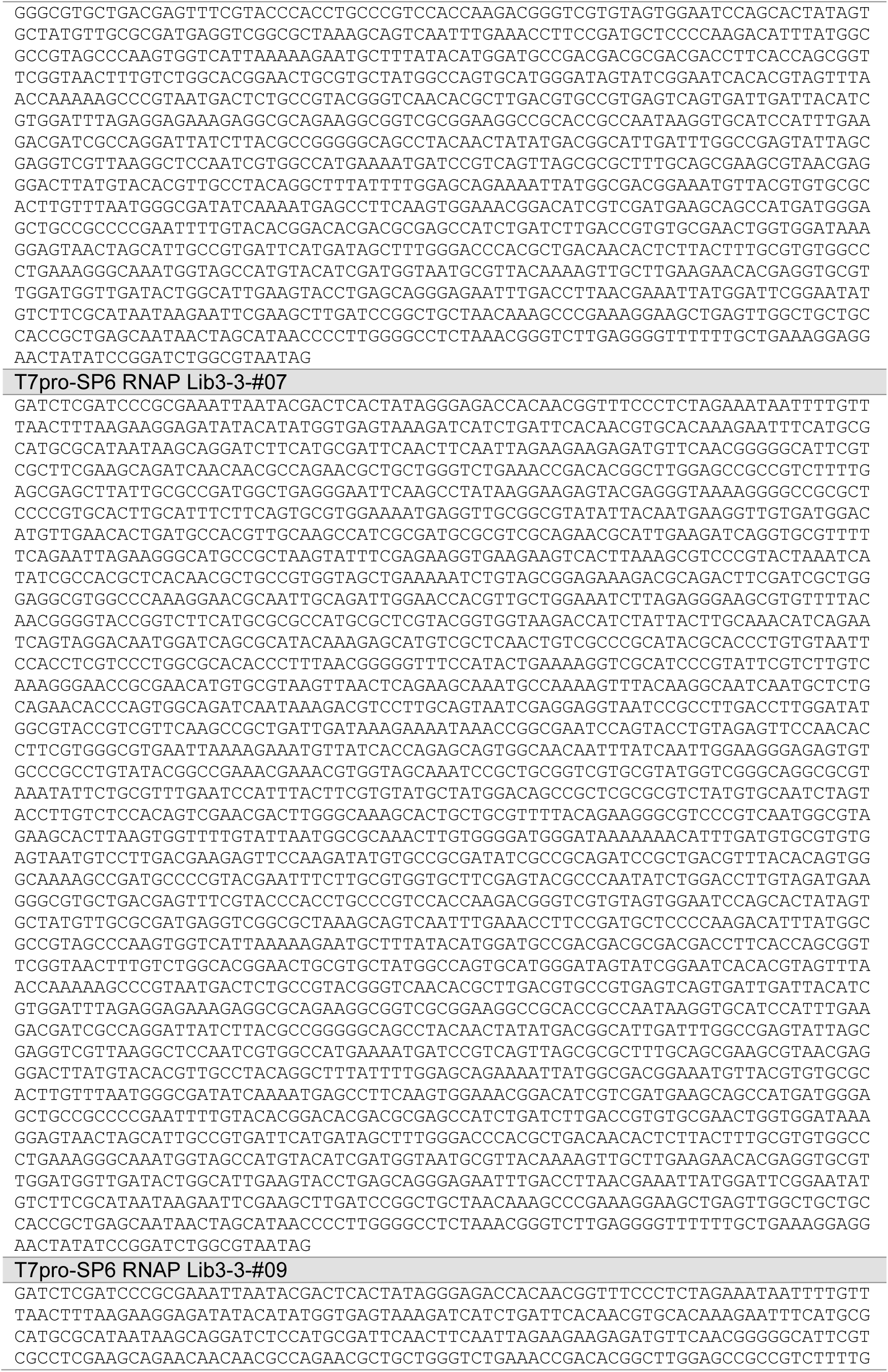
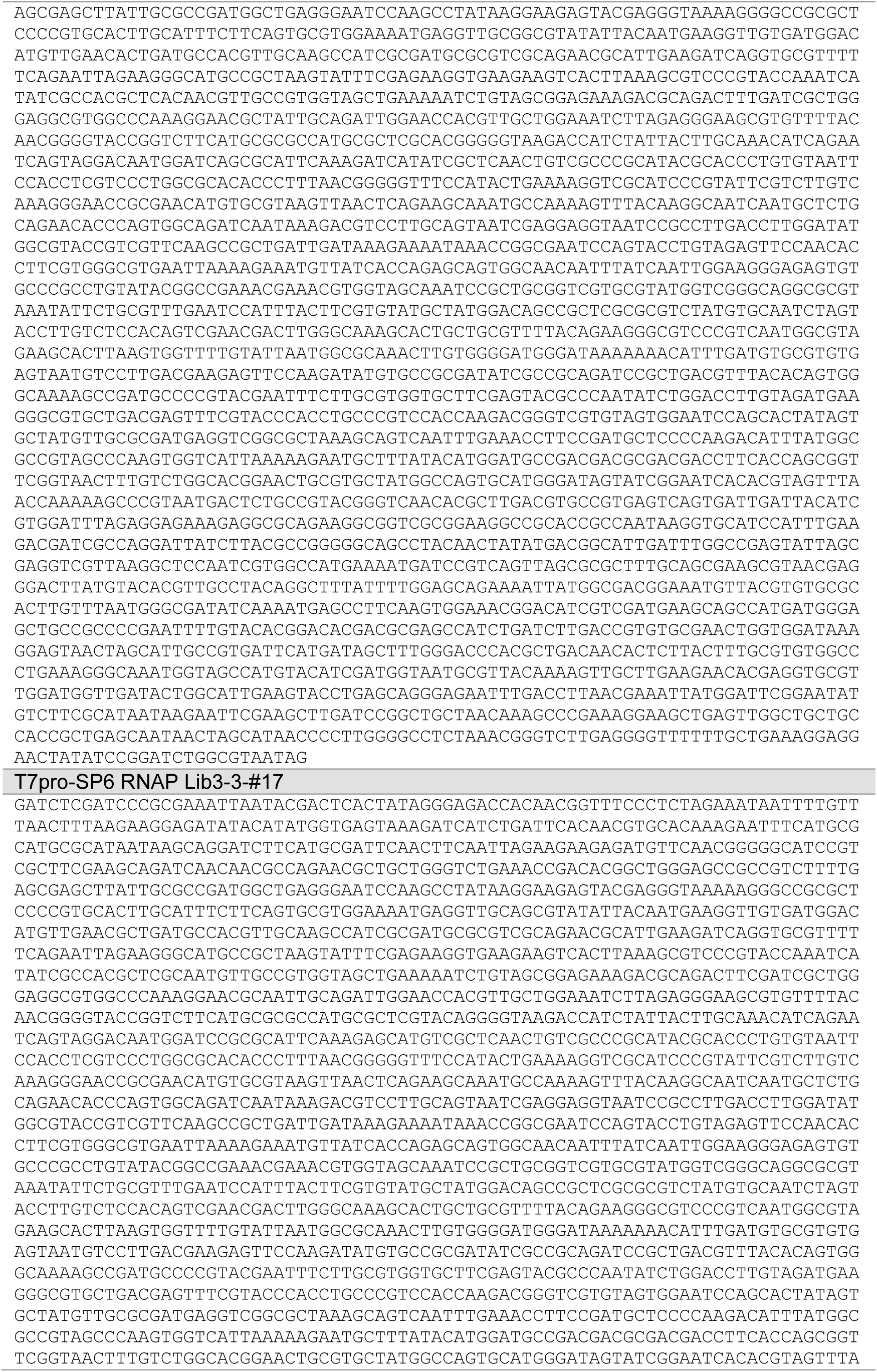
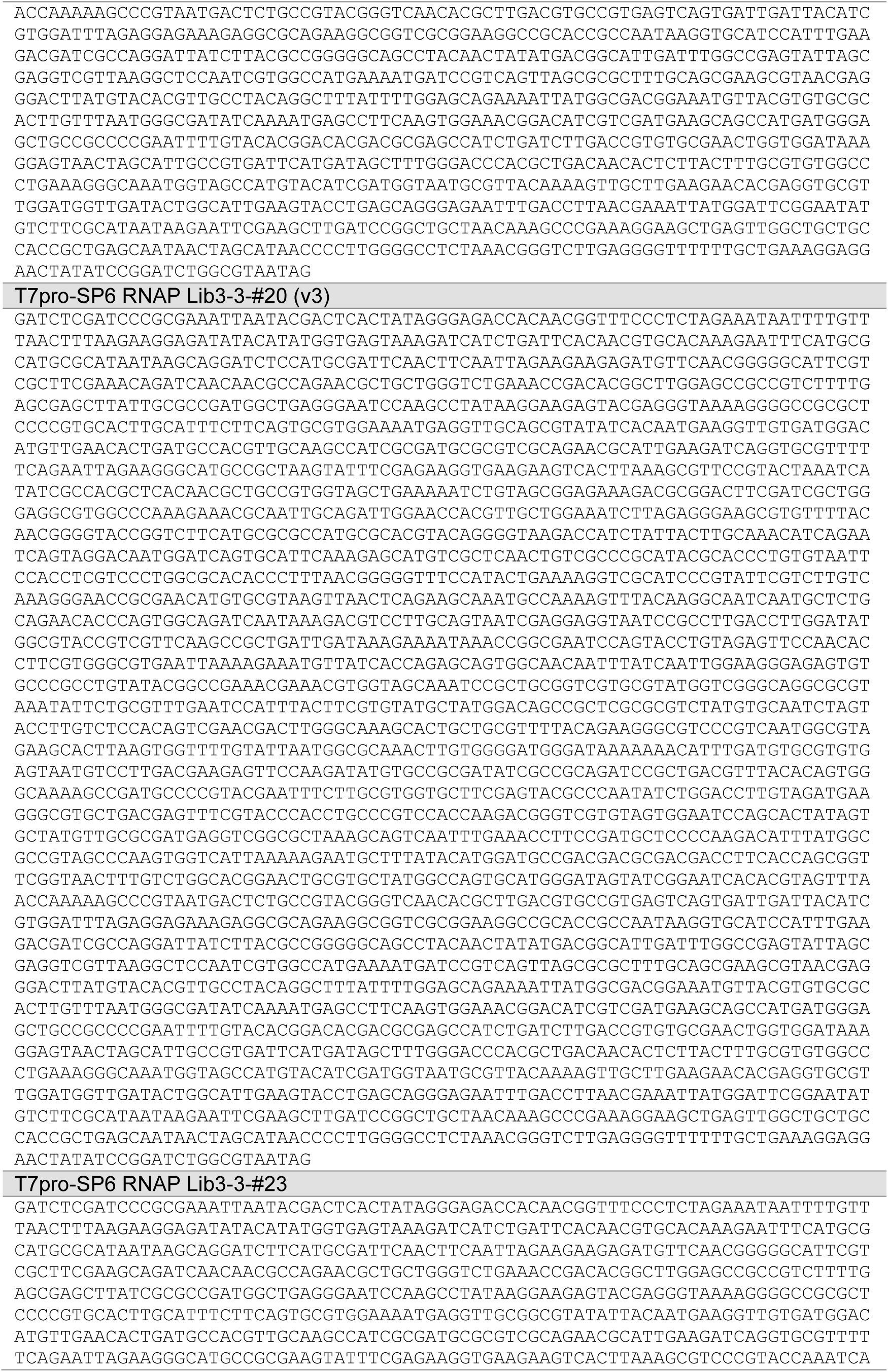
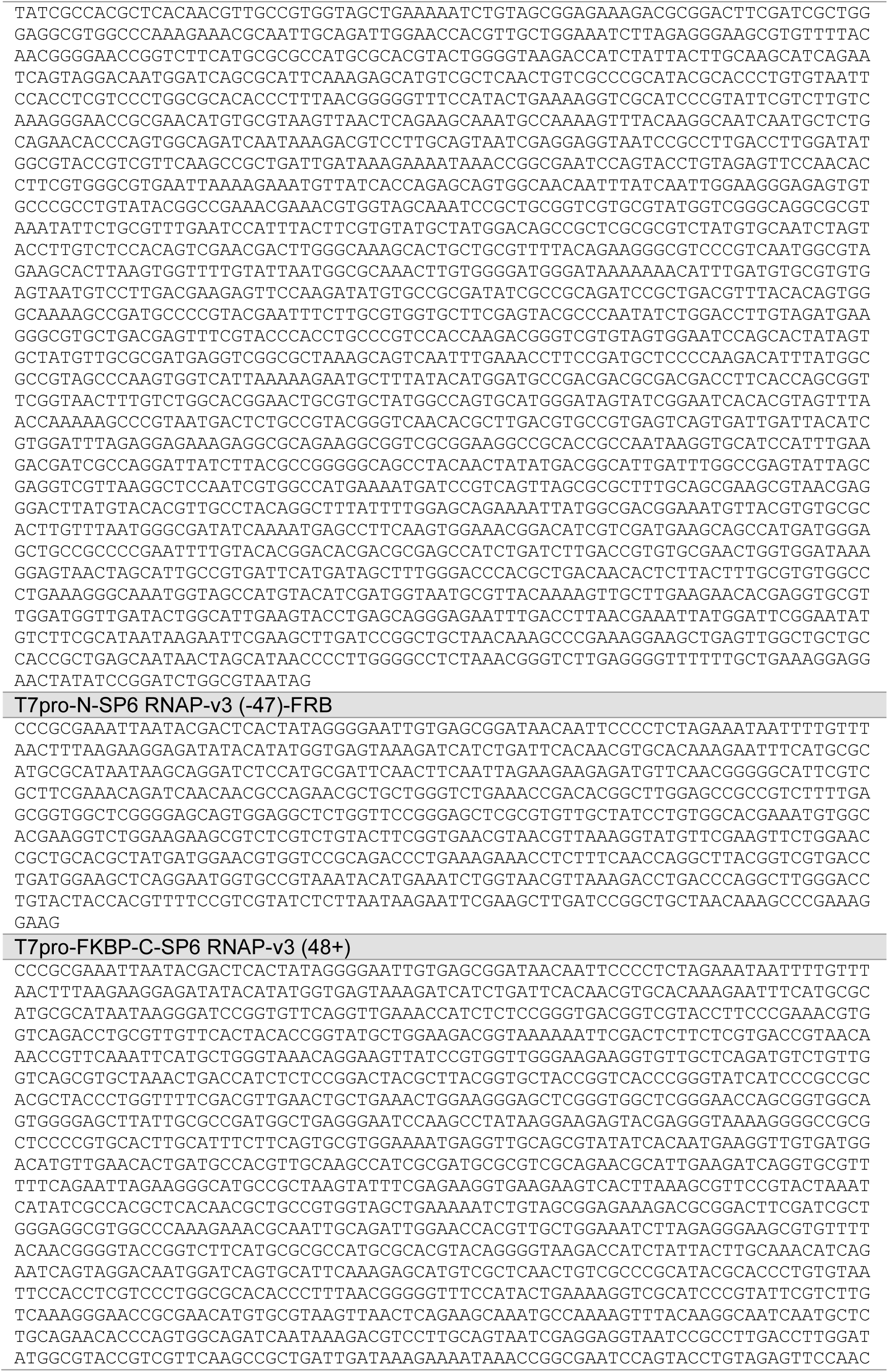
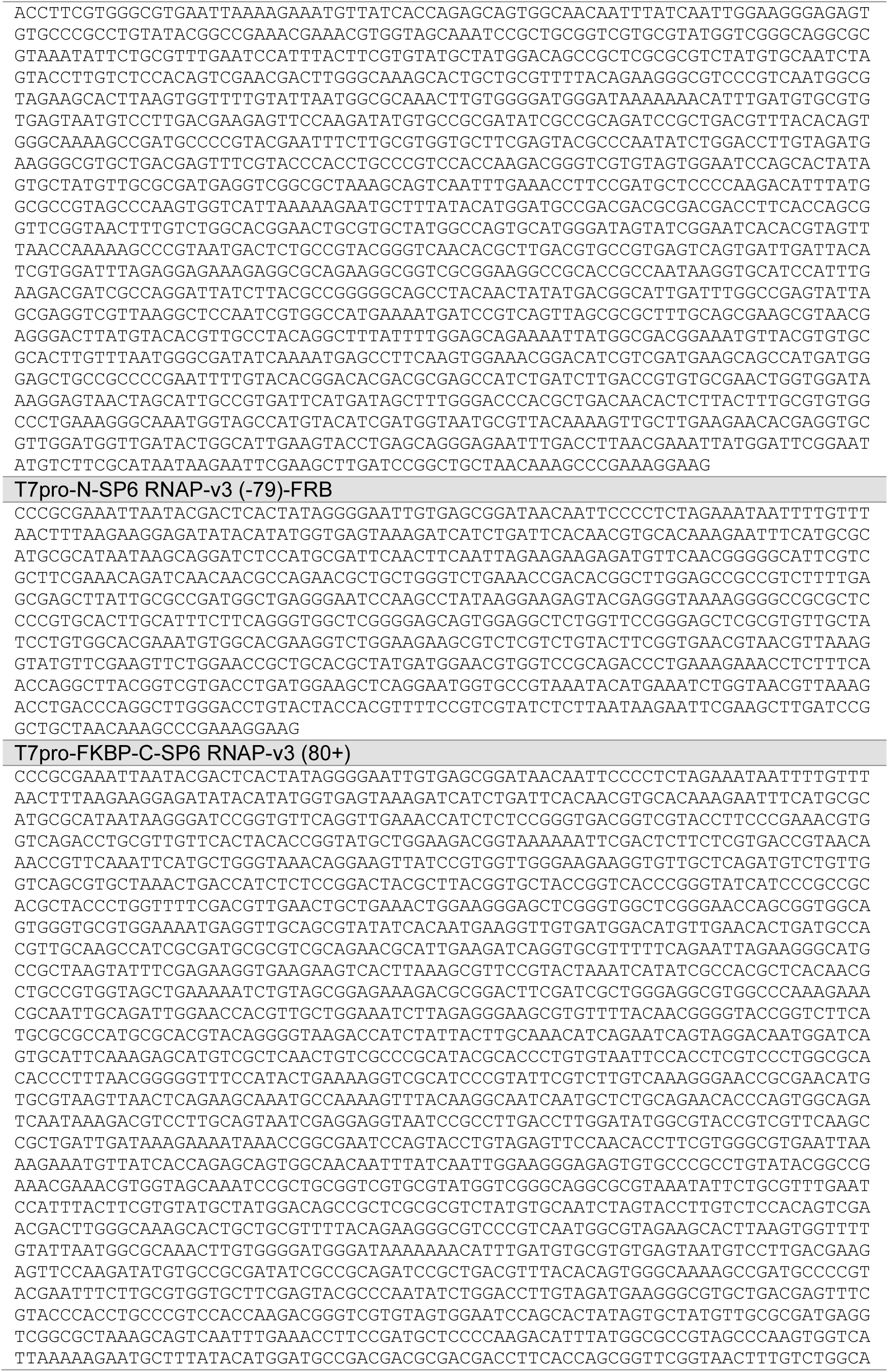
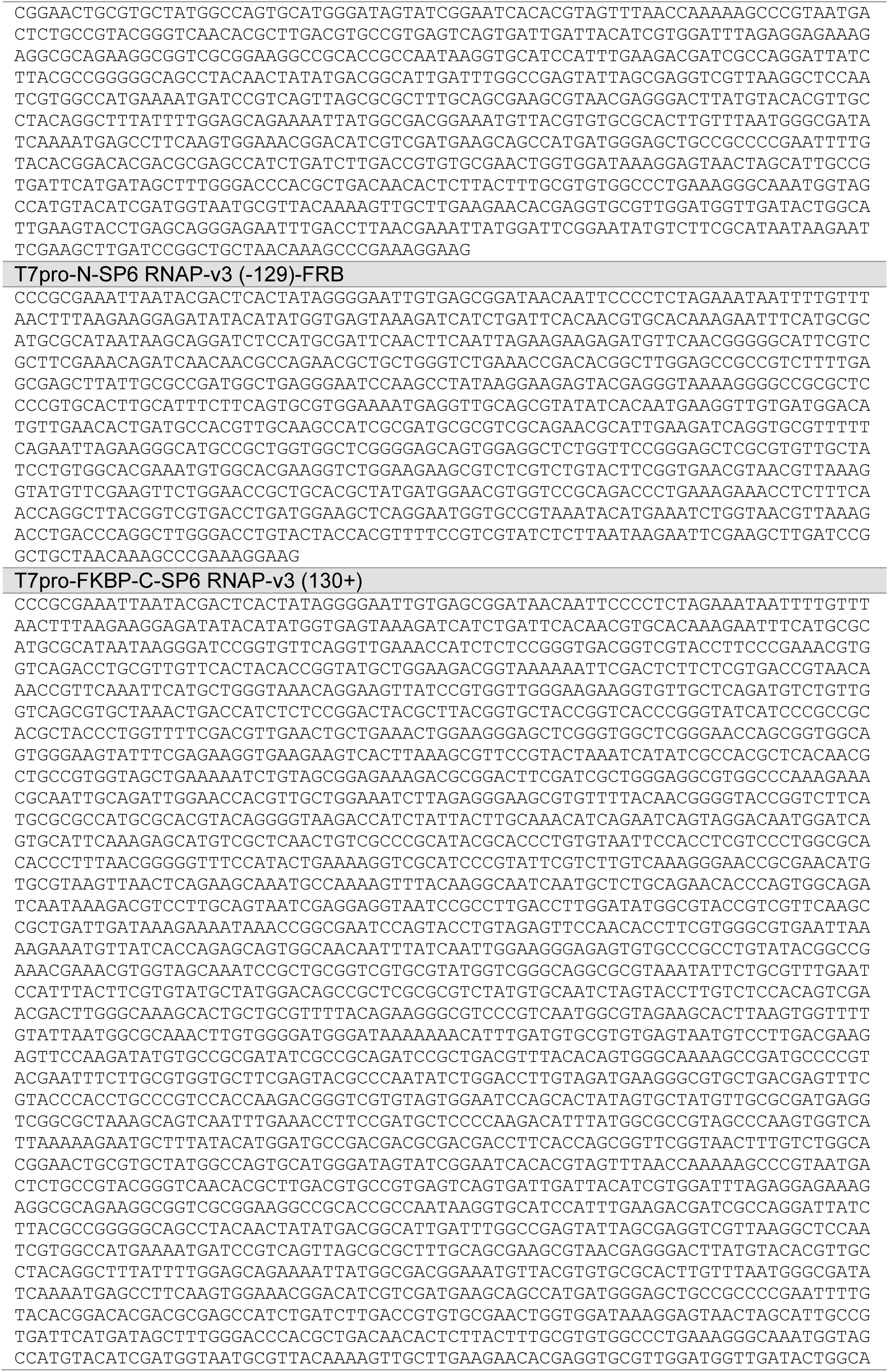
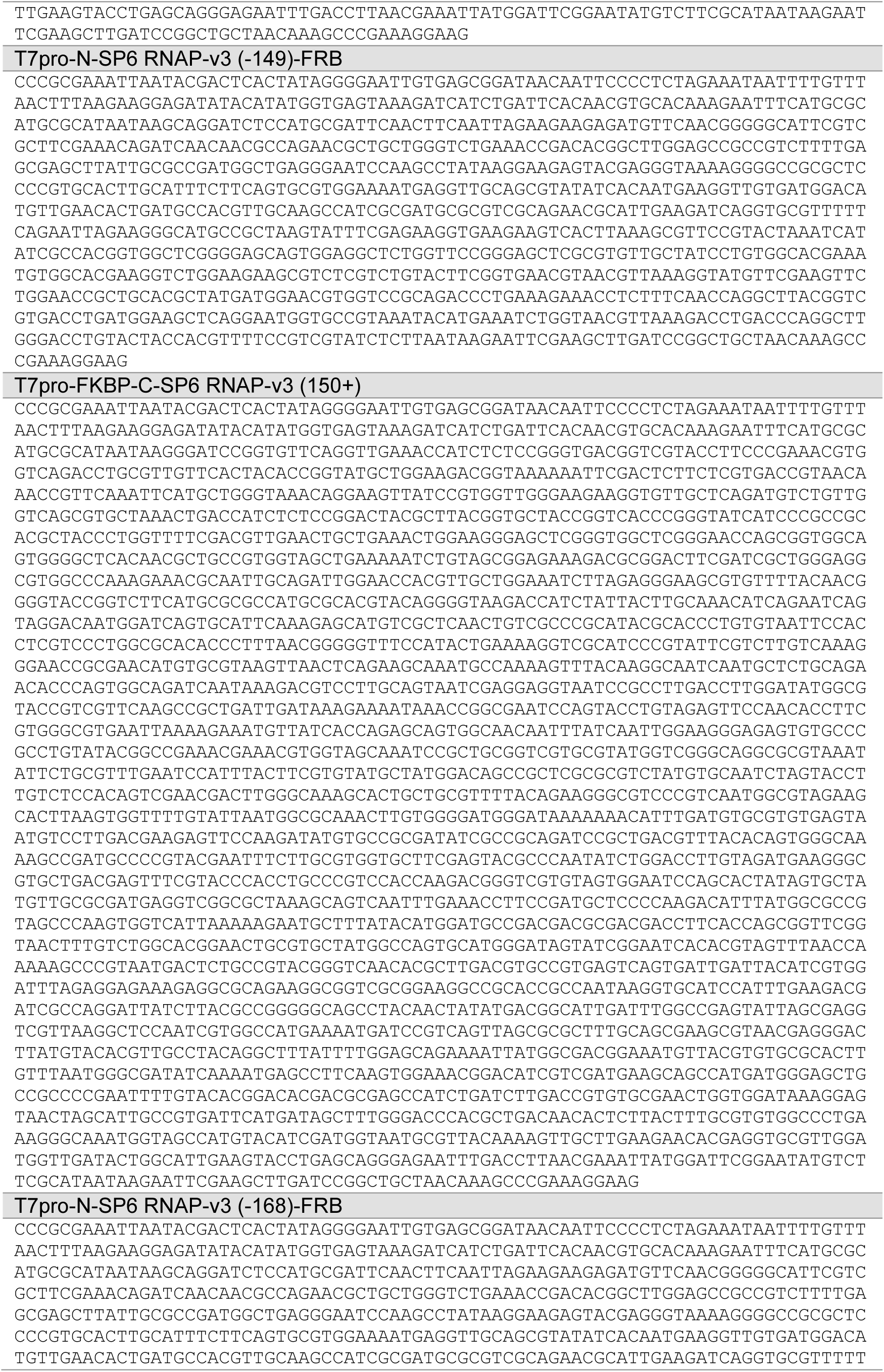
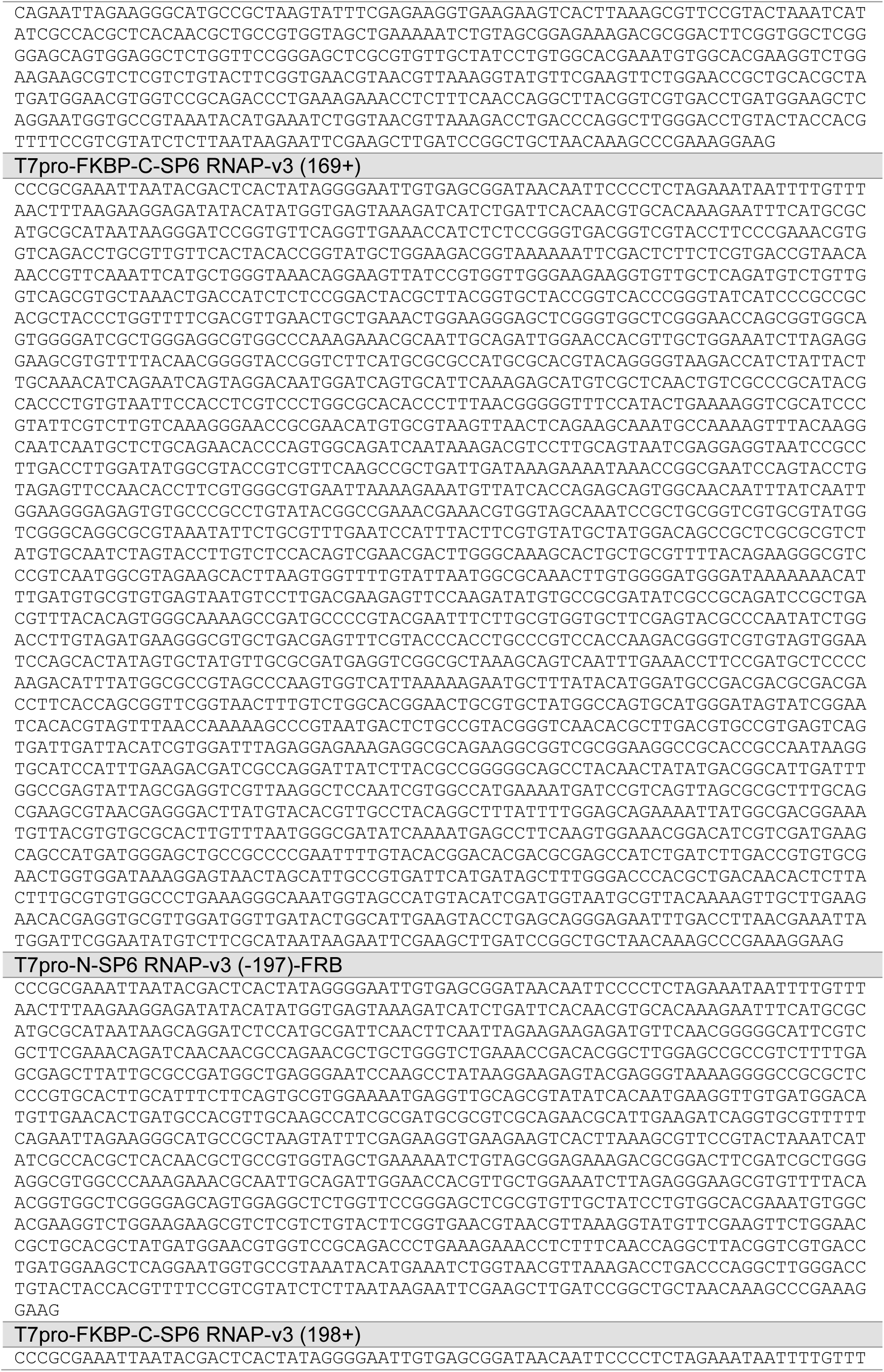
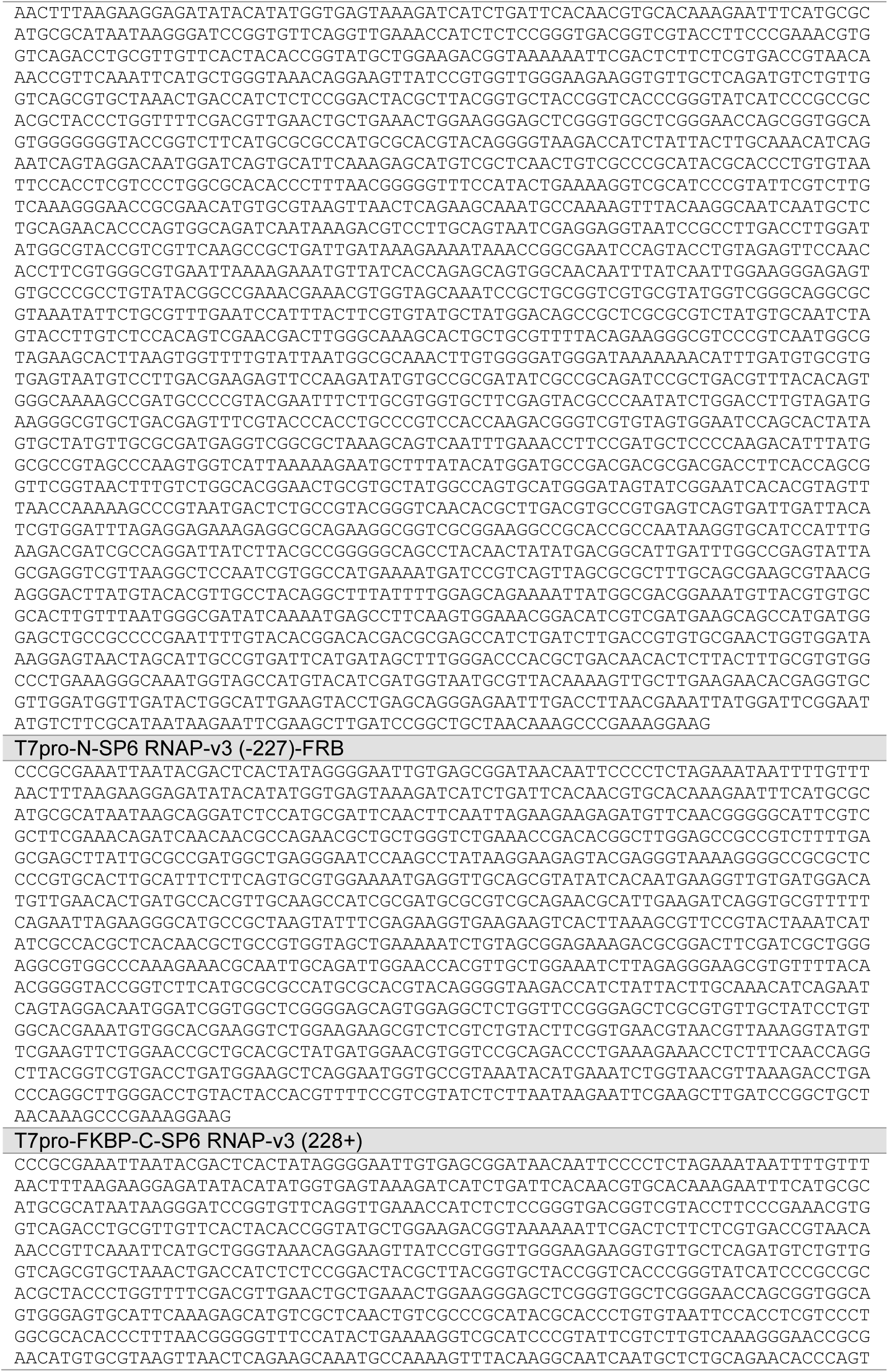
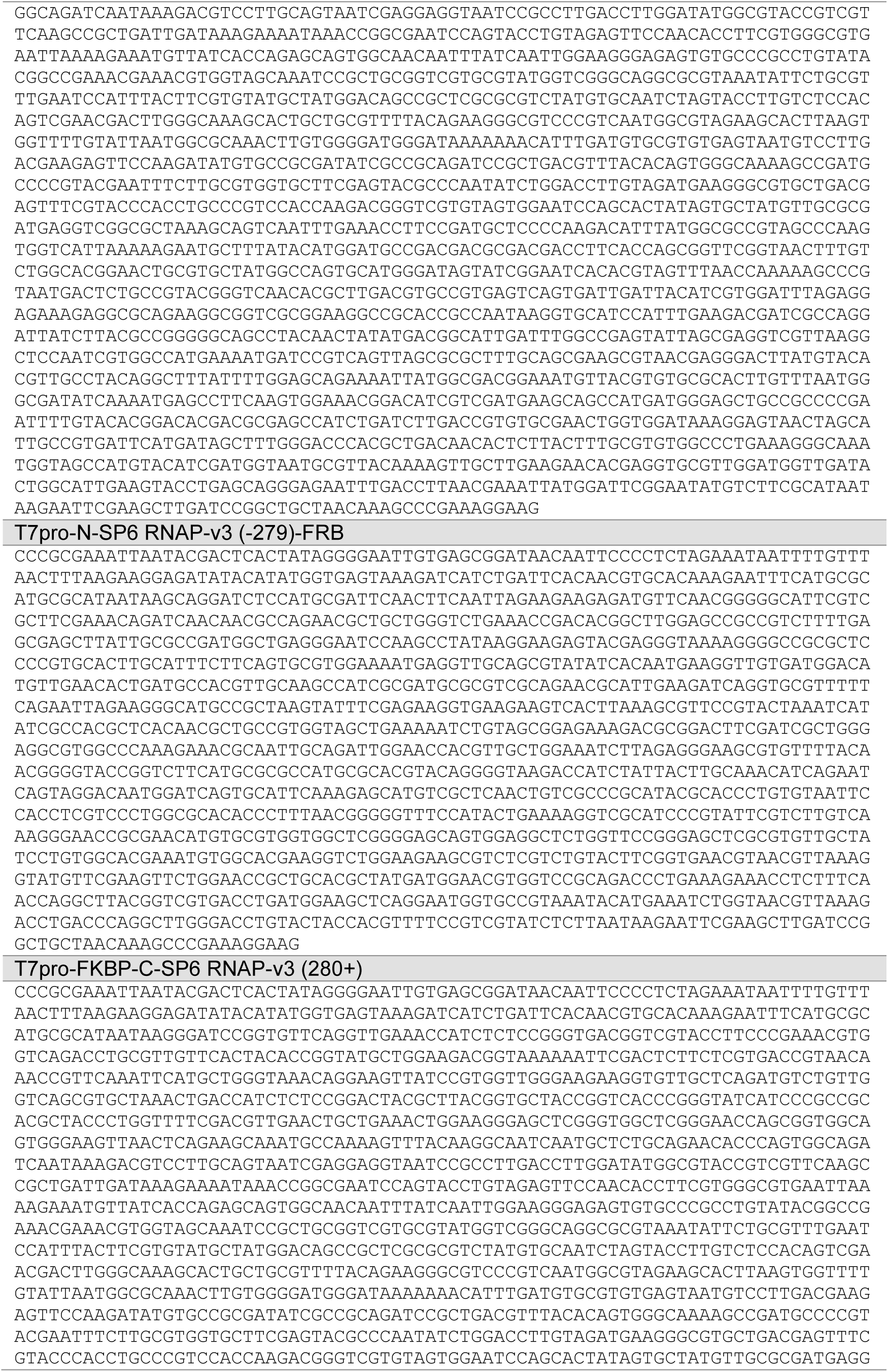
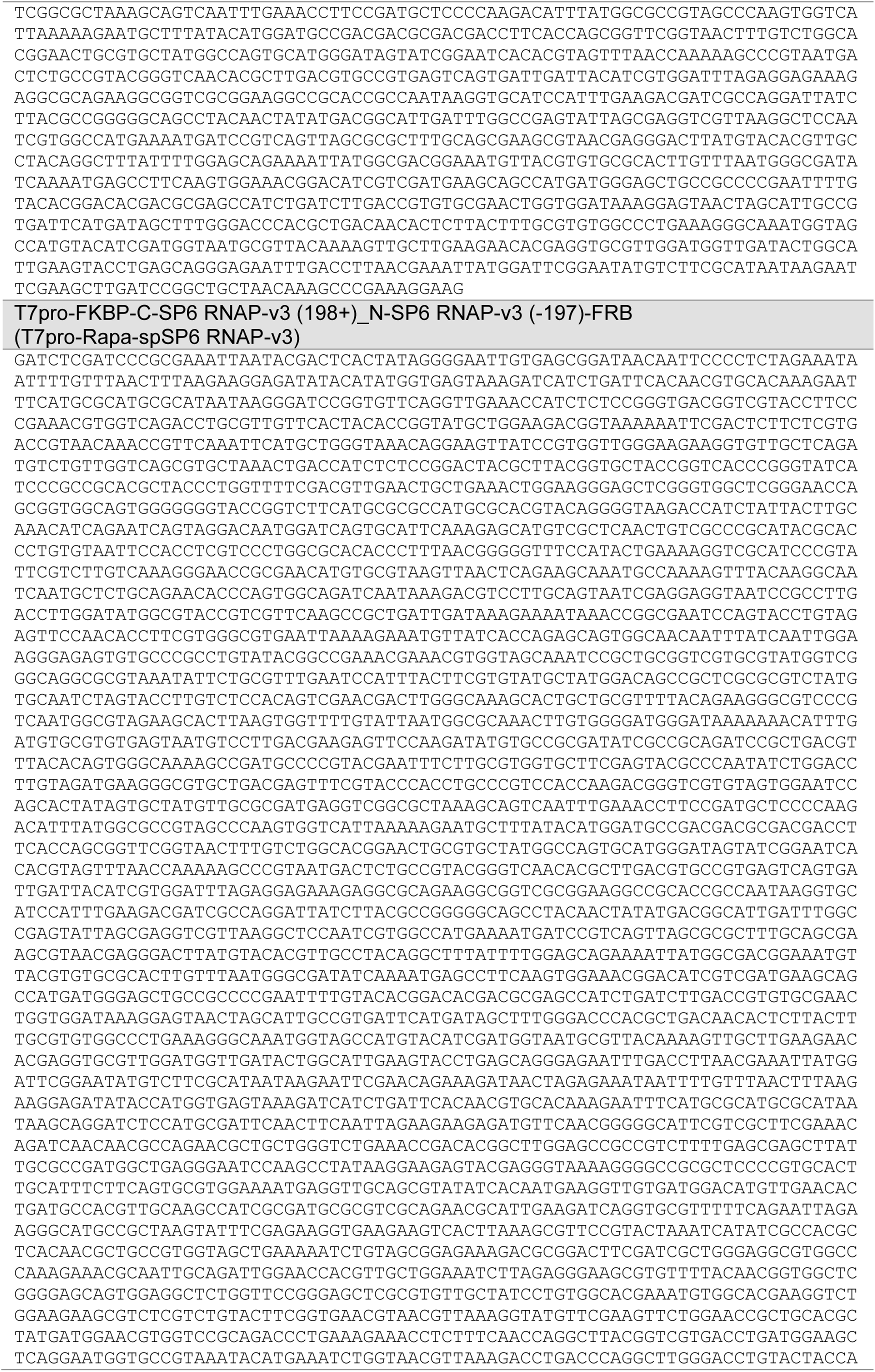
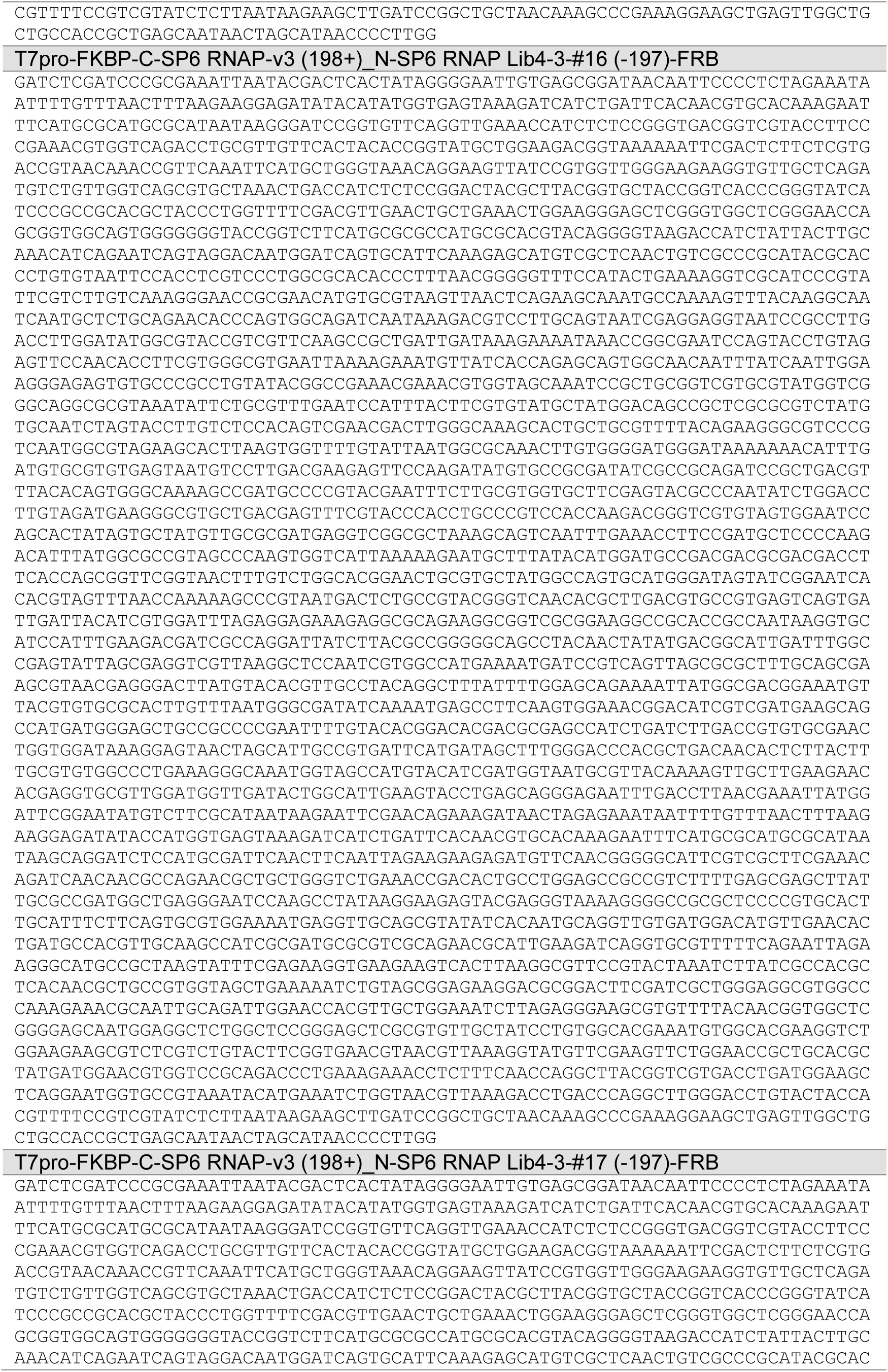
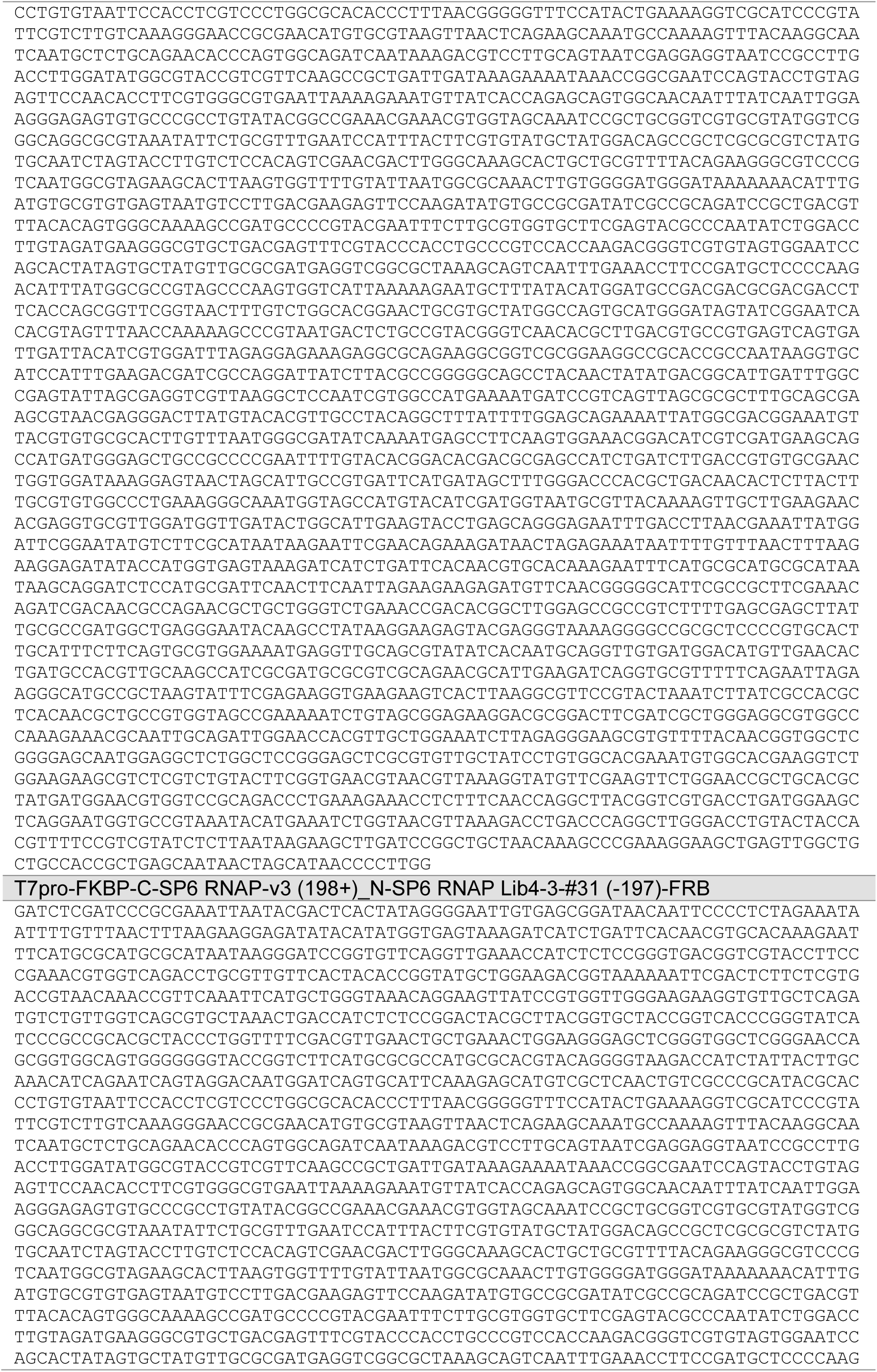
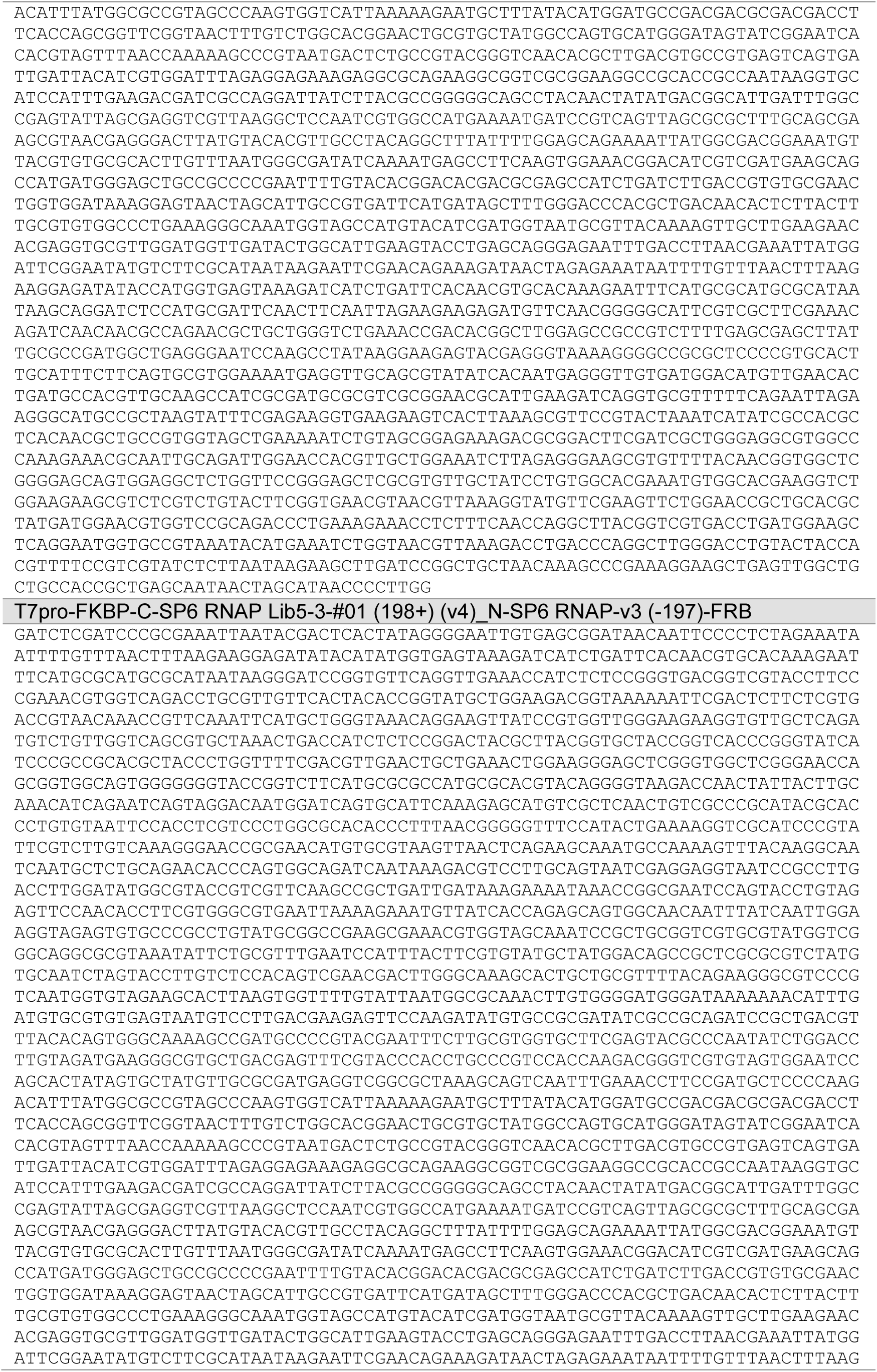
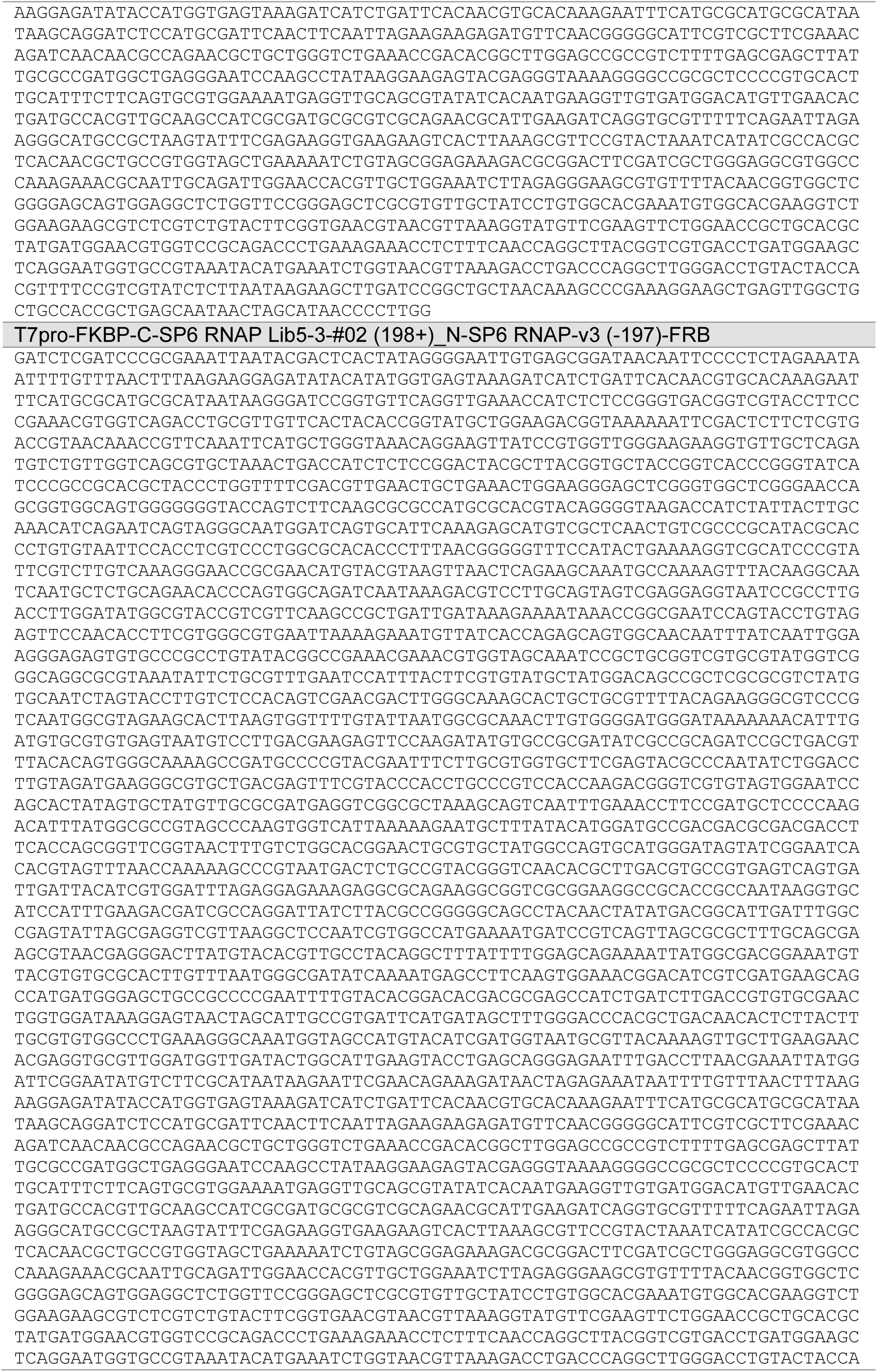
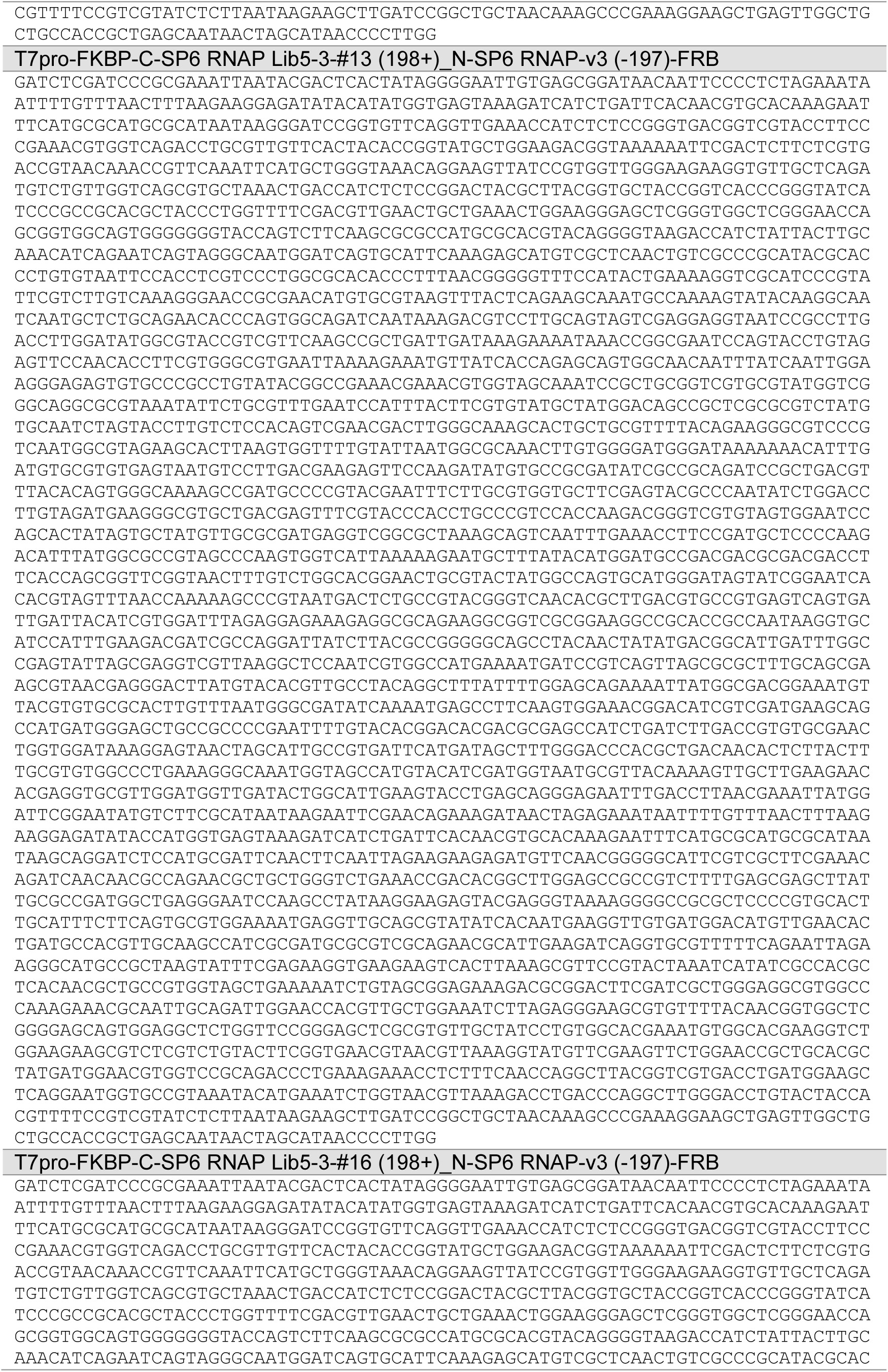
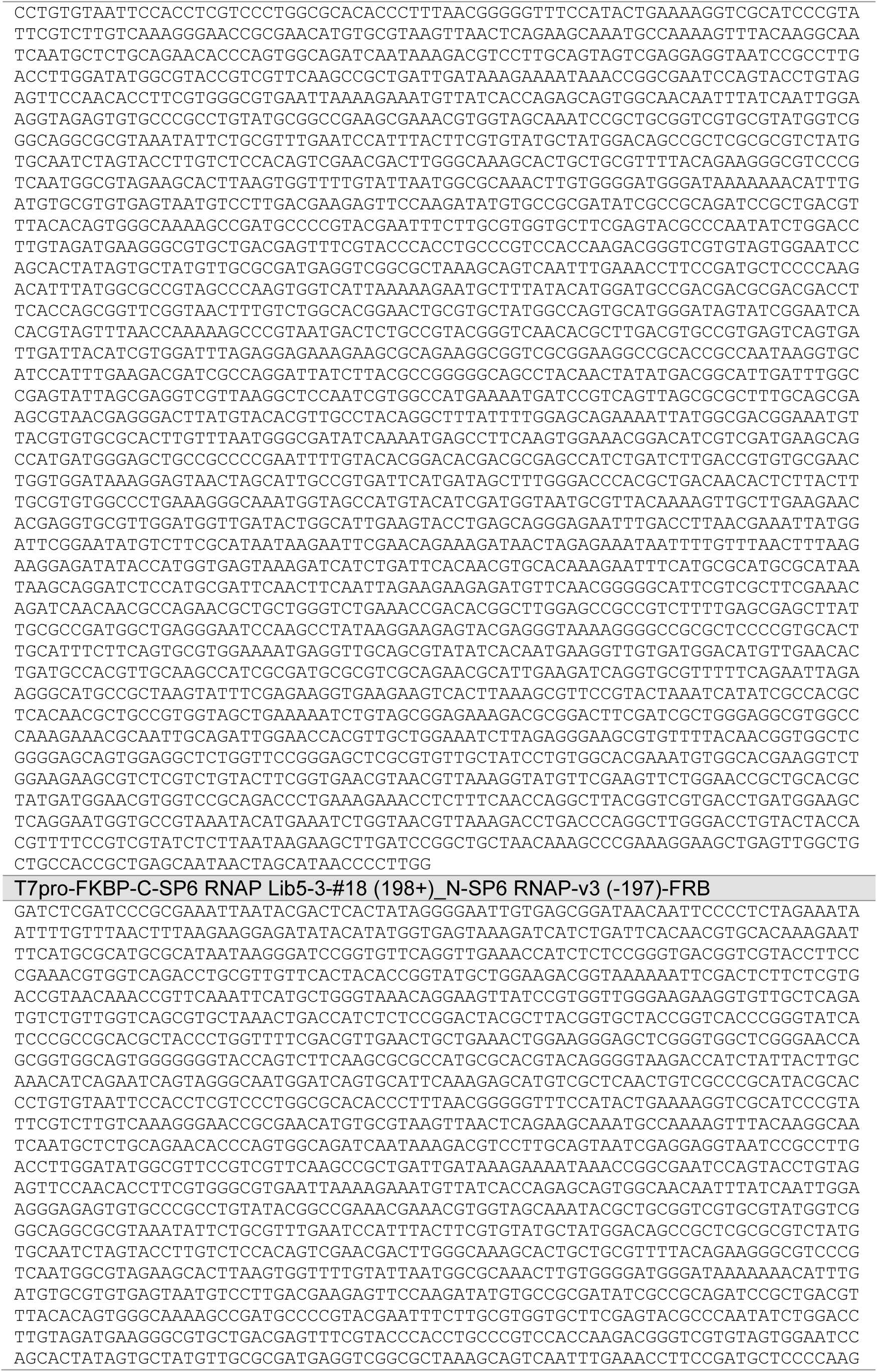
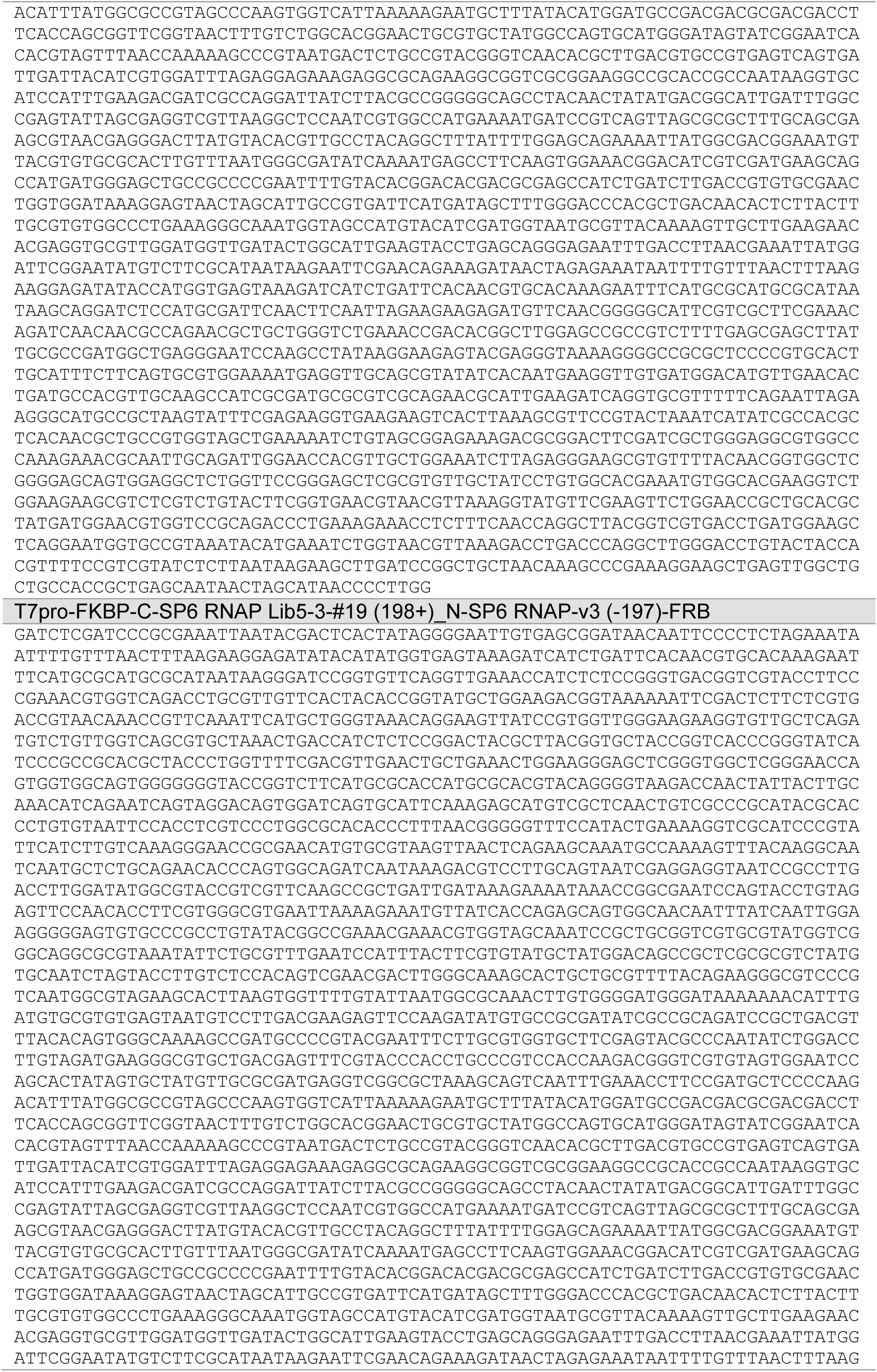
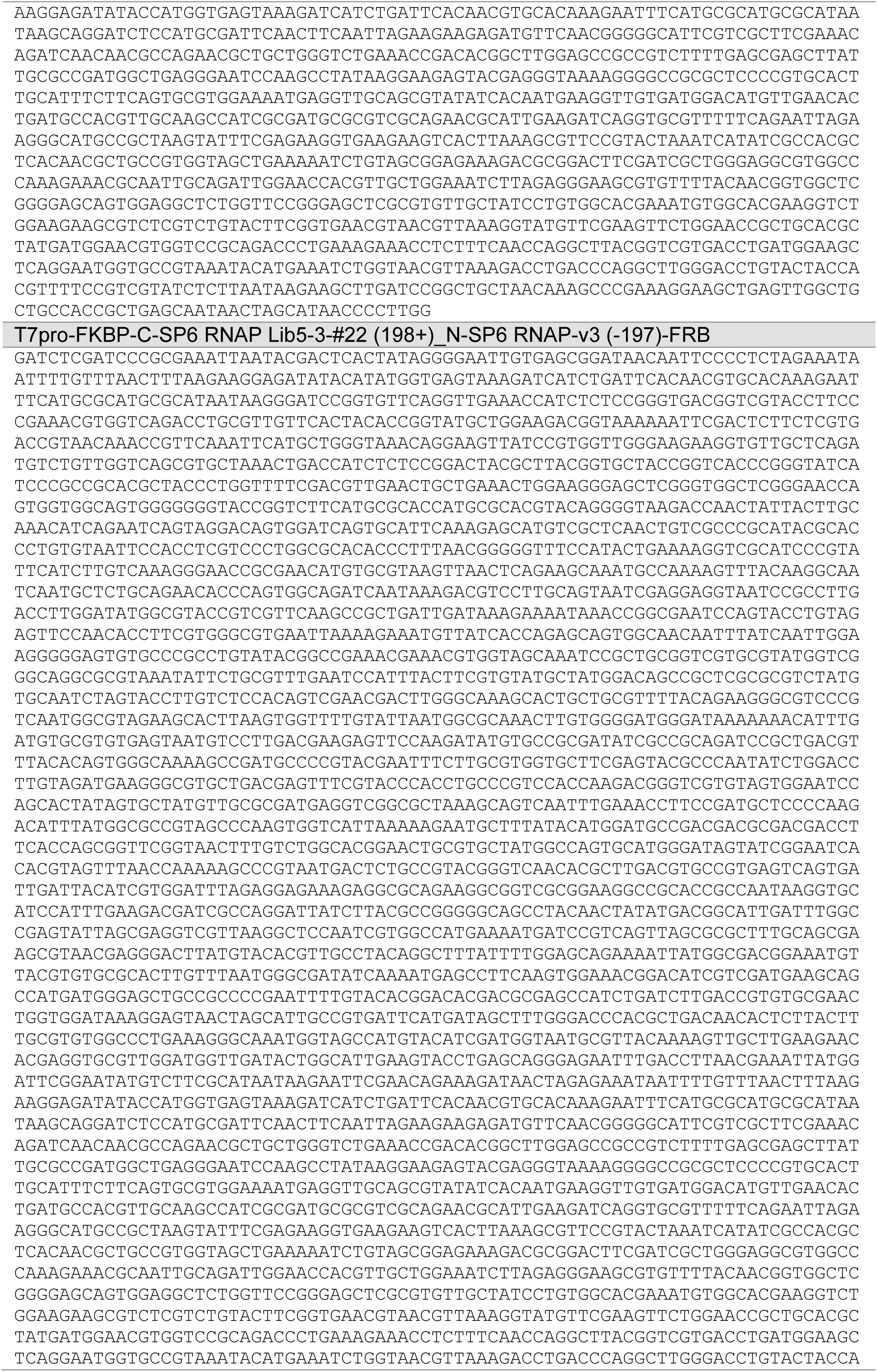
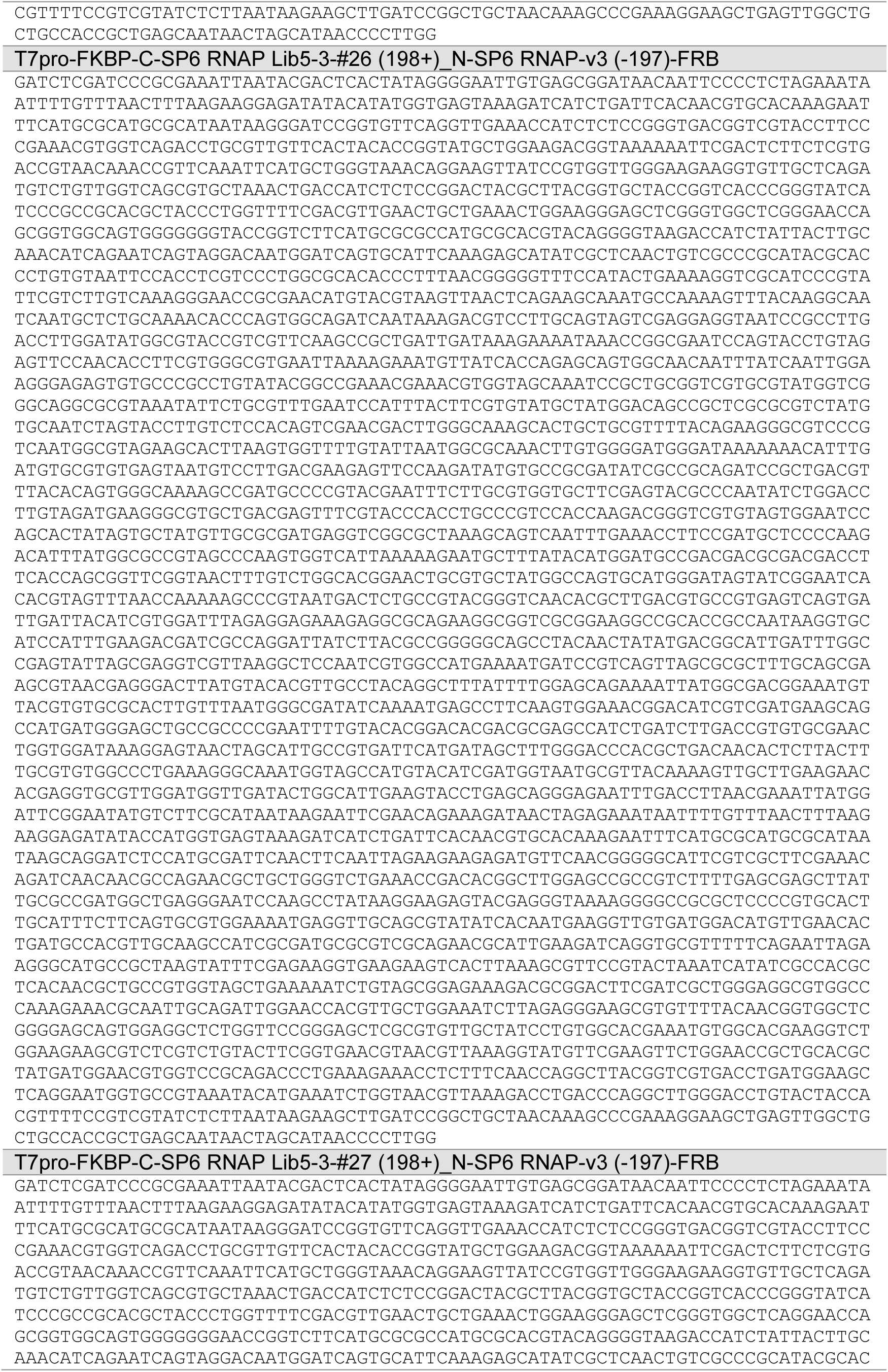
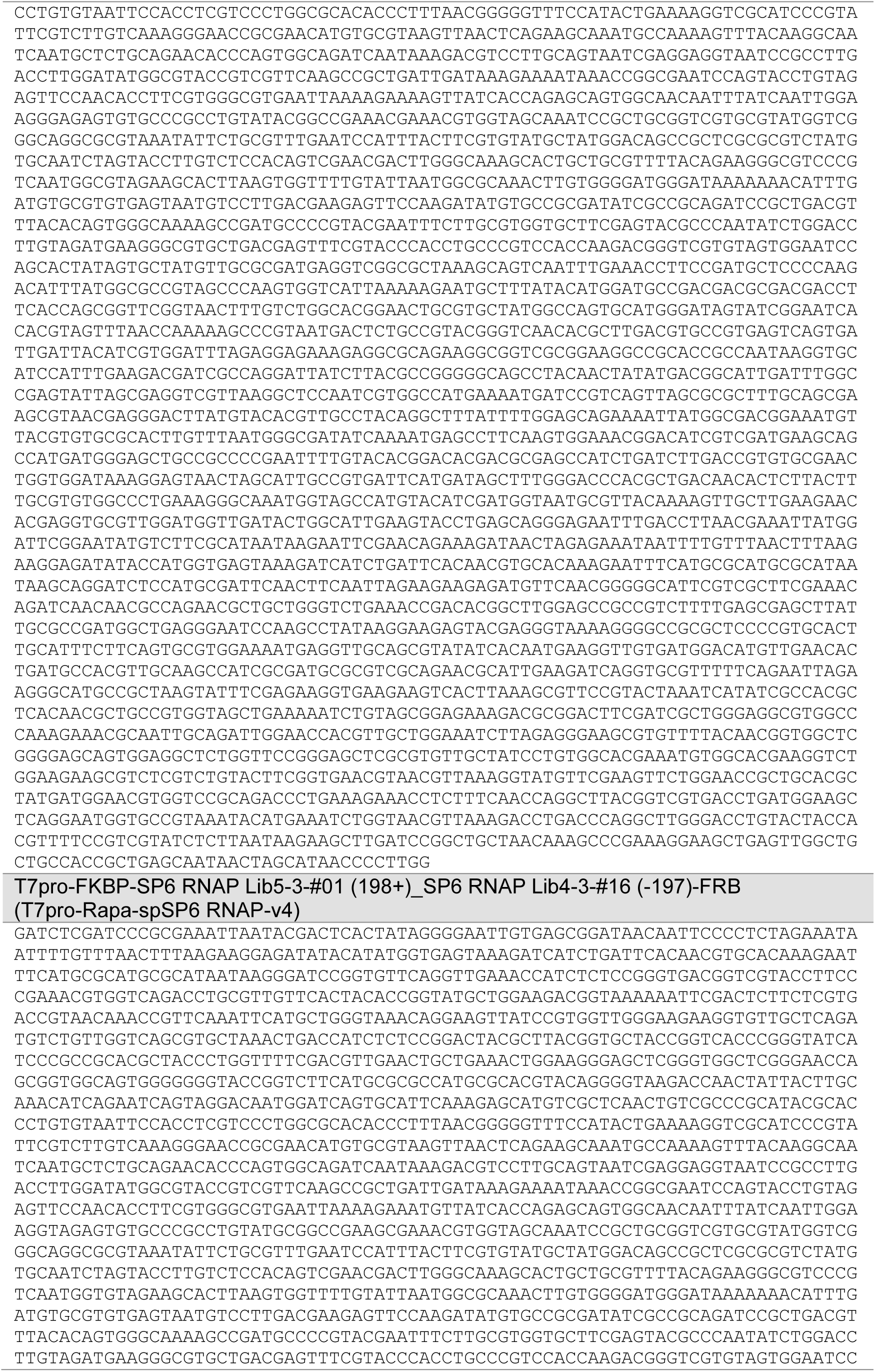
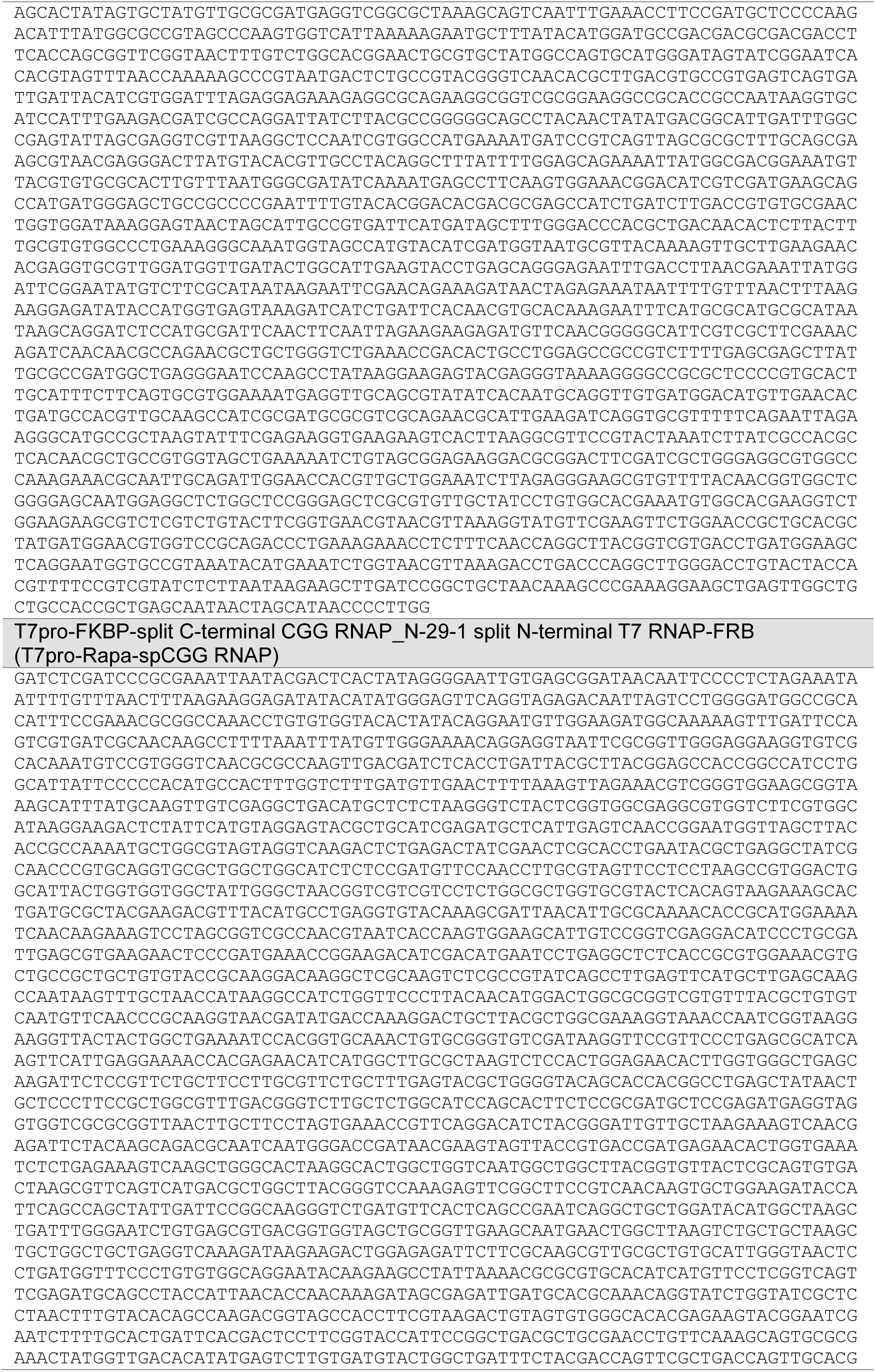
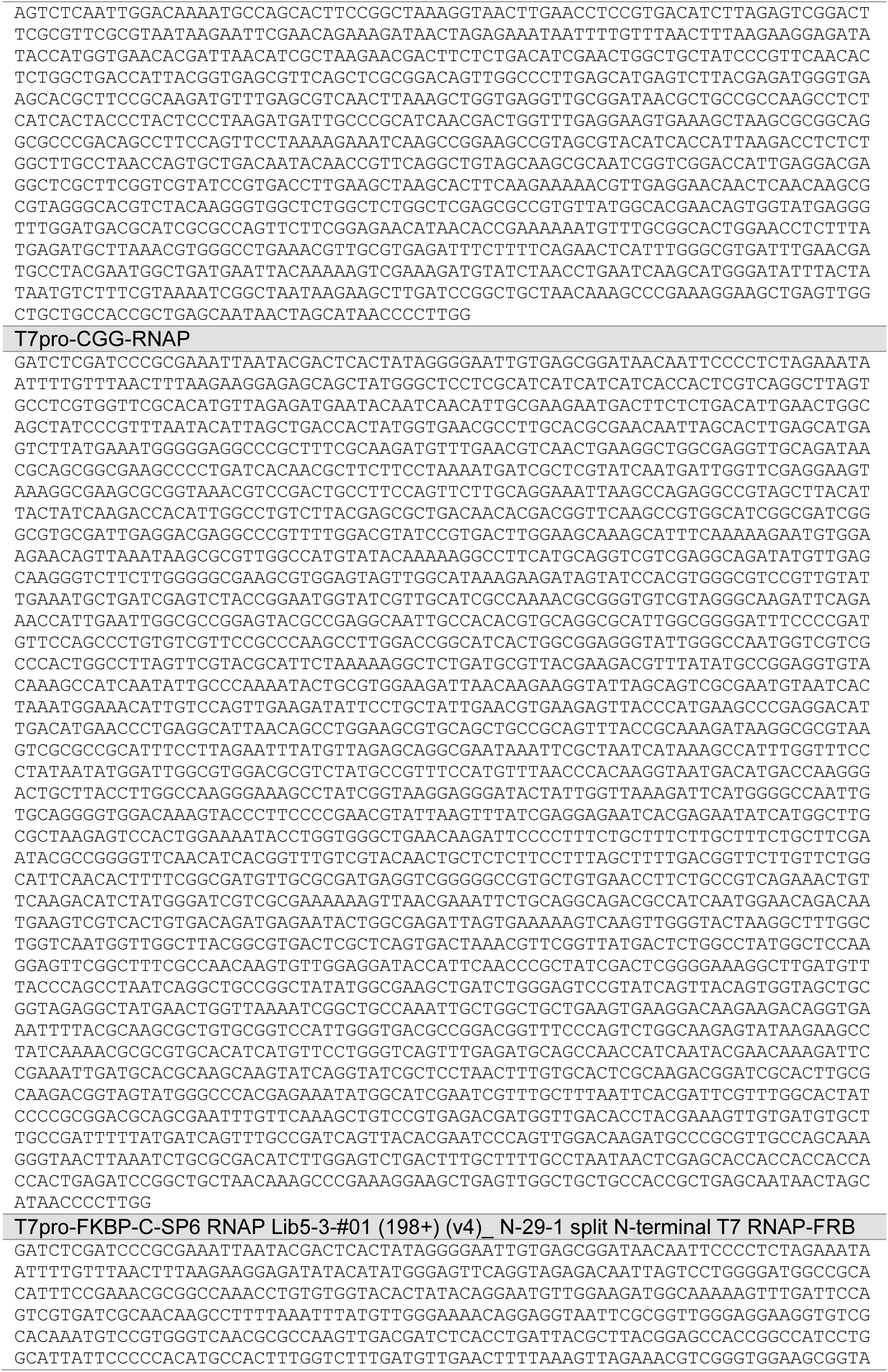
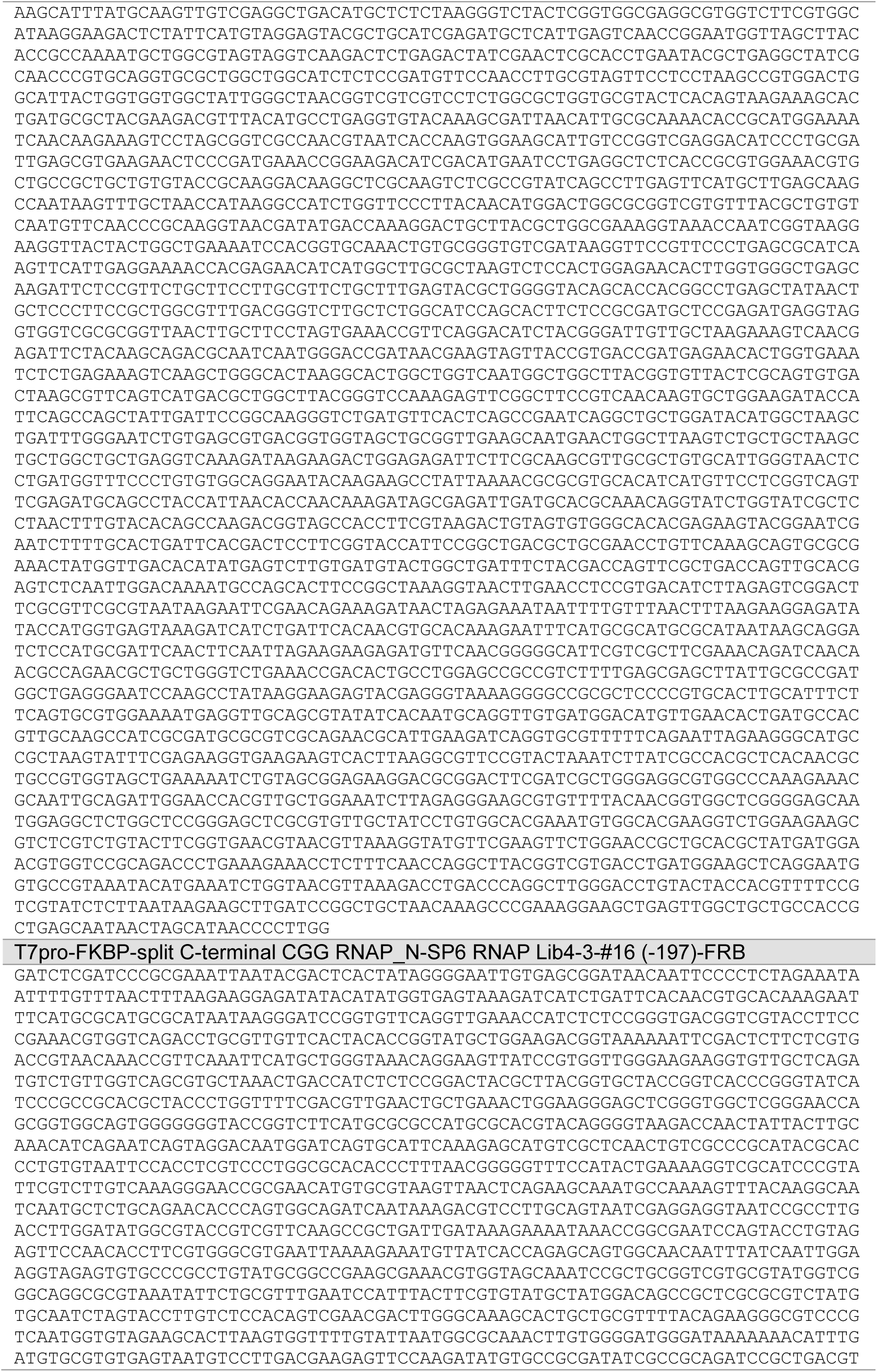
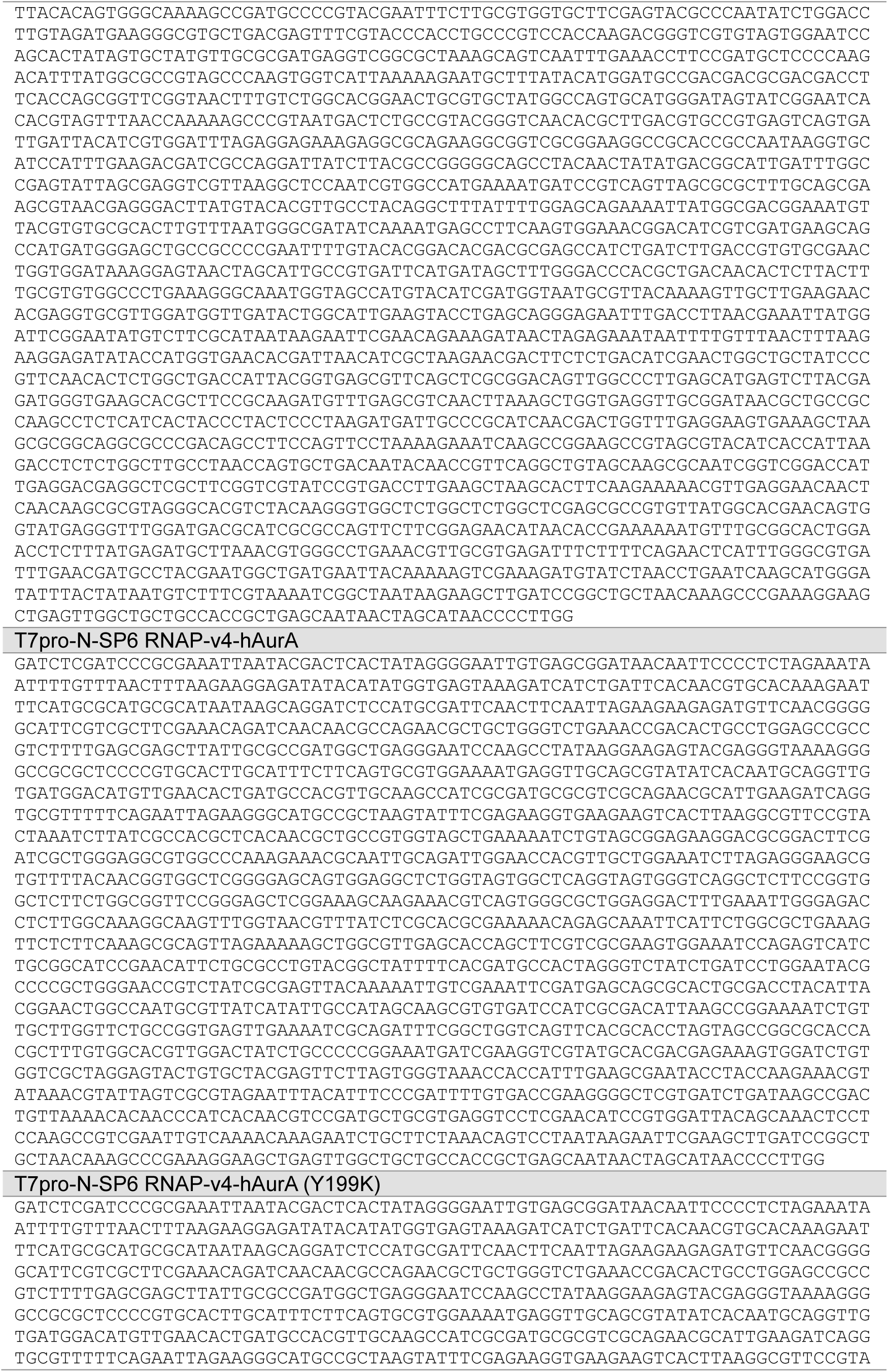
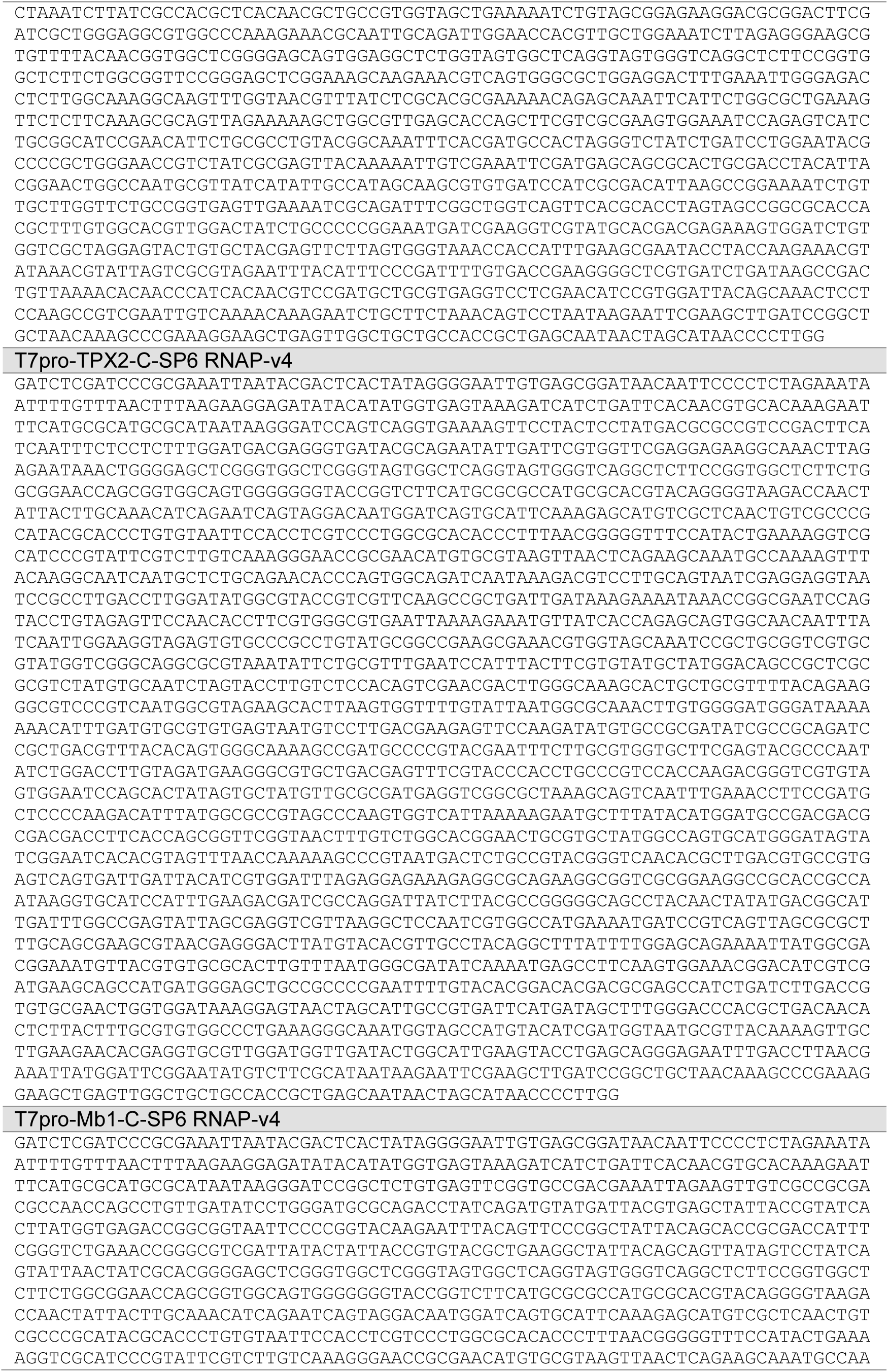
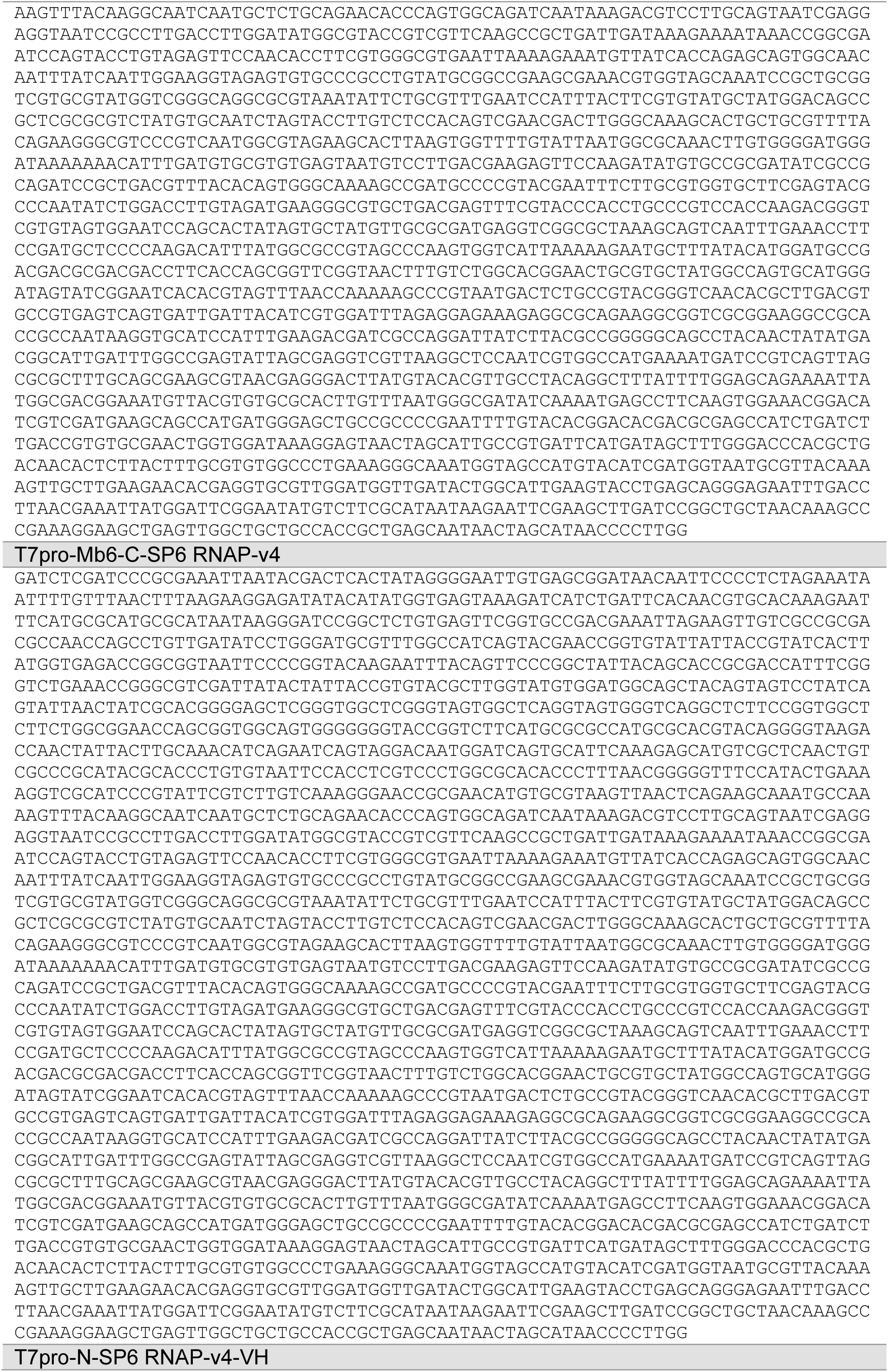
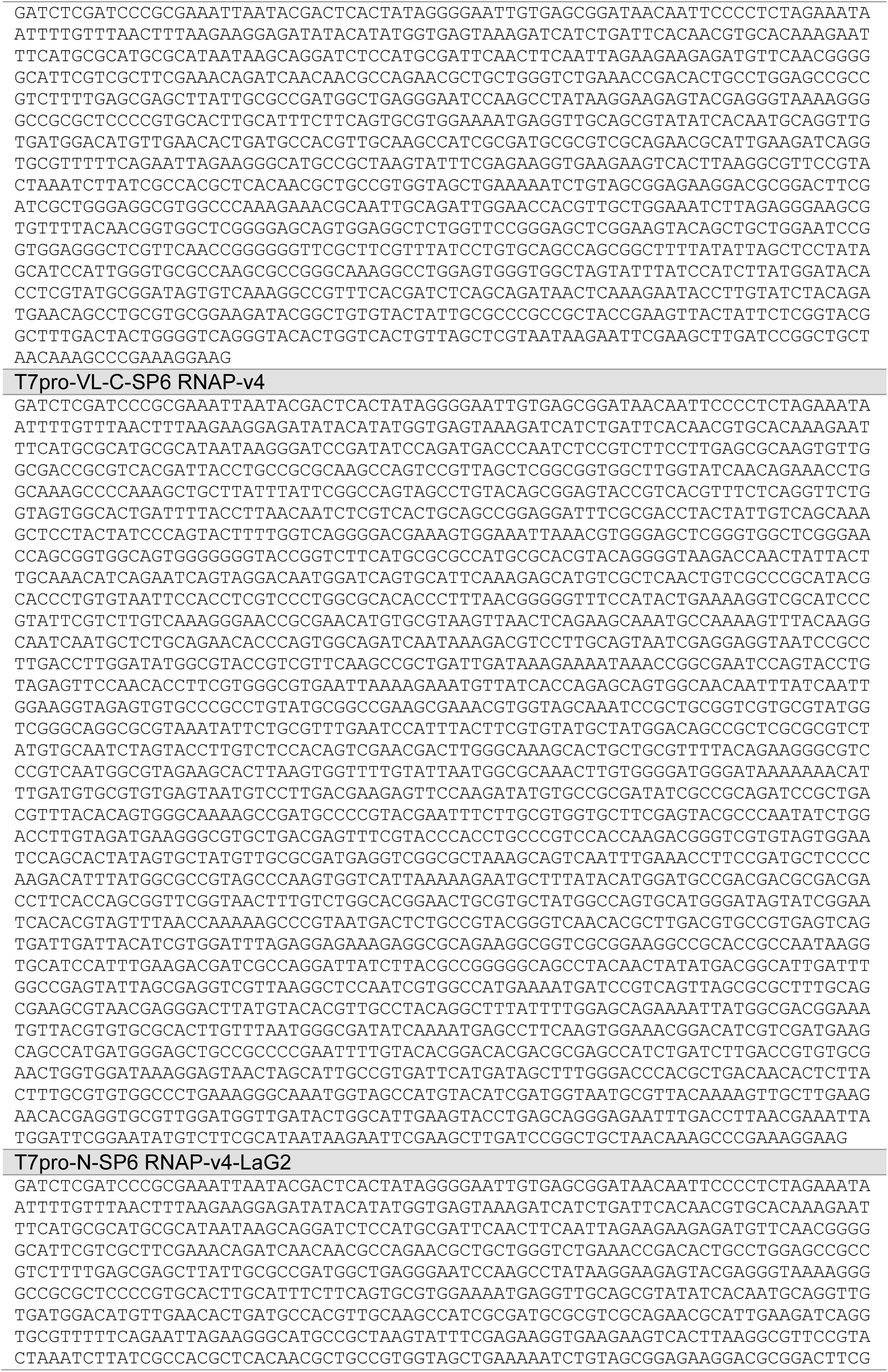
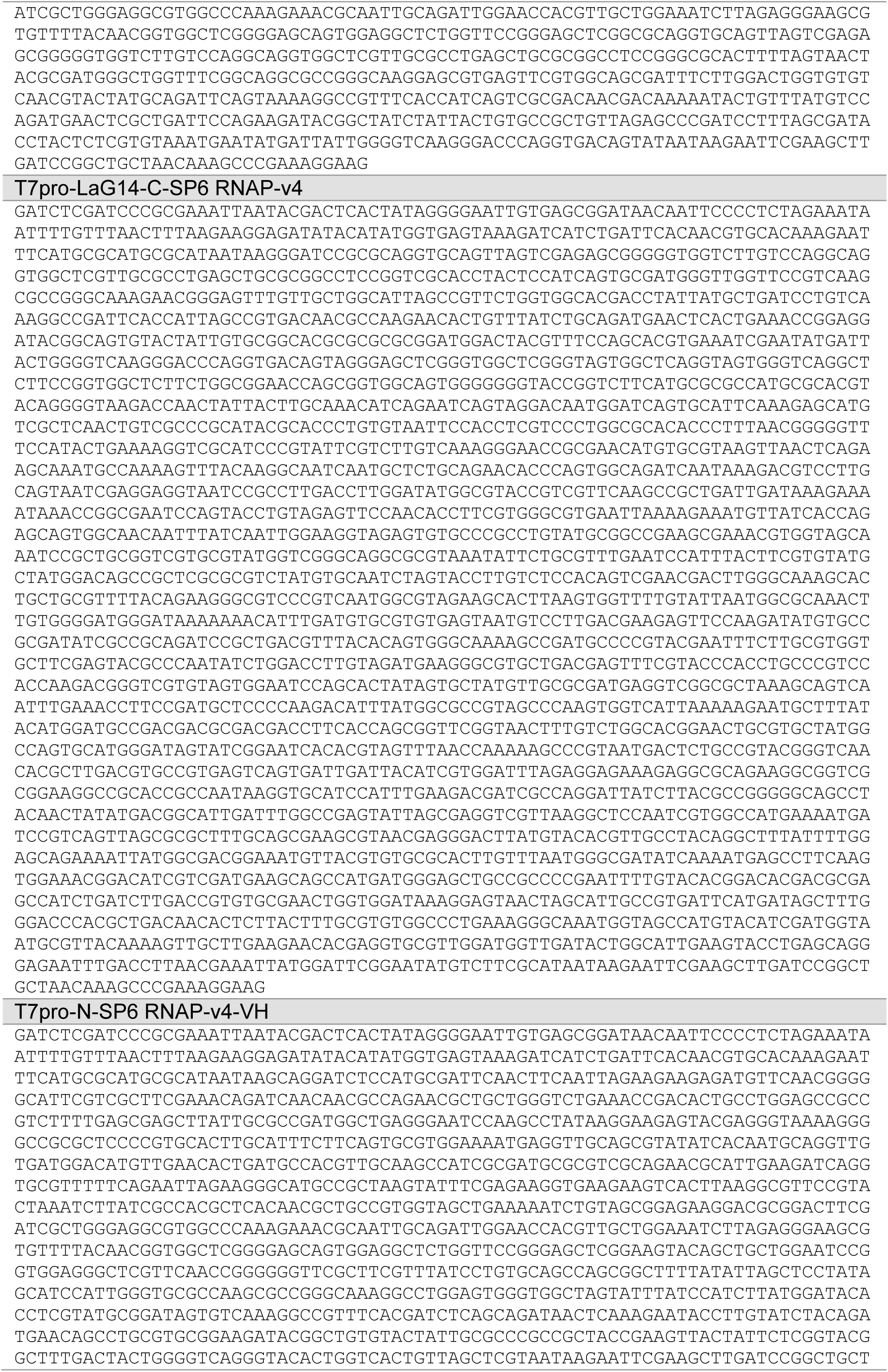
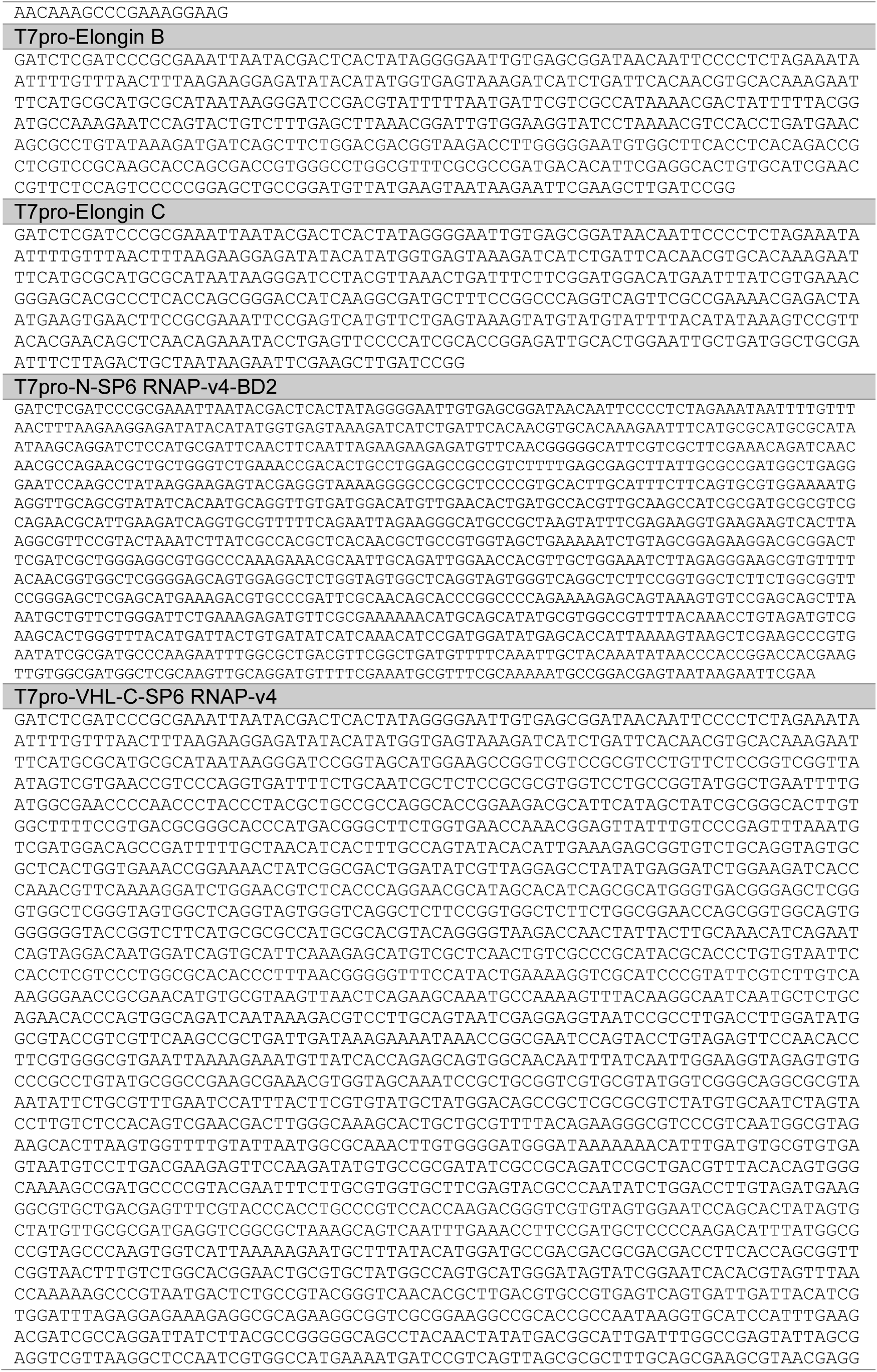
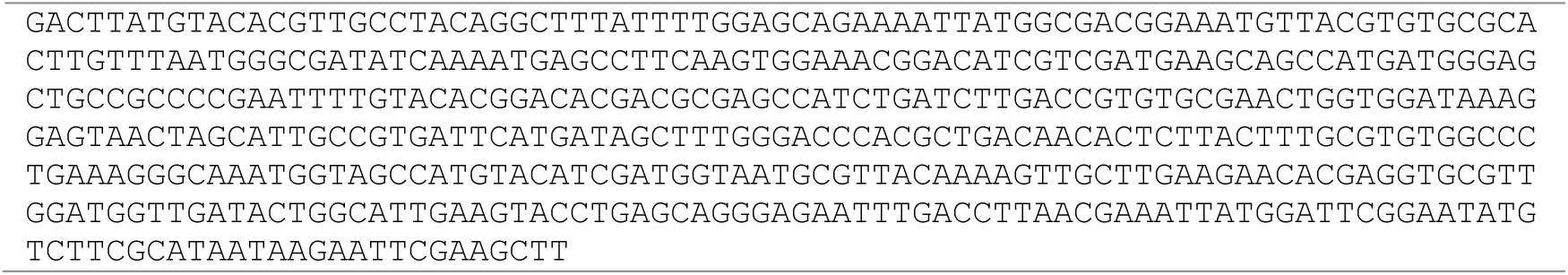
Nucleotide sequences of DNA templates for cell-free translation.

**Supplementary Table 4:**
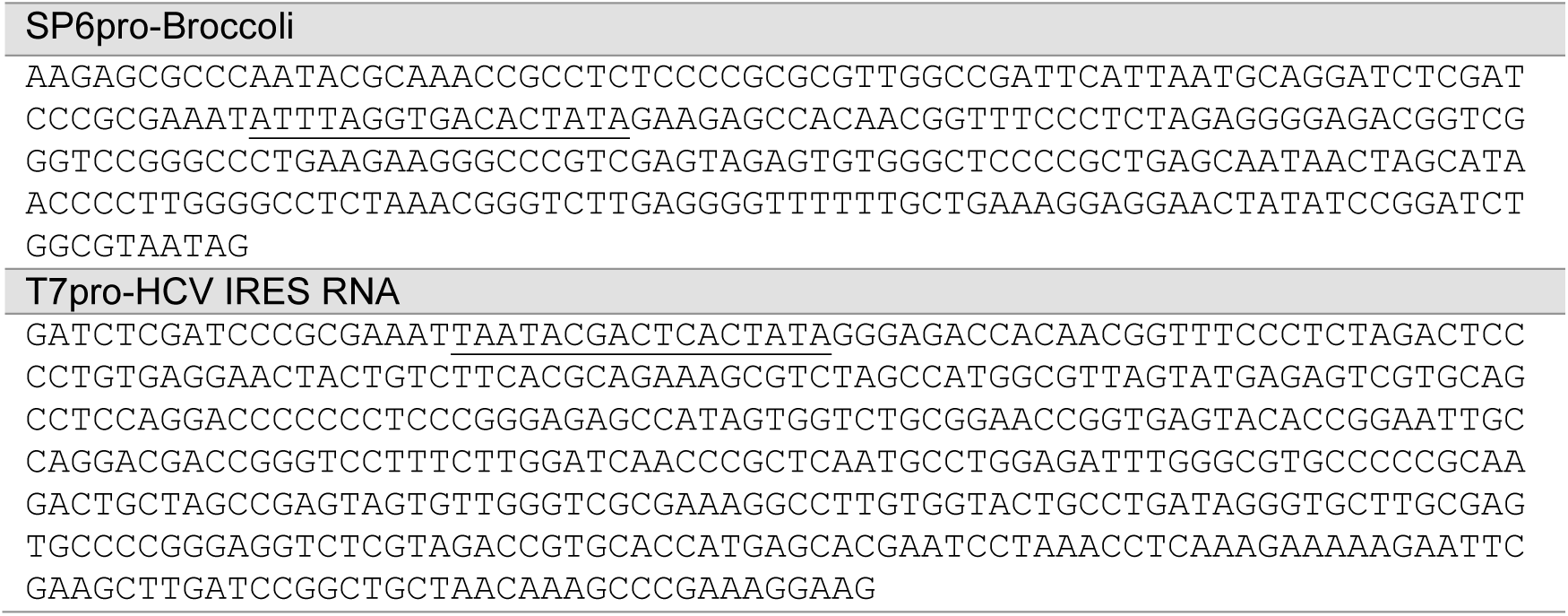
Nucleotide sequences of DNA templates for *in vitro* transcription. Transcription promoter sequences are underlined.

### Extended Data Figures

**Extended Data Fig. 1:**
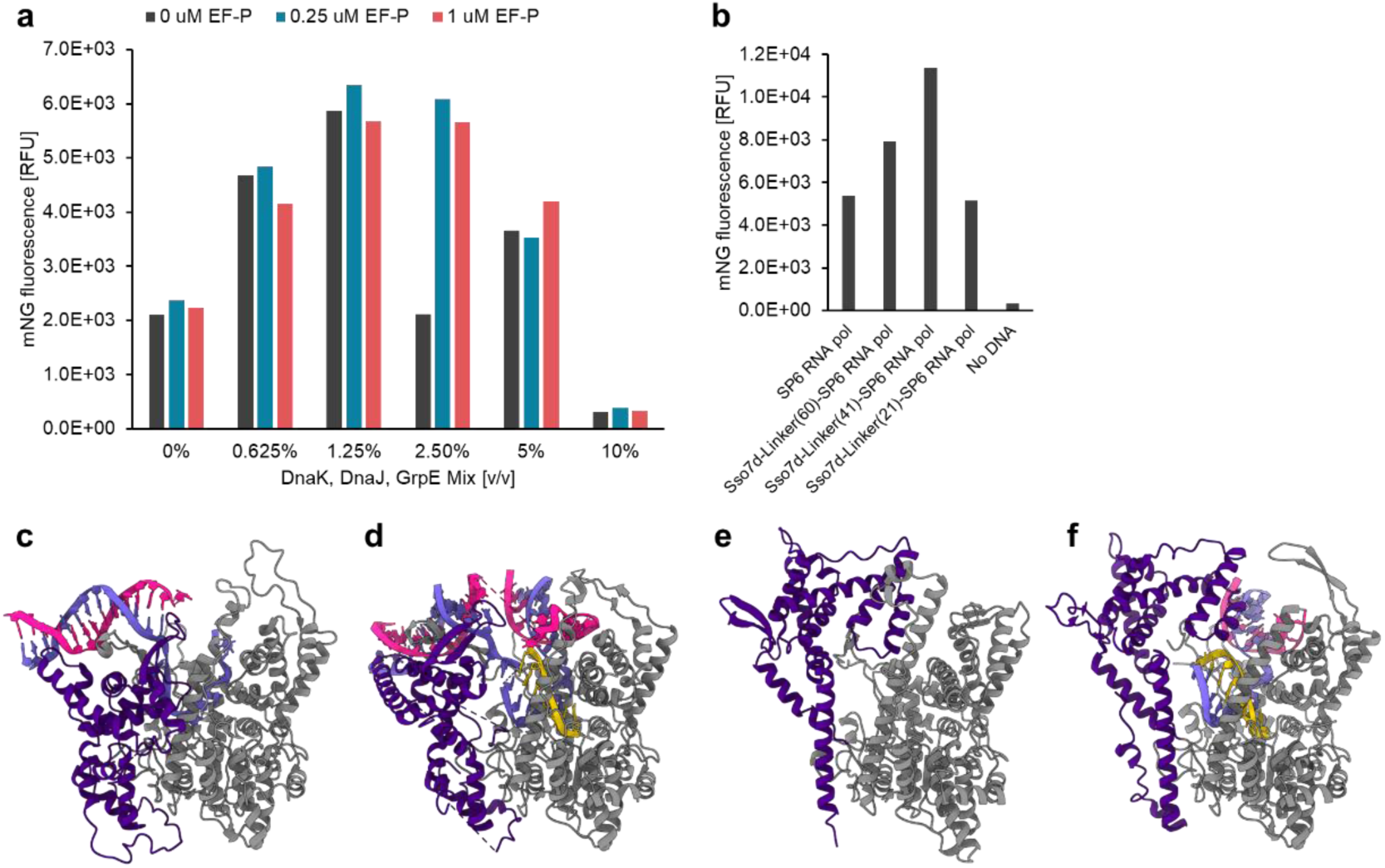
Optimization of IVTT reaction to express SP6 RNA polymerase. **(a)** Optimization of elongation factor P (EF-P) and chaperone conditions. **(b)** Optimization of the linker length between Sso7d and SP6 RNA polymerase. **(c)** Crystal structure of T7 RNA polymerase initiation state (PDB ID: 2PI4). **(d)** Crystal structure of T7 RNA polymerase intermediate state (PDB ID: 3E2E). **(e)** Crystal structure of T7 RNA polymerase elongation state (PDB ID: 1H38). **(f)** Predicted structure of SP6 RNA polymerase by ColabFold. N-terminal domains of RNA polymerases (Residues 1-240 in SP6 RNAP, residues 1-266 in T7 RNAP) are colored in purple, other regions are colored in gray. RNAs are colored in yellow and template DNAs are colored light purple, and non-templated DNAs are in pink.

**Extended Data Fig. 2:**
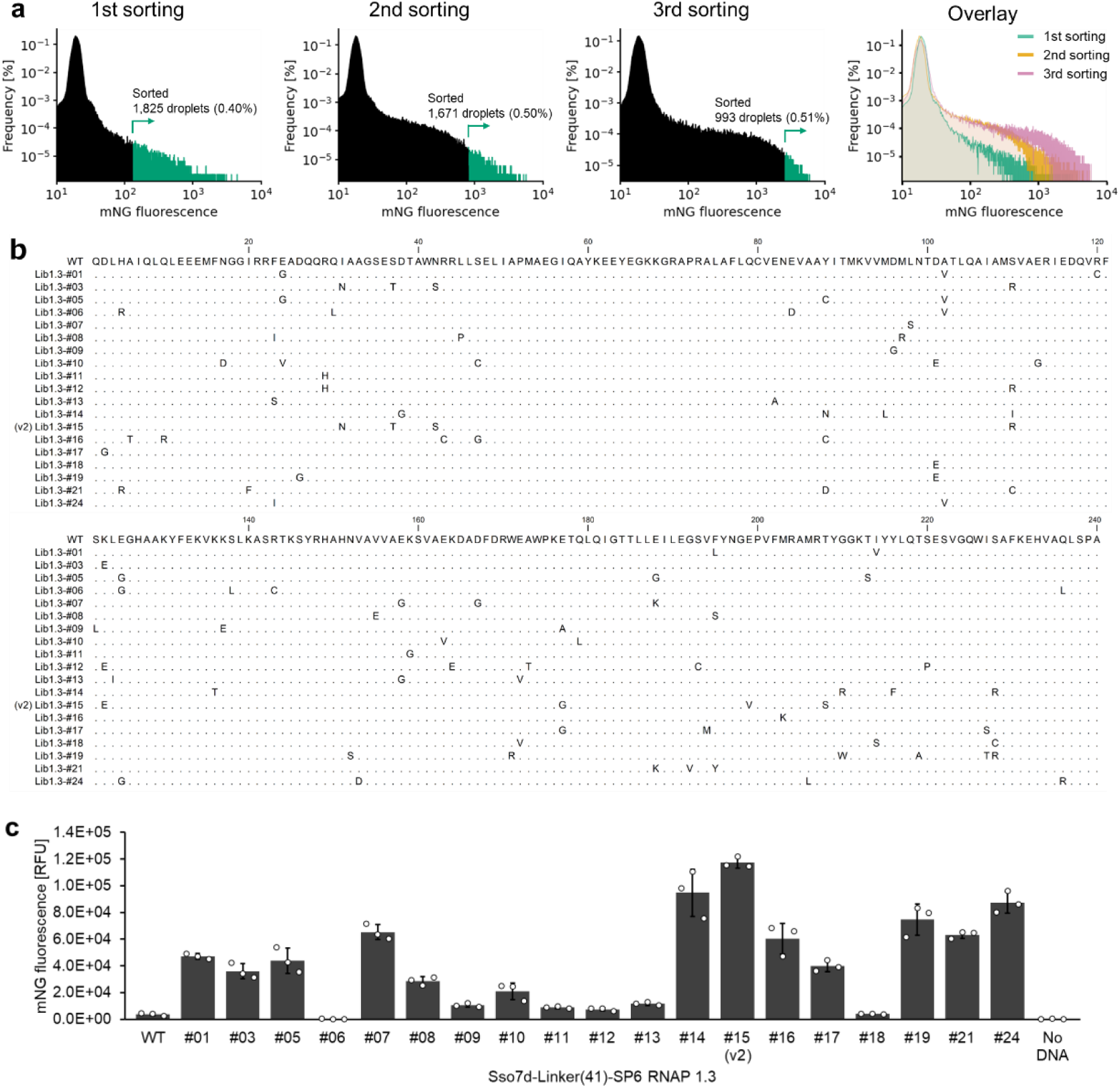
Directed evolution of SP6 RNAP from library 1. **(a)** Histograms of the fluorescence signal distribution of droplets screened for SP6 RNA polymerase in fluorescence-activated droplet sorting (FADS). The overlaid histogram is the same as Fig. 1c. **(b)** Sequence alignment of evolved variants identified from Library 1-3. Residue numbers correspond to SP6 RNAP-wt (UniProt: P06221). The same residues relative to SP6 RNAP-wt are shown in dots. **(c)** Fluorescence of mNeonGreen expressed by Sso7d-SP6 RNAP variants identified from Library 1.3 in boosted PURE system. The data are presented as the mean ± SD from three independent experiments.

**Extended Data Fig. 3:**
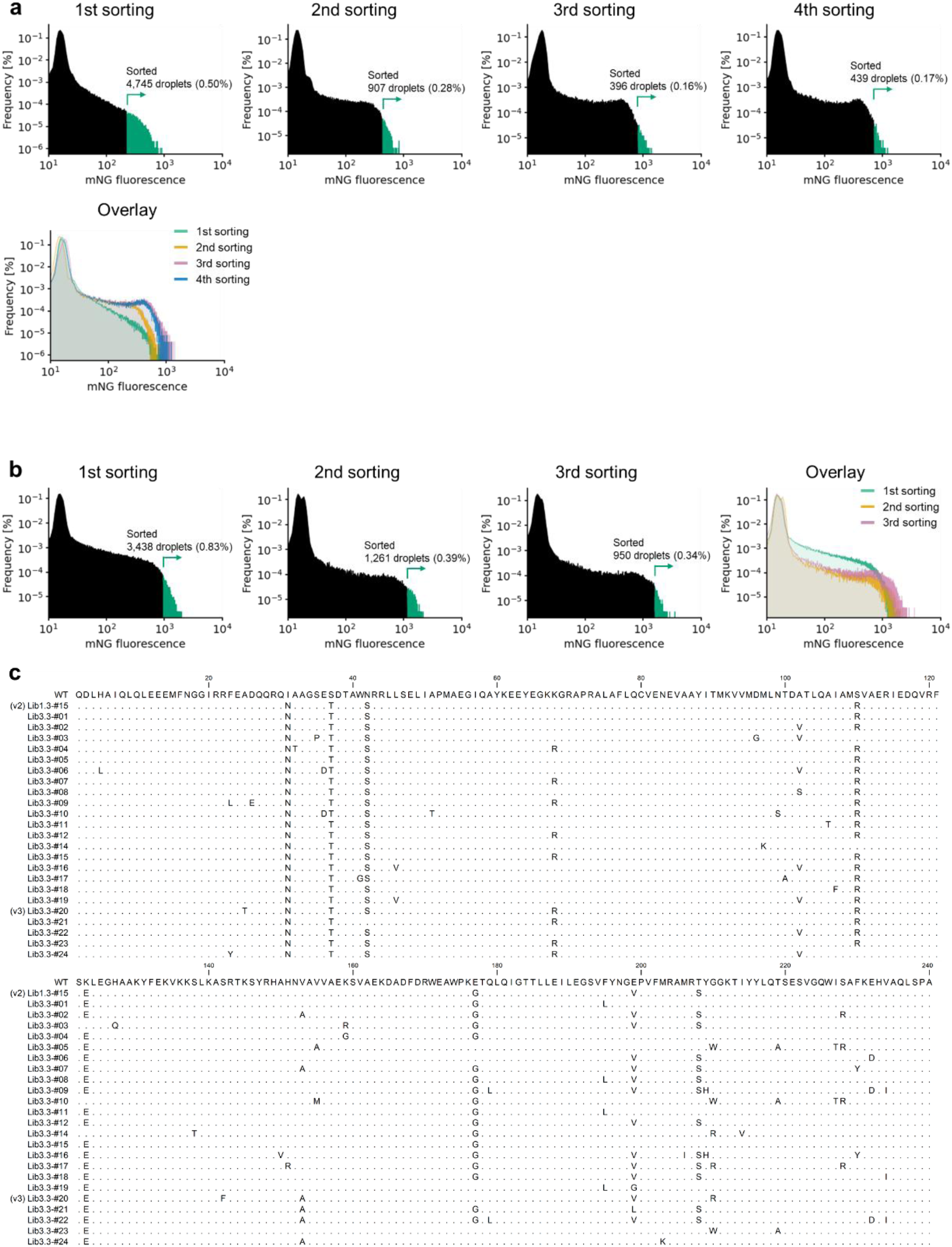
Directed evolution of SP6 RNAP from Library 2 and 3. **(a-b)** Histograms of the fluorescence signal distribution of droplets screened for SP6 RNA polymerase Library 2 or 3 in FADS. Overlaid histograms are the same as those in Fig. 1d and e. **(c)** Sequence alignment of evolved variants identified from Library 3.3. Residue numbers correspond to SP6 RNAP-wt (UniProt: P06221). The same residues relative to SP6 RNAP-wt are shown in dots.

**Extended Data Fig. 4:**
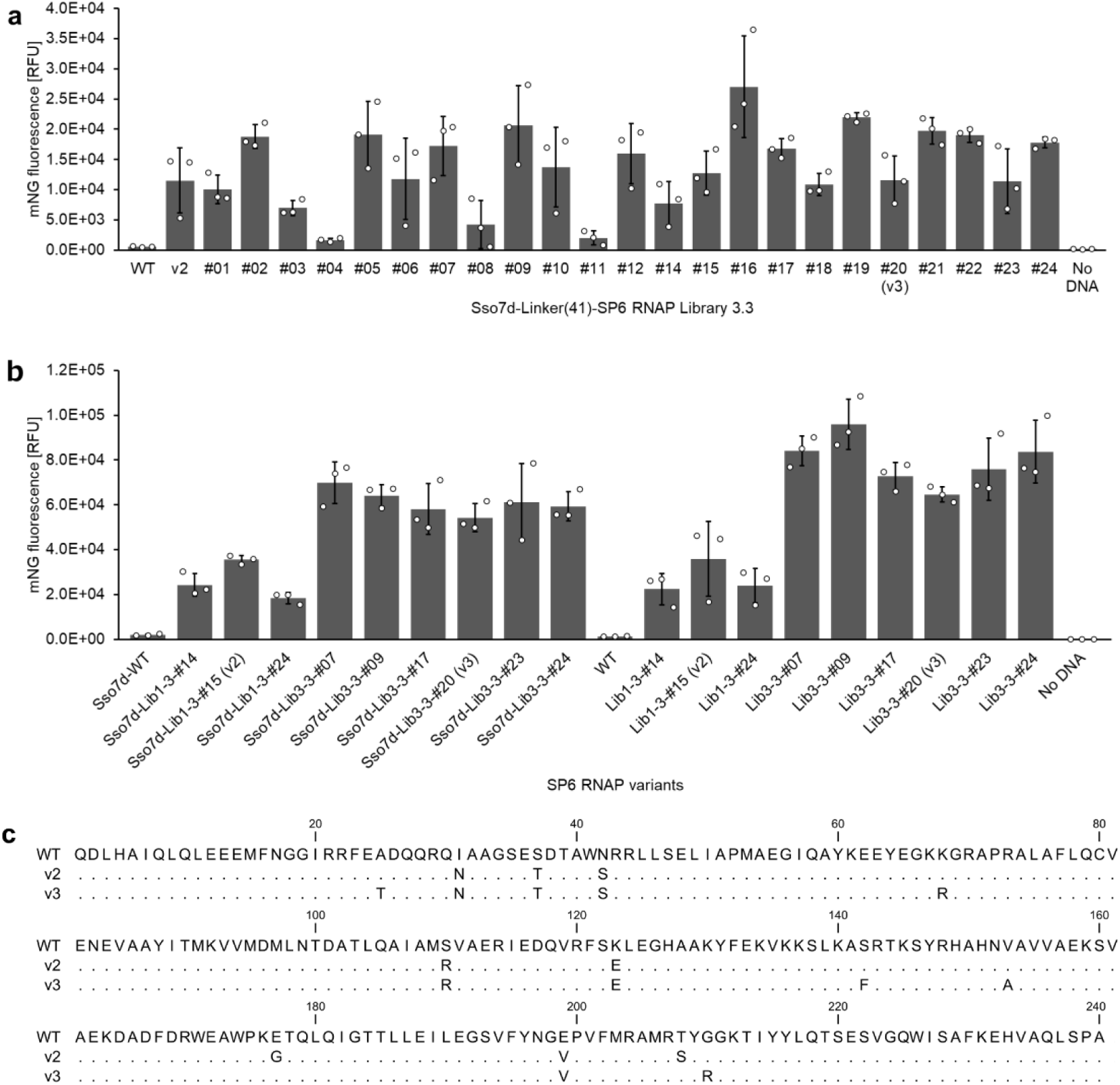
Evaluation of evolved SP6 RNAP. **(a)** The activity measurement of SP6 RNAP variants identified from Library 3.3 by mNG reporter assay in boosted PURE system. **(b)** The activity measurement of SP6 RNAP variants with or without Sso7d by mNG reporter assay in boosted PURE system. **(c)** Sequence alignment of evolved variants identified from Library 3.3. Residue numbers correspond to SP6 RNAP-wt (UniProt: P06221). The same residues relative to SP6 RNAP-wt are shown in dots. The data are presented as the mean ± SD from three independent experiments.

**Extended Data Fig. 5:**
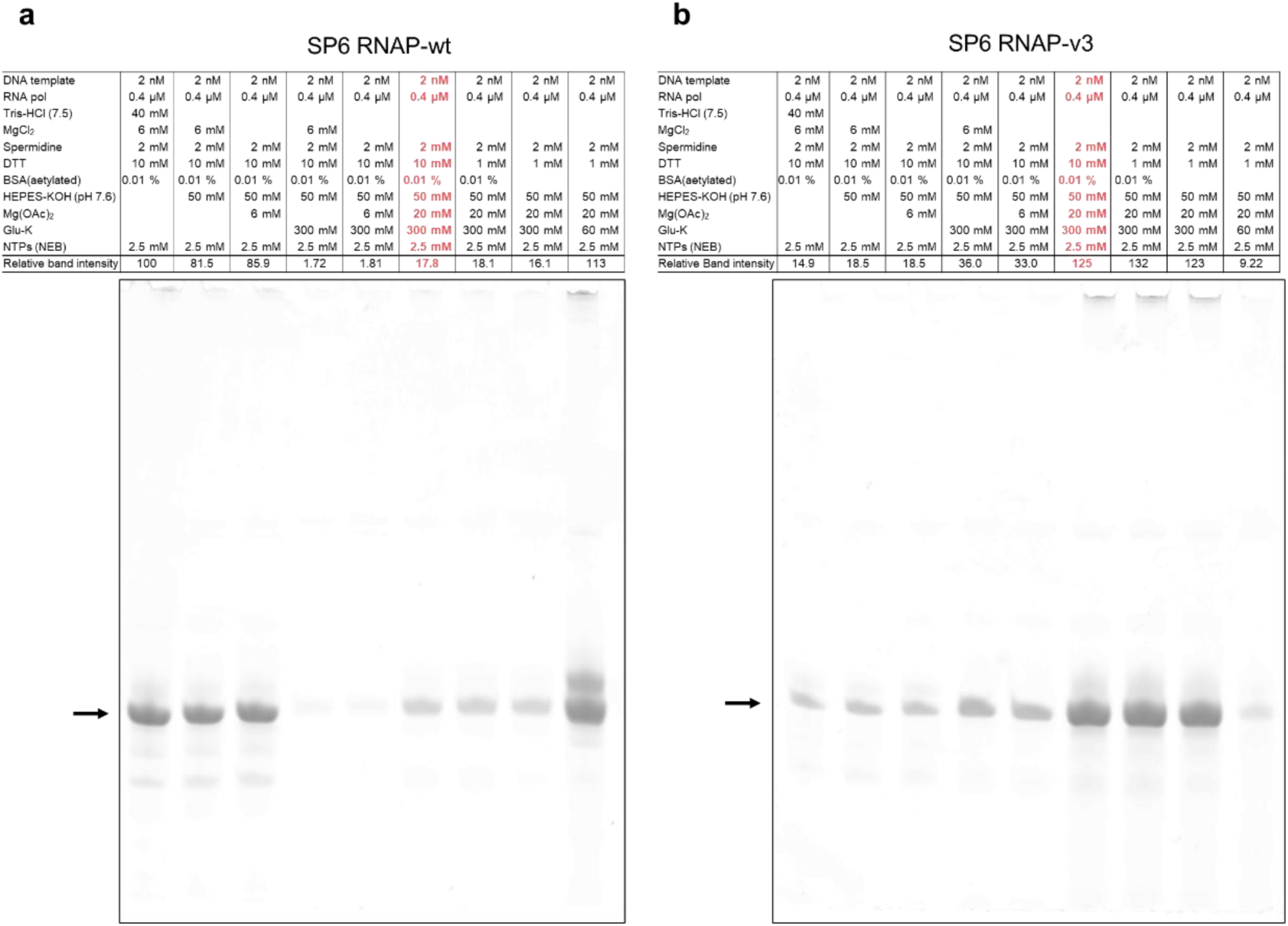
*In vitro* transcription by SP6 RNAP. RNA products transcribed *in vitro* by SP6 RNAP-wt (**a**) or SP6 RNAP-v3 (**b**) in different buffer conditions were analyzed by denaturing PAGE. Band intensities of desired RNAs, indicated by black arrows, were quantified and normalized to that of RNA produced by SP6 RNAP-wt in typical transcription buffer. The conditions mimicking the PURE system are colored red.

**Extended Data Fig. 6:**
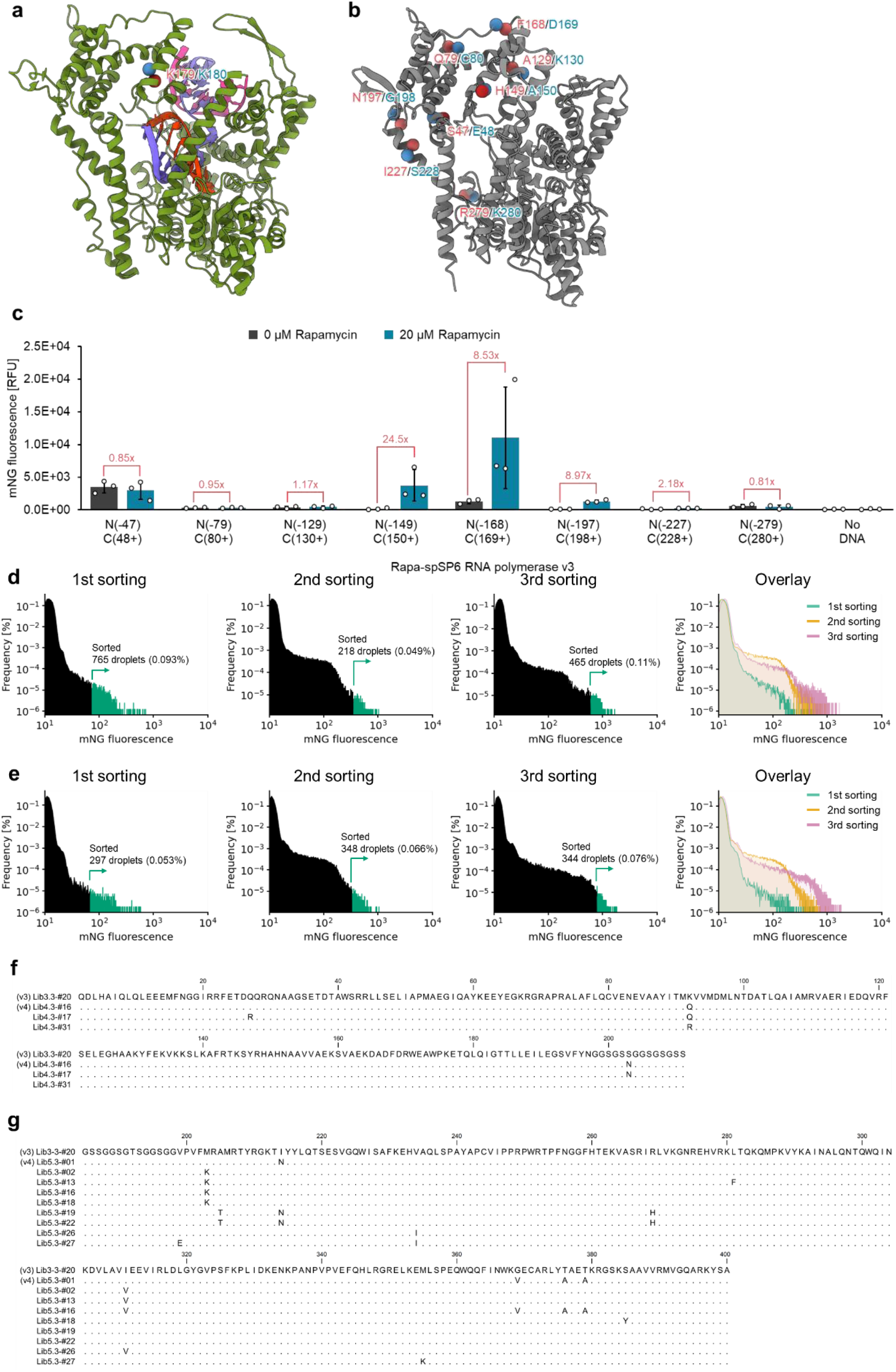
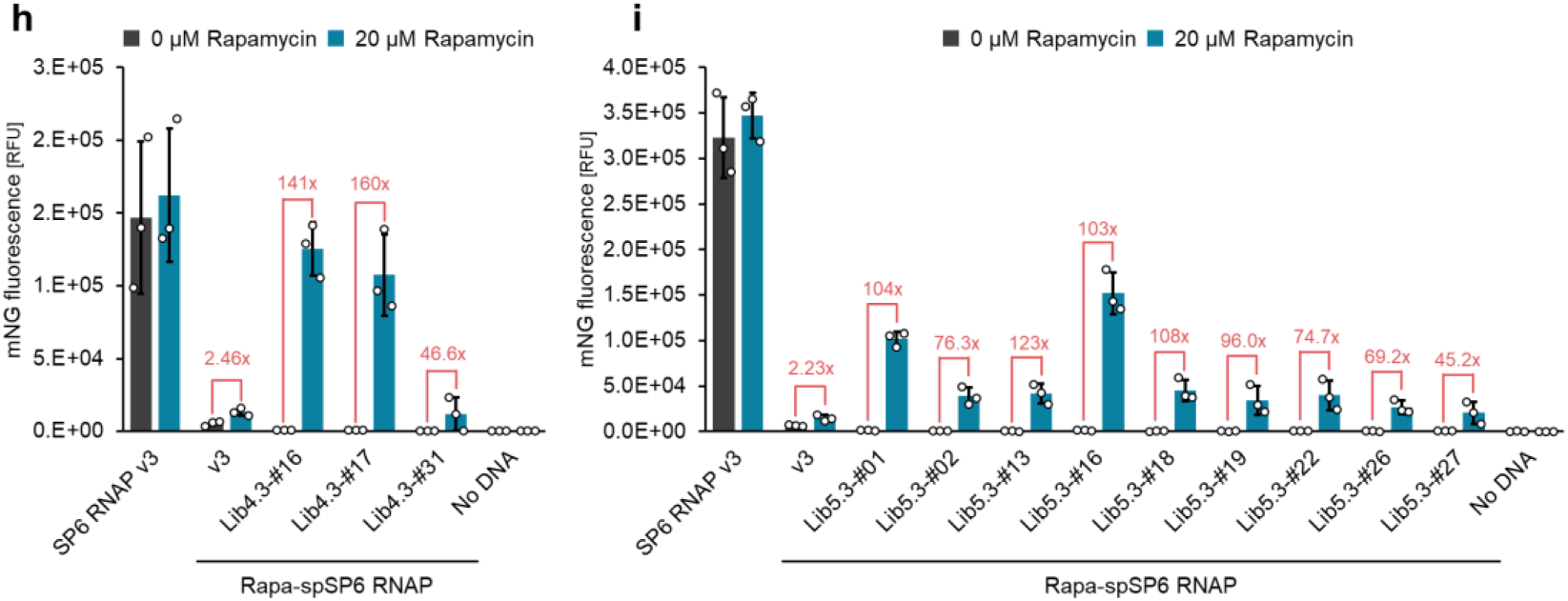
Cell-free evolution of spSP6 RNA polymerase. **(a)** Crystal structure of T7 RNA polymerase elongation state (PDB ID: 1H38). Split site of previously reported spT7 RNAP is between K179 (red sphere) and K180 (blue sphere). Protein is colored in green, RNAs are colored orange, template DNAs are colored light purple, and non-templated DNAs are in pink. **(b)** Predicted structure of SP6 RNA polymerase by ColabFold. Split sites tried in this study are shown in red and blue spheres. **(c)** Rapamycin-induced spSP6 RNAP activities measured by mNG reporter assay in PURE system. The data are presented as the mean ± SD from three independent experiments. **(d, e)** Histograms of the fluorescence signal distribution of droplets screened for spSP6 RNAP Library 4 (**d**) or Library 5 (**e**) in FADS. Overlaid histograms are the same as Fig. 3b or 3c. **(f, g)** Sequence alignment of evolved variants identified from Library 4.3 (**f**) and Library 5.3 (**g**). Residue numbers correspond to SP6 RNAP-wt (UniProt: P06221). The same residues relative to SP6 RNAP-v3 are shown in dots. (**h, i**) Rapamycin-induced spSP6 RNAPs’ activity measured by mNG reporter assay in PURE system. The data are presented as the mean ± SD from three independent experiments.

**Extended Data Fig. 7:**
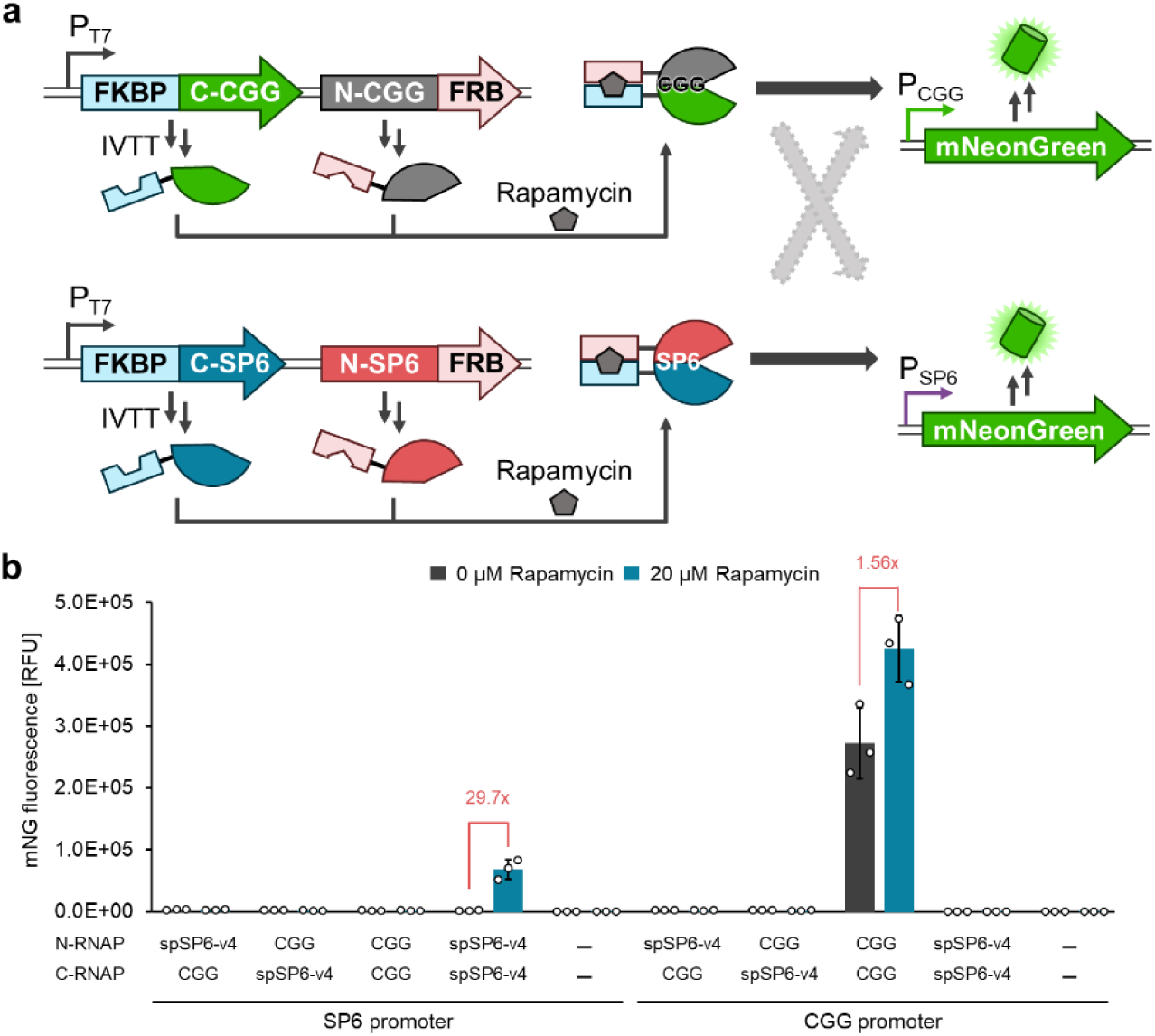
Comparison of evolved spSP6 RNA polymerase and previously reported split T7 RNA polymerase variant. **(a)** Schematic illustration of orthogonality between spCGG RNAP and spSP6 RNAP measured by mNG reporter assay in PURE system. **(b)** Rapamycin-induced split RNAPs activity measured by mNG reporter assay in PURE system. The data are presented as the mean ± SD from three independent experiments.

**Extended Data Fig. 8:**
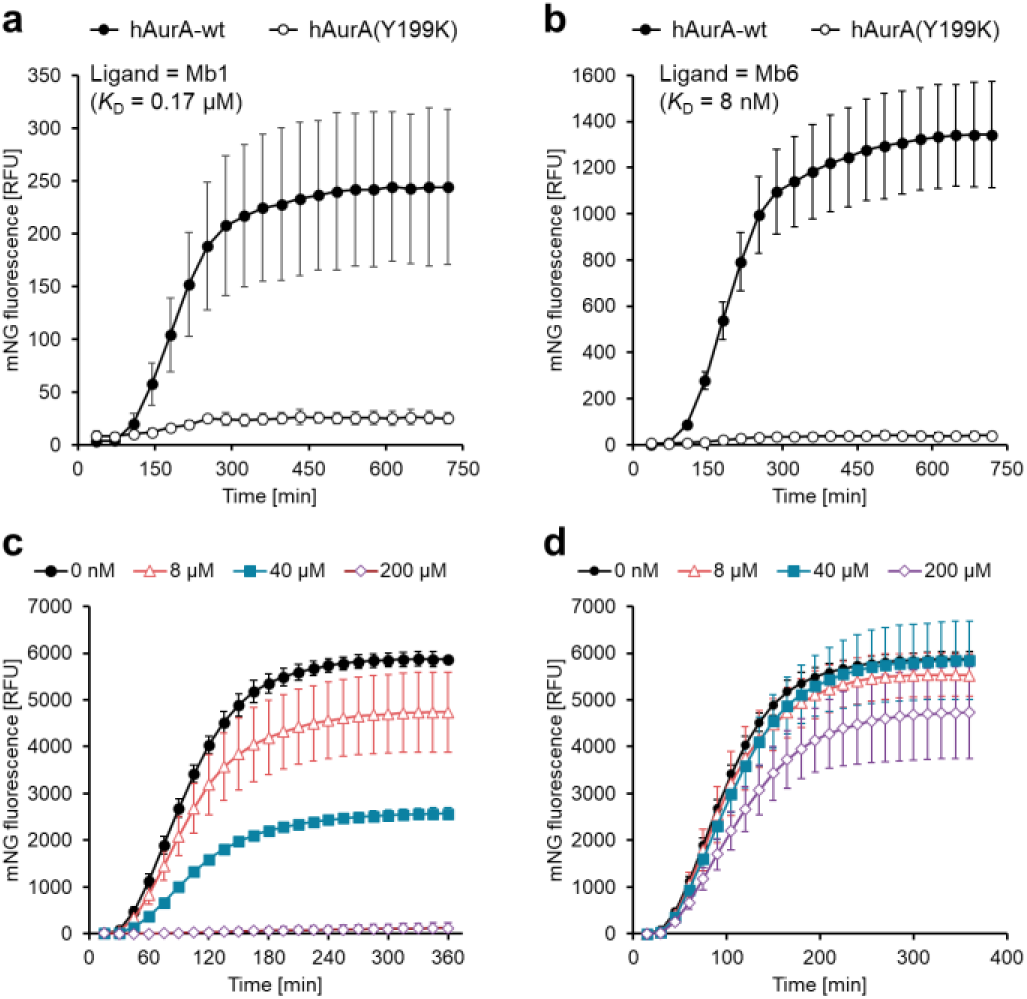
Detection of molecular interactions by spSP6 RNAP-v4. The expression level of reporter fluorescent protein under the SP6 promoter was monitored in a time-dependent manner. **(a, b)** Human Aurora A kinase (hAurA) or its negative mutant (Y199K) was fused to the N-terminal fragment of spSP6 RNAP-v4 as a target, and monobody Mb1 (**a**) or Mb6 (**b**) was fused to the C-terminal fragments of spSP6 RNAP-v4. **(c, d)** Human VHL and Brd4 bromodomain 2 (Brd4^BD2^) were fused to N- and C-terminal fragments of spSP6 RNAP-v4. In the presence of 4 μM MZ1 molecular glue, bromodomain binder (+)-JQ1 (**c**) or VHL binder VH032 (**d**) were added to inhibit the molecular glue activity of MZ1. The data are presented as the mean ± SD from three independent experiments.

**Extended Data Fig. 9:**
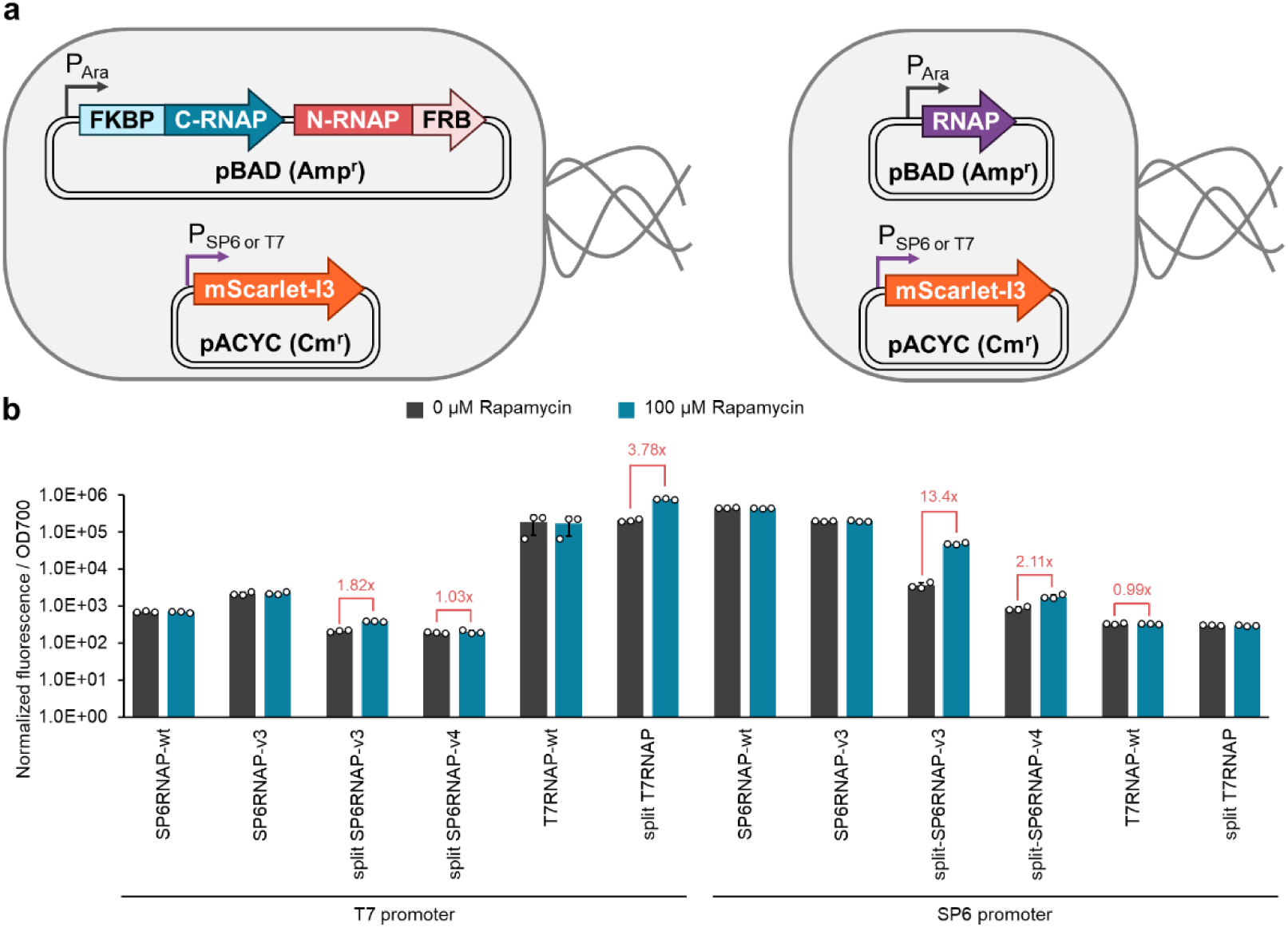
RNA polymerase activity in bacterial cells. **(a)** Schematic illustration of RNAP activity measurement in *E. coli*. **(b)** Rapamycin-induced RNAP activities measured by mScarlet-I3 reporter assay in *E. coli*. The data are presented as the mean ± SD from three independent experiments.

**Extended Data Fig. 10:**
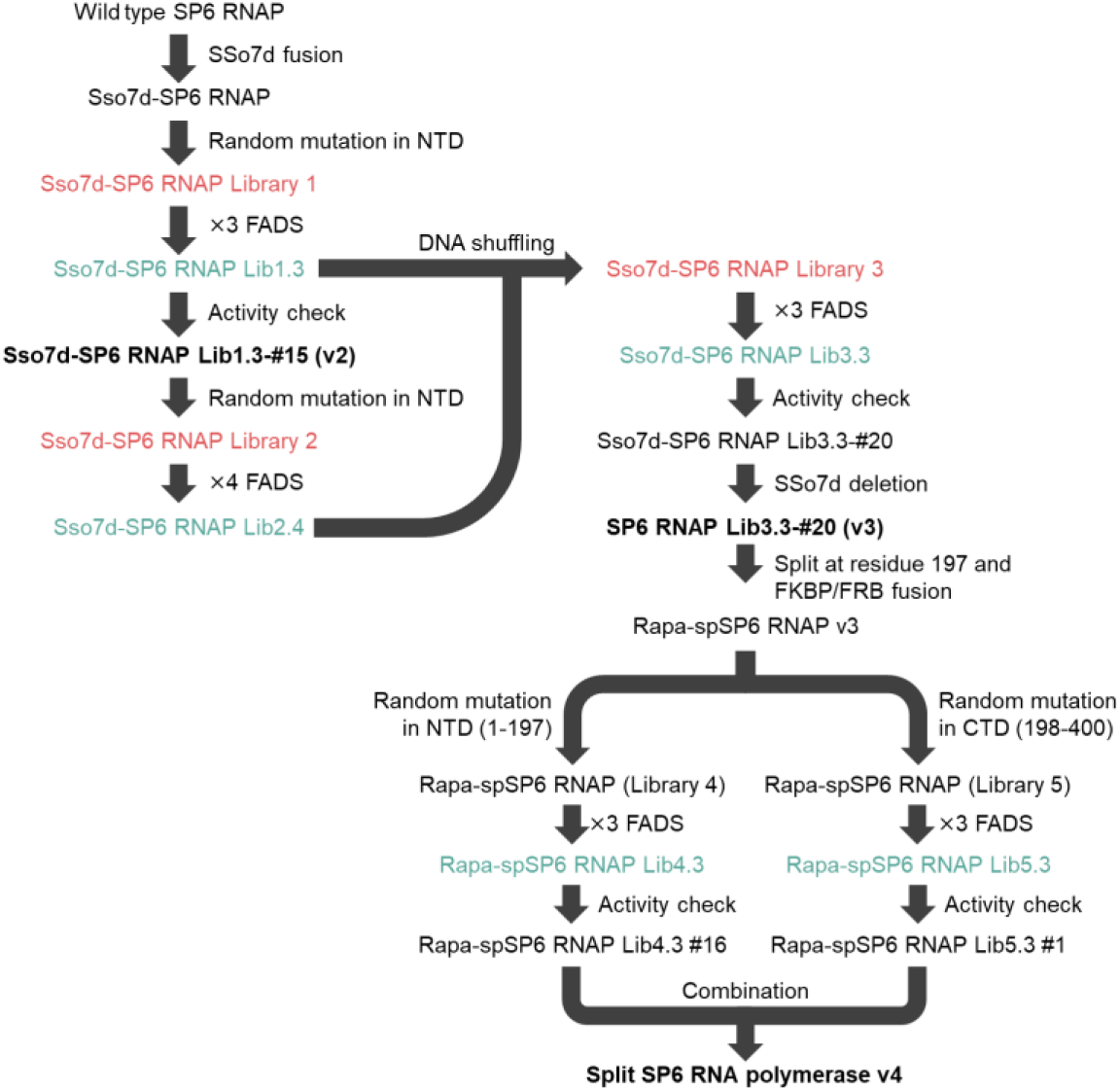
Overview of SP6 RNA polymerase engineering and evolution in this study. Initial libraries for each round are colored red, libraries after the evolution are colored blue, and evolved RNAPs identified after activity measurement are in black bold letters.

## References

1 Y. J. Wang, P. Xue, M. F. Cao, T. H. Yu, S. T. Lane & H. M. Zhao. Directed Evolution: Methodologies and Applications. Chem. Rev. 121, 12384–12444 (2021).

2 A. Jain, S. Stavrakis & A. deMello. Droplet-based microfluidics and enzyme evolution. Curr. Opin. Biotechnol. 87, 103097 (2024).

3 C. Olagnon, J. F. Wardman, F. Liu, H. M. Chen, H. Moon, S. A. Nasseri, D. Seale, P. Rahfeld, S. J. Hallam, J. N. Kizhakkedathu & S. G. Withers. Ultrahigh-Throughput Single Emulsion Droplet Screening for the Discovery of New B Antigen Cleaving Enzymes. ACS Catal. 14, 12884–12894 (2024).

4 A. Autour & C. A. Merten. Fluorescence-activated droplet sequencing (FAD-seq) directly provides sequences of screening hits in antibody discovery. Proc. Natl. Acad. Sci. USA 121, e2405342121 (2024).

5 E. L. Medina, V. A. Maola, M. Hajjar, G. K. Ko, E. J. Ho, A. R. Horton, N. Chim & J. C. Chaput. Rapid evolution of a highly efficient RNA polymerase by homologous recombination. Nat. Chem. Biol. (2026).

6 A. C. Hunt, B. J. Rasor, K. Seki, H. M. Ekas, K. F. Warfel, A. S. Karim & M. C. Jewett. Cell-Free Gene Expression: Methods and Applications. Chem. Rev. 125, 91–149 (2024).

7 Y. Shimizu, A. Inoue, Y. Tomari, T. Suzuki, T. Yokogawa, K. Nishikawa & T. Ueda. Cell-free translation reconstituted with purified components. Nat. Biotechnol. 19, 751–755 (2001).

8. T. Furubayashi, N. Terasaka, K. Tajima & H. Noji. Boosted cell-free gene expression for robust signal readout from a single-copy DNA template in microdroplets. bioRxiv (2026).

9 J. M. Holstein, C. Gylstorff & F. Hollfelder. Cell-free Directed Evolution of a Protease in Microdroplets at Ultrahigh Throughput. ACS Synth. Biol. 10, 252–257 (2021).

10 T. Tabuchi & Y. Yokobayashi. High-throughput screening of cell-free riboswitches by fluorescence-activated droplet sorting. Nucleic Acids Res. 50, 3535–3550 (2022).

11 A. Chen, W. Xu, X. D. Zhang, J. Lao, X. Zhao, K. Milcic & D. A. Weitz. Ultrahigh-Throughput Multiplexed Screening of Purified Protein from Cell-Free Expression Using Droplet Microfluidics. J. Am. Chem. Soc. 147, 28758–28772 (2025).

12 A. R. Abate, T. Hung, P. Mary, J. J. Agresti & D. A. Weitz. High-throughput injection with microfluidics using picoinjectors. Proc. Natl. Acad. Sci. USA 107, 19163–19166 (2010).

13 F. Diehl, M. Li, Y. P. He, K. W. Kinzler, B. Vogelstein & D. Dressman. BEAMing: single-molecule PCR on microparticles in water-in-oil emulsions. Nat. Methods 3, 551–559 (2006).

14 J. Pu, J. Zinkus-Boltz & B. C. Dickinson. Evolution of a split RNA polymerase as a versatile biosensor platform. Nat. Chem. Biol. 13, 432–438 (2017).

15 M. A. McSweeney, A. T. Patterson, K. Loeffler, R. Cuellar Lelo de Larrea, M. P. McNerney, R. S. Kane & M. P. Styczynski. A modular cell-free protein biosensor platform using split T7 RNA polymerase. Sci. Adv. 11, eado6280 (2025).

16 S. Komatsu, H. Ohno & H. Saito. Target-dependent RNA polymerase as universal platform for gene expression control in response to intracellular molecules. Nat. Commun. 14, 7256 (2023).

17 E. T. Butler & M. J. Chamberlin. Bacteriophage SP6-specific RNA polymerase. I. Isolation and characterization of the enzyme. J. Biol. Chem. 257, 5772–5778 (1982).

18 N. C. Shaner, G. G. Lambert, A. Chammas, Y. Ni, P. J. Cranfill, M. A. Baird, B. R. Sell, J. R. Allen, R. N. Day, M. Israelsson, M. W. Davidson & J. Wang. A bright monomeric green fluorescent protein derived from Branchiostoma lanceolatum. Nat. Methods 10, 407–409 (2013).

19 Y. Wang, D. E. Prosen, L. Mei, J. C. Sullivan, M. Finney & P. B. Vander Horn. A novel strategy to engineer DNA polymerases for enhanced processivity and improved performance in vitro. Nucleic Acids Res. 32, 1197–1207 (2004).

20 M. Mirdita, K. Schutze, Y. Moriwaki, L. Heo, S. Ovchinnikov & M. Steinegger. ColabFold: making protein folding accessible to all. Nat. Methods 19, 679–682 (2022).

21 T. H. Tahirov, D. Temiakov, M. Anikin, V. Patlan, W. T. McAllister, D. G. Vassylyev & S. Yokoyama. Structure of a T7 RNA polymerase elongation complex at 2.9 A resolution. Nature 420, 43–50 (2002).

22 A. Dousis, K. Ravichandran, E. M. Hobert, M. J. Moore & A. E. Rabideau. An engineered T7 RNA polymerase that produces mRNA free of immunostimulatory byproducts. Nat. Biotechnol. 41, 560–568 (2023).

23 Y. Kazuta, T. Matsuura, N. Ichihashi & T. Yomo. Synthesis of milligram quantities of proteins using a reconstituted in vitro protein synthesis system. J. Biosci. Bioeng. 118, 554–557 (2014).

24 P. A. Krieg & D. A. Melton. In vitro RNA synthesis with SP6 RNA polymerase. Methods Enzymol. 155, 397–415 (1987).

25 K. Takai, T. Sawasaki & Y. Endo. Practical cell-free protein synthesis system using purified wheat embryos. Nat. Protoc. 5, 227–238 (2010).

26 T. H. Segall-Shapiro, A. J. Meyer, A. D. Ellington, E. D. Sontag & C. A. Voigt. A ‘resource allocator’ for transcription based on a highly fragmented T7 RNA polymerase. Mol. Syst. Biol. 10, 742 (2014).

27 J. W. Ellefson, A. J. Meyer, R. A. Hughes, J. R. Cannon, J. S. Brodbelt & A. D. Ellington. Directed evolution of genetic parts and circuits by compartmentalized partnered replication. Nat. Biotechnol. 32, 97–101 (2014).

28 J. Pu, J. A. Dewey, A. Hadji, J. L. LaBelle & B. C. Dickinson. RNA Polymerase Tags To Monitor Multidimensional Protein-Protein Interactions Reveal Pharmacological Engagement of Bcl-2 Proteins. J. Am. Chem. Soc. 139, 11964–11972 (2017).

29 M. J. Styles, J. A. Pixley, T. Wei, C. Basile, S. S. Lu & B. C. Dickinson. PANCS-Binders: a rapid, high-throughput binder discovery platform. Nat. Methods 22, 1720–1730 (2025).

30 A. Zorba, V. Nguyen, A. Koide, M. Hoemberger, Y. Zheng, S. Kutter, C. Kim, S. Koide & D. Kern. Allosteric modulation of a human protein kinase with monobodies. Proc. Natl. Acad. Sci. USA 116, 13937–13942 (2019).

31 H. Ueda, K. Tsumoto, K. Kubota, E. Suzuki, T. Nagamune, H. Nishimura, P. A. Schueler, G. Winter, I. Kumagai & W. C. Mahoney. Open sandwich ELISA: A novel immunoassay based on the interchain interaction of antibody variable region. Nat. Biotechnol. 14, 1714–1718 (1996).

32 T. W. J. Gadella, Jr., L. van Weeren, J. Stouthamer, M. A. Hink, A. H. G. Wolters, B. N. G. Giepmans, S. Aumonier, J. Dupuy & A. Royant. mScarlet3: a brilliant and fast-maturing red fluorescent protein. Nat. Methods 20, 541–545 (2023).

33 S. A. Fernando & G. S. Wilson. Studies of the Hook Effect in the One-Step Sandwich Immunoassay. J. Immunol. Methods 151, 47–66 (1992).

34 B. Wang, S. Cao & N. Zheng. Emerging strategies for prospective discovery of molecular glue degraders. Curr. Opin. Struct. Biol. 86, 102811 (2024).

35 M. S. Gadd, A. Testa, X. Lucas, K. H. Chan, W. Chen, D. J. Lamont, M. Zengerle & A. Ciulli. Structural basis of PROTAC cooperative recognition for selective protein degradation. Nat. Chem. Biol. 13, 514–521 (2017).

36 K. M. Riching, E. A. Caine, M. Urh & D. L. Daniels. The importance of cellular degradation kinetics for understanding mechanisms in targeted protein degradation. Chem. Soc. Rev. 51, 6210–6221 (2022).

37 A. Hecht, D. Endy, M. Salit & M. S. Munson. When Wavelengths Collide: Bias in Cell Abundance Measurements Due to Expressed Fluorescent Proteins. ACS Synth. Biol. 5, 1024–1027 (2016).

38 C. G. England, E. B. Ehlerding & W. Cai. NanoLuc: A Small Luciferase Is Brightening Up the Field of Bioluminescence. Bioconjug. Chem. 27, 1175–1187 (2016).

39 Y. M. Long, A. Mora, F. Z. Li, E. Gürsoy, K. E. Johnston & F. H. Arnold. LevSeq: Rapid Generation of Sequence-Function Data for Directed Evolution and Machine Learning. ACS Synth. Biol. 14, 230–238 (2024).

40 W. Lu, N. Terasaka, Y. Sakaguchi, T. Suzuki, T. Suzuki & H. Suga. An anticodon-sensing T-boxzyme generates the elongator nonproteinogenic aminoacyl-tRNA in situ of a custom-made translation system for incorporation. Nucleic Acids Res. 52, 3938–3949 (2024).

41 D. A. Wong, Z. M. Shaver, M. D. Cabezas, M. Daniel-Ivad, K. F. Warfel, D. V. Prasanna, S. E. Sobol, R. Fernandez, F. Tobias, S. K. Filip, S. W. Hulbert, P. Faull, R. Nicol, M. P. DeLisa, E. P. Balskus, A. S. Karim & M. C. Jewett. Characterizing and engineering post-translational modifications with high-throughput cell-free expression. Nat. Commun. 16, 7215 (2025).

42 C. K. Schissel, H. Roberts-Mataric, I. J. Garcia, H. Kang, R. Mowzoon-Mogharrabi, M. B. Francis & A. Schepartz. Peptide Backbone Editing via Post-Translational O to C Acyl Shift. J. Am. Chem. Soc. 147, 6503–6513 (2025).

43 A. Sanjeev, C. R. Summers, J. Politi, P. J. Beuning & C. S. P. Cheng. Engineered chimeric T7 RNA polymerase improves salt tolerance and reduces dsRNA impurity generation during in vitro transcription of mRNA. Nucleic Acids Res. 53 (2025).

44 V. Noireaux, R. Bar-Ziv & A. Libchaber. Principles of cell-free genetic circuit assembly. Proc. Natl. Acad. Sci. USA 100, 12672–12677 (2003).

45 J. Bae, J. Kim, J. Choi, H. Lee & M. Koh. Split Proteins and Reassembly Modules for Biological Applications. ChemBioChem 25, e202400123 (2024).

46 Z. Abil, A. M. R. Sierra & C. Danelon. Clonal Amplification-Enhanced Gene Expression in Synthetic Vesicles. ACS Synth. Biol. 12, 1187–1203 (2023).

47 Y. Zhang, Y. Minagawa, H. Kizoe, K. Miyazaki, R. Iino, H. Ueno, K. V. Tabata, Y. Shimane & H. Noji. Accurate high-throughput screening based on digital protein synthesis in a massively parallel femtoliter droplet array. Sci. Adv. 5, eaav8185 (2019).

48 Y. Azuma, T. G. W. Edwardson, N. Terasaka & D. Hilvert. Modular Protein Cages for Size-Selective RNA Packaging in Vivo. J. Am. Chem. Soc. 140, 566–569 (2018).

49 G. S. Filonov, C. W. Kam, W. Song & S. R. Jaffrey. In-gel imaging of RNA processing using broccoli reveals optimal aptamer expression strategies. Chem. Biol. 22, 649–660 (2015).

50 H. M. Zhao, L. Giver, Z. X. Shao, J. A. Affholter & F. H. Arnold. Molecular evolution by staggered extension process (StEP) in vitro recombination. Nat. Biotechnol. 16, 258–261 (1998).

